# euka: Robust detection of eukaryotic taxa from modern and ancient environmental DNA using pangenomic reference graphs

**DOI:** 10.1101/2023.04.04.535531

**Authors:** Nicola Alexandra Vogel, Joshua Daniel Rubin, Mikkel Swartz, Juliette Vlieghe, Peter Wad Sackett, Anders Gorm Pedersen, Mikkel Winther Pedersen, Gabriel Renaud

## Abstract

1. Ancient environmental DNA (eDNA) is a crucial source of in-formation for past environmental reconstruction. However, the com-putational analysis of ancient eDNA involves not only the inherited challenges of ancient DNA (aDNA) but also the typical difficulties of eDNA samples, such as taxonomic identification and abundance esti-mation of identified taxonomic groups. Current methods for ancient eDNA fall into those that only perform mapping followed by taxo-nomic identification and those that purport to do abundance estima-tion. The former leaves abundance estimates to users, while methods for the latter are not designed for large metagenomic datasets and are often imprecise and challenging to use.

2. Here, we introduce euka, a tool designed for rapid and accurate characterisation of ancient eDNA samples. We use a taxonomy-based pangenome graph of reference genomes for robustly assigning DNA sequences and use a maximum-likelihood framework for abundance estimation. At the present time, our database is restricted to mito-chondrial genomes of tetrapods and arthropods but can be expanded in future versions.

3. We find euka to outperform current taxonomic profiling tools as well as their abundance estimates. Crucially, we show that regardless of the filtering threshold set by existing methods, euka demonstrates higher accuracy. Furthermore, our approach is robust to sparse data, which is idiosyncratic of ancient eDNA, detecting a taxon with an average of fifty reads aligning. We also show that euka is consistent with competing tools on empirical samples and about ten times faster than current quantification tools.

4. euka’s features are fine-tuned to deal with the challenges of ancient eDNA, making it a simple-to-use, all-in-one tool. It is available on GitHub: https://github.com/grenaud/vgan. euka enables re-searchers to quickly assess and characterise their sample, thus allowing it to be used as a routine screening tool for ancient eDNA.

## 1 Introduction

Over the last forty years, investigation of ancient DNA (aDNA) has allowed researchers to study and understand extinct species and their past environments. However, the idiosyncrasies of aDNA render its proper analysis exceptionally challenging. Firstly, DNA degrades in the environment after the cell leaves the host as DNA repair mechanisms cease to work. This results in DNA fragmentation, where ancient DNA typically tends to be particularly short (*<* 100bp) (Pääbo 1989; Hofreiter, Serre, et al. 2001). Second, DNA fragments accumulate post-mortem chemical damages (such as deamination of cytosines causes observed damage patterns of cytosine to thymine or guanine to adenine) (Briggs et al. 2007; Prüfer et al. 2010). Ancient DNA damage and short fragment lengths can cause problems aligning the sequences to reference genomes and taxonomic profile aDNA samples (Poullet and Orlando 2020; Jónsson et al. 2013). Finally, contamination from either ancient microbes or present-day sources often represents the majority of the extracted DNA. Such contamination can align to the reference genome spuriously or due to the phylogenetic proximity of the contaminant (Peyŕegne and Peter 2020). They are often seen to be piling up in specific genomic locations instead of evening distributed across the reference genome. While the field of aDNA mostly has focussed on the isolation of DNA from a fossil source, more recently, ancient and modern environmental DNA (eDNA) has become an increasingly used method to track, monitor and reconstruct organismal assemblages. However, the greatest computational challenge of both ancient and modern eDNA is to correctly assign every read taxonomically and, following estimate, the corresponding abundance of the taxa (Ruppert, Kline, and Rahman 2019; Beng and Corlett 2020).

Following the decomposition of biological remains, such as animals and plants, traces of extracellular DNA remain preserved in sediments (Haile et al. 2007; Yoccoz et al. 2012; Haouchar et al. 2014; Willerslev, Hansen, et al. 2003; Hofreiter, Mead, et al. 2003; Lydolph et al. 2005; Willerslev and Cooper 2005). Ancient eDNA fragments from eukaryotic species combined with other characteristics of sediments provided a novel way to study past ecosystems, the timing of speciation and extinction events, population structure and migration dynamics (Jørgensen et al. 2012; Graham et al. 2016; Ficetola et al. 2018; Dussex et al. 2021; Pansu et al. 2015; Zavala et al. 2021; Mikkel W Pedersen et al. 2016). In the analysis of ancient eDNA, mitochondrial DNA (mtDNA) is frequently used for species identification (Zavala et al. 2021; Murchie, Monteath, et al. 2021; Slon et al. 2017), as the high abundance and smaller genome of the mitochondrion compared to the nucleus allows for more accessible and faster species characterisations (Elyasigorji et al. 2022; Crampton-Platt et al. 2016; Gómez-Rodrıguez et al. 2015).

The challenges associated with ancient eDNA are the characteristic problems of aDNA combined with the computational difficulties of eDNA. There have been many approaches to tackling both sets of problems. The use of taxonomy-specific primers, otherwise referred to as metabarcoding, facilitates species identification in eDNA samples. It has also been used for ancient eDNA analysis (Giguet-Covex et al. 2014; Crump et al. 2021; Rijal et al. 2020; Weyrich et al. 2017). However, metabarcoding is biased towards longer, more modern DNA fragments (Beng and Corlett 2020; Orlando et al. 2021). A more effective alternative for sequencing ancient eDNA is shotgun-sequencing metagenomes (Slon et al. 2017; Mikkel Winther Pedersen et al. 2021; Lammers, Heintzman, and Alsos 2021; Gelabert et al. 2021). It am-plifies all DNA fragments, independent of their size (Armbrecht, Eisenhofer, et al. 2021; Taberlet et al. 2012). Capture enrichment via bait/probe sets designed to retain specific DNA sequences indicative of the presence of certain species is often used in combination with shotgun-metagenomics (Slon et al. 2017; Armbrecht, Herrando-Pérez, et al. 2020; Murchie, Kuch, et al. 2021; Kjær et al. 2022). However, to efficiently use capture enrichment, the target species must be known *a priori*. This leaves the task of taxonomically char-acterising present species and estimating their abundance (e.g. 70% brown bears, 20% beetles, etc.) of the aDNA in a metagenomic sample.

There is a pressing need for more accurate taxonomic identification and abundance estimation of ancient eDNA data. Some current methods focus on mapping and taxonomic classification, leaving post-processing and quan-tification estimations to the user. The most common of these include MALT (Megan Alignment Tool) (Herbig et al. 2017) and Bowtie2 (Langmead and Salzberg 2012) combined with ngsLCA (Wang et al. 2022) which is part of the HOLI pipeline (Mikkel W Pedersen et al. 2016). Both methods align against a reference database and subsequently run the naive lowest common ances-tor (LCA) algorithm (Bender et al. 2005) to taxonomically classify every alignment in the input file.

It is to our knowledge only HAYSTAC that performs mapping and reports abundance estimation for ancient eDNA (Dimopoulos et al. 2022). It presents an alternative to LCA inference and provides post-processing steps to ascer-tain the authenticity of the ancient eDNA (e.g. ancient damage estimation, coverage evenness test). However, HAYSTAC is computationally intensive, of-ten taking several days for a single sample. It is, therefore, impractical to rapidly screen large datasets. All current methods, HAYSTAC included, require the user to build a reference set of expected taxa, potentially overlooking certain taxa if the database is too narrow.

We introduce euka, an easy-to-use command-line tool written in C++ for robust identification of tetrapodic and arthropodic mitochondrial DNA from ancient eDNA samples. euka aligns ancient eDNA sequences to a taxon-based pangenomic reference graph, filters using an ancient-damage-aware maximum-likelihood framework and subsequently estimates the respective abundance using Markov Chain Monte Carlo (MCMC). Crucially, we show that regardless of the filtering approach, euka outperforms existing tools de-tecting taxa in simulated modern and ancient metagenomic environments using euka’s curated database and taxonomic resolution. We demonstrate euka’s accurate abundance estimation for a simulated ancient eDNA sample and its robustness to low-abundance taxa detection with approximately fifty reads aligning. We also find euka to perform consistently with other tools on empirical datasets. euka is roughly 10 times faster than current ancient eDNA abundance estimation tools and is able to process a billion reads in approximately six hours, thus making it suitable for the screening of ancient eDNA samples. euka is a subcommand of the vgan suite of tools for pangenomics, an open-source software under GPLv3. It accepts FASTQ as input and provides the user with a summary of all taxonomic groups present within the given sample, including their respective abundance esti-mates, damage profiles, fragment length distribution and estimated coverage across the pangenome graph.

## 2 Methods

### 2.1 General Workflow

In this section, we introduce euka’s general workflow (Figure 1), which com-prises three steps: 1) mapping to our reference structure, 2) filtering spurious alignments and 3) abundance estimation of detected taxa. First, we describe our curated database and how it is used to construct the taxa-based pangenome reference structure. Then we explain the quality-control filters, including our ancient damage-aware maximum-likelihood framework for quantifying the probability of correct alignments and the test for minimal coverage across our pangenome graph. Lastly, we describe our approach for estimating the abundance of detected taxonomic groups using an MCMC sampling algorithm.

**Figure 1:**
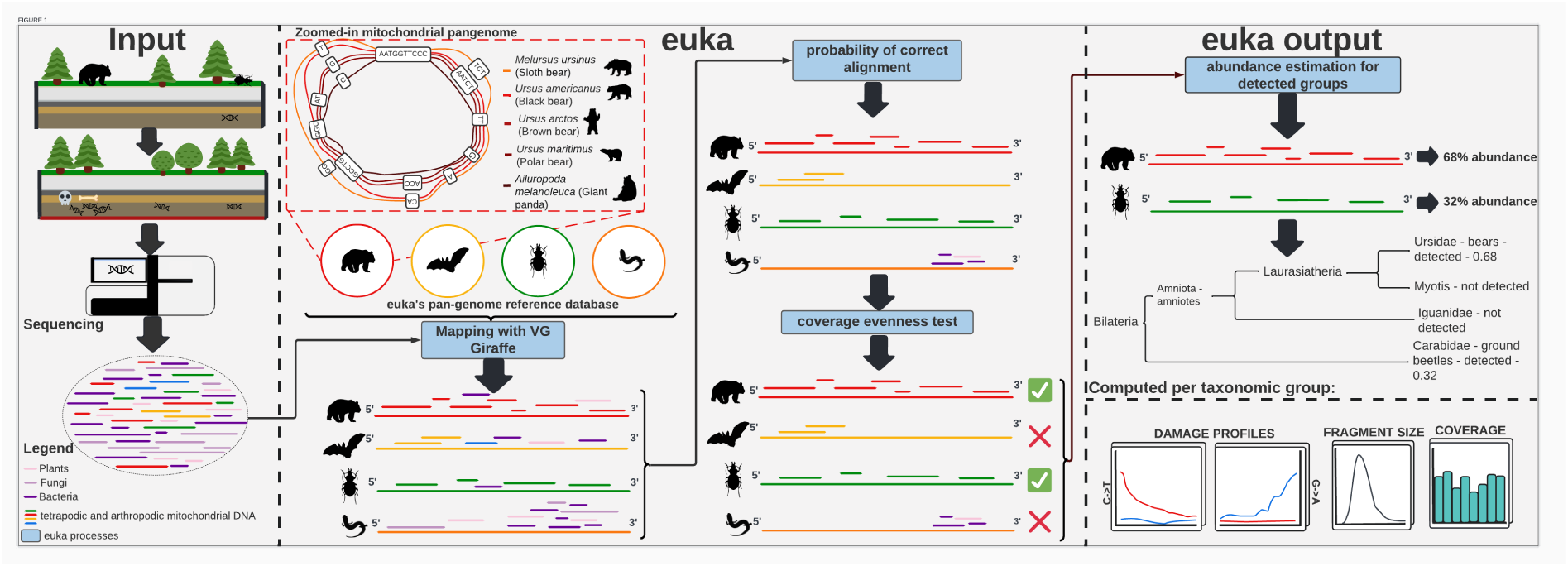
euka’s main workflow: Sediments contain extracellular DNA through the decomposition of biological remains. Sequencing techniques pick up on the ancient DNA fragments and every other source of DNA in sedimentary samples. The resulting data contains a mix of unknown DNA sequences (FASTQ), used as input for euka. euka maps the unknown samples against the taxa-based mitochondrial pangenome graph to determine which taxa are present in a metagenomic sample. To refine its findings euka uses a maximum likelihood framework, which accounts for the effects of damage typically seen in aDNA, to filter spurious alignments. Additionally, euka filters taxa based on a minimal breadth of coverage across the pangenome. The abundance of the detected taxa is estimated, and the output is visualised to provide a detailed summary of the sample.

### 2.1 Database Content

euka’s first workflow step involves mapping metagenomic sequence data to our pangenomic graph reference structure (find an explanation in Supplementary section 1), which is provided via FTP as described in euka’s installation guide^1^. Our reference structure is built from a curated database comprising 5847 mitochondrial genomes from 336 defined taxonomic groups within the phylum Arthropoda and subphylum Tetrapoda of the kingdom Animalia (see Supplementary Table 2 and Supplementary section 2 for a detailed description of the construction, curation and an explanation of the taxonomic resolution). The mitogenomes are clustered into taxa according to their sequence identity, and each taxon is transformed into a single circular graph (see Figure 1 for an example using bears as a taxon). All the connected components are merged together into a single graph to form euka’s pangenomic reference graph used for mapping. For further details on how the pangenomic graph is built from the mitochondrial database, please refer to section 3 of the Supplementary material.

### 2.3 Mapping and Post-processing

euka loads its pangenomic reference graph at runtime, together with the user-provided FASTQ file for mapping. We assume that the FASTQ input has been pre-processed (low-complexity filtering and PCR duplicate removal) prior to running euka. euka calls the subcommand giraffe of the vg toolkit (Siŕen et al. 2021; Hickey et al. 2020) to map the input FASTQ against the mitogenomes embedded in our graph (see Supplementary section 4 for mapping parameters). In the first post-processing phase, we seek to determine if an alignment is genuine, i.e. that it stems from the correct taxa and that the position of the aligned DNA fragment is correct. To achieve this, we compute the ratio of two models. The first posits that the alignment is correct, and the second postulates that the alignment is spurious. Using simulations, we determined that this ratio effectively minimises the number of incorrect alignments (Supplementary Figure 1). Additionally, we discard alignments with a mapping quality threshold under 30 on a PHRED scale calculated by vg giraffe (Supplementary Figure 2). A detailed explanation of our framework is provided in the Supplementary section 5. In the second post-processing phase, we restrict the analysis to taxa where sequenced DNA fragments sufficiently covered the mitogenomes in the graph and were evenly distributed. With this step, we aim to avoid the detection of taxa where multiple DNA sequences align to only one part of the genome, especially in regions of low complexity, otherwise referred to as read pile-ups. To mitigate the impact of such pile-ups, a coverage binning system was developed to estimate the minimal breadth of coverage of each taxon’s connected component. A taxon is only considered detected if it has a minimum of one read mapped to each bin of the taxon pangenome graph, has a minimum of six bins with an entropy score of 1.17 and, in total, a minimum of ten mapped reads. We deliver a more detailed explanation in the Supplementary section 6.

### 2.4 Abundance Estimation

We now have completed the alignment post-processing; every remaining taxon and associated alignment is believed to be of high confidence. The next step is the abundance estimation of detected taxa. To do this, we first define an initial abundance vector for the sample as the proportion of post-processed alignments in each detected taxon compared to the total number of high-confidence alignments. We use an MCMC sampling algorithm to obtain a posterior estimate of our abundance vector as well as confidence intervals (95% and 85%). We describe our abundance estimation in more detail in the Supplementary section 7.

### 2.5 Ancient DNA authentication

For every taxon that passes our filtering steps, we check alignments to the taxon for signs of ancient damage. If there is an ancient damage event (*C → U* mutations seen as *C → T* or *G → A* mutations on the reverse strand), we store these occurrences and output the probability of observing ancient damage in a given position as a damage profile for the taxon. Along-side the damage profiles, we also output a tab-separated file detailing the coverage and fragment length distribution, completing the post-processing of the alignment file. We provide R scripts to plot damage rates at each end of the DNA sequence, fragment size distribution as well as the coverage of the mitogenomes on a per-bin basis.

### 2.6 Test Data Preparation

To test the accuracy of all tools when it comes to taxa detection as well as individual taxonomic read assignments, we generated three *in silico* ancient eDNA sample environments. These simulations are designed to mimic the abundance of different kingdoms in an empirical environment from previously published studies as closely as possible. This is done to realistically simulate the difficulties that environmental samples represent, such as high false-positive mapping from non-eukaryotic species to eukaryotic genomes. A description of the simulated samples and our simulation workflow can be found in the Supplementary section 8.

We used three empirical samples for benchmarking against existing meth-ods. Two samples were already pre-processed (filtered for low-complexity reads and PCR duplicates removed), a 15kya Mexican cave sediment sample (Mikkel Winther Pedersen et al. 2021; Ardelean et al. 2020) and a 2-million-year-old ancient eDNA sample from Greenland (Kjær et al. 2022). Further-more, we had a 30kya permafrost sample from Yukon, Canada (Murchie, Monteath, et al. 2021), where we used the raw empirical data, did adapter trimming and merging with leeHom (Renaud, Stenzel, and Kelso 2014) using ancient parameters and filtered low-complexity reads using sga preprocess-ing^2^.

### 2.7 Software Limitations

In euka’s current version, the database is restricted to tetrapodic and arthropodic taxa to reduce disk space and mapping time requirements. However, as compression techniques for pangenome graphs continue to improve (De-orowicz, Danek, and Li 2023), we plan to update our database to cover all eukaryotes in the near future. An additional limitation of euka is that we only process DNA from the mitochondria. The size of nuclear genomes would make the pangenome construction across many species intractable.

## 3 Results

First, we demonstrate euka’s performance compared to existing tools using three simulated ancient eDNA environments, where present taxa and exact abundances are known. We benchmark against Bowtie2 + ngsLCA and MALT by mapping and analysing the different taxa without explicit abundance quantification. We subsequently compare euka’s performance against HAYSTAC, which specifically quantifies abundance. We provide a feature comparison in Supplementary Table 1. Finally, we present euka’s results compared to all four tools on three empirical samples. All experiments used euka’s curated database as a reference to evaluate the computational method *per se*. Owing to our taxonomic limitations, we only considered alignments to taxa defined in Supplementary Table 2. The exact commands are provided in the Supplementary section 10.

### 3.1 Simulated Data

#### 3.1.1 Mapping Accuracy

In our first benchmark, we evaluated the accuracy of read assignments for four different tools, namely euka, MALT, Bowtie2+ngsLCA, and HAYSTAC, using simulated environments. euka only considers alignments with high confidence as defined in section 2.4. For the remaining tools, we set the filtersas recommended by the authors in the original publication (Supplementary section 10). In our simulations, we know the ground truth assignments and can therefore compute the accuracy of assignments to taxa. Accuracy is computed as the number of correctly assigned reads divided by the total number of alignments to these taxa. Figure 2A indicates that our approach produced the highest mean accuracy across all environments and damage rates, demonstrating its effectiveness (for the accuracy measure from 1 to 0, see Supplementary figure 3). Figure 2B shows the rates of simulated damage. In comparison, Bowtie2+ngsLCA’s strict filters followed by HAYSTAC’s Bayesian approach resulted in lower accuracy. MALT, with its heuristic alignment ap-proach, produced the lowest accuracy.

**Figure 2:**
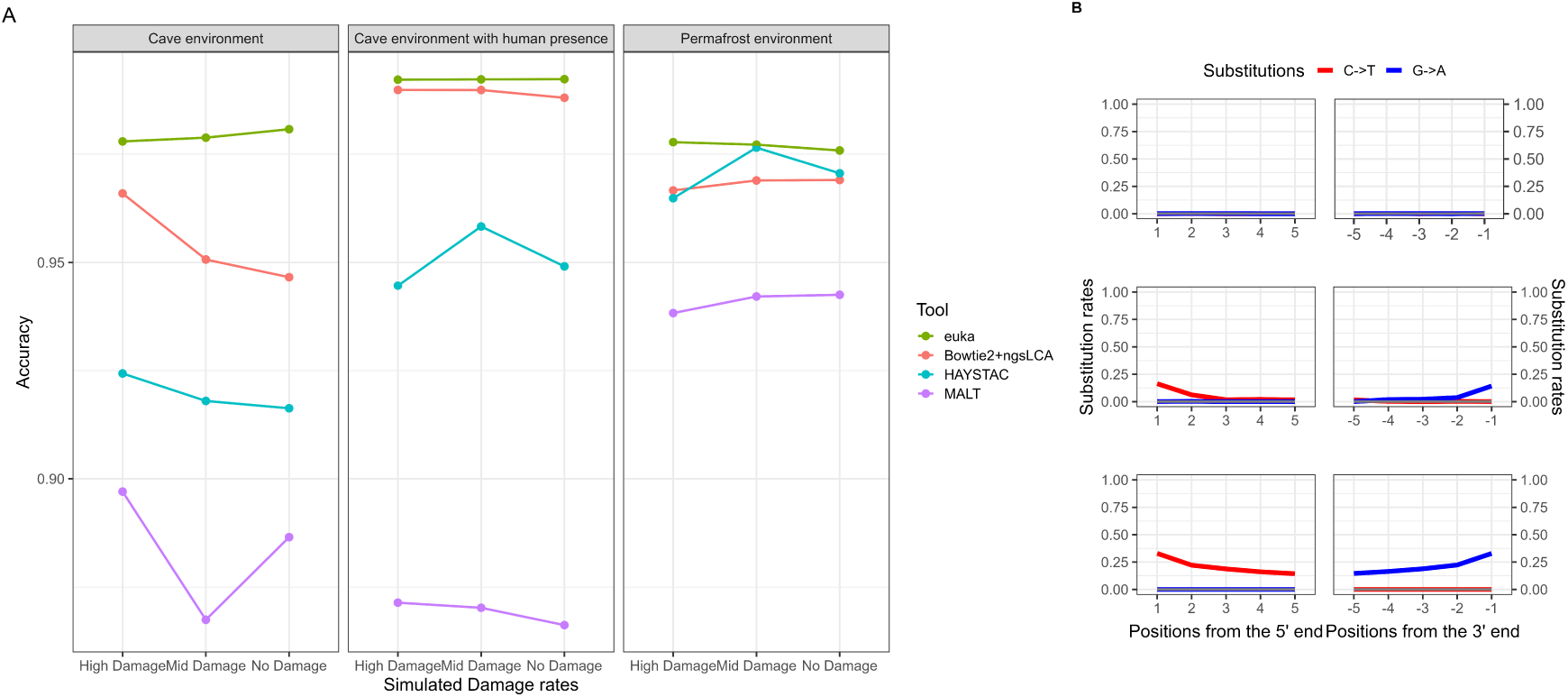
A Comparison of sensitivity on simulated reads. True positive mappings to all known taxa were assessed within three simulated ancient metagenomic samples on three levels of simulated damage. Each tool was provided with the same database for mapping and compared on euka’s defined taxonomic levels. B Damage profiles for the three levels of simulated damage within the metagenomic samples. The top panel shows no simulated damage; the middle panel shows the profile for our mid-simulated damage samples and the bottom panel for our high-damage simulations.

**Figure 3:**
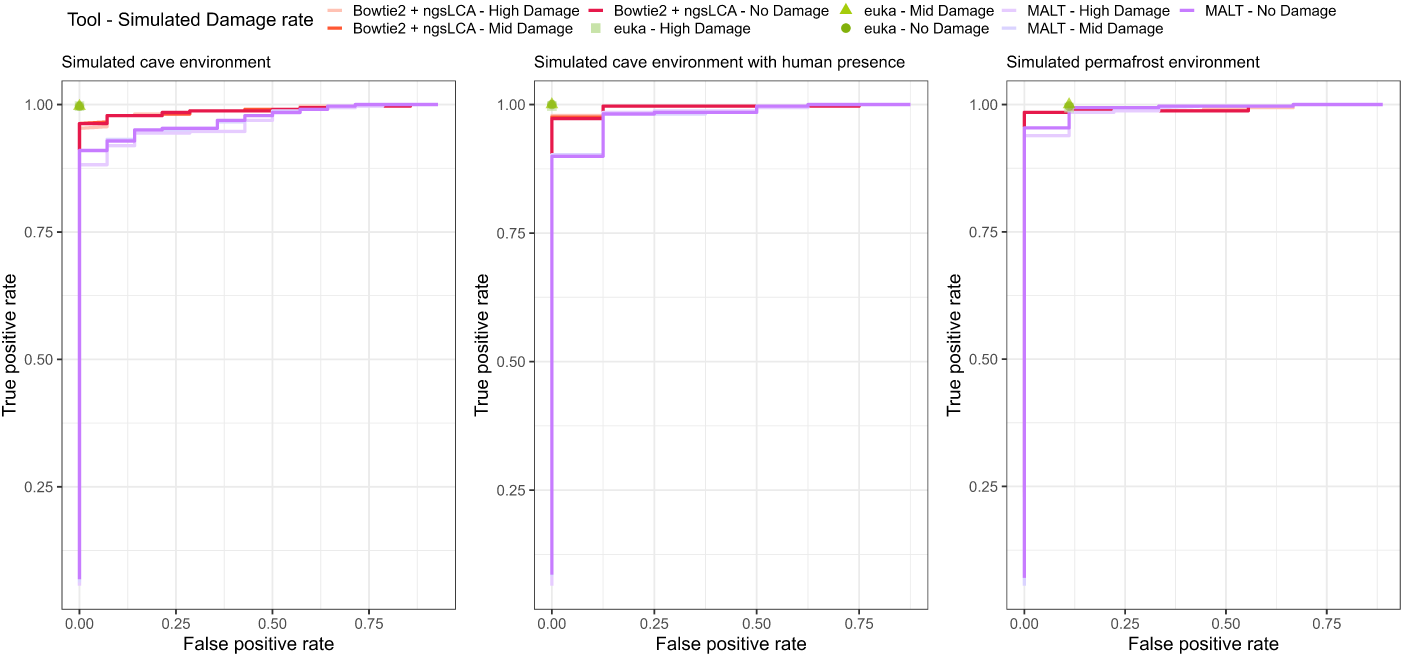
Comparing euka against Bowtie2 + ngsLCA and MALT on taxa abundance estimation. Because Bowtie2 + ngsLCA and MALT do not provide abundance estimations for their detected taxa, the taxa detection comparison between Bowtie2 + ngsLCA and MALT and euka is computed by setting each number between 1 and the maximum number of reads assigned to one taxon as a threshold and comparing the true positive rate against the false positive rate. A list of detected taxa and their abundance estimate is provided by euka; it, therefore, allows us to show it a single dot.

#### 3.1.2 Correctness of Abundance

We next evaluated the estimated taxa abundances for each simulated environ-ment. Both Bowtie2+ngsLCA and MALT are predicated on the LCA algorithm and output every taxon with at least one alignment and do not provide abundance estimates. Therefore, to compare euka with Bowtie2+ngsLCA and MALT, we used as a criterion for detecting a taxon a threshold ranging from a single alignment to the maximum number of alignments to a given taxon. We assessed false-positives against true-positives (Figure 3). Especially for the cave environments, euka significantly outperforms both competing tools. For the permafrost environment, euka is more accurate than existing meth-ods; however, this environment holds the challenge of detecting the ant taxon (Formicidae). With standard parameters, we cannot detect the Formicidae as their mitogenomes consist of too many low-entropy regions.

We then compared HAYSTAC and euka abundance estimation precision using Bray-Curtis dissimilarity (Bray and Curtis 1957; Ricotta and Podani 2017). Figure 4 displays mean dissimilarity between euka and HAYSTAC compared to the ground truth, showing that euka consistently outperforms HAYSTAC. Importantly, HAYSTAC overestimates abundance by assigning a default value to all present taxa, regardless of filtering criteria, leading to significant bias in larger reference sets. In contrast, euka’s taxon filtering method ensures more accurate abundance estimation. We excluded the ant family (Formicidae) from the permafrost sample abundance calculation due to their low-entropy genomes and high likelihood of spurious alignments.

**Figure 4:**
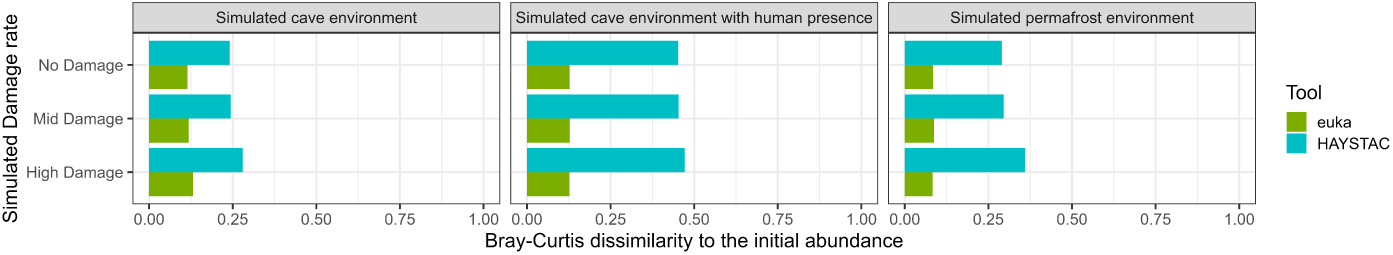
euka against HAYSTAC comparison of the average Bray-Curtis dissimilarity of the output abundance vectors. The Bray-Curtis dissimilarity estimates values from 0 to 1, with 0 showing perfect similarity. The Bray-Curtis dissimilarity between euka’s and HAYSTAC’s abundance vectors to the initial input vector of the simulated metagenomic sample was calculated. This figure emphasises euka’s robustness to ancient damage compared to HAYSTAC.

Furthermore, we compared runtimes of the quantification tools, namely euka and HAYSTAC for our simulated environments. Figure 5 demonstrates that euka is significantly faster ( 8.3*x* faster on user time and 12*x* faster on wall clock time). euka requires about 46Gb of RAM, which is feasible even for modern laptops.

**Figure 5:**
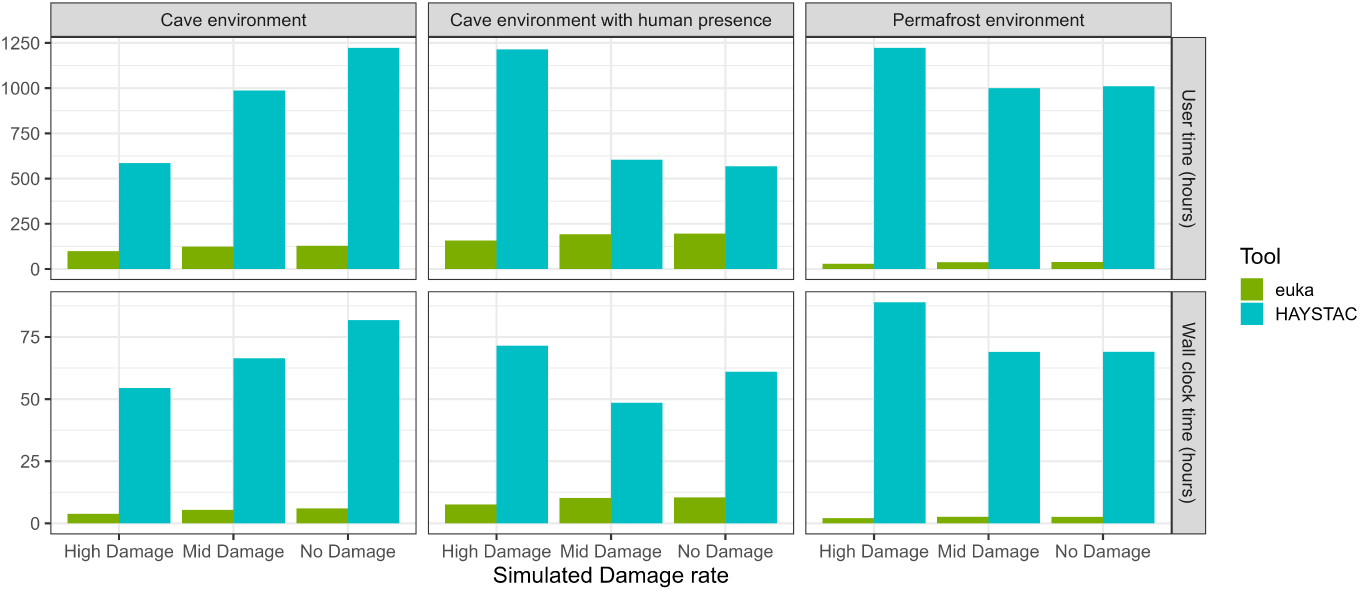
User time in hours (top) and wall clock time in hours (bottom) comparison between euka and HAYSTAC on three simulated ancient metagenomic samples under three levels of simulated damage rates. On average, euka is about 8.3x faster in user time than HAYSTAC. On a metagenomic sample with approximately 1 billion fragments, euka has an average runtime of 5.66h compared to HAYSTAC’s average runtime of 67,85h (multi-threaded at 20 cores), making euka about 12x faster.

#### 3.1.3 Robustness of Abundance

To evaluate euka’s robustness to low coverage depth, we generated a FASTQ file containing three taxa (Bovidae, Myotis, Ursidae) in low, medium and high abundance, respectively. We downsampled reads to 75%, 50%, 25%, 10% and 1% of the original sample for all three simulated ancient DNA damage ranges (Figure 2B)). Our results (Figure 6) demonstrate that euka is highly accurate and can estimate abundances down to 10% of the original input for high-and medium-abundant taxa. Even in low-abundance scenarios, euka remains accurate down to 75% of the original input level, equivalent to approximately fifty alignments ( 0.2*X* coverage). In contrast, HAYSTAC’s abundance estimation fails to recover the ground truth, owing to the same issue shown in the environment simulation experiment. We repeated the experiment for medium damage and no damage, finding no discernible differences to Figure 6 (see Supplementary Figures 4 and 5, respectively).

**Figure 6:**
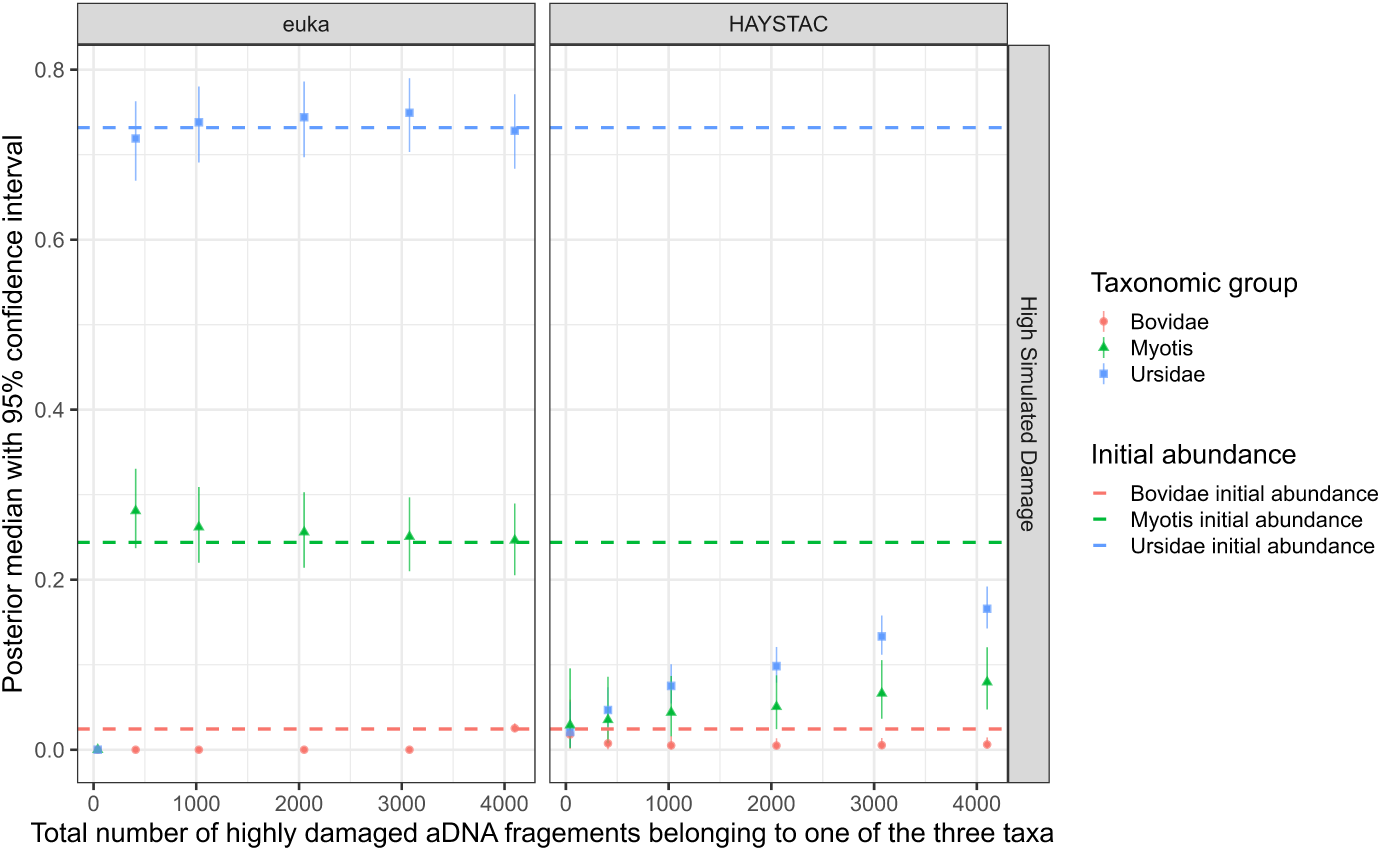
euka and HAYSTAC’s abundance estimation for a downsampled ancient sample of three taxa. To show euka’s robustness, we downsampled the original sample, considering where euka would be unable to detect the different taxa. The initial abundance of the three taxa was represented with a dotted line. On average, euka detects taxa with approx. 50 alignments. The robustness test was repeated for a medium simulated damage rate (Supplementary Figure4 and with no ancient damage (Supplementary Figure 5); however, the results for euka and HAYSTAC were consistent with different levels of damage.

### 3.2 Empirical Data

We also benchmarked on empirical samples. Here the ground truth is unknown, so we evaluated consistency while manually verifying the results. We used two empirical ancient eDNA samples to compare against existing methods and one ancient eDNA sample to demonstrate euka’s detailed output summary.

#### 3.2.1 Analysis of a 15kya Mexican Cave Sample

First, we used the UE1210 Mexican cave layer sample from Mikkel Winther Pedersen et al. (2021) (ENA projects PRJEB42692). The sample was sequenced using Illumina HiSeq and NovaSeq shotgun sequencing and reported the presence of the American Black Bear (Mikkel Winther Pedersen et al. 2021). In the previous study, which describes the cave environment Ardelean et al. (2020), the tribe Marmotini was also detected. To set a threshold on the number of aligned reads to mark a taxon as being detected for Bowtie2+ngsLCA and MALT, we used the threshold obtained from our ROC curves, which has the highest true positive value (all known taxa are detected), with the lowest false-positive value, as described in section 3.1.2. As for HAYSTAC, we used every taxon with coverage evenness value less than ten as recommended in Dimopoulos et al. (2022). Since HAYSTAC classifies at the species level, we aggregated reads from the same taxon as defined in euka’s database (Supplementary Table 2). We considered a taxon detected for HAYSTAC if thirty reads were mapped.

Figure 7 illustrates euka’s ability to detect both the highly-abundant bears and the low-abundance taxon Xerinae (the tribe Marmotini is contained within the subfamily Xerinae in euka’s database). All three competing tools report a number of taxa unseen in the original publication. For example, we see a recurring taxon of skinks (Scincidae) (displayed in red in Figure 7) predicted by Bowtie2 + ngsLCA, HAYSTAC, and MALT. However, euka does not predict the presence of this taxon. Further inspection of the alignments shows DNA pileups over a single locus in the skink mitogenome (Supplementary Figures 6, 7, 8). The alignments are, therefore, almost certainly spurious.

**Figure 7:**
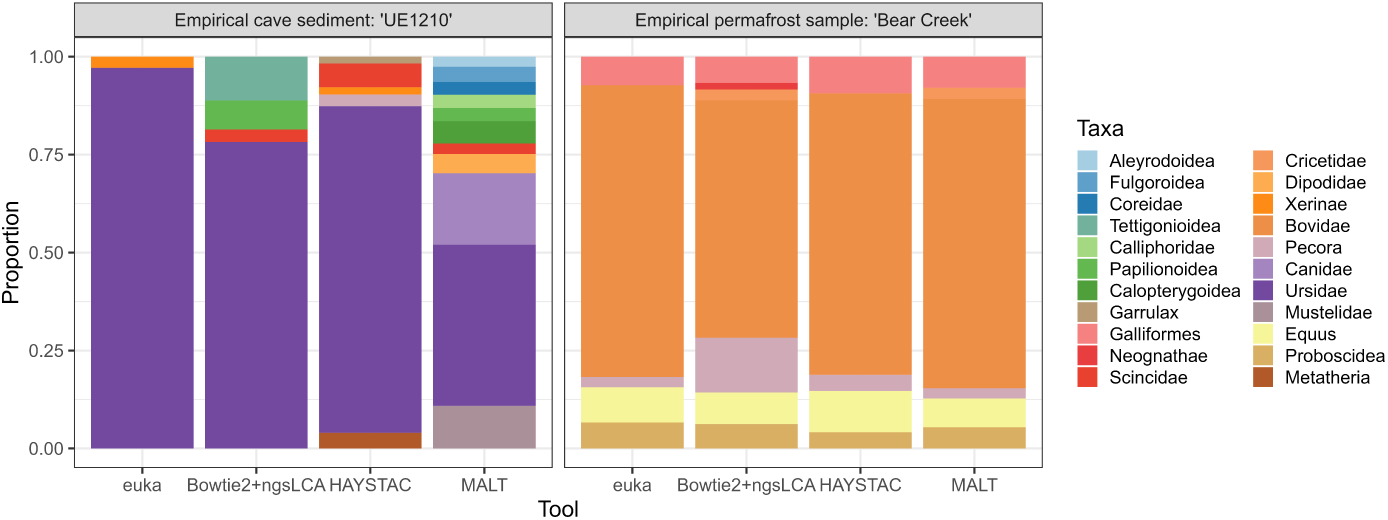
Taxa identification for two empirical datasets. Considering only taxa in HAYSTAC with coverage evenness *<* 10. We used the best threshold estimated for the mimicked simulated sample from Figure 3 for Bowtie2 + ngsLCA and MALT, excluding every taxonomic rank above superfamily for the ’UE1210’ sample. For the ’BearCreek’ sample, we used a 10% fraction of the best ROC threshold due to our high coverage simulation.

#### 3.2.2 Analysis of a 30 kya permafrost sample from Canada

We used the ”Bear Creek” sample from Murchie, Monteath, et al. (2021) (BioProject: PRJNA722670, Accessions: SRR14265632 – SRR14265692), in which the original publication detected the families Phasianidae (e.g. pheasants), Elephantidae (e.g. mammoths), Equidae (e.g. horses), Bovidae (e.g. bison), Cricetidae (e.g. lemmings, muskrats) and Cervidae (e.g. deers). The sample was used for target enrichment (Murchie, Kuch, et al. 2021) and sequenced with Illumina HiSeq (Murchie, Monteath, et al. 2021).

As our empirical data represented only 10% of the simulated data, we set a threshold on the number of aligned reads set at 10% of that obtained on simulations. The detection threshold for HAYSTAC remained the same as in subsection 3.2.1. Analysis of the permafrost sample with all four tools is shown in Figure 7: euka and HAYSTAC report all taxa discovered in the original publication except for the family Cricetidae, which does not pass euka’s minimum breath of coverage filter and HAYSTAC’s minimum number of reads. The missing taxon Cricetidae could be caused by a different read length threshold in the original publication. Murchie, Monteath, et al. (2021) allowed every read with a minimum length of 24 bp, where we set the minimum read length to 30 bp for this re-analysis. Bowtie2+ngsLCA reports a small number of alignments from Galliformes on a higher taxonomic rank (Neognathae).

#### 3.2.3 Re-analysis of a 2-million-year-old eDNA sample from Green-land

Our final empirical sample is the 2-million-year-old sediment samples from the Kap Københaven Formation in Greenland, published by Kjær et al. (2022) (ENA: PRJEB55522). The samples were shotgun-sequenced using Illumina HiSeq and NovaSeq, using mtDNA capture enrichment (Kjær et al. 2022). The original publication detected five mammalian families: Anati-dae (e.g. swans, ducks), Cervidae (e.g. deers), Cricetidae (e.g. lemmings, muskrats), Elephantidae (e.g. mastodons) and Leporidae (e.g. rabbits).

We reanalyzed the dataset to demonstrate its ancient eDNA screening utility. All mammalian families from the publication were detected (Supplementary Figures 9–14), highlighting euka’s effectiveness with low-abundance, damaged data. An unreported taxon, Folivora (sloths), was found (Supplementary Figure 11). However, visual inspection revealed an unusual fragment length distribution, suggesting potential exogenous contamination, as the maximum occurs above 80 bp, atypical for authentic aDNA fragments.

## 4 Conclusion

This study indicates that euka supplements currently available tools in effectively detecting taxonomic groups and quantifying their corresponding abun-dance from simulated eDNA and ancient eDNA samples, subject to euka’s database limitations as discussed above (Subsection 2.7). Our results, tested on euka’s taxonomic resolution level, demonstrate that euka’s ancient-aware maximum-likelihood framework is more precise in identifying correct taxo-nomic assignments than contemporary methodologies. Although our filters are effective, detecting arthropodic taxa should still be approached with caution due to the abundance of low-complexity regions in arthropodic genomes and a dearth of reference genomes, making spurious mappings more prob-able. Additionally, we minimise the false detection of taxa in a sample by implementing multiple filters, which otherwise would require the use of sepa-rate tools, as evidenced by our benchmarking experiments on simulated and empirical data. Furthermore, our tool outperforms existing quantification tools regarding both accurate abundance estimation and runtime.

Our user-friendly command-line tool features a comprehensive tetrapod and arthropod database, enabling rapid profiling of large ancient eDNA datasets. It offers detailed sample summaries, including abundance esti-mates, damage profiles, fragment length distributions, and coverage assessments for each detected taxon. Visual output aids in identifying non-ancient taxa or potential modern contamination. euka’s all-in-one workflow simplifies and enhances initial ancient eDNA sample characterisation.

## 5 Main Figures

## 6 Declarations

### 6.1 Software Versions

We used vg version 1.44.0 - ’Solara’, ODGI version v0.6.3-52-g0d7f950 - ’Pulizia’, EMBOSS version 6.6.0, PRANK version v.170427 for the construction of euka’s pangenome graph structure and any mapping done with euka. For the bench-marking tests we used MALT version 0.5.3, MEGAN6 version 6.24.1, Bowtie2 version 2.4.4, ngsLCA version 950f5af (htslib: 1.13) and HAYSTAC version v0.4.8. Our simulated environments were created using Snakemake version 5.10.0, ART version 2.5.8 and leeHom version 1.2.15. All plots were produced using R version 4.1.2 ’Bird Hippie’.

## 6.2 Acknowledgements

We would like to thank Tyler Murchie and Hendrik Poinar for providing us with empirical data to test euka. We would also like to thank Giulia Zampirolo for testing our tool and Ana Duggan for reading and commenting on our manuscript. At last, we want to thank Haoming Yang for building euka’s progress bars.

## 6.4 Authors’ Contributions

The method was developed and implemented by NAV, JDR and GR. Additional subcommands of the method were implemented by MS and JV. MWP and AGP supported the implementation and interpretation of the data. NAV performed all tests. IT infrastructure support for the project was given by PWS. All authors read and approved the final manuscript.

## 6.5 Data Availability

vgan can be built from source or downloaded as a static binary from https://github.com/grenaud/vgan. It is also available on BioConda https://bioconda.github.io/recipes/vgan/README.html and as a Docker image.

Scripts and data pertaining to the benchmarking experiments are available at https://github.com/nicolaavogel/eukaPaperData.git.

## Footnotes

1 (https://github.com/grenaud/vgan)

2 (https://github.com/jts/sga)

