## Supplementary Material for "euka: Robust detection of eukaryotic taxa from modern and ancient environmental DNA using pangenomic reference graphs"

### Contents

**1 Pangenome Graph Description** **4**

**2 Database Construction and Curation** **5**

2.1 Database Construction . . . . . . . . . . . . . . . . . . . . . . . 5

2.2 Database Curation . . . . . . . . . . . . . . . . . . . . . . . . 7

**3 Taxonomic Pangenome Graph Construction** **9**

**4 Mapping Parameters** **10**

**5 Ancient-aware-likelihood Framework** **10**

**6 Minimum Breadth of Coverage Estimation** **13**

**7 Abundance Estimation using MCMC Sampling** **15**

7.1 Metropolis-Hastings algorithm for sampling from a distribution over V . . . . . . . . . . . . . . . . . . . . . . . . . . . . 18

**8 Creation of Simulated Metagenomic Environments** **20**

**9 Feature comparison between existing methods** **21**

**10 Benchmarking Commands** **23**

**11 Supplementary Figures 25**

### 1 Pangenome Graph Description

A pangenome graph, otherwise known as a variation graph, is a single data structure that represents a collection of genomes and their interrelation. In particular, a pangenome graph is a tuple.

$$G = (V, P, E)$$

Here $V$ are the graph's nodes, which contain bases of a nucleotide sequence. These nodes have a continuous identification number (node ID) used to determine the exact position in the graph space. To reflect the double-stranded nature of DNA, these nodes have two possible orientations, in the sense that the node sequence in one orientation is the reverse complement of the node sequence in the opposite orientation. $P$ represent the graph's embedded paths, which are sequences that can be reconstructed by walking the graph across a series of nodes in a given orientation in a particular way. Finally, $E$ denotes the edges of the graph. For an arbitrary pair of nodes $(V_1, V_2) \in G$, an edge exists between $V_1$ and $V_2$ only if at least one embedded path traversing those nodes (in either order) exists without traversing any intermediate node (A. M. Novak et al. 2017; Paten, A. Novak, and Haussler 2014; Garrison et al. 2018).

A pangenome graph can consist of multiple connected components. A connected component exists if the embedded sequences show significant differences and a single data structure is unable to represent all sequences. Furthermore, variation graphs can be circularized. This is advantageous when representing naturally circular sequences such as mitochondrial or viral genomes (Garrison et al. 2018).

### 2 Database Construction and Curation

#### 2.1 Database Construction

`euka`'s database construction and curation includes the following steps. First, we extracted all Tetrapod and Arthropod genus names from the NCBI Taxonomy downloaded on 26/09/2022 with the `ete` toolkit version v3.1.2 (Huerta-Cepas, Serra, and Bork 2016) and downloaded every available complete mitogenome for each name from NCBI RefSeq (O'Leary et al. 2016). In total, there were 45959 Arthropod and 5594 Tetrapod genera. The mitogenomes are grouped in their respective genera and stored in multi-FASTA files. We aimed for one mitochondrial genome per species/ sub-species. If there was a choice among multiple references, we ranked the mitogenomes based on the number of "N" nucleobases present and only chose the highest ranking. We made sure to exclude duplicate samples. Finally, we downloaded the complete taxonomic lineage for each mitogenome, which is used for sorting and indexing the database (Huerta-Cepas, Serra, and Bork 2016).

After the initial download, the database consists of 4078 genera: 2487 arthropods and 1591 tetrapods. We use this database to define large enough taxonomic groups for our pangenome structures. The definition of our taxonomic groups is based on a merging algorithm, which uses the taxonomic lineage files. We count every mitogenome in every multi-FASTA file for each genus. If a file has more than ten mitogenomes, the genera are kept as is. If the genera have less than ten reference sequences, they will be sorted into their next highest taxonomic group. This process is repeated three times. During the second run, we lower the number of necessary reference sequences to seven, and in the third run, we lower it to five. We end up with a database consisting of 336 taxonomic groups. The groups have a taxonomic level from genus to order. Our goal was to be able to characterise an ancient eDNA sample as specifically as possible. However, we depend on the data availability and size for different taxonomic groups. For example, the genus *Rattus* is a model organism and, therefore, has well-curated reference genomes. Further, the genus consists of enough species to form a valuable group for our graph construction. There is a dearth of reference mitogenomes for some genera, especially within the Arthropods. Unfortunately, this leaves us at a higher taxonomic resolution of some taxa than other current taxonomic profilers. Still, we believe our resolution is sufficient to screen a sample and optimise downstream analysis. Our reference database for graph construction can only be as specific as the available data; as more reference mitogenomes become available, we will update our graph accordingly.

#### 2.2 Database Curation

The first step of our database curation is a check for contaminated sequences. We use the `conterminator` tool[1] (Steinegger and Salzberg 2020) with default settings. We did not identify any contaminated sequences.

We then rotated the mitogenome sequences in each taxonomic group. Mitogenomes are circular, and for most eukaryotes, there is no agreed-upon junction from which the sequence begins, as there is in the case of humans. This leads to shifted and reversed mitogenome sequences for different species in a taxonomic group. We rotate the sequences by identifying the longest common substring of each genome and using this as the common starting point.

Our second curation step involves testing the similarity of each group. Due to the fact that we have groups as high as the order rank, we need to ensure that the mitogenomes in the group are sufficiently similar to construct a reasonable variation graph. We use the Needleman-Wunsch pairwise alignment algorithm (Needleman and Wunsch 1970) from `EMBOSS` (Rice, Longden,

[1]https://github.com/martin-steinegger/conterminator

and Bleasby 2000) to compare each sequence in a group against the rest. The following criteria must be fulfilled by a mitogenome to be kept in the group: the summed score of similarity must be between 50 and 99%; the gap score can not be higher than 15%; and the Needleman-Wunsch score must be higher than half of the maximum score of the group. This curation step ensures we remove duplicate sequences, only include sequences of that particular taxonomic group, and remove reference sequences from a taxonomic group that are too divergent from the rest of the sequences. These curation steps are necessary to create a working pangenome graph. We were left with 336 taxonomic groups. If a group was left with only genomes belonging to the same lower taxonomic class, the group's name implied it was changed. The curated and rotated multi-FASTA file for each group were used to construct a multiple sequence alignment (MSA) with `PRANK` (Löytynoja and Russell 2014). We use this MSA to calculate the mean edit distance between all group sequences. The mean distance will increase with the diversity of a taxonomic group. We stored the mean edit distance with the taxonomic name of the group and their ID. This group information is essential for correctly identifying a sample with `euka`. Table 2 lists all 336 defined taxonomic groups with their sorted reference genomes and NCBI accession numbers.

### 3 Taxonomic Pangenome Graph Construction

After the database construction and curation, we used the 336 taxa and their associated reference sequences to construct the taxa-based pangenome graph with 336 connected components, 10812444 edges and 6925364 nodes using the `vg` toolkit (Garrison et al. 2018; Hickey et al. 2020), and `ODGI` (Guarracino et al. 2022). The first step of constructing our graph is to build an individual pangenome graph per taxon using `vg` subcommand `construct` for each MSA. The individual graphs are circularised to reflect the circular nature of the mitochondrial genome using `vg circularize`. Afterwards, their node IDs are manipulated to be continuous throughout all taxa using `vg ids`. The IDs of the first and last node of each connected component are used as an index for the taxonomic identification process (e.g. the taxon of earless seals (Phocidae) use node 5140148 to 5153433). We then run the `VG combine` subcommand to unite all individual graphs into a single graph structure, in which each taxon constitutes exactly one connected component. We index the graph's embedded mitogenomes using `vg gbwt` and create the requisite distance and snarl index files using `vg index` to allow for mapping with `vg giraffe` (Sirén et al. 2021). Finally, we transform the `vg` structure into `ODGI` format using `vg convert` to be able to efficiently load the taxa-based pangenome graph with `euka` at runtime.

### 4 Mapping Parameters

`euka` accepts FASTQ input files for mapping. The provided input is mapped against the embedded mitogenome paths in our taxa-based pangenome graph. We use the following parameters for mapping with `giraffe`: `-s 100 -D 400 -w 40 -v 2 -r 30`. `vg giraffe` needs a minimizer file that stores a set of k-mers stemming from the embedded paths (here: mitogenomes) in the reference graph. Our minimizer file was constructed with a window size of 9 base pairs and a k-mer size of 21 base pairs. These parameters allow us to map all reads with a minimum length of 29 base pairs which is the window size plus k-mer size minus one as per `giraffe`'s documentation (Sirén et al. 2021). A read-length cutoff of 30bp is a standard practice in ancient eDNA studies. To reduce false positives, we do not allow multi-mapping (i.e. mapping a sequence to multiple locations) in `giraffe`. After the mapping is complete, we have an internal GAM (Graph Alignment / Map) file that stores all the alignment information and is used for `euka`'s post-processing steps.

### 5 Ancient-aware-likelihood Framework

`euka`'s workflow post-processes the internally generated GAM file from the mapping with `giraffe`. `euka` considers one alignment at a time. Based on the indexing system using node IDs described in section 3, we link each alignment to a taxon in the graph. We then assess the likelihood of the alignment being correct with our ancient-aware-likelihood framework. For this assessment, we compute the likelihood ratio of two models: 1) model 1 posits that the alignment is correct, i.e. the fragment has been mapped to the correct taxonomic group and at the correct location, whereas 2) model 2 proposes that the alignment was spurious. The likelihood for each read is calculated by inferring the probability for each base, given if the base is a match or mismatch to the reference sequence. We consider that a DNA fragment aligned to a specific taxon contains matches or mismatches of a single base and insertion, deletions or softclips (i.e. unaligned portions) to a path in the graph. To quantify the probability that the DNA fragment stems from the specific taxon where it aligned (model 1), we consider that any match is the result of a lack of mutation, sequencing error or ancient damage, whereas a mismatch can be explained by either of the three.

For instance, let us say that at a given position, $r$, in the graph, we have observed the base $b$ in the DNA fragment. Let us assume that $b$ has a predicted sequencing error rate of $\epsilon$ (derived from the per base quality score) and a probability of a base substitution due to aDNA damage of $\delta$, which depends on the position of the base along the fragment and the type of substitution. Furthermore, we assume that the base has some probability of mutation, $\mu$, with respect to the reference, which is taken to be the same as the mean edit distance, divided by the length of the alignment, within the given taxonomic group to which $r$ belongs. If we assume that substitution occurs according to the Kimura 2-parameter model, the probability that $b$ aligns at that position under model 1 is then:

$$\begin{cases} (1-\mu)(1-\epsilon)(1-\delta) & (b = r), \\ (1-\mu)\frac{\epsilon}{3} + \mu\,\kappa_{tv}(1-\epsilon) + (1-\mu)(1-\epsilon)\,\delta & (b \neq r) \end{cases} \tag{1}$$

where $\kappa_{tv}$ is the transition/transversion rate ratio, which is here taken to be $\frac{1}{22}$.

As for model 2, the alignment to the graph is due to chance meaning that the fragment does not stem from the mitogenomes of the taxon. In order to compute the probability of a match or a mismatch for a random sequence of DNA, we simulated 1M random reads using the nucleotide frequency of mitogenomes (namely 36% As, 21% Cs, 12% Gs and 31% Ts) with read lengths ranging from 20bp to 150bp. We mapped these random sequences against our graph and estimated the average number of mismatches. We estimated that, on average, 25.5% of bases had a mismatch. Therefore, under model 2, each base has a mismatch probability of $\zeta = 0.255$ and a match probability of $(1-\zeta)$. The probability of base $b$ aligning to the graph base $r$ by chance is:

$$\begin{cases} (1-\zeta) & (b = r), \\ (\zeta) & (b \neq r) \end{cases} \tag{2}$$

For both models, we assign a standard probability of 0.02 in case of an insertion or deletion similar to the findings from Laricchia et al. 2022. Unresolved bases (i.e. Ns) are treated as a sequencing error and incur a probability of 0.25 as this represents the probability of a mismatch for a random DNA base. Soft-clipped bases are treated as a match in 25% of the cases and a mismatch in 75%. A soft clip interpreted as a mismatch has the probability of the base quality, a match of the probability $1-$ base quality. This entails that soft-clipped regions of the DNA sequence with high quality will be more unlikely than regions with poor quality. Once we have the likelihoods for both mutually exclusive models, we calculate the ratio between them. The log-likelihood ratio quantifies the degree to which we believe the fragment aligned with the correct taxon. If the log-likelihood ratio is less than 1 and, therefore, model 2 is much more likely than model 1, we believe the alignment to be spurious and discard it.

### 6 Minimum Breadth of Coverage Estimation

In our second post-processing step, we use the binning system to estimate the minimum breadth of coverage for each component in the pangenome graph. For our binning system, we first identified chunks of 1500 bases for each taxon pangenome graph using `vg`'s subcommand `chunk` (Hickey et al. 2020). The subcommand divides every embedded path (here: every embedded mitogenome of the taxonomic group) into `vg` structures of the set sequence length. We iterated through every chunk for every path recording the head node ID and the tail node ID with the `VG stats` subcommand to build a bin index. Based on our mitogenome rotation prior to the graph construction, we ensure that each taxonomic component begins and ends with the same node ID. For any intermediate node IDs defining the start and end of different bins, we took the minimum and maximum node ID for each chunk between all embedded mitogenomes. The Shannon entropy (Shannon 1948) is computed for each bin to measure the base diversity represented in the 1500bp. It is defined as follows:

$$H = -\sum_{i=1}^{n} p_i \log_2 p_i \tag{3}$$

where $H$ is the entropy score, $p_i$ is the frequency of a base $i$ at a given position, and the summation is over all possible bases. A high Shannon entropy represents a diverse sequence composition, while a low Shannon entropy score represents sequences with low complexity (e.g. high A/T content). Therefore, the lower the entropy, the more uncertain we become about an alignment in this genomic region as the probability of a spurious mapping increases with low-complexity sequences. We set a global threshold of 1.17, meaning we do not trust the alignments in a region with lower entropy. When filtering on an individual taxon, the coverage of all bins connected to it is examined to ensure coverage uniformity, with at least one alignment per bin. In the standard settings of `euka`, the number of allowed bins with an entropy score lower than 1.17 is restricted to six, reducing the incidence of falsely positive detected taxa. However, these parameters can be tailored to facilitate the detection of distinct taxonomic groups. For instance, the subgraph of the ant family (Formicidae) primarily comprises low-entropy bins that fall below the standard threshold. Consequently, users must either decrease the entropy threshold or permit fewer bins with a low-entropy score to detect this taxon. A lot of the arthropod taxa have low-complexity genomes, and spurious mappings can accrue more often than for tetrapod taxa. Our entropy threshold and minimum bin filter are to ensure a minimal amount of false positive detection for the arthropods. However, results for these taxa, especially with a large amount of low-entropy sequences, should still be considered carefully. We provide a list of all taxa in `euka`'s database with their assigned bins (denoted by the first node ID of the bin and the last node ID of the bin) and the bin's entropy score named euka_db.bins on our GitHub `https://github.com/grenaud/vgan` within the share/euka_dir directory.

### 7 Abundance Estimation using MCMC Sampling

Every taxon and their associated alignments that passed our post-processing steps are believed to be of high confidence. The number of high-confidence alignments associated with the different taxa contains information about their relative abundances. Naively, the relative abundance of some taxon, $i$, can simply be estimated as the number of alignments associated with taxon $i$, divided by the total number of alignments. However, the true underlying frequencies in the original biological environment can, of course, have differed from these estimates. First, the number of sequence reads can be considered to be a random multinomial sample from the DNA library, which is itself a random sample from the original environment. Secondly, it is possible that an alignment has been erroneously associated with a taxon due to a mapping error (typically caused by repetitive sequences) or spurious sequence similarity. In order to estimate the relative abundances while accounting for uncertainty, we have here used Markov chain Monte Carlo (MCMC) sampling to obtain a probability distribution (and corresponding credible intervals) over the taxon frequencies. We have designed the method to put more weight on high-confidence and unambiguous alignments.

The goal is to estimate a vector of frequencies, $V = (v_1, v_2, \ldots, v_n)$, where $v_i$ is the relative abundance of taxon $i$, and $n$ is the number of detected taxa. By definition $0 < v_i < 1$ and $\sum_{i=1}^{n} v_i = 1$. Specifically, we use the Metropolis-Hastings algorithm to obtain a probability distribution over possible values of $V$ in a way where we put more weight on alignments where we have high confidence that mapping is correct and alignment is not spurious. We then take the following to be the likelihood of a read $r$ being aligned to taxon $i$:

$$P(r = i|V) = v_i \left[ \left( \frac{M1_{r,i}}{M1_{r,i} + M2_{r,i}} \right) MQ_{r,i} \right] + (1 - v_i) e_{non-i} (\frac{1}{n-1}) \qquad (4)$$

Here, the subscript $r, i$ represents the specific combination of read and taxon for our parameters, $\frac{M1_{r,i}}{M1_{r,i}+M2_{r,i}}$ is the posterior probability of Model 1 being correct for the read $r$ that is mapped to taxon $i$, and $MQ_{r,i}$ is the mapping quality on the probability scale (not the PHRED scale). The term $e_{non-i}$ is the probability of mapping any read, $s$ that is not from taxon $i$, to the wrong taxon and is defined as:

$$e_{non-i} = \frac{\sum_{s \neq r, j \neq i} [1 - MQ_{s,j} \frac{M1_{s,j}}{M1_{s,j}+M2_{s,j}}]}{n_{j \neq i}} \qquad (5)$$

where $n_{j \neq i}$ is the number of reads not mapped to taxon $i$. The total likelihood for all reads is found by multiplying the individual likelihoods. Let $\mathbb{R}$ be the set of all reads. The total likelihood for all reads is defined as:

$$P(\mathbb{R}|V) = \prod_{j=1}^{k} P(r = j|V) \qquad (6)$$

where $k$ is the total number of reads.

#### 7.1 Metropolis-Hastings algorithm for sampling from a distribution over V

In the following, we describe the individual steps of our MCMC sampling algorithm:

1. For each taxon, $i$, compute $e_{non-i}$. This is the average value of $(1 - MQ\frac{M1}{M1+M2})$ for all those reads, $s$, that are not mapped to taxon $i$ and are found according to equation (5). These values are calculated before starting the MCMC.

2. Initialize $V_1 = (v_1, v_2, \ldots, v_n)$ such that $v_i = \frac{c_i}{k}$, where $c_i$ is the count of alignments associated with taxon $i$.

3. For $t = 1, 2, \ldots, N_{\text{samples}}$ repeat the steps $4 - 7$ below to generate the $t$'th sample.

4. Propose new vector $V^*$ based on the current value $V_t$ as follows:

   (a) Convert the frequency vector $V_t$ to a real-valued vector $U_t$ using the additive log-ratio transform:

   $$U_t = \left(\log\left(\frac{v_1}{v_n}\right), \log\left(\frac{v_2}{v_n}\right), \ldots, \log\left(\frac{v_{n-1}}{v_n}\right)\right) \quad (7)$$

   Note that $V_t$ has length $n$, while $U_t$ has length $n - 1$.

(b) Compute the perturbed vector $U^*$ by adding random, zero-centered terms to each element:

$$U^* = (u_1 + \epsilon_1, u_2 + \epsilon_2, \ldots, u_{n-1} + \epsilon_{n-1}) \tag{8}$$

where $\epsilon_i \sim \mathcal{N}(\mu = 0, \sigma^2 = 0.02)$

(c) Convert $U^*$ back to a length-n frequency vector $V^*$:

$$V^* = \left( \frac{e^{u_1}}{\sum_i e^{u_i} + 1}, \ldots, \frac{e^{u_{n-1}}}{\sum_i e^{u_i} + 1}, \frac{1}{\sum_i e^{u_i} + 1} \right) \tag{9}$$

5. Compute the likelihoods $P(\mathbb{B}|V_t)$ and $P(\mathbb{B}|V^*)$

6. If $P(\mathbb{B}|V^*) > P(\mathbb{B}|V_t)$ : Accept the proposed move in parameter space and set $V_{t+1} = V^*$

7. If $P(\mathbb{B}|V^*) < P(\mathbb{B}|V_t)$ : Accept the move with probability $\alpha = \frac{P(\mathbb{B}|V^*)}{P(\mathbb{B}|V_t)}$:

   (a) Draw a random number $r$ in $[0, 1]$

   (b) Accept the proposal if $r < \alpha$, and set $V_{t+1} = V^*$

   (c) Reject the proposal if $r > \alpha$, and set $V_{t+1} = V_t$

By default, we repeat this process 10,000 times and discard the first 100 samples; the user can customise those parameters. However, we have seen that 10,000 iterations seem to be sufficient to explore the probability landscape around the global maximum, even if our initial vector was chosen randomly.

After the last iteration, we sort the individual abundances per taxa to extract the posterior median and the 85% and 95% credible intervals. We report our estimates in the TSV files with the suffixes "abundance" and "detected" alongside their respective taxa and the total number of alignments, completing `euka`'s final workflow step.

### 8 Creation of Simulated Metagenomic Environments

To test `euka`'s accuracy against current methods, we simulated three metagenomic environments where the taxa and their abundances are known. We compared all the results based on the taxa that `euka` defines in its database table 2. The first environment is based on Pedersen et al. (2021) and Ardelean et al. (2020) cave sediment samples representing a complex environment with fourteen represented taxa. The second environment is based on Murchie, Karpinski, et al. (2022) and Murchie, Monteath, et al. (2021) permafrost sample and simulates high-abundance taxa after performing mitochondrial capture enrichment. Finally, the third environment is based on the study by Gelabert et al. (2021) and represents a sedimentary cave environment with ancient human presence. In all three environments, the proportion of detectable mtDNA was around 0.01% of all reads. Additionally, we added nuclear DNA from the genomes of the taxa we used for the environment, comprising roughly 4.99% of the metagenomic sample. The remaining 95% of the environment comprises bacteria (80%), fungi (6%), plants (1,6%), viruses (0.4%) and archaea (7%) according to Tkacz, Hortala, and Poole 2018.

Three simulated environments were created using the following workflow: We downloaded mitochondrial and nuclear reference genomes from NCBI RefSeq (O'Leary et al. 2016 for our known taxa. Additionally, we downloaded a variety of sequences from plants, fungi, bacteria, viruses and archaea. Fragments from the different genomes were produced using `gargammel` (Renaud, Hanghøj, et al. 2017) with three levels of simulated ancient damage: no damage, medium damage (substitution rates taken from (Olalde et al. 2014)) and high damage (substitution rates taken from (Günther et al. 2015)). The fragments were merged with `leeHom` (with ancient parameters) (Renaud, Stenzel, and Kelso 2014). Afterwards, we filtered out low-complexity reads using `sga` preprocessing [2] with a dust filter of 1. The `Snakemake` pipeline used for creating our simulated environments, including a list of NCBI accession numbers for the sampled genomes, can be found `https://github.com/nicolaavogel/eukaPaperData.git`.

### 9 Feature comparison between existing methods

For an overview of the tools, we created a feature comparison in Table 1.

[2](https://github.com/jts/sga)

| Feature | euka | HAYSTAC | Bowtie2 + ngsLCA | MALT |
| --- | --- | --- | --- | --- |
| Output visualisation | ✓ | ✓ | ✓ | ✓ |
| Database provided | ✓ | ✓ | X | X |
| Abundance estimation | ✓ | ✓ | X | X |
| Coverage evenness test | ✓ | ✓ | X | X |
| Ancient damage profile estimation | ✓ | ✓ | X | X |
| Fragment length estimation | ✓ | ✓ | X | X |
| No taxonomic limitations | X | ✓ | ✓ | ✓ |

Table 1: Feature comparison across euka, HAYSTAC, Bowtie2 + ngsLCA and MALT

### 10 Benchmarking Commands

#### 10.1 MALT

```
MALT-0.5.3/malt-build -i [databse_sequences].fa.gz -d index
--sequenceType DNA
malt-run -m BlastN -d [index] -i [environment.fq.gz] -o [output_dir]
-t 20 -top 20 -e 0.00001 -mq 1 -mif -id 95 -sup 3 -wlca -lcp 80
```

#### 10.2 Bowtie2

```
bowtie2 --threads 20 -k 1000 -D 15 -R 2 -N 1 -L 22 -i S,1,1.15 --np 1
--mp "1,1" --rdg "0,1" --rfg "0,1" --score-min "L,0,-0.1" -x [database]
-U  [environment.fq.gz]  --no-unal | samtools view -bS - > [input].bam
samtools sort -n -O bam -o [input].sort.bam -@ 20 [input].bam
```

#### 10.3 ngsLCA

```
ngsLCA -simscorelow 0.95 -simscorehigh 1.0
-names ncbi_tax_dmp/names.dmp -nodes ncbi_tax_dmp/nodes.dmp
-acc2tax ncbi_tax_dmp/nucl_gb.accession2taxid.gz
-bam [input].sort.bam -outnames [output]
```

#### 10.4 HAYSTAC

```
haystac sample --output [environment_dir] --fastq [environment.fq.gz]
--core 20 --trim-adapter False
haystac analyse --mode abundances --database [database]
--sample [environment_dir] --output [ouput_dir] --bowtie2-threads 20
--aDNA --cores 20
```

### 11 Supplementary Figures

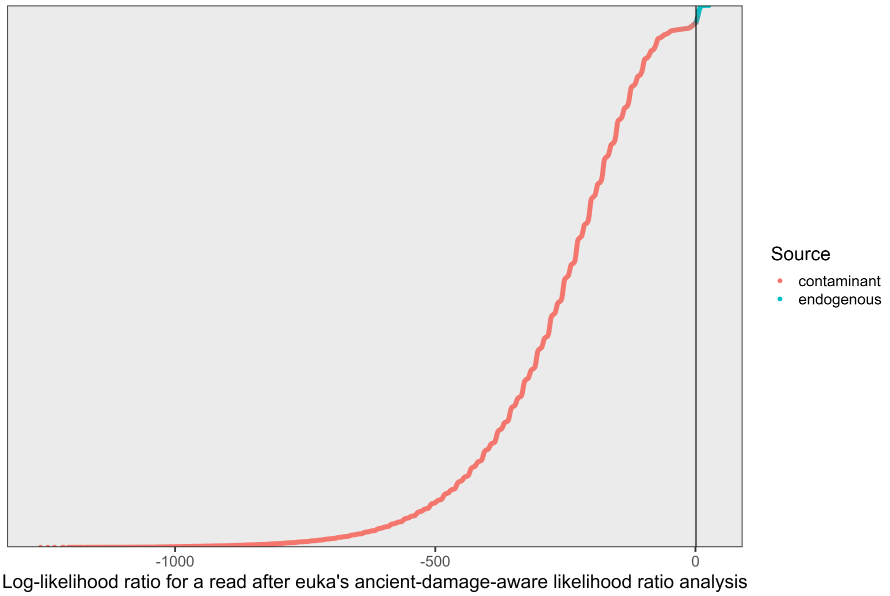

Figure 1: Log-likelihood assessment of every mapped read from our simulated cave environment with high simulated damage rates. The vertical line drawn at position 1 represents the log-likelihood needed for model 1 (the mapped read stems from the taxon) to be favoured. We coloured the reads by their source: endogenous (simulated mitochondrial reads) and contaminants (reads simulated from bacteria, viruses, archaea, fungi, plants or nuclear genomes). Every read with a log-likelihood lower than 1 is disregarded by `euka`'s ancient-aware maximum-likelihood framework.

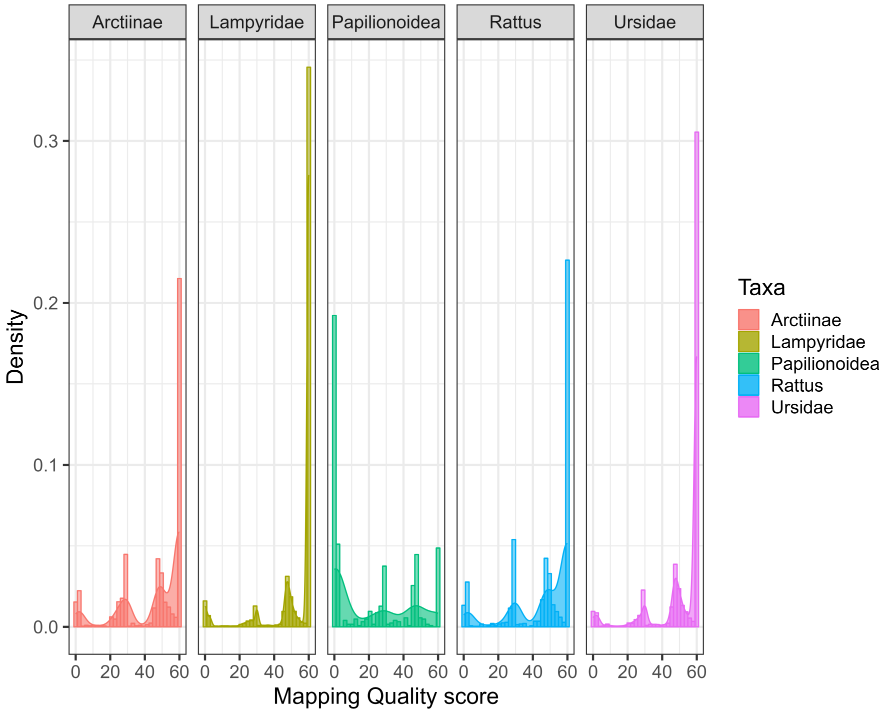

Figure 2: Assessment of Mapping Quality threshold between five different taxa: the taxa Arctiinae, Lampyridae, Rattus, and Ursidae are true positives in the simulated environment, whereas Papilionoidea attracts alignments due to its low-entropy genome. The comparison shows the optional cutoff mapping quality to filter out as many false-positive Papilionoidea fragments as possible.

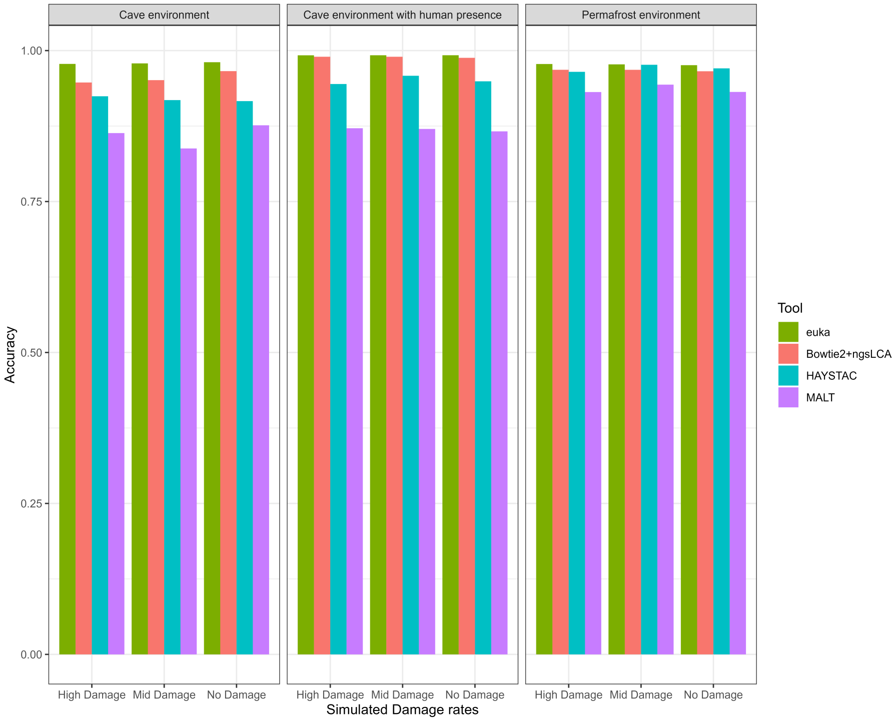

Figure 3: Uncapped accuracy comparison between all four tools (`euka`, `Bowtie2` + `ngsLCA`, `HAYSTAC`, `MALT`) in three different simulated environments over three different levels of simulated damage.

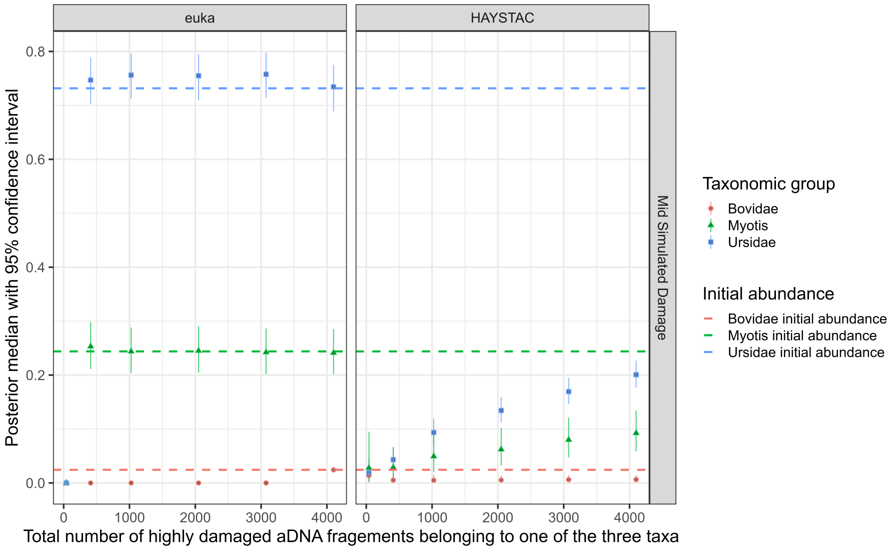

Figure 4: `euka`'s and `HAYSTAC`'s abundance estimation for a downsampled ancient metagenomic sample of three taxa with a medium damage rate. To show `euka`'s robustness, we downsampled the original sample estimating the point where `euka` would be unable to detect the different taxa. The initial abundance of the three taxa was represented with a dotted line. On average, taxon detection can be confidently determined with `euka` around 50 reads present. `euka` abundance estimation stayed robust until it was downsampled to 1% of its original input.

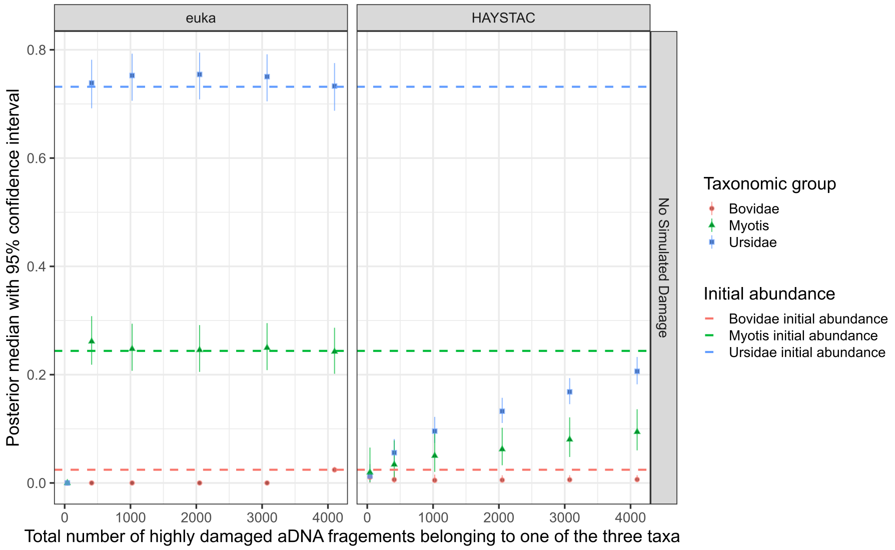

Figure 5: `euka`'s and `HAYSTAC`'s abundance estimation for a downsampled modern metagenomic sample of three taxa. To show `euka`'s robustness, we downsampled the original sample estimating the point where `euka` would be unable to detect the different taxa. The initial abundance of the three taxa was represented with a dotted line. On average, taxon detection can be confidently determined with `euka` around 50 reads present. `euka` abundance estimation stayed robust until it was downsampled to 1% of its original input.

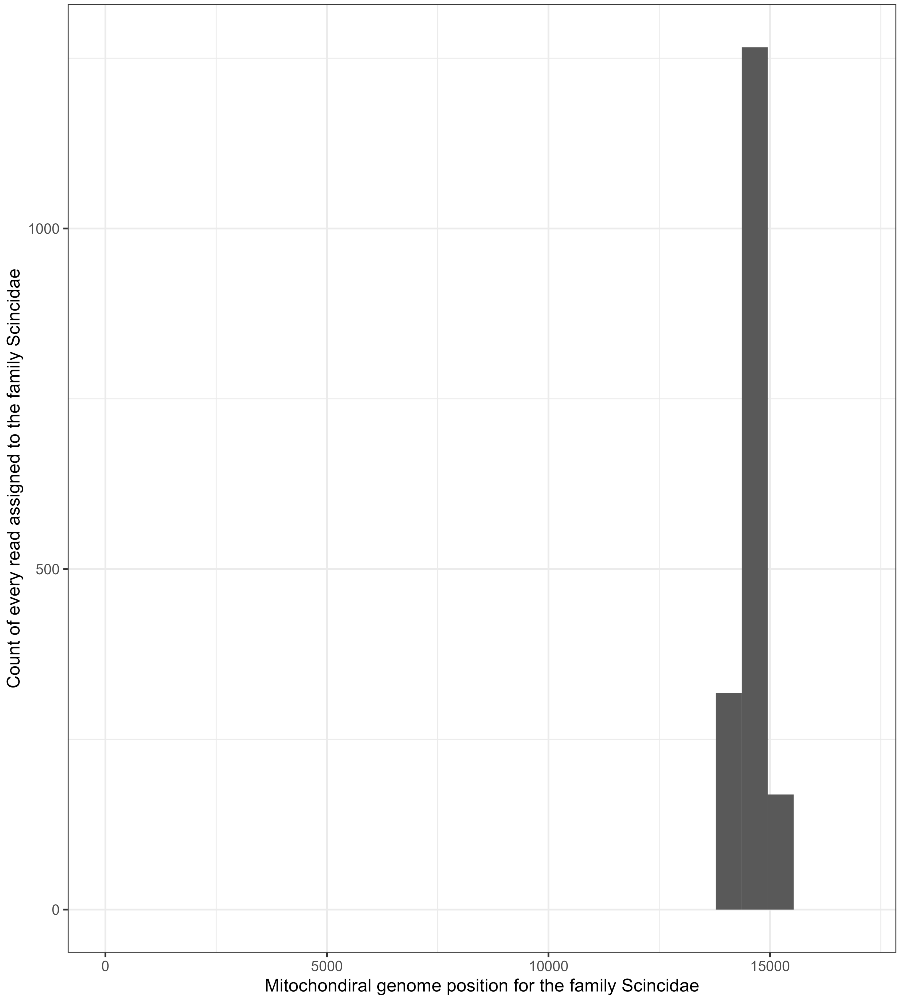

Figure 6: `Bowitie2` + `ngsLCA`'s resulting positions in the genome for all reads taxonomically assigned to the family Scincidae across the average length of a mitochondrial genome for the family Scincidae (approximately 16.500 base pairs).

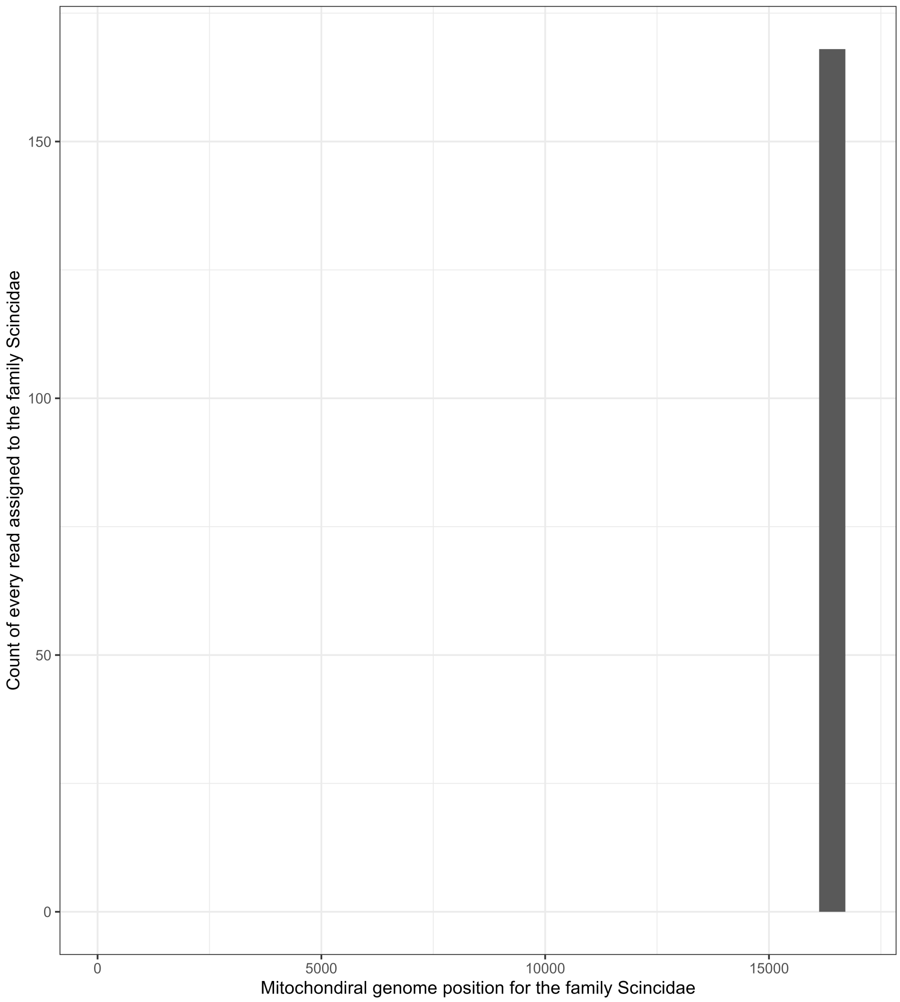

Figure 7: `HAYSTAC`'s resulting positions in the genome for all reads taxonomically assigned to the family Scincidae across the average length of a mitochondrial genome for the family Scincidae (approximately 16.500 base pairs).

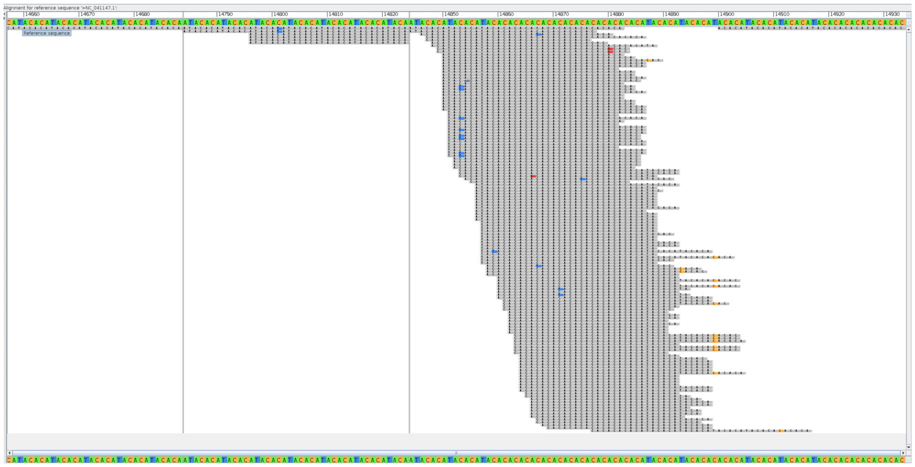

Figure 8: Picture from `MEGAN` alignment visualisation for all the reads mapped to *Isopachys gyldenstolpei* with `MALT` from the family Scincidae. This species was chosen as a representative as it had the most alignments from all members of the family. The picture focuses on the genomic positions between 14660 and 14930 base pairs, where all reads recorded are mapped.

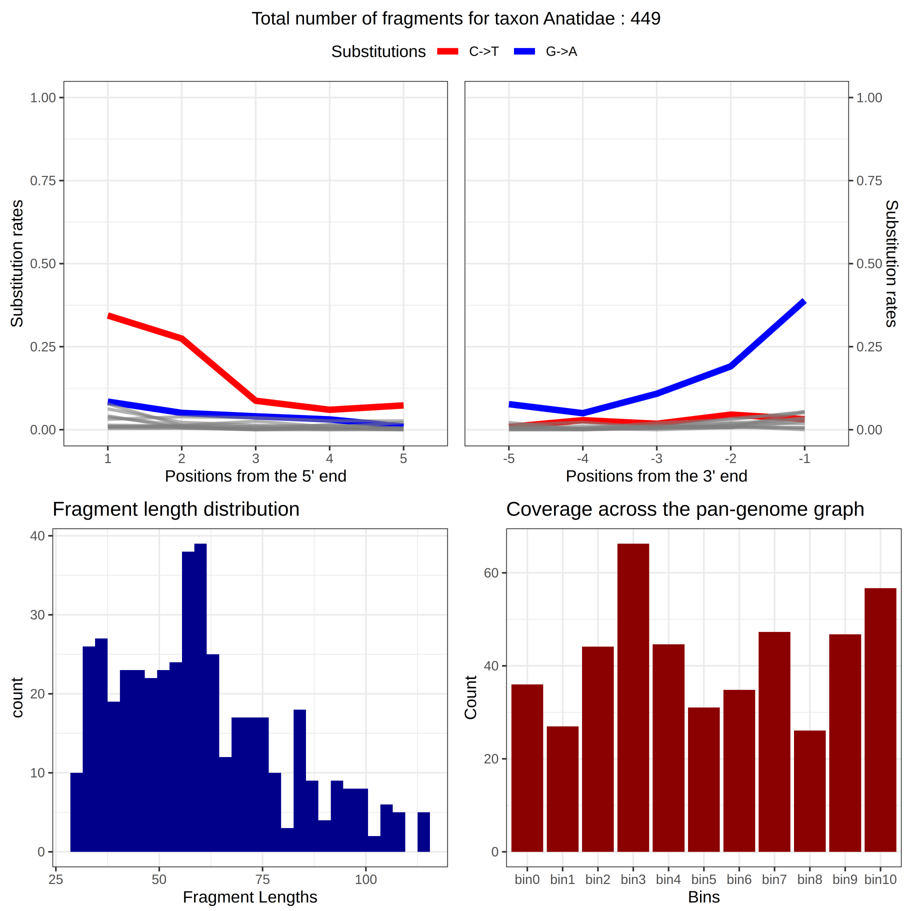

Figure 9: `euka`'s output graphic for the family Anatidae. The top panels show our damage rate estimation for the 5' end (top right) and the 3' end (top left). On the bottom right, the fragment length distribution is plotted. The bottom right is a visualisation of the minimal breadth of coverage for a taxon.

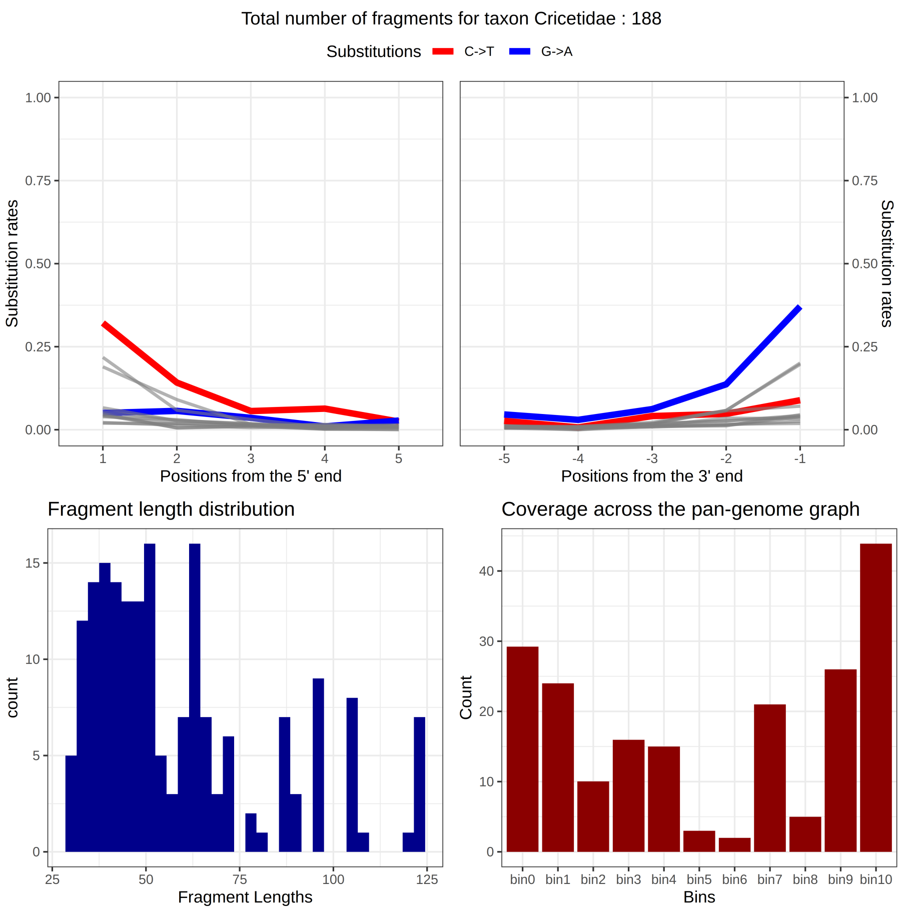

Figure 10: `euka`'s output graphic for the family Cricetidae. The top panels show our damage rate estimation for the 5' end (top right) and the 3' end (top left). On the bottom right, the fragment length distribution is plotted. The bottom right is a visualisation of the minimal breadth of coverage for a taxon.

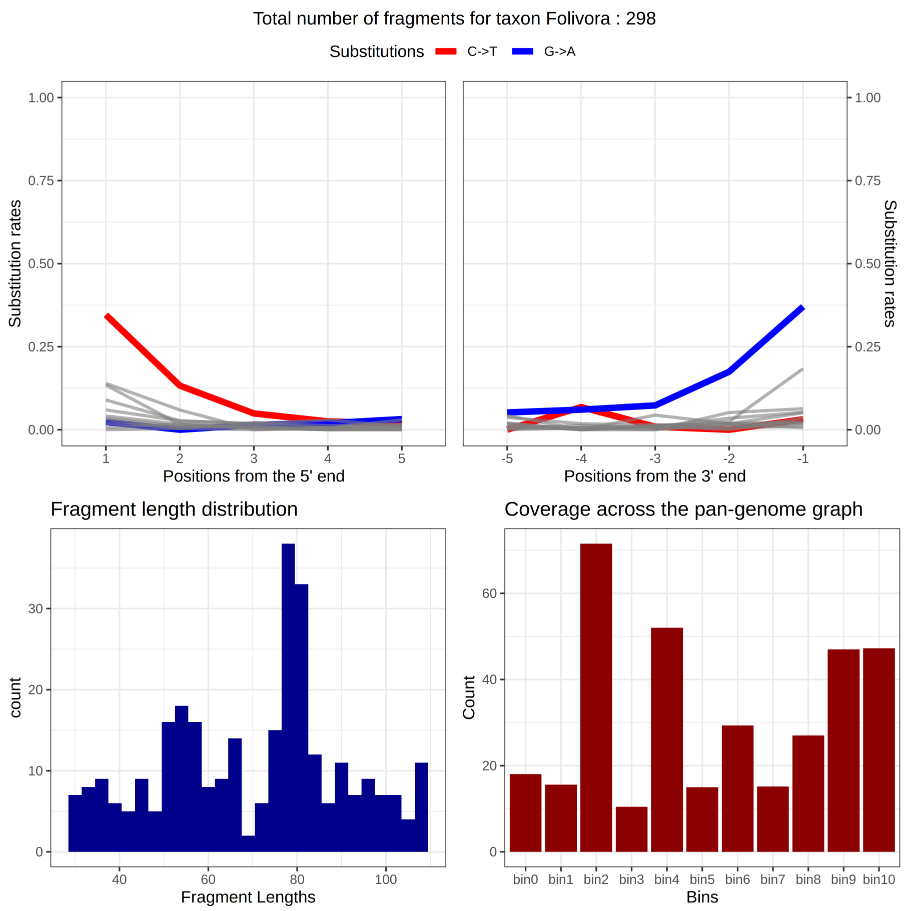

Figure 11: `euka`'s output graphic for the family Folivora. The top panels show our damage rate estimation for the 5' end (top right) and the 3' end (top left). On the bottom right, the fragment length distribution is plotted. The bottom right is a visualisation of the minimal breadth of coverage for a taxon.

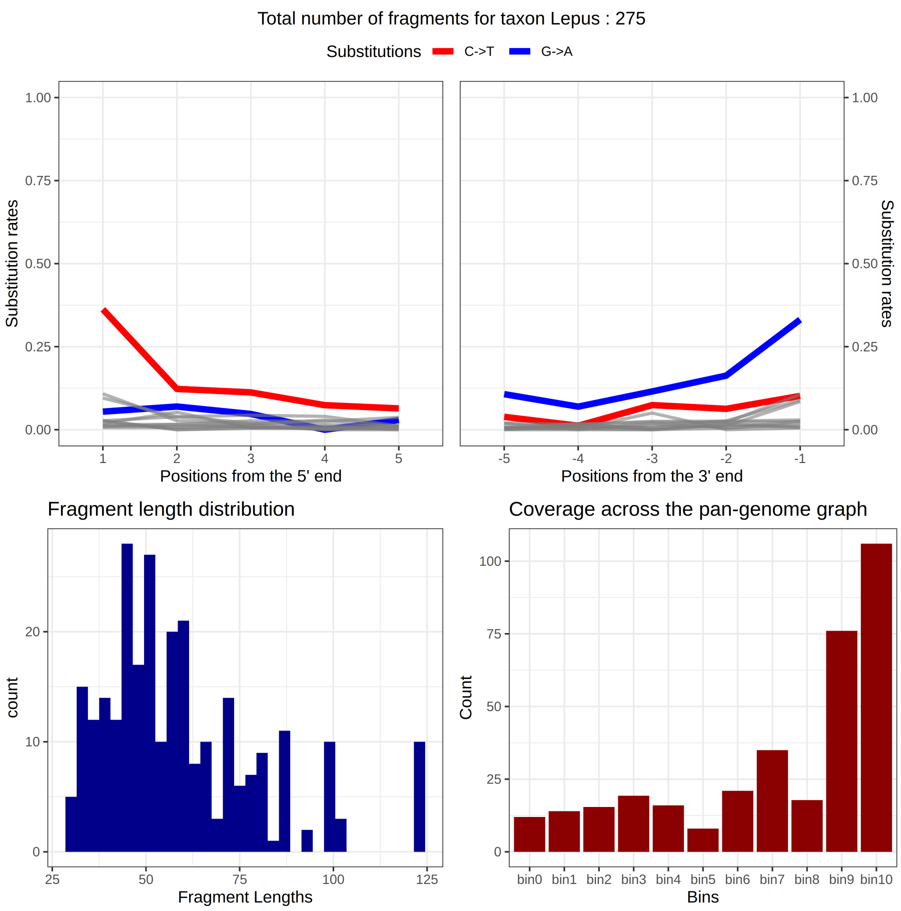

Figure 12: `euka`'s output graphic for the family Lepus. The top panels show our damage rate estimation for the 5' end (top right) and the 3' end (top left). On the bottom right, the fragment length distribution is plotted. The bottom right is a visualisation of the minimal breadth of coverage for a taxon.

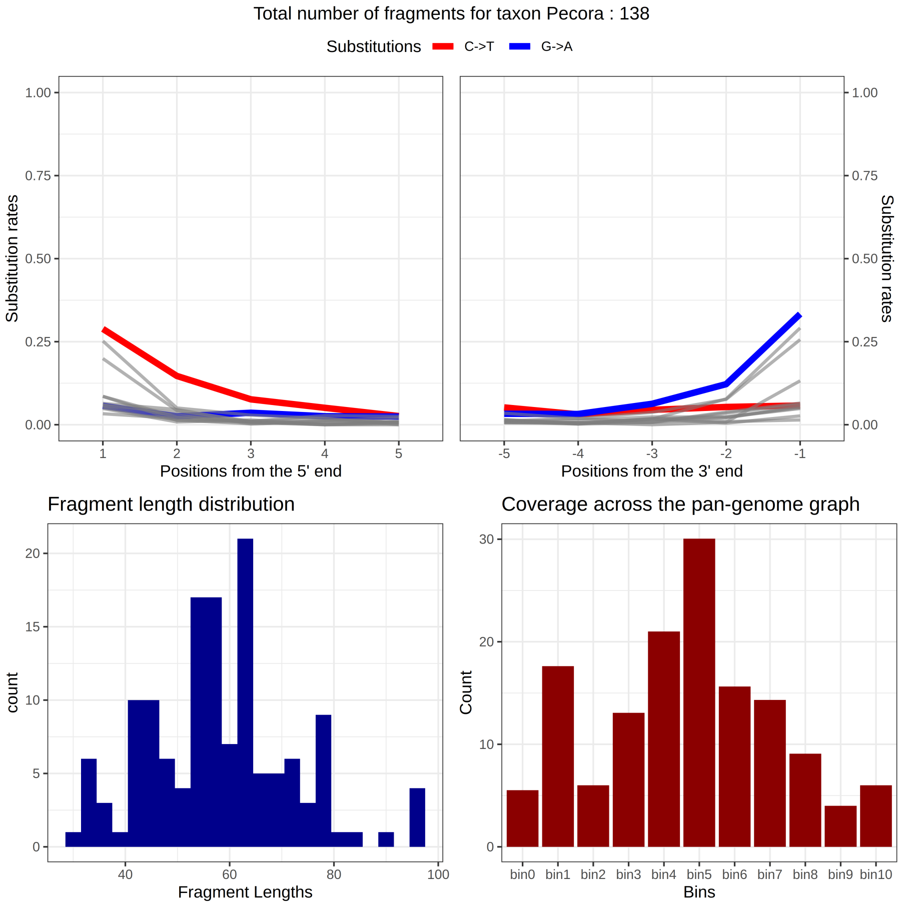

Figure 13: `euka`'s output graphic for the family Pecora. The top panels show our damage rate estimation for the 5' end (top right) and the 3' end (top left). On the bottom right, the fragment length distribution is plotted. The bottom right is a visualisation of the minimal breadth of coverage for a taxon.

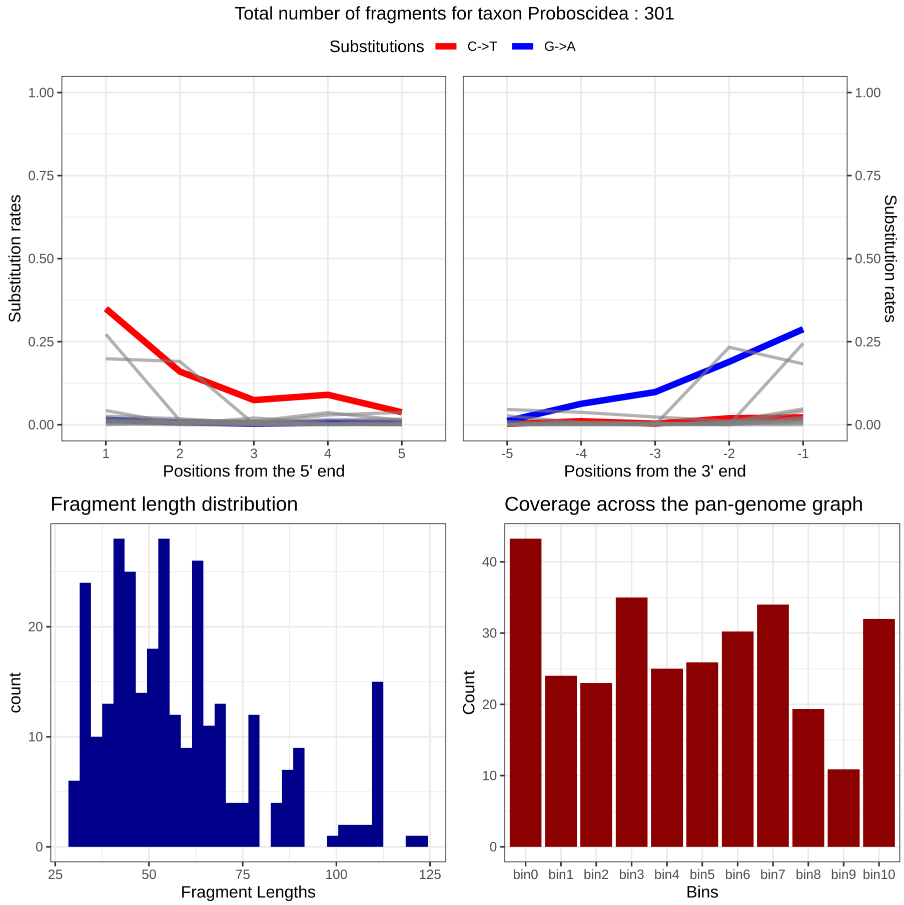

Figure 14: `euka`'s output graphic for the family Proboscidea. The top panels show our damage rate estimation for the 5' end (top right) and the 3' end (top left). On the bottom right, the fragment length distribution is plotted. The bottom right is a visualisation of the minimal breadth of coverage for a taxon.

| **Taxon** | **NCBI Accession Number** | ***Species name*** |
|---|---|---|
| Acalyptratae | NC_059897.1 | *Nothybus sumatranus* |
| Acalyptratae | NC_034924.1 | *Spaniocelyphus pilosus* |
| Acalyptratae | NC_034922.1 | *Cestrotus liui* |
| Acalyptratae | NC_060541.1 | *Homoneura interstincta* |
| Acalyptratae | NC_034923.1 | *Pachycerina decemlineata* |
| Acalyptratae | NC_016865.1 | *Fergusonina taylori* |
| Acalyptratae | NC_063098.1 | *Dryomyza anilis* |
| Acalyptratae | NC_026866.1 | *Nemopoda mamaevi* |
| Acalyptratae | NC_065365.1 | *Chyliza chikuni* |
| Acalyptratae | NC_065364.1 | *Chyliza bambusae* |
| Acalyptratae | NC_065363.1 | *Chamaepsila testudinaria* |
| Acalyptratae | NC_065368.1 | *Loxocera sinica* |
| Acalyptratae | NC_065367.1 | *Loxocera planivena* |
| Acalyptratae | NC_065366.1 | *Loxocera lunata* |
| Acalyptratae | NC_030246.1 | *Melanagromyza sojae* |
| Acalyptratae | NC_041083.1 | *Liriomyza chinensis* |
| Acalyptratae | NC_016716.1 | *Liriomyza huidobrensis* |
| Acalyptratae | NC_016713.1 | *Liriomyza bryoniae* |
| Acalyptratae | NC_015926.1 | *Liriomyza sativae* |
| Acalyptratae | NC_014283.1 | *Liriomyza trifolii* |
| Acaroidea | NC_038058.1 | *Rhizoglyphus robini* |
| Acaroidea | NC_023778.1 | *Aleuroglyphus ovatus* |

| | | |
|---|---|---|
| Acaroidea | NC_028725.1 | *Tyrophagus longior* |
| Acaroidea | NC_026079.1 | *Tyrophagus putrescentiae* |
| Accipitridae | NC_045364.1 | *Accipiter trivirgatus* |
| Accipitridae | NC_026082.1 | *Accipiter virgatus* |
| Accipitridae | NC_025580.1 | *Accipiter nisus* |
| Accipitridae | NC_011818.1 | *Accipiter gentilis* |
| Accipitridae | NC_045042.1 | *Aquila nipalensis* |
| Accipitridae | NC_035806.1 | *Aquila heliaca* |
| Accipitridae | NC_024087.1 | *Aquila chrysaetos* |
| Accipitridae | NC_029377.1 | *Buteo hemilasius* |
| Accipitridae | NC_029189.1 | *Buteo lagopus* |
| Accipitridae | NC_052805.1 | *Circaetus pectoralis* |
| Accipitridae | NC_035801.1 | *Circus melanoleucos* |
| Accipitridae | NC_036050.1 | *Gyps fulvus* |
| Accipitridae | NC_040858.1 | *Haliaeetus albicilla* |
| Accipitridae | NC_038195.1 | *Milvus migrans* |
| Accipitridae | NC_007599.1 | *Nisaetus alboniger* |
| Accipitridae | NC_007598.1 | *Nisaetus nipalensis* |
| Accipitridae | NC_015887.1 | *Spilornis cheela* |
| Accipitridae | NC_052803.1 | *Spizaetus tyrannus* |
| Accipitriformes | NC_063526.1 | *Cathartes burrovianus* |
| Accipitriformes | NC_007628.1 | *Cathartes aura* |
| Accipitriformes | NC_063525.1 | *Coragyps atratus* |

| | | |
|---|---|---|
| Accipitriformes | NC_063527.1 | *Sarcoramphus papa* |
| Accipitriformes | NC_058600.1 | *Vultur gryphus* |
| Accipitriformes | NC_008550.1 | *Pandion haliaetus* |
| Achelata | NC_052750.1 | *Panulirus penicillatus* |
| Achelata | NC_052749.1 | *Panulirus longipes* |
| Achelata | NC_039671.1 | *Panulirus argus* |
| Achelata | NC_028024.1 | *Panulirus cygnus* |
| Achelata | NC_028627.1 | *Panulirus versicolor* |
| Achelata | NC_016015.1 | *Panulirus homarus* |
| Achelata | NC_014854.1 | *Panulirus ornatus* |
| Achelata | NC_014339.1 | *Panulirus stimpsoni* |
| Achelata | NC_004251.1 | *Panulirus japonicus* |
| Achelata | NC_049031.1 | *Linuparus trigonus* |
| Achelata | NC_041155.1 | *Puerulus angulatus* |
| Achelata | NC_041153.1 | *Ibacus alticrenatus* |
| Achelata | NC_025581.1 | *Ibacus ciliatus* |
| Achelata | NC_044424.1 | *Parribacus antarcticus* |
| Achelata | NC_041156.1 | *Remiarctus bertholdii* |
| Achelata | NC_052010.1 | *Scyllarides haanii* |
| Achelata | NC_044425.1 | *Scyllarides squammosus* |
| Achelata | NC_020022.1 | *Scyllarides latus* |
| Achelata | NC_021753.1 | *Palinurellus wieneckii* |
| Acraea | NC_029514.1 | *Acraea kalinzu* |

| | | |
|---|---|---|
| Acraea | NC_029513.1 | *Acraea vestalis* |
| Acraea | NC_029512.1 | *Acraea parrhasia* |
| Acraea | NC_029511.1 | *Acraea alcinoe* |
| Acraea | NC_029510.1 | *Acraea serena* |
| Acraea | NC_029508.1 | *Acraea lycoa* |
| Acraea | NC_029507.1 | *Acraea perenna* |
| Acraea | NC_029505.1 | *Acraea penelope* |
| Acraea | NC_029504.1 | *Acraea poggei* |
| Acraea | NC_029503.1 | *Acraea rogersi* |
| Acraea | NC_029502.1 | *Acraea bonasia* |
| Acraea | NC_029501.1 | *Acraea pharsalus* |
| Acraea | NC_029500.1 | *Acraea circeis* |
| Acraea | NC_029499.1 | *Acraea zetes* |
| Acraea | NC_029498.1 | *Acraea acerata* |
| Acraea | NC_029497.1 | *Acraea egina* |
| Acraea | NC_029496.1 | *Acraea jodutta* |
| Acraea | NC_013604.1 | *Acraea issoria* |
| Acraea | NC_029509.1 | *Acraea polis* |
| Acrididae | NC_034674.1 | *Aiolopus thalassinus* |
| Acrididae | NC_052715.1 | *Anapodisma miramae* |
| Acrididae | NC_025946.1 | *Angaracris rhodopa* |
| Acrididae | NC_046527.1 | *Apalacris nigrogeniculata* |
| Acrididae | NC_039962.1 | *Arcyptera meridionalis* |

| | | |
|---|---|---|
| Acrididae | NC_013805.1 | *Arcyptera coreana* |
| Acrididae | NC_052731.1 | *Bryodema kozlovi* |
| Acrididae | NC_046554.1 | *Bryodemella tuberculata diluta* |
| Acrididae | NC_053659.1 | *Caryandoides hunanica* |
| Acrididae | NC_059683.1 | *Ceracris hainanensis* |
| Acrididae | NC_059852.1 | *Ceracris hoffmanni* |
| Acrididae | NC_043956.1 | *Ceracris fasciata fasciata* |
| Acrididae | NC_025285.1 | *Ceracris versicolor* |
| Acrididae | NC_019994.1 | *Ceracris kiangsu* |
| Acrididae | NC_019993.1 | *Chondracris rosea* |
| Acrididae | NC_056786.1 | *Chorthippus parallelus erythropus* |
| Acrididae | NC_048465.1 | *Chorthippus fallax* |
| Acrididae | NC_011095.1 | *Chorthippus chinensis* |
| Acrididae | NC_029408.1 | *Compsorhipis davidiana* |
| Acrididae | NC_041412.1 | *Dasyhippus barbipes* |
| Acrididae | NC_042904.1 | *Diabolocatantops pinguis* |
| Acrididae | NC_039408.1 | *Dnopherula yuanmowensis* |
| Acrididae | NC_054195.1 | *Eclipophleps carinata* |
| Acrididae | NC_046556.1 | *Emeiacris maculata* |
| Acrididae | NC_052732.1 | *Epacromius coerulipes* |
| Acrididae | NC_045237.1 | *Euchorthippus unicolor* |
| Acrididae | NC_014449.1 | *Euchorthippus fusigeniculatus* |
| Acrididae | NC_046557.1 | *Euthystira luteifemora* |

| | | |
|---|---|---|
| Acrididae | NC_053658.1 | *Fer nigripennis* |
| Acrididae | NC_011114.1 | *Gastrimargus marmoratus* |
| Acrididae | NC_046411.1 | *Gesonula punctifrons* |
| Acrididae | NC_014349.1 | *Gomphocerippus rufus* |
| Acrididae | NC_021103.1 | *Gomphocerus sibiricus* |
| Acrididae | NC_015478.1 | *Gomphocerus sibiricus tibetanus* |
| Acrididae | NC_013847.1 | *Gomphocerus licenti* |
| Acrididae | NC_029205.1 | *Gonista bicolor* |
| Acrididae | NC_046537.1 | *Heteropternis respondens* |
| Acrididae | NC_030587.1 | *Hieroglyphus tonkinensis* |
| Acrididae | NC_014891.1 | *Locusta migratoria manilensis* |
| Acrididae | NC_015624.1 | *Locusta migratoria tibetensis* |
| Acrididae | NC_001712.1 | *Locusta migratoria* |
| Acrididae | NC_011119.1 | *Locusta migratoria migratoria* |
| Acrididae | NC_036994.1 | *Longchuanacris curvifurculus* |
| Acrididae | NC_036062.1 | *Nomadacris japonica* |
| Acrididae | NC_052734.1 | *Oedaleus manjius* |
| Acrididae | NC_046538.1 | *Oedaleus abruptus* |
| Acrididae | NC_029327.1 | *Oedaleus infernalis* |
| Acrididae | NC_011115.1 | *Oedaleus decorus asiaticus* |
| Acrididae | NC_013701.1 | *Ognevia longipennis* |
| Acrididae | NC_046560.1 | *Omocestus viridulus* |
| Acrididae | NC_023467.1 | *Orinhippus tibetanus* |

| | | |
|---|---|---|
| Acrididae | NC_045883.1 | *Oxya agavisa* |
| Acrididae | NC_045928.1 | *Oxya hainanensis* |
| Acrididae | NC_043773.1 | *Oxya japonica* |
| Acrididae | NC_032076.1 | *Oxya hyla* |
| Acrididae | NC_010219.1 | *Oxya chinensis* |
| Acrididae | NC_053745.1 | *Oxytauchira flange* |
| Acrididae | NC_046570.1 | *Oxytauchira brachyptera* |
| Acrididae | NC_053660.1 | *Paratoacris reticulipennis* |
| Acrididae | NC_061318.1 | *Phlaeoba antennata* |
| Acrididae | NC_031506.1 | *Phlaeoba infumata* |
| Acrididae | NC_029150.1 | *Phlaeoba tenebrosa* |
| Acrididae | NC_011827.1 | *Phlaeoba albonema* |
| Acrididae | NC_013835.1 | *Prumna arctica* |
| Acrididae | NC_025765.1 | *Pseudoxya diminuta* |
| Acrididae | NC_035227.1 | *Pternoscirta caliginosa* |
| Acrididae | NC_064683.1 | *Schistocerca gregaria* |
| Acrididae | NC_063967.1 | *Schistocerca nitens* |
| Acrididae | NC_063966.1 | *Schistocerca cancellata* |
| Acrididae | NC_061052.1 | *Schistocerca americana* |
| Acrididae | NC_046564.1 | *Sphingonotus yenchihensis* |
| Acrididae | NC_046563.1 | *Sphingonotus ningsianus* |
| Acrididae | NC_046550.1 | *Sphingonotus menglaensis* |
| Acrididae | NC_052717.1 | *Stenocatantops mistshenkoi* |

| | | |
|---|---|---|
| Acrididae | NC_046565.1 | *Traulia orchotibialis* |
| Acrididae | NC_046551.1 | *Traulia lofaoshana* |
| Acrididae | NC_041114.1 | *Traulia nigritibialis* |
| Acrididae | NC_036063.1 | *Traulia minuta* |
| Acrididae | NC_013826.1 | *Traulia szetschuanensis* |
| Acrididae | NC_027179.1 | *Trilophidia annulata* |
| Acrididae | NC_053942.1 | *Uvaroviola multispinosa* |
| Acrididae | NC_021609.1 | *Xenocatantops brachycerus* |
| Acridomorpha | NC_014887.1 | *Acrida cinerea* |
| Acridomorpha | NC_011303.1 | *Acrida willemsei* |
| Acridomorpha | NC_046539.1 | *Parapleurus alliaceus* |
| Acridomorpha | NC_046534.1 | *Pseudoeoscyllina brevipennisoides* |
| Acridomorpha | NC_046544.1 | *Calliptamus barbarus* |
| Acridomorpha | NC_030626.1 | *Calliptamus abbreviatus* |
| Acridomorpha | NC_011305.1 | *Calliptamus italicus* |
| Acridomorpha | NC_029135.1 | *Peripolus nepalensis* |
| Acridomorpha | NC_036750.1 | *Caryanda elegans* |
| Acridomorpha | NC_030165.1 | *Caryanda sp. ZH-2016* |
| Acridomorpha | NC_041116.1 | *Choroedocus capensis* |
| Acridomorpha | NC_034673.1 | *Choroedocus violaceipes* |
| Acridomorpha | NC_046531.1 | *Shirakiacris yunkweiensis* |
| Acridomorpha | NC_021610.1 | *Shirakiacris shirakii* |
| Acridomorpha | NC_020775.1 | *Lithidiopsis carinatus* |

| | | |
|---|---|---|
| Acridomorpha | NC_031397.1 | *Curvipennis wixiensis* |
| Acridomorpha | NC_031817.1 | *Fruhstorferiola tonkinensis* |
| Acridomorpha | NC_031379.1 | *Fruhstorferiola huayinensis* |
| Acridomorpha | NC_026716.1 | *Fruhstorferiola kulinga* |
| Acridomorpha | NC_046545.1 | *Fruhstorferiola omei* |
| Acridomorpha | NC_046529.1 | *Indopodisma kingdoni* |
| Acridomorpha | NC_023920.1 | *Kingdonella bicollina* |
| Acridomorpha | NC_046530.1 | *Paratonkinacris vittifemoralis* |
| Acridomorpha | NC_046561.1 | *Pedopodisma emeiensis* |
| Acridomorpha | NC_027187.1 | *Qinlingacris taibaiensis* |
| Acridomorpha | NC_032716.1 | *Tonkinacris sinensis* |
| Acridomorpha | NC_046533.1 | *Xiangelilacris zhongdianensis* |
| Acridomorpha | NC_030586.1 | *Yunnanacris yunnaneus* |
| Acridomorpha | NC_020778.1 | *Ommexecha virens* |
| Acridomorpha | NC_023919.1 | *Pacris xizangensis* |
| Acridomorpha | NC_020776.1 | *Pyrgacris descampsi* |
| Acridomorpha | NC_046532.1 | *Spathosternum prasiniferum prasiniferum* |
| Acridomorpha | NC_020773.1 | *Tristira magellanica* |
| Acridomorpha | NC_014491.1 | *Physemacris variolosa* |
| Acridomorpha | NC_020777.1 | *Tanaocerus koebelei* |
| Acridomorpha | NC_046528.1 | *Conophymacris viridis* |
| Acridomorpha | NC_046555.1 | *Dericorys annulata* |
| Acridomorpha | NC_020774.1 | *Lentula callani* |

| | | |
|---|---|---|
| Acridomorpha | NC_057646.1 | *Melanoplus differentialis* |
| Acridomorpha | NC_014490.1 | *Xyleus modestus* |
| Acrodonta | NC_012452.1 | *Chamaeleo calcaricarens* |
| Acrodonta | NC_012445.1 | *Chamaeleo arabicus* |
| Acrodonta | NC_012444.1 | *Chamaeleo zeylanicus* |
| Acrodonta | NC_012443.1 | *Chamaeleo monachus* |
| Acrodonta | NC_012436.1 | *Chamaeleo dilepis* |
| Acrodonta | NC_012427.1 | *Chamaeleo chamaeleon* |
| Acrodonta | NC_012422.1 | *Chamaeleo africanus* |
| Acrodonta | NC_012420.1 | *Chamaeleo calyptratus* |
| Acrodonta | NC_013603.2 | *Pseudotrapelus sinaitus* |
| Acrodonta | NC_008065.1 | *Xenagama taylori* |
| Acrodonta | NC_009421.1 | *Chlamydosaurus kingii* |
| Acrodonta | NC_006922.1 | *Pogona vitticeps* |
| Acrodonta | NC_053755.1 | *Calotes mystaceus* |
| Acrodonta | NC_009683.1 | *Calotes versicolor* |
| Acrodonta | NC_056342.1 | *Diploderma micangshanensis* |
| Acrodonta | NC_039453.1 | *Pseudocalotes microlepis* |
| Acrodonta | NC_014178.1 | *Hydrosaurus amboinensis* |
| Acrodonta | NC_024179.1 | *Leiolepis reevesii* |
| Acrodonta | NC_014179.1 | *Leiolepis guttata* |
| Adephaga | NC_011329.1 | *Trachypachus holmbergi* |
| Adephaga | NC_036271.1 | *Amphizoa insolens* |

| | | |
|---|---|---|
| Adephaga | NC_057538.1 | *Hydrocolus stagnalis* |
| Adephaga | NC_057541.1 | *Hydroporus signatus* |
| Adephaga | NC_057540.1 | *Hydroporus planus* |
| Adephaga | NC_057539.1 | *Hydroporus analis* |
| Adephaga | NC_057542.1 | *Iberoporus pluto* |
| Adephaga | NC_057543.1 | *Leconectes striatellus* |
| Adephaga | NC_057544.1 | *Lioporeus pilatei* |
| Adephaga | NC_057545.1 | *Mystonectes coelamboides* |
| Adephaga | NC_057546.1 | *Nebrioporus vagrans* |
| Adephaga | NC_057547.1 | *Nectoporus subrotundus* |
| Adephaga | NC_057549.1 | *Oreodytes scitulus* |
| Adephaga | NC_057548.1 | *Oreodytes davisii* |
| Adephaga | NC_037752.1 | *Paroster microsturtensis* |
| Adephaga | NC_037751.1 | *Paroster mesosturtensis* |
| Adephaga | NC_037750.1 | *Paroster macrosturtensis* |
| Adephaga | NC_057550.1 | *Porhydrus obliquesignatus* |
| Adephaga | NC_057551.1 | *Rhithrodytes bimaculatus* |
| Adephaga | NC_057553.1 | *Scarodytes savinensis* |
| Adephaga | NC_057552.1 | *Scarodytes malickyi* |
| Adephaga | NC_030593.1 | *Hygrobia hermanni* |
| Adephaga | NC_054236.1 | *Dineutus mellyi* |
| Adephaga | NC_013249.1 | *Macrogyrus oblongus* |
| Adephaga | NC_065263.1 | *Mastax latefasciata* |

| | | |
|---|---|---|
| Adephaga | NC_038191.1 | *Cicindela anchoralis* |
| Adephaga | NC_035714.1 | *Manticora tibialis* |
| Adephaga | NC_064369.1 | *Notiophilus quadripunctatus* |
| Adephaga | NC_064368.1 | *Omophron limbatum* |
| Adephaga | NC_060736.1 | *Scarites subterraneus* |
| Adephaga | NC_036261.1 | *Tachyta nana* |
| Adephaga | NC_054235.1 | *Eretes sticticus* |
| Adephaga | NC_037749.1 | *Limbodessus palmulaoides* |
| Adephaga | NC_063608.1 | *Hyphydrus ovatus* |
| Afrotheria | NC_010304.1 | *Eremitalpa granti* |
| Afrotheria | NC_010301.1 | *Dendrohyrax dorsalis* |
| Afrotheria | NC_004919.1 | *Procavia capensis* |
| Afrotheria | NC_004026.1 | *Macroscelides proboscideus* |
| Afrotheria | NC_003314.1 | *Dugong dugon* |
| Afrotheria | NC_010302.1 | *Trichechus manatus* |
| Afrotheria | NC_002631.2 | *Echinops telfairi* |
| Afrotheria | NC_002078.1 | *Orycteropus afer* |
| Aleocharinae | NC_061260.1 | *Aleochara curtula* |
| Aleocharinae | NC_059899.1 | *Aleochara postica* |
| Aleocharinae | NC_028598.1 | *Callicerus obscurus* |
| Aleocharinae | NC_028602.1 | *Liogluta microptera* |
| Aleocharinae | NC_028600.1 | *Euryusa optabilis* |
| Aleocharinae | NC_028604.1 | *Myrmecocephalus concinnus* |

| | | |
|---|---|---|
| Aleocharinae | NC_028606.1 | *Oxypoda acuminata* |
| Aleyrodoidea | NC_029155.1 | *Aleurocanthus spiniferus* |
| Aleyrodoidea | NC_056299.1 | *Aleyrodes shizuokensis* |
| Aleyrodoidea | NC_006279.1 | *Bemisia tabaci* |
| Aleyrodoidea | NC_024056.1 | *Bemisia afer* |
| Aleyrodoidea | NC_050930.1 | *Crenidorsum turpiniae* |
| Aleyrodoidea | NC_006292.1 | *Tetraleurodes acaciae* |
| Aleyrodoidea | NC_005939.1 | *Aleurodicus dugesii* |
| Alpheoidea | NC_041151.1 | *Alpheus inopinatus* |
| Alpheoidea | NC_038116.1 | *Alpheus japonicus* |
| Alpheoidea | NC_038068.1 | *Alpheus hoplocheles* |
| Alpheoidea | NC_014883.1 | *Alpheus distinguendus* |
| Alpheoidea | NC_047307.1 | *Synalpheus microneptunus* |
| Alpheoidea | NC_045223.1 | *Lebbeus groenlandicus* |
| Alpheoidea | NC_064049.1 | *Lysmata boggessi* |
| Alpheoidea | NC_050676.1 | *Lysmata amboinensis* |
| Alpheoidea | NC_050677.1 | *Saron marmoratus* |
| Ambystoma | NC_039182.1 | *Ambystoma talpoideum* |
| Ambystoma | NC_027501.1 | *Ambystoma bishopi* |
| Ambystoma | NC_014572.1 | *Ambystoma unisexual* |
| Ambystoma | NC_014571.1 | *Ambystoma texanum* |
| Ambystoma | NC_014568.1 | *Ambystoma barbouri* |
| Ambystoma | NC_006889.1 | *Ambystoma dumerilii* |

| | | |
|---|---|---|
| Ambystoma | NC_006888.1 | *Ambystoma andersoni* |
| Ambystoma | NC_006887.1 | *Ambystoma tigrinum tigrinum* |
| Ambystoma | NC_006890.1 | *Ambystoma californiense* |
| Ambystoma | NC_006330.1 | *Ambystoma laterale* |
| Americhelydia | NC_000886.1 | *Chelonia mydas* |
| Americhelydia | NC_012398.1 | *Eretmochelys imbricata* |
| Americhelydia | NC_028634.1 | *Lepidochelys olivacea* |
| Americhelydia | NC_018550.1 | *Natator depressa* |
| Americhelydia | NC_011198.1 | *Chelydra serpentina* |
| Americhelydia | NC_009260.1 | *Macrochelys temminckii* |
| Americhelydia | NC_014577.1 | *Kinosternon leucostomum* |
| Americhelydia | NC_017607.1 | *Sternotherus carinatus* |
| Amphisbaenia | NC_006284.1 | *Amphisbaena schmidti* |
| Amphisbaenia | NC_006285.1 | *Geocalamus acutus* |
| Amphisbaenia | NC_006288.1 | *Bipes canaliculatus* |
| Amphisbaenia | NC_006286.1 | *Bipes tridactylus* |
| Amphisbaenia | NC_006287.1 | *Bipes biporus* |
| Amphisbaenia | NC_012433.1 | *Blanus cinereus* |
| Amphisbaenia | NC_006282.1 | *Rhineura floridana* |
| Amphisbaenia | NC_006283.1 | *Diplometopon zarudnyi* |
| Anareolatae | NC_014695.1 | *Phraortes illepidus* |
| Anareolatae | NC_014705.1 | *Phraortes sp.* |
| Anareolatae | NC_014673.1 | *Micadina phluctainoides* |

| | | |
|---|---|---|
| Anareolatae | NC_058255.1 | *Eurycantha calcarata* |
| Anareolatae | NC_014688.1 | *Megacrania alpheus adan* |
| Anareolatae | NC_017748.1 | *Extatosoma tiaratum* |
| Anareolatae | NC_014694.1 | *Entoria okinawaensis* |
| Anareolatae | NC_013185.1 | *Ramulus hainanense* |
| Anareolatae | NC_014702.1 | *Ramulus irregulariterdentatus* |
| Anatidae | NC_009684.1 | *Anas platyrhynchos* |
| Anatidae | NC_050973.1 | *Anas penelope* |
| Anatidae | NC_028346.1 | *Anas clypeata* |
| Anatidae | NC_024631.1 | *Anas acuta* |
| Anatidae | NC_023352.1 | *Anas falcata* |
| Anatidae | NC_022452.1 | *Anas crecca* |
| Anatidae | NC_015482.1 | *Anas formosa* |
| Anatidae | NC_039888.1 | *Anser albifrons frontalis* |
| Anatidae | NC_025654.1 | *Anser indicus* |
| Anatidae | NC_007011.1 | *Branta canadensis* |
| Anatidae | NC_012843.1 | *Cygnus atratus* |
| Anatidae | NC_027096.1 | *Cygnus olor* |
| Anatidae | NC_017604.1 | *Cygnus columbianus bewickii* |
| Anguimorpha | NC_005958.1 | *Abronia graminea* |
| Anguimorpha | NC_048977.1 | *Anguis graeca* |
| Anguimorpha | NC_048976.1 | *Anguis colchica* |
| Anguimorpha | NC_030273.1 | *Anguis cephallonica* |

| | | |
|---|---|---|
| Anguimorpha | NC_012431.1 | *Anguis fragilis* |
| Anguimorpha | NC_048956.1 | *Anguis veronensis* |
| Anguimorpha | NC_057268.1 | *Dopasia hainanensis* |
| Anguimorpha | NC_030369.1 | *Dopasia gracilis* |
| Anguimorpha | NC_008776.1 | *Heloderma suspectum* |
| Anguimorpha | NC_005959.1 | *Shinisaurus crocodilurus* |
| Anomaluromorpha | NC_038173.1 | *Anomalurus derbianus* |
| Anomaluromorpha | NC_038172.1 | *Anomalurus pelii* |
| Anomaluromorpha | NC_009056.1 | *Anomalurus sp.* |
| Anomaluromorpha | NC_038174.1 | *Anomalurus pusillus* |
| Anomaluromorpha | NC_038171.1 | *Anomalurus beecrofti* |
| Anomaluromorpha | NC_038162.1 | *Idiurus macrotis* |
| Anomopoda | NC_064987.1 | *Daphnia tibetana* |
| Anomopoda | NC_045243.1 | *Daphnia laevis* |
| Anomopoda | NC_044415.1 | *Daphnia similis* |
| Anomopoda | NC_028152.1 | *Daphnia carinata* |
| Anomopoda | NC_026914.1 | *Daphnia magna* |
| Anomopoda | NC_000844.1 | *Daphnia pulex* |
| Anopheles | NC_064612.1 | *Anopheles darlingi* |
| Anopheles | NC_064611.1 | *Anopheles coustani* |
| Anopheles | NC_064610.1 | *Anopheles nili* |
| Anopheles | NC_064608.1 | *Anopheles moucheti* |
| Anopheles | NC_064607.1 | *Anopheles marshallii* |

| | | |
|---|---|---|
| Anopheles | NC_064606.1 | *Anopheles maculipalpis* |
| Anopheles | NC_064605.1 | *Anopheles aquasalis* |
| Anopheles | NC_064604.1 | *Anopheles coluzzii* |
| Anopheles | NC_064603.1 | *Anopheles funestus* |
| Anopheles | NC_057652.1 | *Anopheles ininii* |
| Anopheles | NC_057251.1 | *Anopheles sacharovi* |
| Anopheles | NC_057101.1 | *Anopheles anthropophagus* |
| Anopheles | NC_056264.1 | *Anopheles funestus* |
| Anopheles | NC_054312.1 | *Anopheles annulipes* |
| Anopheles | NC_037789.1 | *Anopheles medialis* |
| Anopheles | NC_044647.1 | *Anopheles cruzii* |
| Anopheles | NC_039540.1 | *Anopheles aconitus* |
| Anopheles | NC_039397.1 | *Anopheles splendidus* |
| Anopheles | NC_037815.1 | *Anopheles rondoni* |
| Anopheles | NC_037805.1 | *Anopheles parvus* |
| Anopheles | NC_037829.1 | *Anopheles pseudotibiamaculatus* |
| Anopheles | NC_037827.1 | *Anopheles kompi* |
| Anopheles | NC_037824.1 | *Anopheles pristinus* |
| Anopheles | NC_037820.1 | *Anopheles lutzii* |
| Anopheles | NC_037818.1 | *Anopheles fluminensis* |
| Anopheles | NC_037817.1 | *Anopheles antunesi* |
| Anopheles | NC_037816.1 | *Anopheles guarani* |
| Anopheles | NC_037814.1 | *Anopheles galvaoi* |

| | | |
|---|---|---|
| Anopheles | NC_037813.1 | *Anopheles forattinii* |
| Anopheles | NC_037811.1 | *Anopheles nimbus* |
| Anopheles | NC_037808.1 | *Anopheles strodei* |
| Anopheles | NC_037807.1 | *Anopheles argyritarsis* |
| Anopheles | NC_037806.1 | *Anopheles arthuri* |
| Anopheles | NC_037803.1 | *Anopheles gilesi* |
| Anopheles | NC_037802.1 | *Anopheles minor* |
| Anopheles | NC_037801.1 | *Anopheles striatus* |
| Anopheles | NC_037800.1 | *Anopheles triannulatus* |
| Anopheles | NC_037799.1 | *Anopheles lanei* |
| Anopheles | NC_037798.1 | *Anopheles sawyeri* |
| Anopheles | NC_037795.1 | *Anopheles evansae* |
| Anopheles | NC_037794.1 | *Anopheles costai* |
| Anopheles | NC_037793.1 | *Anopheles oswaldoi* |
| Anopheles | NC_037792.1 | *Anopheles atacamensis atacamensis* |
| Anopheles | NC_037791.1 | *Anopheles braziliensis* |
| Anopheles | NC_037790.1 | *Anopheles peryassui* |
| Anopheles | NC_037788.1 | *Anopheles marajoara* |
| Anopheles | NC_037787.1 | *Anopheles benarrochi* |
| Anopheles | NC_037786.1 | *Anopheles rangeli* |
| Anopheles | NC_036263.1 | *Anopheles dirus* |
| Anopheles | NC_030718.1 | *Anopheles albitarsis* |
| Anopheles | NC_030717.1 | *Anopheles janconnae* |

| | | |
|---|---|---|
| Anopheles | NC_030716.1 | *Anopheles albitarsis* |
| Anopheles | NC_030715.1 | *Anopheles oryzalimnetes* |
| Anopheles | NC_030250.1 | *Anopheles laneanus* |
| Anopheles | NC_030249.1 | *Anopheles bellator* |
| Anopheles | NC_030248.1 | *Anopheles homunculus* |
| Anopheles | NC_000875.1 | *Anopheles quadrimaculatus* |
| Anopheles | NC_028223.1 | *Anopheles stephensi* |
| Anopheles | NC_028222.1 | *Anopheles punctulatus* |
| Anopheles | NC_028221.1 | *Anopheles minimus* |
| Anopheles | NC_028220.1 | *Anopheles merus* |
| Anopheles | NC_028219.1 | *Anopheles melas* |
| Anopheles | NC_028218.1 | *Anopheles maculatus* |
| Anopheles | NC_028217.1 | *Anopheles epiroticus* |
| Anopheles | NC_028216.1 | *Anopheles culicifacies* |
| Anopheles | NC_028214.1 | *Anopheles christyi* |
| Anopheles | NC_028213.1 | *Anopheles atroparvus* |
| Anopheles | NC_027502.1 | *Anopheles culicifacies* |
| Anopheles | NC_024740.1 | *Anopheles cruzii* |
| Anopheles | NC_020769.1 | *Anopheles hinesorum* |
| Anopheles | NC_020768.1 | *Anopheles cracens* |
| Anopheles | NC_020663.1 | *Anopheles deaneorum* |
| Anopheles | NC_037821.1 | *Anopheles nr. costai* |
| Anopheles | NC_056265.1 | *Anopheles rivulorum* |

| | | |
|---|---|---|
| Anopheles | NC_028016.1 | *Anopheles sinensis* |
| Anopheles | NC_037810.1 | *Anopheles goeldii* |
| Anopheles | NC_057651.1 | *Anopheles crucians* |
| Anopheles | NC_057959.1 | *Anopheles maverlius* |
| Anopheles | NC_020770.1 | *Anopheles farauti* |
| Anostraca | NC_042147.1 | *Artemia sinica* |
| Anostraca | NC_001620.1 | *Artemia franciscana* |
| Anostraca | NC_021383.1 | *Artemia tibetiana* |
| Anostraca | NC_057109.1 | *Artemia salina* |
| Anostraca | NC_050310.1 | *Eubranchipus grubii* |
| Anostraca | NC_026704.1 | *Streptocephalus sirindhornae* |
| Anostraca | NC_046688.1 | *Streptocephalus cafer* |
| Anostraca | NC_026710.1 | *Phallocryptus tserensodnomi* |
| Anura | NC_021477.1 | *Bombina lichuanensis* |
| Anura | NC_021476.1 | *Bombina microdeladigitora* |
| Anura | NC_006689.1 | *Bombina orientalis* |
| Anura | NC_009258.1 | *Bombina variegata* |
| Anura | NC_011049.1 | *Bombina maxima* |
| Anura | NC_006402.1 | *Bombina fortinuptialis* |
| Anura | NC_027072.1 | *Leiopelma hochstetteri* |
| Anura | NC_014691.1 | *Leiopelma archeyi* |
| Anura | NC_057468.1 | *Leptobrachium liui* |
| Anura | NC_031411.1 | *Leptobrachium leishanense* |

| | | |
|---|---|---|
| Anura | NC_024427.1 | *Leptobrachium boringii* |
| Anura | NC_060625.1 | *Megophrys sp.* |
| Anura | NC_056343.1 | *Oreolalax schmidti* |
| Anura | NC_049862.1 | *Oreolalax omeimontis* |
| Anura | NC_037382.1 | *Oreolalax multipunctatus* |
| Anura | NC_030627.1 | *Oreolalax lichuanensis* |
| Anura | NC_030605.1 | *Oreolalax major* |
| Anura | NC_031426.1 | *Scutiger ningshanensis* |
| Anura | NC_006688.1 | *Alytes obstetricans pertinax* |
| Anura | NC_006690.1 | *Discoglossus galganoi* |
| Anura | NC_008144.1 | *Pelobates cultripes* |
| Anura | NC_051953.1 | *Pelobates fuscus* |
| Anura | NC_037377.1 | *Scaphiopus holbrookii* |
| Anura | NC_020000.1 | *Pelodytes cf. punctatus* |
| Anura | NC_015620.1 | *Rhinophrynus dorsalis* |
| Anura | NC_015615.1 | *Hymenochirus boettgeri* |
| Anura | NC_061926.1 | *Pipa myersi* |
| Anura | NC_061925.1 | *Pipa snethlageae* |
| Anura | NC_015617.1 | *Pipa carvalhoi* |
| Anura | NC_015618.1 | *Pseudhymenochirus merlini* |
| Aphidinae | NC_011594.1 | *Acyrthosiphon pisum* |
| Aphidinae | NC_064371.1 | *Acyrthosiphon caraganae* |
| Aphidinae | NC_053819.1 | *Aphis spiraecola* |

| | | |
|---|---|---|
| Aphidinae | NC_052865.1 | *Aphis aurantii* |
| Aphidinae | NC_045236.1 | *Aphis glycines* |
| Aphidinae | NC_043903.1 | *Aphis citricidus* |
| Aphidinae | NC_039988.1 | *Aphis fabae mordvilkoi* |
| Aphidinae | NC_031387.1 | *Aphis craccivora* |
| Aphidinae | NC_024581.1 | *Aphis gossypii* |
| Aphidinae | NC_056270.1 | *Brevicoryne brassicae* |
| Aphidinae | NC_022682.1 | *Cavariella salicicola* |
| Aphidinae | NC_022727.1 | *Diuraphis noxia* |
| Aphidinae | NC_050904.1 | *Hyalopterus pruni* |
| Aphidinae | NC_045897.1 | *Indomegoura indica* |
| Aphidinae | NC_064372.1 | *Macrosiphum rosae* |
| Aphidinae | NC_063971.1 | *Macrosiphum albifrons* |
| Aphidinae | NC_029727.1 | *Myzus persicae* |
| Aphidinae | NC_057970.1 | *Neotoxoptera formosana* |
| Aphidinae | NC_062327.1 | *Rhopalosiphum rufiabdominalis* |
| Aphidinae | NC_046740.1 | *Rhopalosiphum nymphaeae* |
| Aphidinae | NC_006158.1 | *Schizaphis graminum* |
| Aphidinae | NC_024683.1 | *Sitobion avenae* |
| Aphidomorpha | NC_035316.1 | *Floraphis meitanensis* |
| Aphidomorpha | NC_036065.1 | *Melaphis rhois* |
| Aphidomorpha | NC_035313.1 | *Nurudea yanoniella* |
| Aphidomorpha | NC_035311.1 | *Nurudea ibofushi* |

| | | |
|---|---|---|
| Aphidomorpha | NC_035301.1 | *Nurudea shiraii* |
| Aphidomorpha | NC_059063.1 | *Schlechtendalia peitan* |
| Aphidomorpha | NC_033410.1 | *Mindarus keteleerifoliae* |
| Aphidomorpha | NC_024926.1 | *Cervaphis quercus* |
| Aphidomorpha | NC_054157.1 | *Eutrichosiphum pasaniae* |
| Aphidomorpha | NC_048525.1 | *Greenidea ficicola* |
| Aphidomorpha | NC_041198.1 | *Greenidea psidii* |
| Aphidomorpha | NC_054348.1 | *Mollitrichosiphum tenuicorpus* |
| Aphidomorpha | NC_053790.1 | *Stomaphis sinisalicis* |
| Aphidomorpha | NC_060838.1 | *Tuberolachnus salignus* |
| Aphidomorpha | NC_033352.1 | *Eriosoma lanigerum* |
| Aphidomorpha | NC_045103.1 | *Paracolopha morrisoni* |
| Aphidomorpha | NC_063091.1 | *Ceratovacuna keduensis* |
| Aphidomorpha | NC_044640.1 | *Pseudoregma bambucicola* |
| Aphidomorpha | NC_050942.1 | *Hamamelistes spinosus* |
| Aphidomorpha | NC_029495.1 | *Hormaphis betulae* |
| Aphidomorpha | NC_053624.1 | *Schizoneuraphis gallarum* |
| Apidae | NC_061380.1 | *Apis mellifera carnica* |
| Apidae | NC_014295.1 | *Apis cerana* |
| Apidae | NC_001566.1 | *Apis mellifera ligustica* |
| Apidae | NC_038114.1 | *Apis nigrocincta* |
| Apidae | NC_037709.1 | *Apis dorsata* |
| Apidae | NC_036235.1 | *Apis nuluensis* |

| | | |
|---|---|---|
| Apidae | NC_035883.1 | *Apis mellifera sahariensis* |
| Apidae | NC_021401.1 | *Apis florea* |
| Apidae | NC_051932.1 | *Apis mellifera* |
| Apidae | NC_057194.1 | *Amegilla calceifera* |
| Apidae | NC_051485.1 | *Habropoda radoszkowskii* |
| Apidae | NC_057952.1 | *Bombus longipennis* |
| Apidae | NC_011923.1 | *Bombus hypocrita sapporensis* |
| Apidae | NC_045178.1 | *Bombus terrestris lusitanicus* |
| Apidae | NC_045179.1 | *Bombus terrestris terrestris* |
| Apidae | NC_057192.1 | *Thyreus decorus* |
| Apidae | NC_026198.1 | *Melipona scutellaris* |
| Apidae | NC_066054.1 | *Tetragonula pagdeni* |
| Apidae | NC_060989.1 | *Tetrapedia diversipes* |
| Apidae | NC_064404.1 | *Ceratina smaragdula* |
| Apidae | NC_057193.1 | *Ceratina okinawana* |
| Apocrita | NC_025289.1 | *Orthogonalys pulchella* |
| Apocrita | NC_027830.1 | *Taeniogonalos taihorina* |
| Apodiformes | NC_033406.1 | *Amazilia brevirostris* |
| Apodiformes | NC_033405.1 | *Amazilia millerii* |
| Apodiformes | NC_033404.1 | *Amazilia rondoniae* |
| Apodiformes | NC_024156.1 | *Amazilia versicolor* |
| Apodiformes | NC_010094.1 | *Archilochus colubris* |
| Apodiformes | NC_030286.1 | *Calliphlox amethystina* |

| | | |
|---|---|---|
| Apodiformes | NC_025786.1 | *Chrysolampis mosquitus* |
| Apodiformes | NC_030287.1 | *Florisuga fusca* |
| Apodiformes | NC_027455.1 | *Florisuga mellivora* |
| Apodiformes | NC_033413.1 | *Glaucis hirsutus* |
| Apodiformes | NC_030285.1 | *Heliodoxa aurescens* |
| Apodiformes | NC_027453.1 | *Hylocharis cyanus* |
| Apodiformes | NC_033414.1 | *Lophornis magnificus* |
| Apodiformes | NC_027454.1 | *Oreotrochilus melanogaster* |
| Apodiformes | NC_030288.1 | *Phaethornis malaris* |
| Apodiformes | NC_054244.1 | *Aerodramus fuciphagus* |
| Apodiformes | NC_028545.1 | *Chaetura pelagica* |
| Apodiformes | NC_034933.1 | *Cypseloides fumigatus* |
| Apoditrysia | NC_023212.1 | *Carposina sasakii* |
| Apoditrysia | NC_032683.1 | *Monema flavescens* |
| Apoditrysia | NC_041304.1 | *Narosa nigrisigna* |
| Apoditrysia | NC_034993.1 | *Parasa consocia* |
| Apoditrysia | NC_061059.1 | *Thosea sinensis* |
| Apoditrysia | NC_047243.1 | *Phauda flammans* |
| Arachnida | NC_023450.1 | *Cryptocellus narino* |
| Arachnida | NC_023451.1 | *Pseudocellus gertschi* |
| Arachnida | NC_009985.1 | *Pseudocellus pearsei* |
| Arachnida | NC_023452.1 | *Ricinoides karschii* |
| Arachnida | NC_009984.1 | *Nothopuga sp.* |

| | | |
|---|---|---|
| Arадidae | NC_030361.1 | *Aneurus sublobatus* |
| Aradidae | NC_030360.1 | *Aneurus similis* |
| Aradidae | NC_030362.1 | *Aradus compar* |
| Aradidae | NC_030363.1 | *Libiocoris heissi* |
| Aradidae | NC_062724.1 | *Brachyrhynchus triangulus* |
| Aradidae | NC_022670.1 | *Brachyrhynchus hsiaoi* |
| Aradidae | NC_012459.1 | *Neuroctenus parus* |
| Aradidae | NC_063144.1 | *Neuroctenus yunnanensis* |
| Araneae | NC_010777.1 | *Hypochilus thorelli* |
| Araneae | NC_020323.1 | *Liphistius erawan* |
| Araneae | NC_059873.1 | *Atypus karschi* |
| Araneae | NC_020322.1 | *Phyxioschema suthepium* |
| Araneae | NC_062668.1 | *Brachypelma albiceps* |
| Araneae | NC_053738.1 | *Cyriopagopus hainanus* |
| Araneae | NC_044081.1 | *Harpactocrates apennicola* |
| Araneae | NC_044099.1 | *Parachtes romandiolae* |
| Araneae | NC_040861.1 | *Mesabolivar sp.* |
| Araneae | NC_040859.1 | *Mesabolivar sp.* |
| Araneae | NC_020324.1 | *Pholcus phalangioides* |
| Araneae | NC_042902.1 | *Loxosceles similis* |
| Archaeognatha | NC_051491.1 | *Pedetontus zhejiangensis* |
| Archaeognatha | NC_011717.1 | *Pedetontus silvestrii* |
| Archaeognatha | NC_007688.1 | *Petrobius brevistylis* |

| | | |
|---|---|---|
| Archaeognatha | NC_021384.1 | *Songmachilis xinxiangensis* |
| Archaeognatha | NC_010532.1 | *Trigoniophthalmus alternatus* |
| Archaeognatha | NC_006895.1 | *Nesomachilis australica* |
| Arctiinae | NC_057559.1 | *Arctia plantaginis* |
| Arctiinae | NC_037512.1 | *Aglaomorpha histrio* |
| Arctiinae | NC_027094.1 | *Callimorpha dominula* |
| Arctiinae | NC_065732.1 | *Asura megala* |
| Arctiinae | NC_065307.1 | *Brunia dorsalis* |
| Arctiinae | NC_037515.1 | *Paraona staudingeri* |
| Arctiinae | NC_026844.1 | *Vamuna virilis* |
| Arctiinae | NC_062185.1 | *Nyctemera adversata* |
| Arctiinae | NC_062176.1 | *Amerila alberti* |
| Arctiinae | NC_062088.1 | *Pareuchaetes insulata* |
| Arctiinae | NC_014058.1 | *Hyphantria cunea* |
| Arctiinae | NC_026692.1 | *Lemyra melli* |
| Arctiinae | NC_062183.1 | *Phragmatobia fuliginosa* |
| Arctiinae | NC_060594.1 | *Spilarctia casigneta* |
| Arctiinae | NC_050385.1 | *Spilosoma lubricipeda* |
| Arctiinae | NC_021416.1 | *Amata formosae* |
| Argasidae | NC_023340.1 | *Antricola mexicanus* |
| Argasidae | NC_060373.1 | *Carios vespertilionis* |
| Argasidae | NC_037524.1 | *Carios faini* |
| Argasidae | NC_005291.1 | *Carios capensis* |

| | | |
|---|---|---|
| Argasidae | NC_033900.1 | *Nothoaspis amazoniensis* |
| Argasidae | NC_023370.1 | *Otobius megnini* |
| Astacidea | NC_033504.1 | *Austropotamobius torrentium* |
| Astacidea | NC_026560.1 | *Austropotamobius pallipes* |
| Astacidea | NC_033509.1 | *Pacifastacus leniusculus* |
| Astacidea | NC_033507.1 | *Cambarus robustus* |
| Astacidea | NC_033510.1 | *Procambarus acutus* |
| Astacidea | NC_020021.1 | *Procambarus fallax* |
| Astacidea | NC_016926.1 | *Procambarus clarkii* |
| Astacidea | NC_028447.1 | *Procambarus alleni* |
| Astacidea | NC_033506.1 | *Cambaroides japonicus* |
| Astacidea | NC_033505.1 | *Cambaroides dauricus* |
| Astacidea | NC_016925.1 | *Cambaroides similis* |
| Astacidea | NC_025592.1 | *Enoplometopus debelius* |
| Astacidea | NC_020027.1 | *Enoplometopus occidentalis* |
| Astacidea | NC_015607.1 | *Homarus americanus* |
| Astigmata | NC_062715.1 | *Blomia tropicalis* |
| Astigmata | NC_062716.1 | *Lepidoglyphus destructor* |
| Astigmata | NC_048990.1 | *Carpoglyphus lactis* |
| Astigmata | NC_038207.1 | *Histiostoma feroniarum* |
| Athyma | NC_060740.1 | *Athyma zeroca* |
| Athyma | NC_039833.1 | *Athyma gutama* |
| Athyma | NC_039697.1 | *Athyma kanwa* |

| | | |
|---|---|---|
| Athyma | NC_039699.1 | *Athyma jina* |
| Athyma | NC_039886.1 | *Athyma libnites* |
| Athyma | NC_039883.1 | *Athyma ranga* |
| Athyma | NC_039880.1 | *Athyma fortuna* |
| Athyma | NC_039879.1 | *Athyma nefte* |
| Athyma | NC_039876.1 | *Athyma disjuncta* |
| Athyma | NC_039873.1 | *Athyma recurva* |
| Athyma | NC_039872.1 | *Athyma punctata* |
| Athyma | NC_039870.1 | *Athyma pravara* |
| Athyma | NC_024394.1 | *Athyma kasa* |
| Athyma | NC_024393.1 | *Athyma selenophora* |
| Athyma | NC_017744.1 | *Athyma sulpitia* |
| Athyma | NC_024418.1 | *Athyma opalina* |
| Athyma | NC_024395.1 | *Athyma cama* |
| Athyma | NC_024410.1 | *Athyma asura* |
| Athyma | NC_024397.1 | *Athyma perius* |
| Atkinsoniella | NC_064507.1 | *Atkinsoniella nigrita* |
| Atkinsoniella | NC_062853.1 | *Atkinsoniella xanthoabdomena* |
| Atkinsoniella | NC_062852.1 | *Atkinsoniella wui* |
| Atkinsoniella | NC_062851.1 | *Atkinsoniella warpa* |
| Atkinsoniella | NC_062850.1 | *Atkinsoniella uniguttata* |
| Atkinsoniella | NC_062849.1 | *Atkinsoniella tiani* |
| Atkinsoniella | NC_062848.1 | *Atkinsoniella thaloidea* |

| | | |
|---|---|---|
| Atkinsoniella | NC_062846.1 | *Atkinsoniella longiuscula* |
| Atkinsoniella | NC_062845.1 | *Atkinsoniella flavipenna* |
| Atkinsoniella | NC_062844.1 | *Atkinsoniella curvata* |
| Atkinsoniella | NC_062843.1 | *Atkinsoniella aurantiaca* |
| Atkinsoniella | NC_062842.1 | *Atkinsoniella yunnanana* |
| Atkinsoniella | NC_062847.1 | *Atkinsoniella thalia* |
| Atyoidea | NC_039593.1 | *Caridina indistincta* |
| Atyoidea | NC_038067.1 | *Caridina multidentata* |
| Atyoidea | NC_030219.1 | *Caridina cf. nilotica* |
| Atyoidea | NC_024751.1 | *Caridina gracilipes* |
| Atyoidea | NC_008413.1 | *Halocaridina rubra* |
| Atyoidea | NC_035412.1 | *Halocaridinides fowleri* |
| Atyoidea | NC_043865.1 | *Neocaridina heteropoda koreana* |
| Atyoidea | NC_023823.1 | *Neocaridina denticulata* |
| Atyoidea | NC_027603.1 | *Paratya australiensis* |
| Atyoidea | NC_035411.1 | *Stygiocaris stylifera* |
| Atyoidea | NC_035404.1 | *Stygiocaris lancifera* |
| Azeliinae | NC_041089.1 | *Hydrotaea chalcogaster* |
| Azeliinae | NC_042952.1 | *Hydrotaea aenescens* |
| Azeliinae | NC_047403.1 | *Hydrotaea dentipes* |
| Azeliinae | NC_042951.1 | *Hydrotaea spinigera* |
| Azeliinae | NC_037195.1 | *Hydrotaea ignava* |
| Azeliinae | NC_053670.1 | *Muscina pascuorum* |

| | | |
|---|---|---|
| Azeliinae | NC_034805.1 | *Muscina angustifrons* |
| Azeliinae | NC_026292.1 | *Muscina stabulans* |
| Azeliinae | NC_029487.1 | *Muscina levida* |
| Azeliinae | NC_042953.1 | *Synthesiomyia nudiseta* |
| Bactrocera | NC_062139.1 | *Bactrocera neohumeralis* |
| Bactrocera | NC_062138.1 | *Bactrocera frauenfeldi* |
| Bactrocera | NC_008748.1 | *Bactrocera dorsalis* |
| Bactrocera | NC_005333.1 | *Bactrocera oleae* |
| Bactrocera | NC_053983.1 | *Bactrocera thailandica* |
| Bactrocera | NC_046521.1 | *Bactrocera rubigina* |
| Bactrocera | NC_042712.1 | *Bactrocera biguttula* |
| Bactrocera | NC_038164.1 | *Bactrocera tsuneonis* |
| Bactrocera | NC_037723.1 | *Bactrocera ritsemai* |
| Bactrocera | NC_037722.1 | *Bactrocera limbifera* |
| Bactrocera | NC_028347.1 | *Bactrocera diaphora* |
| Bactrocera | NC_029468.1 | *Bactrocera umbrosa* |
| Bactrocera | NC_029467.1 | *Bactrocera melastomatos* |
| Bactrocera | NC_029466.1 | *Bactrocera latifrons* |
| Bactrocera | NC_028327.1 | *Bactrocera arecae* |
| Bactrocera | NC_027725.1 | *Bactrocera zonata* |
| Bactrocera | NC_018787.1 | *Bactrocera correcta* |
| Bactrocera | NC_014402.1 | *Bactrocera minax* |
| Bactrocera | NC_009772.1 | *Bactrocera carambolae* |

| | | |
|---|---|---|
| Bactrocera | NC_027290.1 | *Bactrocera tau* |
| Bactrocera | NC_027254.1 | *Bactrocera scutellata* |
| Balanomorpha | NC_029168.1 | *Acasta sulcata* |
| Balanomorpha | NC_024525.1 | *Amphibalanus amphitrite* |
| Balanomorpha | NC_053649.1 | *Fistulobalanus albicostatus* |
| Balanomorpha | NC_029167.1 | *Armatobalanus allium* |
| Balanomorpha | NC_024526.1 | *Striatobalanus amaryllis* |
| Balanomorpha | NC_056392.1 | *Balanus trigonus* |
| Balanomorpha | NC_056162.1 | *Megabalanus tintinnabulum* |
| Balanomorpha | NC_024636.1 | *Megabalanus ajax* |
| Balanomorpha | NC_006293.1 | *Megabalanus volcano* |
| Balanomorpha | NC_023945.1 | *Nobia grandis* |
| Balanomorpha | NC_050836.1 | *Semibalanus cariosus* |
| Balanomorpha | NC_039849.1 | *Semibalanus balanoides* |
| Bifurcata | NC_020038.1 | *Phyllodactylus unctus* |
| Bifurcata | NC_012366.1 | *Tarentola mauritanica* |
| Bifurcata | NC_053655.1 | *Teratoscincus roborowskii* |
| Bifurcata | NC_008774.1 | *Coleonyx variegatus* |
| Bifurcata | NC_033383.1 | *Eublepharis macularius* |
| Bifurcata | NC_026105.1 | *Goniurosaurus luii* |
| Bifurcata | NC_018368.1 | *Hemitheconyx caudicinctus* |
| Bifurcata | NC_035150.1 | *Aeluroscalabotes felinus* |
| Bifurcata | NC_062789.1 | *Hoplodactylus duvaucelii* |

| | | |
|---|---|---|
| Bifurcata | NC_024557.1 | *Aprasia parapulchella* |
| Blattodea | NC_059069.1 | *Cryptocercus laojunensis* |
| Blattodea | NC_059068.1 | *Cryptocercus pudacuoensis* |
| Blattodea | NC_059067.1 | *Cryptocercus weixiensis* |
| Blattodea | NC_059066.1 | *Cryptocercus changbaiensis* |
| Blattodea | NC_059065.1 | *Cryptocercus tianbaensis* |
| Blattodea | NC_059064.1 | *Cryptocercus sanchaensis* |
| Blattodea | NC_037496.1 | *Cryptocercus meridianus* |
| Blattodea | NC_030191.1 | *Cryptocercus kyebangensis* |
| Blattodea | NC_018132.1 | *Cryptocercus relictus* |
| Blattodea | NC_034842.1 | *Neostylopyga rhombifolia* |
| Blattodea | NC_039940.1 | *Periplaneta brunnea* |
| Blattodea | NC_034841.1 | *Periplaneta australasiae* |
| Blattodea | NC_016956.1 | *Periplaneta americana* |
| Blattodea | NC_006076.1 | *Periplaneta fuliginosa* |
| Blattodea | NC_030003.1 | *Shelfordella lateralis* |
| Blattodea | NC_018122.1 | *Microhodotermes viator* |
| Blattodea | NC_018120.1 | *Mastotermes darwiniensis* |
| Bombycidae | NC_003395.1 | *Bombyx mandarina* |
| Bombycidae | NC_002355.1 | *Bombyx mori* |
| Bombycidae | NC_037149.1 | *Bombyx lemeepauli* |
| Bombycidae | NC_026518.1 | *Bombyx huttoni* |
| Bombycidae | NC_038104.1 | *Ernolatia moorei* |

| | | |
|---|---|---|
| Bombycidae | NC_062709.1 | *Gunda ochracea* |
| Bombycidae | NC_038087.1 | *Ocinara albicollis* |
| Bombycidae | NC_062712.1 | *Penicillifera lactea* |
| Bombycidae | NC_062710.1 | *Penicillifera tamsi* |
| Bombycidae | NC_021962.1 | *Rondotia menciana* |
| Bombycidae | NC_045528.1 | *Rotunda rotundapex* |
| Bombycidae | NC_036484.1 | *Triuncina daii* |
| Bombycidae | NC_062711.1 | *Valvaribifidum huananense* |
| Bostrichiformia | NC_063826.1 | *Anthrenus museorum* |
| Bostrichiformia | NC_053877.1 | *Dermestes ater* |
| Bostrichiformia | NC_053876.1 | *Dermestes lardarius* |
| Bostrichiformia | NC_044851.1 | *Dermestes coarctatus* |
| Bostrichiformia | NC_044850.1 | *Dermestes frischii* |
| Bostrichiformia | NC_044849.1 | *Dermestes tessellatocollis* |
| Bostrichiformia | NC_037200.1 | *Dermestes maculatus* |
| Bostrichiformia | NC_053875.1 | *Trogoderma granarium* |
| Bostrichiformia | NC_060290.1 | *Gibbium aequinoctiale* |
| Bostrichiformia | NC_036678.1 | *Stegobium paniceum* |
| Bostrichiformia | NC_038197.1 | *Lasioderma serricorne* |
| Bostrichiformia | NC_013582.1 | *Apatides fortis* |
| Bostrichiformia | NC_042820.1 | *Rhyzopertha dominica* |
| Bovidae | NC_020674.1 | *Addax nasomaculatus* |
| Bovidae | NC_009510.1 | *Ammotragus lervia* |

| | | |
|---|---|---|
| Bovidae | NC_020678.1 | *Antidorcas marsupialis* |
| Bovidae | NC_012098.1 | *Antilope cervicapra* |
| Bovidae | NC_033873.1 | *Bison schoetensacki* |
| Bovidae | NC_027233.1 | *Bison priscus* |
| Bovidae | NC_014044.1 | *Bison bonasus* |
| Bovidae | NC_044933.1 | *Bootherium bombifrons* |
| Bovidae | NC_006380.3 | *Bos grunniens* |
| Bovidae | NC_006853.1 | *Bos taurus* |
| Bovidae | NC_005971.1 | *Bos indicus* |
| Bovidae | NC_036020.1 | *Bos frontalis* |
| Bovidae | NC_020614.1 | *Boselaphus tragocamelus* |
| Bovidae | NC_060307.1 | *Bubalus quarlesi* |
| Bovidae | NC_057438.1 | *Bubalus arnee Assam* |
| Bovidae | NC_006295.1 | *Bubalus carabanensis* |
| Bovidae | NC_020615.1 | *Bubalus depressicornis* |
| Bovidae | NC_043930.1 | *Budorcas taxicolor taxicolor* |
| Bovidae | NC_039686.1 | *Budorcas taxicolor tibetana* |
| Bovidae | NC_005044.2 | *Capra hircus* |
| Bovidae | NC_020683.1 | *Capra caucasica* |
| Bovidae | NC_020625.1 | *Capra pyrenaica* |
| Bovidae | NC_020624.1 | *Capra nubiana* |
| Bovidae | NC_020623.1 | *Capra ibex* |
| Bovidae | NC_020622.1 | *Capra falconeri* |

| | | |
|---|---|---|
| Bovidae | NC_020626.1 | *Capra sibirica* |
| Bovidae | NC_045205.1 | *Capricornis rubidus* |
| Bovidae | NC_030179.1 | *Capricornis sp.* |
| Bovidae | NC_023457.1 | *Capricornis milneedwardsii* |
| Bovidae | NC_012096.1 | *Capricornis crispus* |
| Bovidae | NC_010640.1 | *Capricornis swinhoei* |
| Bovidae | NC_020701.1 | *Dorcatragus megalotis* |
| Bovidae | NC_020702.1 | *Eudorcas rufifrons* |
| Bovidae | NC_039669.1 | *Eudorcas thomsonii* |
| Bovidae | NC_020710.1 | *Gazella subgutturosa* |
| Bovidae | NC_020709.1 | *Gazella spekei* |
| Bovidae | NC_020708.1 | *Gazella leptoceros* |
| Bovidae | NC_020706.1 | *Gazella erlangeri* |
| Bovidae | NC_020705.1 | *Gazella dorcas* |
| Bovidae | NC_020703.1 | *Gazella bennettii* |
| Bovidae | NC_020628.1 | *Hemitragus jemlahicus* |
| Bovidae | NC_020621.1 | *Hemitragus jayakari* |
| Bovidae | NC_062362.1 | *Hippotragus niger niger* |
| Bovidae | NC_062361.1 | *Hippotragus niger roosevelti* |
| Bovidae | NC_020712.1 | *Hippotragus equinus* |
| Bovidae | NC_035309.1 | *Hippotragus leucophaeus* |
| Bovidae | NC_020716.1 | *Litocranius walleri* |
| Bovidae | NC_020718.1 | *Madoqua saltiana* |

| | | |
|---|---|---|
| Bovidae | NC_020717.1 | *Madoqua kirkii SUN* |
| Bovidae | NC_042943.1 | *Myotragus balearicus* |
| Bovidae | NC_021381.1 | *Naemorhedus goral* |
| Bovidae | NC_020723.1 | *Naemorhedus griseus* |
| Bovidae | NC_020722.1 | *Naemorhedus baileyi* |
| Bovidae | NC_013751.1 | *Naemorhedus caudatus* |
| Bovidae | NC_020726.1 | *Nanger soemmerringii* |
| Bovidae | NC_020724.1 | *Nanger dama* |
| Bovidae | NC_020725.1 | *Nanger granti* |
| Bovidae | NC_020728.1 | *Neotragus moschatus* |
| Bovidae | NC_020727.1 | *Neotragus batesi* |
| Bovidae | NC_020630.1 | *Oreamnos americanus* |
| Bovidae | NC_058019.1 | *Oryx dammah* |
| Bovidae | NC_020793.1 | *Oryx beisa* |
| Bovidae | NC_016422.1 | *Oryx gazella* |
| Bovidae | NC_020733.1 | *Ourebia ourebi* |
| Bovidae | NC_020631.1 | *Ovibos moschatus* |
| Bovidae | NC_001941.1 | *Ovis aries* |
| Bovidae | NC_047196.1 | *Ovis ammon ammon* |
| Bovidae | NC_039432.1 | *Ovis dalli* |
| Bovidae | NC_039431.1 | *Ovis nivicola lydekkeri* |
| Bovidae | NC_026064.1 | *Ovis vignei* |
| Bovidae | NC_020656.1 | *Ovis ammon* |

| | | |
|---|---|---|
| Bovidae | NC_015889.1 | *Ovis canadensis* |
| Bovidae | NC_007441.1 | *Pantholops hodgsonii* |
| Bovidae | NC_020738.1 | *Procapra gutturosa* |
| Bovidae | NC_020632.1 | *Pseudois nayaur* |
| Bovidae | NC_016689.1 | *Pseudois schaeferi* |
| Bovidae | NC_020616.1 | *Pseudoryx nghetinhensis* |
| Bovidae | NC_020741.1 | *Raphicerus campestris* |
| Bovidae | NC_020789.1 | *Rupicapra pyrenaica* |
| Bovidae | NC_020746.1 | *Saiga tatarica* |
| Bovidae | NC_020617.1 | *Syncerus caffer* |
| Bovidae | NC_020618.1 | *Taurotragus derbianus* |
| Bovidae | NC_020788.1 | *Tetracerus quadricornis* |
| Bovidae | NC_038064.1 | *Tragelaphus buxtoni* |
| Bovidae | NC_020751.1 | *Tragelaphus scriptus* |
| Bovidae | NC_020750.1 | *Tragelaphus oryx* |
| Bovidae | NC_020749.1 | *Tragelaphus eurycerus* |
| Bovidae | NC_020748.1 | *Tragelaphus angasii* |
| Bovidae | NC_020620.1 | *Tragelaphus spekii* |
| Bovidae | NC_020619.1 | *Tragelaphus imberbis* |
| Bovidae | NC_020752.1 | *Tragelaphus strepsiceros* |
| Bovidae | NC_020675.1 | *Aepyceros melampus* |
| Bovidae | NC_020676.1 | *Alcelaphus buselaphus* |
| Bovidae | NC_023542.1 | *Beatragus hunteri* |

| | | |
|---|---|---|
| Bovidae | NC_020699.1 | *Connochaetes taurinus* |
| Bovidae | NC_020698.1 | *Connochaetes gnou* |
| Bovidae | NC_023543.1 | *Damaliscus lunatus* |
| Bovidae | NC_020627.1 | *Damaliscus pygargus* |
| Bovidae | NC_020735.1 | *Philantomba maxwellii* |
| Bovidae | NC_020736.1 | *Philantomba monticola* |
| Bovidae | NC_020747.1 | *Sylvicapra grimmia* |
| Bovidae | NC_020734.1 | *Pelea capreolus* |
| Bovidae | NC_020715.1 | *Kobus ellipsiprymnus* |
| Bovidae | NC_018603.1 | *Kobus leche* |
| Bovidae | NC_020794.1 | *Redunca arundinum* |
| Bovidae | NC_020742.1 | *Redunca fulvorufula* |
| Braconidae | NC_045903.1 | *Asobara japonica* |
| Braconidae | NC_064332.1 | *Euurobracon yokahamae* |
| Braconidae | NC_062620.1 | *Virgulibracon endoxylaphagus* |
| Braconidae | NC_060869.1 | *Chelonus formosanus* |
| Braconidae | NC_014278.1 | *Spathius agrili* |
| Braconidae | NC_053259.1 | *Meteorus pulchricornis* |
| Braconidae | NC_063945.1 | *Cotesia flavipes* |
| Braconidae | NC_014272.1 | *Cotesia vestalis* |
| Bucerotiformes | NC_038152.1 | *Anthracoceros coronatus* |
| Bucerotiformes | NC_038201.1 | *Buceros bicornis* |
| Bucerotiformes | NC_015201.1 | *Bycanistes brevis* |

| | | |
|---|---|---|
| Bucerotiformes | NC_039950.1 | *Rhyticeros undulatus* |
| Buprestidae | NC_064400.1 | *Agrilus ornatus* |
| Buprestidae | NC_064324.1 | *Agrilus sichuanus* |
| Buprestidae | NC_030758.1 | *Agrilus planipennis* |
| Buprestidae | NC_064326.1 | *Coraebus diminutus* |
| Buprestidae | NC_064325.1 | *Coraebus cloueti* |
| Buprestidae | NC_060321.1 | *Coraebus cavifrons* |
| Buprestidae | NC_064327.1 | *Meliboeus sinae* |
| Buprestidae | NC_064328.1 | *Sambus femoralis* |
| Buprestidae | NC_060322.1 | *Trachys variolaris* |
| Buprestidae | NC_046045.1 | *Trachys auricollis* |
| Buprestidae | NC_057197.1 | *Melanophila acuminata* |
| Buprestidae | NC_012765.1 | *Chrysochroa fulgidissima* |
| Buprestidae | NC_063127.1 | *Dicerca corrugata* |
| Buprestidae | NC_062816.1 | *Chalcophora japonica* |
| Buprestidae | NC_013580.1 | *Acmaeodera sp.* |
| Buprestidae | NC_063147.1 | *Coomaniella dentata* |
| Buprestidae | NC_063146.1 | *Coomaniella copipes* |
| Calliphoridae | NC_026996.1 | *Aldrichina grahami* |
| Calliphoridae | NC_053679.1 | *Calliphora nigribarbis* |
| Calliphoridae | NC_053677.1 | *Calliphora uralensis* |
| Calliphoridae | NC_053662.1 | *Calliphora sinensis* |
| Calliphoridae | NC_029215.1 | *Calliphora chinghaiensis* |

| | | |
|---|---|---|
| Calliphoridae | NC_028411.1 | *Calliphora vomitoria* |
| Calliphoridae | NC_019639.1 | *Calliphora vicina* |
| Calliphoridae | NC_059913.1 | *Lucilia shenyangensis* |
| Calliphoridae | NC_053672.1 | *Lucilia papuensis* |
| Calliphoridae | NC_009733.1 | *Lucilia sericata* |
| Calliphoridae | NC_029486.1 | *Lucilia coeruleiviridis* |
| Calliphoridae | NC_019637.1 | *Lucilia porphyrina* |
| Calliphoridae | NC_019573.1 | *Lucilia cuprina* |
| Calliphoridae | NC_028057.1 | *Lucilia caesar* |
| Calliphoridae | NC_028056.1 | *Lucilia illustris* |
| Calliphoridae | NC_053676.1 | *Polleniopsis mongolica* |
| Calliphoridae | NC_050875.1 | *Triceratopyga calliphoroides* |
| Callosciurinae | NC_035817.1 | *Callosciurus finlaysonii* |
| Callosciurinae | NC_035816.1 | *Callosciurus prevostii* |
| Callosciurinae | NC_035815.1 | *Callosciurus notatus* |
| Callosciurinae | NC_025550.1 | *Callosciurus erythraeus* |
| Callosciurinae | NC_030071.1 | *Callosciurus adamsi* |
| Callosciurinae | NC_048991.1 | *Dremomys everetti* |
| Callosciurinae | NC_035577.1 | *Dremomys pernyi* |
| Callosciurinae | NC_026442.1 | *Dremomys rufigenis* |
| Callosciurinae | NC_030072.1 | *Exilisciurus exilis* |
| Callosciurinae | NC_030070.1 | *Lariscus insignis* |
| Callosciurinae | NC_049034.1 | *Sundasciurus hippurus* |

| | | |
|---|---|---|
| Callosciurinae | NC_049032.1 | *Sundasciurus altitudinis* |
| Callosciurinae | NC_035813.1 | *Sundasciurus lowii* |
| Callosciurinae | NC_035812.1 | *Sundasciurus brookei* |
| Callosciurinae | NC_049033.1 | *Sundasciurus robinsoni* |
| Callosciurinae | NC_029325.1 | *Tamiops maritimus* |
| Callosciurinae | NC_026875.1 | *Tamiops swinhoei* |
| Calopterygoidea | NC_027181.1 | *Atrocalopteryx atrata* |
| Calopterygoidea | NC_036749.1 | *Atrocalopteryx melli* |
| Calopterygoidea | NC_057643.1 | *Mnais tenuis* |
| Calopterygoidea | NC_065214.1 | *Neurobasis longipes* |
| Calopterygoidea | NC_042442.1 | *Psolodesmus mandarinus* |
| Calopterygoidea | NC_023233.1 | *Vestalis melania* |
| Calopterygoidea | NC_065215.1 | *Euphaea ochracea* |
| Calopterygoidea | NC_014493.1 | *Euphaea formosa* |
| Calopterygoidea | NC_020636.1 | *Pseudolestes mirabilis* |
| Calyptratae | NC_053661.1 | *Fannia scalaris* |
| Calyptratae | NC_059850.1 | *Cephenemyia stimulator* |
| Calyptratae | NC_045881.1 | *Cephenemyia trompe* |
| Calyptratae | NC_031381.1 | *Chrysomya phaonis* |
| Calyptratae | NC_028412.1 | *Chrysomya nigripes* |
| Calyptratae | NC_025338.1 | *Chrysomya pinguis* |
| Calyptratae | NC_019635.1 | *Chrysomya saffranea* |
| Calyptratae | NC_019632.1 | *Chrysomya bezziana* |

| | | |
|---|---|---|
| Calyptratae | NC_019631.1 | *Chrysomya albiceps* |
| Calyptratae | NC_019634.1 | *Chrysomya rufifacies* |
| Calyptratae | NC_002697.1 | *Chrysomya putoria* |
| Calyptratae | NC_019633.1 | *Chrysomya megacephala* |
| Calyptratae | NC_006378.1 | *Dermatobia hominis* |
| Calyptratae | NC_042781.1 | *Gasterophilus nasalis* |
| Calyptratae | NC_042780.1 | *Gasterophilus inermis* |
| Calyptratae | NC_029834.1 | *Gasterophilus intestinalis* |
| Calyptratae | NC_029812.1 | *Gasterophilus pecorum* |
| Calyptratae | NC_042379.1 | *Gyrostigma rhinocerontis* |
| Calyptratae | NC_046479.1 | *Cephalopina titillator* |
| Calyptratae | NC_013932.1 | *Hypoderma lineatum* |
| Calyptratae | NC_059851.1 | *Oestrus ovis* |
| Calyptratae | NC_045882.1 | *Rhinoestrus usbekistanicus* |
| Calyptratae | NC_053684.1 | *Pollenia pediculata* |
| Calyptratae | NC_059870.1 | *Senotainia albifrons* |
| Calyptratae | NC_028226.1 | *Delia antiqua* |
| Calyptratae | NC_042770.1 | *Fucellia costalis* |
| Calyptratae | NC_063908.1 | *Hylemya nigrimana* |
| Calyptratae | NC_050317.1 | *Hylemya vagans DM814* |
| Calyptratae | NC_050312.1 | *Pegoplata infirma DM976* |
| Calyptratae | NC_038210.1 | *Graphomya rufitibia* |
| Calyptratae | NC_063930.1 | *Mesembrina meridiana* |

| | | |
|---|---|---|
| Calyptratae | NC_037910.1 | *Musca sorbens C065* |
| Calyptratae | NC_024855.1 | *Musca domestica* |
| Calyptratae | NC_007102.1 | *Haematobia irritans irritans* |
| Calyptratae | NC_024856.1 | *Scathophaga stercoraria* |
| Calyptratae | NC_019638.1 | *Hemipyrellia ligurriens* |
| Calyptratae | NC_002660.1 | *Cochliomyia hominivorax* |
| Calyptratae | NC_026668.1 | *Phormia regina* |
| Calyptratae | NC_019636.1 | *Protophormia terraenovae* |
| Calyptratae | NC_056374.1 | *Hamaxiella brunnescens* |
| Calyptratae | NC_019640.1 | *Rutilia goerlingiana* |
| Calyptratae | NC_062963.1 | *Halidaya aurea* |
| Calyptratae | NC_056373.1 | *Periscepsia handlirschi* |
| Calyptratae | NC_060867.1 | *Cylindromyia intermedia* |
| Calyptratae | NC_050938.1 | *Ectophasia rotundiventris* |
| Calyptratae | NC_050880.1 | *Subclytia rotundiventris* |
| Calyptratae | NC_059868.1 | *Phrosinella nasuta* |
| Calyptratae | NC_063942.1 | *Protomiltogramma cincta* |
| Calyptratae | NC_059871.1 | *Goniophyto honshuensis* |
| Calyptratae | NC_053730.1 | *Sarcophila latifrons* |
| Calyptratae | NC_059869.1 | *Sarcotachina subcylindrica* |
| Calyptratae | NC_056896.1 | *Compsilura concinnata* |
| Calyptratae | NC_044409.1 | *Exorista japonica* |
| Calyptratae | NC_039824.1 | *Exorista civilis* |

| | | |
|---|---|---|
| Calyptratae | NC_014704.1 | *Exorista sorbillans* |
| Calyptratae | NC_039963.1 | *Clemelis pullata* |
| Calyptratae | NC_018118.1 | *Elodia flavipalpis* |
| Calyptratae | NC_041657.1 | *Palesisa nudioculata* |
| Calyptratae | NC_039823.1 | *Nemorilla maculosa* |
| Calyptratae | NC_065138.1 | *Winthemia sumatrana* |
| Calyptratae | NC_063609.1 | *Lydina aenea* |
| Calyptratae | NC_063086.1 | *Peleteria iavana* |
| Canidae | NC_062617.1 | *Canis rufus* |
| Canidae | NC_062616.1 | *Canis simensis* |
| Canidae | NC_002008.4 | *Canis lupus familiaris* |
| Canidae | NC_027956.1 | *Canis anthus* |
| Canidae | NC_010340.2 | *Canis lupus chanco* |
| Canidae | NC_008093.1 | *Canis latrans* |
| Canidae | NC_024172.1 | *Chrysocyon brachyurus* |
| Canidae | NC_013445.1 | *Cuon alpinus* |
| Canidae | NC_028427.1 | *Lycaon pictus* |
| Canidae | NC_013700.1 | *Nyctereutes procyonoides* |
| Canidae | NC_036369.1 | *Otocyon megalotis* |
| Canidae | NC_053974.1 | *Speothos venaticus* |
| Canidae | NC_026723.1 | *Urocyon cinereoargenteus* |
| Canidae | NC_023958.1 | *Vulpes corsac* |
| Canidae | NC_023459.1 | *Vulpes zerda* |

| | | |
|---|---|---|
| Canidae | NC_008434.1 | *Vulpes vulpes* |
| Caniformia | NC_011124.1 | *Ailurus fulgens* |
| Caniformia | NC_009691.1 | *Ailurus fulgens styani* |
| Caniformia | NC_042596.1 | *Conepatus chinga* |
| Caniformia | NC_010497.1 | *Spilogale putorius* |
| Caniformia | NC_004029.2 | *Odobenus rosmarus rosmarus* |
| Caniformia | NC_066722.1 | *Bassaricyon gabbii* |
| Caniformia | NC_053977.1 | *Potos flavus* |
| Caniformia | NC_009126.1 | *Procyon lotor* |
| Carabinae | NC_018339.1 | *Calosoma sp.* |
| Carabinae | NC_046469.1 | *Carabus changeonleei* |
| Carabinae | NC_036507.1 | *Carabus lafossei* |
| Carabinae | NC_044759.1 | *Carabus granulatus* |
| Cardinalidae | NC_041667.1 | *Piranga rubriceps* |
| Cardinalidae | NC_041664.1 | *Piranga ludoviciana* |
| Cardinalidae | NC_041663.1 | *Piranga olivacea* |
| Cardinalidae | NC_041661.1 | *Piranga flava* |
| Cardinalidae | NC_041660.1 | *Piranga lutea* |
| Cardinalidae | NC_041659.1 | *Piranga rubra* |
| Cardinalidae | NC_041658.1 | *Piranga roseogularis* |
| Cardinalidae | NC_041665.1 | *Piranga bidentata* |
| Cardinalidae | NC_041666.1 | *Piranga erythrocephala* |
| Caridea | NC_018778.1 | *Alvinocaris chelys* |

| | | |
|---|---|---|
| Caridea | NC_051948.1 | *Chorocaris paulexa* |
| Caridea | NC_054368.1 | *Mirocaris indica* |
| Caridea | NC_021971.1 | *Nautilocaris saintlaurentae* |
| Caridea | NC_020311.1 | *Opaepele loihi* |
| Caridea | NC_027116.1 | *Rimicaris exoculata* |
| Caridea | NC_020310.1 | *Rimicaris kairei* |
| Caridea | NC_037487.1 | *Shinkaicaris leurokolos* |
| Caridea | NC_029372.1 | *Rhynchocinetes durbanensis* |
| Caridea | NC_059935.1 | *Notostomus gibbosus* |
| Caridea | NC_059714.1 | *Oplophorus spinosus* |
| Caridea | NC_040856.1 | *Bitias brevis* |
| Caridea | NC_035828.1 | *Chlorotocus crassicornis* |
| Caridea | NC_040855.1 | *Heterocarpus ensifer* |
| Carnivora | NC_053980.1 | *Atilax paludinosus* |
| Carnivora | NC_053970.1 | *Bdeogale nigripes* |
| Carnivora | NC_053966.1 | *Crossarchus platycephalus* |
| Carnivora | NC_053972.1 | *Galerella sanguinea* |
| Carnivora | NC_053978.1 | *Herpestes naso* |
| Carnivora | NC_006835.1 | *Herpestes javanicus* |
| Carnivora | NC_056368.1 | *Herpestes ichneumon* |
| Carnivora | NC_053967.1 | *Ichneumia albicauda* |
| Carnivora | NC_053963.1 | *Mungos mungo* |
| Carnivora | NC_045900.1 | *Suricata suricatta* |

| | | |
|---|---|---|
| Carnivora | NC_053971.1 | *Chrotogale owstoni* |
| Carnivora | NC_038159.1 | *Parahyaena brunnea* |
| Carnivora | NC_053964.1 | *Proteles cristata* |
| Carnivora | NC_024567.1 | *Nandinia binotata* |
| Carnivora | NC_043895.1 | *Arctictis binturong albifrons* |
| Carnivora | NC_053969.1 | *Arctictis binturong* |
| Carnivora | NC_029403.1 | *Paguma larvata* |
| Carnivora | NC_039591.1 | *Paradoxurus hermaphroditus* |
| Carnivora | NC_024569.1 | *Prionodon pardicolor* |
| Carnivora | NC_033378.1 | *Civettictis civetta* |
| Carnivora | NC_024568.1 | *Genetta servalina* |
| Carnivora | NC_053976.1 | *Viverra zibetha* |
| Carnivora | NC_025296.1 | *Viverricula indica* |
| Carnivora | NC_053961.1 | *Cryptoprocta ferox* |
| Carnivora | NC_053960.1 | *Eupleres goudotii* |
| Carnivora | NC_053959.1 | *Fossa fossana* |
| Carnivora | NC_053958.1 | *Galidia elegans* |
| Carnivora | NC_053957.1 | *Galidictis fasciata* |
| Carnivora | NC_027828.1 | *Mungotictis decemlineata* |
| Carnivora | NC_053956.1 | *Salanoia concolor* |
| Carnivora | NC_005212.1 | *Acinonyx jubatus* |
| Caudata | NC_045857.1 | *Batrachuperus sp.* |
| Caudata | NC_012430.1 | *Batrachuperus yenyuanensis* |

| | | |
|---|---|---|
| Caudata | NC_008091.1 | *Batrachuperus gorganensis* |
| Caudata | NC_008085.1 | *Batrachuperus tibetanus* |
| Caudata | NC_008083.1 | *Batrachuperus pinchonii* |
| Caudata | NC_008077.1 | *Batrachuperus londongensis* |
| Caudata | NC_008090.1 | *Batrachuperus mustersi* |
| Caudata | NC_008081.1 | *Liua tsinpaensis* |
| Caudata | NC_008078.1 | *Liua shihi* |
| Caudata | NC_026854.1 | *Onychodactylus zhaoermii* |
| Caudata | NC_026853.1 | *Onychodactylus zhangyapingi* |
| Caudata | NC_008080.1 | *Pachyhynobius shangchengensis* |
| Caudata | NC_021001.1 | *Pseudohynobius shuichengensis* |
| Caudata | NC_026698.1 | *Pseudohynobius jinfo* |
| Caudata | NC_020634.1 | *Pseudohynobius puxiongensis* |
| Caudata | NC_020635.1 | *Pseudohynobius flavomaculatus* |
| Caudata | NC_004021.1 | *Ranodon sibiricus* |
| Caudata | NC_021106.1 | *Salamandrella tridactyla* |
| Caudata | NC_008082.1 | *Salamandrella keyserlingii* |
| Caudata | NC_007446.1 | *Andrias japonicus* |
| Caudata | NC_004926.1 | *Andrias davidianus* |
| Caudata | NC_023341.1 | *Necturus beyeri* |
| Caudata | NC_023342.1 | *Proteus anguinus* |
| Caudata | NC_036927.1 | *Siren lacertina* |
| Caudata | NC_002756.1 | *Lyciasalamandra atifi* |

| | | |
|---|---|---|
| Cavilabiata | NC_042689.1 | *Acisoma panorpoides* |
| Cavilabiata | NC_026305.1 | *Brachythemis contaminata* |
| Cavilabiata | NC_025758.1 | *Hydrobasileus croceus* |
| Cavilabiata | NC_060538.1 | *Libellula quadrimaculata* |
| Cavilabiata | NC_050696.1 | *Libellula angelina* |
| Cavilabiata | NC_039410.1 | *Nannophya pygmaea* |
| Cavilabiata | NC_053718.1 | *Neurothemis fulvia* |
| Cavilabiata | NC_042732.1 | *Orthetrum melania* |
| Cavilabiata | NC_032050.1 | *Orthetrum testaceum* |
| Cavilabiata | NC_032049.1 | *Orthetrum sabina* |
| Cavilabiata | NC_032048.1 | *Orthetrum chrysis* |
| Cavilabiata | NC_032047.1 | *Orthetrum glaucum* |
| Cavilabiata | NC_056355.1 | *Pantala flavescens* |
| Cavilabiata | NC_053716.1 | *Pseudothemis zonata* |
| Cavilabiata | NC_042691.1 | *Tramea virginia* |
| Cephalophus | NC_050383.1 | *Elaphodus cephalophus cephalophus* |
| Cephalophus | NC_020685.1 | *Cephalophus adersi* |
| Cephalophus | NC_020695.1 | *Cephalophus spadix* |
| Cephalophus | NC_020694.1 | *Cephalophus silvicultor* |
| Cephalophus | NC_020693.1 | *Cephalophus rufilatus* |
| Cephalophus | NC_020692.1 | *Cephalophus ogilbyi* |
| Cephalophus | NC_020691.1 | *Cephalophus nigrifrons* |
| Cephalophus | NC_020690.1 | *Cephalophus natalensis* |

| | | |
|---|---|---|
| Cephalophus | NC_020689.1 | *Cephalophus leucogaster* |
| Cephalophus | NC_020688.1 | *Cephalophus jentinki* |
| Cephalophus | NC_020687.1 | *Cephalophus dorsalis* |
| Cephalophus | NC_020686.1 | *Cephalophus callipygus* |
| Cerambycinae | NC_044698.1 | *Nortia carinicollis* |
| Cerambycinae | NC_053714.1 | *Aromia bungii* |
| Cerambycinae | NC_023937.1 | *Massicus raddei* |
| Cerambycinae | NC_061180.1 | *Nadezhdiella cantori* |
| Cerambycinae | NC_048951.1 | *Neoplocaederus obesus* |
| Cerambycinae | NC_061182.1 | *Xoanodera maculata* |
| Cerambycinae | NC_061058.1 | *Chlorophorus annularis* |
| Cerambycinae | NC_060874.1 | *Turanoclytus namaganensis* |
| Cerambycinae | NC_030782.1 | *Xylotrechus grayii* |
| Cerambycinae | NC_061181.1 | *Allotraeus orientalis* |
| Cerambycinae | NC_051944.1 | *Epipedocera atra heimeitianniu* |
| Cerambycinae | NC_053350.1 | *Anoplistes halodendri* |
| Cerambycinae | NC_045097.1 | *Xystrocera globosa* |
| Cercopithecidae | NC_023965.1 | *Allenopithecus nigroviridis* |
| Cercopithecidae | NC_021943.1 | *Cercocebus chrysogaster* |
| Cercopithecidae | NC_028592.1 | *Cercocebus atys* |
| Cercopithecidae | NC_023964.1 | *Cercocebus torquatus* |
| Cercopithecidae | NC_056341.1 | *Cercopithecus neglectus* |
| Cercopithecidae | NC_021944.1 | *Cercopithecus albogularis* |

| | | |
|---|---|---|
| Cercopithecidae | NC_023963.1 | *Cercopithecus diana* |
| Cercopithecidae | NC_023962.1 | *Cercopithecus lhoesti* |
| Cercopithecidae | NC_023961.1 | *Cercopithecus mitis* |
| Cercopithecidae | NC_008066.1 | *Chlorocebus sabaeus* |
| Cercopithecidae | NC_034277.1 | *Chlorocebus djamdjamensis* |
| Cercopithecidae | NC_034276.1 | *Chlorocebus aethiops* |
| Cercopithecidae | NC_024933.1 | *Chlorocebus cynosuros* |
| Cercopithecidae | NC_007009.1 | *Chlorocebus aethiops* |
| Cercopithecidae | NC_009748.1 | *Chlorocebus tantalus* |
| Cercopithecidae | NC_006901.1 | *Colobus guereza* |
| Cercopithecidae | NC_021947.1 | *Erythrocebus patas* |
| Cercopithecidae | NC_021954.1 | *Lophocebus aterrimus* |
| Cercopithecidae | NC_028442.1 | *Mandrillus leucophaeus* |
| Cercopithecidae | NC_021956.1 | *Mandrillus sphinx* |
| Cercopithecidae | NC_008216.1 | *Nasalis larvatus* |
| Cercopithecidae | NC_061413.1 | *Papio hamadryas hamadryas* |
| Cercopithecidae | NC_020010.2 | *Papio ursinus north* |
| Cercopithecidae | NC_020008.2 | *Papio kindae* |
| Cercopithecidae | NC_020007.2 | *Papio cynocephalus north* |
| Cercopithecidae | NC_008219.1 | *Piliocolobus badius* |
| Cercopithecidae | NC_008217.1 | *Presbytis melalophos* |
| Cercopithecidae | NC_020666.1 | *Procolobus verus* |
| Cercopithecidae | NC_018063.1 | *Pygathrix cinerea 2* |

| | | |
|---|---|---|
| Cercopithecidae | NC_018062.1 | *Pygathrix cinerea 1* |
| Cercopithecidae | NC_018061.1 | *Pygathrix nigripes* |
| Cercopithecidae | NC_008220.1 | *Pygathrix nemaeus* |
| Cercopithecidae | NC_015486.1 | *Rhinopithecus bieti* |
| Cercopithecidae | NC_008218.1 | *Rhinopithecus roxellana* |
| Cercopithecidae | NC_018060.1 | *Rhinopithecus bieti 2* |
| Cercopithecidae | NC_018057.1 | *Rhinopithecus brelichi* |
| Cercopithecidae | NC_015485.1 | *Rhinopithecus avunculus* |
| Cercopithecidae | NC_057099.1 | *Semnopithecus schistaceus* |
| Cercopithecidae | NC_008215.1 | *Semnopithecus entellus* |
| Cercopithecidae | NC_020667.1 | *Simias concolor* |
| Cercopithecidae | NC_019802.1 | *Theropithecus gelada* |
| Cervus | NC_050863.1 | *Cervus canadensis* |
| Cervus | NC_016178.1 | *Cervus nippon kopschi* |
| Cervus | NC_007704.2 | *Cervus elaphus* |
| Cervus | NC_013836.1 | *Cervus elaphus xanthopygus* |
| Cervus | NC_007179.1 | *Cervus nippon yakushimae* |
| Cervus | NC_006993.1 | *Cervus nippon centralis* |
| Cervus | NC_006973.1 | *Cervus nippon yesoensis* |
| Cervus | NC_013840.1 | *Cervus elaphus yarkandensis* |
| Cervus | NC_013834.1 | *Cervus nippon hortulorum* |
| Cervus | NC_008462.1 | *Cervus nippon taiouanus* |
| Cetacea | NC_005271.1 | *Balaenoptera acutorostrata* |

| | | |
|---|---|---|
| Cetacea | NC_007937.1 | *Balaenoptera omurai* |
| Cetacea | NC_007938.1 | *Balaenoptera edeni* |
| Cetacea | NC_006928.1 | *Balaenoptera brydei* |
| Cetacea | NC_006926.1 | *Balaenoptera bonaerensis* |
| Cetacea | NC_006929.1 | *Balaenoptera borealis* |
| Cetacea | NC_001601.1 | *Balaenoptera musculus* |
| Cetacea | NC_001321.1 | *Balaenoptera physalus* |
| Cetacea | NC_005268.1 | *Balaena mysticetus* |
| Cetacea | NC_037444.1 | *Eubalaena glacialis* |
| Cetacea | NC_006931.1 | *Eubalaena japonica* |
| Cetacea | NC_006927.1 | *Megaptera novaeangliae* |
| Cetacea | NC_005270.1 | *Eschrichtius robustus* |
| Cetacea | NC_005269.1 | *Caperea marginata* |
| Charadriiformes | NC_003712.2 | *Arenaria interpres* |
| Charadriiformes | NC_060627.1 | *Calidris alba* |
| Charadriiformes | NC_060626.1 | *Calidris alpina* |
| Charadriiformes | NC_046877.1 | *Calidris pugnax* |
| Charadriiformes | NC_046764.1 | *Calidris tenuirostris* |
| Charadriiformes | NC_040990.1 | *Calidris ruficollis* |
| Charadriiformes | NC_060628.1 | *Calidris subminuta* |
| Charadriiformes | NC_056775.1 | *Chroicocephalus saundersi* |
| Charadriiformes | NC_050864.1 | *Chroicocephalus brunnicephalus* |
| Charadriiformes | NC_052812.1 | *Chroicocephalus maculipennis* |

| | | |
|---|---|---|
| Charadriiformes | NC_034741.1 | *Gallinago stenura* |
| Charadriiformes | NC_036344.1 | *Gelochelidon nilotica* |
| Charadriiformes | NC_023777.1 | *Ichthyaetus relictus* |
| Charadriiformes | NC_029383.1 | *Larus vegae* |
| Charadriiformes | NC_025556.1 | *Larus crassirostris* |
| Charadriiformes | NC_007006.1 | *Larus dominicanus* |
| Charadriiformes | NC_046765.1 | *Limosa lapponica* |
| Charadriiformes | NC_053923.1 | *Limosa haemastica* |
| Charadriiformes | NC_061764.1 | *Numenius minutus* |
| Charadriiformes | NC_059854.1 | *Numenius madagascariensis* |
| Charadriiformes | NC_042233.1 | *Numenius tenuirostris* |
| Charadriiformes | NC_030507.1 | *Numenius phaeopus* |
| Charadriiformes | NC_031347.1 | *Pinguinus impennis* |
| Charadriiformes | NC_062960.1 | *Prosobonia parvirostris* |
| Charadriiformes | NC_025521.1 | *Scolopax rusticola* |
| Charadriiformes | NC_045282.1 | *Sterna paradisaea* |
| Charadriiformes | NC_036345.1 | *Sterna hirundo* |
| Charadriiformes | NC_028176.1 | *Sternula albifrons* |
| Charadriiformes | NC_056273.1 | *Tringa stagnatilis* |
| Charadriiformes | NC_044651.1 | *Tringa nebularia* |
| Charadriiformes | NC_044648.1 | *Tringa totanus* |
| Charadriiformes | NC_039096.1 | *Tringa glareola* |
| Charadriiformes | NC_036016.1 | *Tringa semipalmata* |

| | | |
|---|---|---|
| Charadriiformes | NC_033974.1 | *Tringa ochropus* |
| Charadriiformes | NC_030585.1 | *Tringa erythropus* |
| Charadriiformes | NC_033973.1 | *Xenus cinereus* |
| Charadriiformes | NC_045517.1 | *Aethia cristatella* |
| Charadriiformes | NC_029328.1 | *Synthliboramphus wumizusume* |
| Charadriiformes | NC_007978.1 | *Synthliboramphus antiquus* |
| Charadriiformes | NC_052794.1 | *Burhinus bistriatus* |
| Charadriiformes | NC_061228.1 | *Charadrius dubius* |
| Charadriiformes | NC_059881.1 | *Charadrius mongolus* |
| Charadriiformes | NC_041118.1 | *Charadrius alexandrinus* |
| Charadriiformes | NC_052771.1 | *Charadrius vociferus* |
| Charadriiformes | NC_056257.1 | *Pluvialis squatarola* |
| Charadriiformes | NC_033966.1 | *Pluvialis fulva* |
| Charadriiformes | NC_025514.1 | *Vanellus cinereus* |
| Charadriiformes | NC_034237.1 | *Haematopus ostralegus* |
| Charadriiformes | NC_003713.2 | *Haematopus ater* |
| Charadriiformes | NC_041576.1 | *Hydrophasianus chirurgus* |
| Charadriiformes | NC_024068.1 | *Jacana spinosa* |
| Charadriiformes | NC_027420.1 | *Recurvirostra avosetta* |
| Charadriiformes | NC_026125.1 | *Stercorarius maccormicki* |
| Charadriiformes | NC_052780.1 | *Stercorarius parasiticus* |
| Charadriiformes | NC_059848.1 | *Turnix tanki* |
| Charadriiformes | NC_052815.1 | *Turnix velox* |

| | | |
|---|---|---|
| Chelodina | NC_041292.1 | *Chelodina steindachneri* |
| Chelodina | NC_041291.1 | *Chelodina colliei* |
| Chelodina | NC_041286.1 | *Chelodina canni* |
| Chelodina | NC_041285.1 | *Chelodina burrungandjii* |
| Chelodina | NC_041284.1 | *Chelodina novaeguineae* |
| Chelodina | NC_037387.1 | *Chelodina oblonga* |
| Chelodina | NC_037386.1 | *Chelodina pritchardi* |
| Chelodina | NC_037385.1 | *Chelodina parkeri* |
| Chelodina | NC_037383.1 | *Chelodina expansa* |
| Chelodina | NC_024667.1 | *Chelodina longicollis* |
| Chelodina | NC_037384.1 | *Chelodina mccordi* |
| Chelonoidis | NC_051482.1 | *Chelonoidis microphyes* |
| Chelonoidis | NC_051474.1 | *Chelonoidis darwini* |
| Chelonoidis | NC_051472.1 | *Chelonoidis abingdonii* |
| Chelonoidis | NC_051475.1 | *Chelonoidis donfaustoi* |
| Cherax | NC_026227.1 | *Cherax boesemani* |
| Cherax | NC_026224.1 | *Cherax holthuisi* |
| Cherax | NC_030514.1 | *Cherax sp. HMG-2016* |
| Cherax | NC_026226.1 | *Cherax bicarinatus* |
| Cherax | NC_026559.1 | *Cherax tenuimanus* |
| Cherax | NC_022936.1 | *Cherax cainii* |
| Cherax | NC_022938.1 | *Cherax monticola* |
| Cherax | NC_023480.1 | *Cherax dispar* |

| | | |
|---|---|---|
| Cherax | NC_023479.1 | *Cherax quinquecarinatus* |
| Cherax | NC_023478.1 | *Cherax robustus* |
| Cherax | NC_022939.1 | *Cherax glaber* |
| Cherax | NC_022937.1 | *Cherax quadricarinatus* |
| Cherax | NC_011243.1 | *Cherax destructor* |
| Chironominae | NC_063909.1 | *Axarus fungorum* |
| Chironominae | NC_061966.1 | *Chironomus nipponensis* |
| Chironominae | NC_016167.1 | *Chironomus tepperi* |
| Chironominae | NC_061965.1 | *Glyptotendipes tokunagai* |
| Chironominae | NC_066802.1 | *Microtendipes umbrosus* |
| Chironominae | NC_066803.1 | *Polypedilum henicurum* |
| Chironominae | NC_061259.1 | *Polypedilum nubifer* |
| Chironominae | NC_028015.1 | *Polypedilum vanderplanki* |
| Chironominae | NC_061974.1 | *Stenochironomus zhengi* |
| Chironominae | NC_061972.1 | *Stenochironomus okialbus* |
| Chironominae | NC_061971.1 | *Stenochironomus gibbus* |
| Chrysomus | NC_018813.1 | *Chrysomus cyanopus* |
| Chrysomus | NC_018807.1 | *Chrysomus thilius* |
| Chrysomus | NC_018799.1 | *Chrysomus icterocephalus* |
| Chrysomus | NC_018796.1 | *Chrysomus ruficapillus* |
| Cicadettinae | NC_060717.1 | *Karenia caelatata Karenia Caelatata* |
| Cicadettinae | NC_053870.1 | *Cicadetta abscondita* |
| Cicadettinae | NC_041656.1 | *Magicicada tredecula* |

| | | |
|---|---|---|
| Cicadettinae | NC_041654.1 | *Magicicada tredecassini* |
| Cicadettinae | NC_041652.1 | *Magicicada tredecim* |
| Cicadettinae | NC_041651.1 | *Magicicada neotredecim* |
| Cicadettinae | NC_041653.1 | *Magicicada cassini* |
| Cimicomorpha | NC_027144.1 | *Adelphocoris nigritylus* |
| Cimicomorpha | NC_027143.1 | *Adelphocoris lineolatus* |
| Cimicomorpha | NC_023796.1 | *Adelphocoris fasciaticollis* |
| Cimicomorpha | NC_023083.1 | *Apolygus lucorum* |
| Cimicomorpha | NC_030257.1 | *Creontiades dilutus* |
| Cimicomorpha | NC_037926.1 | *Lygus pratensis* |
| Cimicomorpha | NC_021975.1 | *Lygus lineolaris* |
| Cimicomorpha | NC_065139.1 | *Orius strigicollis* |
| Cimicomorpha | NC_024583.1 | *Orius sauteri* |
| Cimicomorpha | NC_012429.1 | *Orius niger* |
| Cimicomorpha | NC_042679.1 | *Tetraphleps aterrimus* |
| Cimicomorpha | NC_019595.1 | *Gorpis annulatus* |
| Cimicomorpha | NC_037371.1 | *Nabicula flavomarginata* |
| Cimicomorpha | NC_050256.1 | *Nabis ferus* |
| Cimicomorpha | NC_019594.1 | *Nabis apicalis* |
| Cimicomorpha | NC_016432.1 | *Alloeorhynchus bakeri* |
| Cimicomorpha | NC_061220.1 | *Halticus minutus* |
| Cingulata | NC_028559.1 | *Cabassous unicinctus* |
| Cingulata | NC_028557.1 | *Cabassous chacoensis* |

| | | |
|---|---|---|
| Cingulata | NC_028556.1 | *Cabassous centralis* |
| Cingulata | NC_028558.1 | *Cabassous tatouay* |
| Cingulata | NC_028560.1 | *Calyptophractus retusus* |
| Cingulata | NC_028562.1 | *Chaetophractus villosus* |
| Cingulata | NC_028561.1 | *Chaetophractus vellerosus* |
| Cingulata | NC_028563.1 | *Chlamyphorus truncatus* |
| Cingulata | NC_028571.1 | *Euphractus sexcinctus* |
| Cingulata | NC_028573.1 | *Priodontes maximus* |
| Cingulata | NC_028576.1 | *Tolypeutes tricinctus* |
| Cingulata | NC_028575.1 | *Tolypeutes matacus* |
| Cingulata | NC_028577.1 | *Zaedyus pichiy* |
| Cingulata | NC_001821.1 | *Dasypus novemcinctus* |
| Cingulata | NC_028568.1 | *Dasypus sabanicola* |
| Cingulata | NC_028566.1 | *Dasypus kappleri* |
| Cingulata | NC_028565.1 | *Dasypus hybridus* |
| Cingulata | NC_028569.1 | *Dasypus septemcinctus* |
| Cingulata | NC_028567.1 | *Dasypus pilosus* |
| Cingulata | NC_028570.1 | *Dasypus yepesi* |
| Coccinellinae | NC_042417.1 | *Aiolocaria hexaspilota* |
| Coccinellinae | NC_036272.1 | *Anatis ocellata* |
| Coccinellinae | NC_046481.1 | *Hippodamia variegata* |
| Coccinellinae | NC_066052.1 | *Illeis koebelei* |
| Coccinellinae | NC_061950.1 | *Illeis bistigmosa* |

| | | |
|---|---|---|
| Coccinellinae | NC_064320.1 | *Megalocaria dilatata* |
| Coccinellinae | NC_058236.1 | *Oenopia sauzeti* |
| Coccinellinae | NC_066406.1 | *Vibidia duodecimguttata* |
| Coenagrionoidea | NC_054209.1 | *Ceriagrion fallax* |
| Coenagrionoidea | NC_063728.1 | *Ischnura asiatica* |
| Coenagrionoidea | NC_031824.1 | *Ischnura elegans* |
| Coenagrionoidea | NC_021617.1 | *Ischnura pumilio* |
| Coenagrionoidea | NC_027180.1 | *Platycnemis foliacea* |
| Coenagrionoidea | NC_031823.1 | *Megaloprepus caerulatus* |
| Coleoptera | NC_011328.1 | *Tetraphalerus bruchi* |
| Coleoptera | NC_012144.1 | *Hydroscapha granulum* |
| Coleoptera | NC_011322.1 | *Sphaerius sp. BT0074* |
| Coleoptera | NC_036265.1 | *Incoltorrida madagassica* |
| Colubridae | NC_063502.1 | *Elaphe moellendorffi* |
| Colubridae | NC_027605.1 | *Elaphe schrenckii* |
| Colubridae | NC_025643.1 | *Elaphe davidi* |
| Colubridae | NC_024743.1 | *Elaphe bimaculata* |
| Colubridae | NC_012770.1 | *Elaphe poryphyracea* |
| Colubridae | NC_041068.1 | *Elaphe dione E27* |
| Colubridae | NC_024546.1 | *Euprepiophis perlacea* |
| Colubridae | NC_057467.1 | *Gonyosoma frenatum* |
| Colubridae | NC_046046.1 | *Lycodon ruhstrati* |
| Colubridae | NC_028730.1 | *Lycodon flavozonatus* |

| | | |
|---|---|---|
| Colubridae | NC_024559.1 | *Lycodon rufozonatus* |
| Colubridae | NC_052835.1 | *Oligodon chinensis* |
| Colubridae | NC_026083.1 | *Oligodon ningshaanensis* |
| Colubridae | NC_022146.1 | *Oocatochus rufodorsatus* |
| Colubridae | NC_049067.1 | *Orientocoluber spinalis* |
| Colubridae | NC_009769.1 | *Pantherophis slowinskii* |
| Colubridae | NC_030041.1 | *Ptyas mucosa* |
| Colubridae | NC_028049.1 | *Ptyas dhumnades* |
| Colubridae | NC_062677.1 | *Calamaria septentrionalis* |
| Colubridae | NC_015793.1 | *Nerodia sipedon* |
| Colubridae | NC_046823.1 | *Opisthotropis latouchii* |
| Colubridae | NC_056274.1 | *Opisthotropis guangxiensis* |
| Colubridae | NC_056928.1 | *Pseudagkistrodon rudis* |
| Colubridae | NC_030210.1 | *Rhabdophis tigrinus* |
| Colubridae | NC_063962.1 | *Trimerodytes annularis* |
| Colubridae | NC_053908.1 | *Pseudoxenodon stejnegeri* |
| Colubridae | NC_022430.1 | *Sibynophis chinensis* |
| Colubroidea | NC_013981.1 | *Pseudoleptodeira latifasciata* |
| Colubroidea | NC_012816.1 | *Thermophis zhaoermii* |
| Colubroidea | NC_035713.1 | *Thermophis baileyi* |
| Colubroidea | NC_035058.1 | *Thermophis shangrila* |
| Colubroidea | NC_060375.1 | *Myanophis thanlyinensis* |
| Colubroidea | NC_062614.1 | *Aipysurus eydouxii* |

| | | |
|---|---|---|
| Colubroidea | NC_046794.1 | *Hydrophis curtus* |
| Colubroidea | NC_046795.1 | *Hydrophis cyanocinctus* |
| Colubroidea | NC_058002.1 | *Pareas formosensis* |
| Colubroidea | NC_057676.1 | *Pareas stanleyi* |
| Colubroidea | NC_050894.1 | *Pareas boulengeri* |
| Colubroidea | NC_032085.1 | *Achalinus rufescens* |
| Colubroidea | NC_032084.1 | *Achalinus spinalis* |
| Colubroidea | NC_011576.1 | *Achalinus meiguensis* |
| Colubroidea | NC_011393.1 | *Bungarus fasciatus* |
| Colubroidea | NC_011389.1 | *Naja atra* |
| Colubroidea | NC_011394.1 | *Ophiophagus hannah* |
| Colubroidea | NC_054255.1 | *Sinomicrurus macclellandi* |
| Colubroidea | NC_036055.1 | *Laticauda semifasciata* |
| Colubroidea | NC_036054.1 | *Laticauda colubrina* |
| Colubroidea | NC_036053.1 | *Laticauda laticaudata* |
| Colubroidea | NC_061027.1 | *Psammophis lineolatus* |
| Coptotermes | NC_037018.1 | *Coptotermes suzhouensis* |
| Coptotermes | NC_030021.1 | *Coptotermes travians* |
| Coptotermes | NC_030020.1 | *Coptotermes sjoestedti* |
| Coptotermes | NC_030017.1 | *Coptotermes michaelseni* |
| Coptotermes | NC_030016.1 | *Coptotermes kalshoveni* |
| Coptotermes | NC_030015.1 | *Coptotermes heimi* |
| Coptotermes | NC_030014.1 | *Coptotermes gestroi* |

| | | |
|---|---|---|
| Coptotermes | NC_030013.1 | *Coptotermes frenchi* |
| Coptotermes | NC_030012.1 | *Coptotermes elisae* |
| Coptotermes | NC_030011.1 | *Coptotermes amanii* |
| Coptotermes | NC_028722.1 | *Coptotermes testaceus* |
| Coptotermes | NC_018125.1 | *Coptotermes lacteus* |
| Coptotermes | NC_015800.1 | *Coptotermes formosanus* |
| Coptotermes | NC_030018.1 | *Coptotermes remotus* |
| Coptotermes | NC_030019.1 | *Coptotermes sepangensis* |
| Coraciiformes | NC_035868.1 | *Alcedo atthis* |
| Coraciiformes | NC_035746.1 | *Halcyon smyrnensis* |
| Coraciiformes | NC_024198.1 | *Halcyon pileata* |
| Coraciiformes | NC_028177.1 | *Halcyon coromanda* |
| Coraciiformes | NC_052813.1 | *Halcyon senegalensis* |
| Coraciiformes | NC_011712.1 | *Todiramphus sanctus vagans* |
| Coraciiformes | NC_052810.1 | *Brachypteracias leptosomus* |
| Coraciiformes | NC_024280.1 | *Ceryle rudis* |
| Coraciiformes | NC_052793.1 | *Chloroceryle aenea* |
| Coraciiformes | NC_035658.1 | *Megaceryle lugubris* |
| Coraciiformes | NC_011716.2 | *Eurystomus orientalis* |
| Coraciiformes | NC_052798.1 | *Eurystomus gularis* |
| Coraciiformes | NC_034642.1 | *Merops viridis* |
| Coraciiformes | NC_052799.1 | *Todus mexicanus* |
| Coreidae | NC_035509.1 | *Anoplocnemis curvipes* |

| | | |
|---|---|---|
| Coreidae | NC_063143.1 | *Cletomorpha raja* |
| Coreidae | NC_053874.1 | *Cletus rubidiventris* |
| Coreidae | NC_050997.1 | *Cletus punctiger* |
| Coreidae | NC_042806.1 | *Cloresmus pulchellus* |
| Coreidae | NC_046833.1 | *Enoplops potanini* |
| Coreidae | NC_042809.1 | *Leptoglossus membranaceus* |
| Coreidae | NC_042811.1 | *Mictis tenebrosa* |
| Coreidae | NC_042807.1 | *Molipteryx lunata* |
| Coreidae | NC_065112.1 | *Notobitus montanus* |
| Coreidae | NC_037376.1 | *Notopteryx soror* |
| Coreidae | NC_042814.1 | *Pseudomictis brevicornis* |
| Coronuloidea | NC_033393.1 | *Epopella plicata* |
| Coronuloidea | NC_029169.1 | *Chelonibia testudinaria* |
| Coronuloidea | NC_037241.1 | *Tesseropora rosea* |
| Coronuloidea | NC_056911.1 | *Tetraclita kuroshioensis* |
| Coronuloidea | NC_037398.1 | *Tetraclita rufotincta* |
| Coronuloidea | NC_029154.1 | *Tetraclita serrata* |
| Coronuloidea | NC_008974.1 | *Tetraclita japonica* |
| Corvoidea | NC_051467.1 | *Aphelocoma coerulescens* |
| Corvoidea | NC_041197.1 | *Artamus cinereus* |
| Corvoidea | NC_053090.1 | *Cnemophilus loriae* |
| Corvoidea | NC_015824.1 | *Cyanopica cyanus* |
| Corvoidea | NC_045376.1 | *Dendrocitta formosae* |

| | | |
|---|---|---|
| Corvoidea | NC_015810.1 | *Garrulus glandarius* |
| Corvoidea | NC_051558.1 | *Nucifraga caryocatactes* |
| Corvoidea | NC_022839.1 | *Nucifraga columbiana* |
| Corvoidea | NC_020424.1 | *Oriolus chinensis* |
| Corvoidea | NC_014879.1 | *Podoces hendersoni* |
| Corvoidea | NC_025927.1 | *Pyrrhocorax graculus* |
| Corvoidea | NC_037486.1 | *Urocissa caerulea* |
| Corvoidea | NC_020426.1 | *Urocissa erythrorhyncha* |
| Corvoidea | NC_031350.1 | *Callaeas cinereus* |
| Corvoidea | NC_031351.1 | *Heteralocha acutirostris* |
| Corvoidea | NC_029143.1 | *Philesturnus carunculatus* |
| Corvoidea | NC_051017.1 | *Edolisoma coerulescens* |
| Corvoidea | NC_024257.1 | *Pericrocotus ethologus* |
| Corvoidea | NC_052013.1 | *Cinclosoma castanotum* |
| Corvoidea | NC_052012.1 | *Cinclosoma clarum* |
| Corvoidea | NC_052011.1 | *Cinclosoma punctatum* |
| Corvoidea | NC_053092.1 | *Ifrita kowaldi* |
| Corvoidea | NC_050688.1 | *Ptilorrhoa caerulescens* |
| Corvoidea | NC_051006.1 | *Ptilorrhoa leucosticta* |
| Corvoidea | NC_053087.1 | *Struthidea cinerea* |
| Corvoidea | NC_056389.1 | *Dicrurus macrocercus* |
| Corvoidea | NC_051028.1 | *Dicrurus megarhynchus* |
| Corvoidea | NC_043948.1 | *Dicrurus hottentottus* |

| | | |
|---|---|---|
| Corvoidea | NC_053070.1 | *Erythrocercus mccallii* |
| Corvoidea | NC_053094.1 | *Machaerirhynchus nigripectus* |
| Corvoidea | NC_053109.1 | *Chloropsis hardwickii* |
| Corvoidea | NC_053068.1 | *Chloropsis cyanopogon* |
| Corvoidea | NC_051018.1 | *Irena cyanogastra* |
| Corvoidea | NC_051030.1 | *Lanius ludovicianus* |
| Corvoidea | NC_039818.1 | *Lanius tigrinus* |
| Corvoidea | NC_030604.1 | *Lanius schach* |
| Corvoidea | NC_028333.1 | *Lanius cristatus* |
| Corvoidea | NC_027655.1 | *Lanius isabellinus* |
| Corvoidea | NC_053067.1 | *Dryoscopus gambensis* |
| Corvoidea | NC_059916.1 | *Grallina cyanoleuca* |
| Corvoidea | NC_051022.1 | *Myiagra hebetior* |
| Corvoidea | NC_032725.1 | *Terpsiphone atrocaudata* |
| Corvoidea | NC_053086.1 | *Orthonyx spaldingii* |
| Corvoidea | NC_053088.1 | *Aleadryas rufinucha* |
| Corvoidea | NC_053096.1 | *Daphoenositta chrysoptera* |
| Corvoidea | NC_053091.1 | *Eulacestoma nigropectus* |
| Corvoidea | NC_053104.1 | *Falcunculus frontatus* |
| Corvoidea | NC_051019.1 | *Pachycephala philippinensis* |
| Corvoidea | NC_040956.1 | *Rhagologus leucostigma* |
| Corvoidea | NC_028336.1 | *Turnagra capensis* |
| Corvoidea | NC_053056.1 | *Paradisaea raggiana* |

| | | |
|---|---|---|
| Corvoidea | NC_051016.1 | *Vireo altiloquus* |
| Corvoidea | NC_024869.1 | *Vireo olivaceus* |
| Corvus | NC_062298.1 | *Corvus cornix cornix* |
| Corvus | NC_046029.1 | *Corvus dauuricus* |
| Corvus | NC_045381.1 | *Corvus pectoralis* |
| Corvus | NC_034839.1 | *Corvus cryptoleucus* |
| Corvus | NC_034838.1 | *Corvus corax* |
| Corvus | NC_035877.1 | *Corvus coronoides* |
| Corvus | NC_031518.1 | *Corvus moriorum* |
| Corvus | NC_027173.1 | *Corvus macrorhynchos* |
| Corvus | NC_026783.1 | *Corvus hawaiiensis* |
| Corvus | NC_026461.1 | *Corvus brachyrhynchos* |
| Corvus | NC_024607.1 | *Corvus splendens* |
| Corvus | NC_002069.2 | *Corvus frugilegus* |
| Corvus | NC_051471.1 | *Corvus moneduloides* |
| Cossoidea | NC_023936.1 | *Eogystia hippophaecolus* |
| Cossoidea | NC_051865.1 | *Chalcidica minea* |
| Cossoidea | NC_062621.1 | *Endoxyla cinereus* |
| Cossoidea | NC_051866.1 | *Zeuzera multistrigata* |
| Crambidae | NC_061248.1 | *Botyodes principalis* |
| Crambidae | NC_060868.1 | *Cnaphalocrocis patnalis* |
| Crambidae | NC_015985.1 | *Cnaphalocrocis medinalis* |
| Crambidae | NC_042150.1 | *Cydalima perspectalis* |

| | | |
|---|---|---|
| Crambidae | NC_025933.1 | *Glyphodes pyloalis* |
| Crambidae | NC_022699.1 | *Glyphodes quadrimaculalis* |
| Crambidae | NC_027174.1 | *Loxostege sticticalis* |
| Crambidae | NC_024283.1 | *Maruca testulalis* |
| Crambidae | NC_024099.1 | *Maruca vitrata* |
| Crambidae | NC_025764.1 | *Nomophila noctuella* |
| Crambidae | NC_039177.1 | *Omiodes indicata* |
| Crambidae | NC_066444.1 | *Omphisa fuscidentalis* |
| Crambidae | NC_059846.1 | *Ostrinia kasmirica* |
| Crambidae | NC_056248.1 | *Ostrinia furnacalis* |
| Crambidae | NC_054270.1 | *Ostrinia nubilalis* |
| Crambidae | NC_048888.1 | *Ostrinia zealis* |
| Crambidae | NC_048887.1 | *Ostrinia scapulalis* |
| Crambidae | NC_039632.1 | *Palpita hypohomalia* |
| Crambidae | NC_033540.1 | *Pycnarmon lactiferalis* |
| Crambidae | NC_066087.1 | *Pygospila tyres* |
| Crambidae | NC_046050.1 | *Pyrausta despicata* |
| Crambidae | NC_061243.1 | *Sinomphisa plagialis* |
| Crambidae | NC_062118.1 | *Sitochroa verticalis* |
| Crambidae | NC_027443.1 | *Spoladea recurvalis* |
| Crambidae | NC_061245.1 | *Syllepte taiwanalis* |
| Crambidae | NC_030510.1 | *Tyspanodes striata* |
| Crambidae | NC_025569.1 | *Tyspanodes hypsalis* |

| | | |
|---|---|---|
| Crambidae | NC_029716.1 | *Chilo sacchariphagus* |
| Crambidae | NC_015612.1 | *Chilo suppressalis* |
| Crambidae | NC_061606.1 | *Crambus perlellus* |
| Crambidae | NC_013274.1 | *Diatraea saccharalis* |
| Crambidae | NC_029751.1 | *Pseudargyria interruptella* |
| Crambidae | NC_030509.1 | *Evergestis junctalis* |
| Crambidae | NC_050323.1 | *Cataclysta lemnata* |
| Crambidae | NC_021756.1 | *Elophila interruptalis* |
| Crambidae | NC_020094.1 | *Paracymoriza prodigalis* |
| Crambidae | NC_023471.1 | *Paracymoriza distinctalis* |
| Crambidae | NC_031151.1 | *Parapoynx crisonalis* |
| Crambidae | NC_056800.1 | *Heortia vitessoides* |
| Crangonyctoidea | NC_046509.1 | *Bactrurus brachycaudus* |
| Crangonyctoidea | NC_046510.1 | *Stygobromus pizzinii* |
| Crangonyctoidea | NC_030261.1 | *Stygobromus indentatus* |
| Crangonyctoidea | NC_046511.1 | *Stygobromus allegheniensis* |
| Cricetidae | NC_065750.1 | *Chionomys nivalis* |
| Cricetidae | NC_065749.1 | *Chionomys roberti* |
| Cricetidae | NC_056331.1 | *Cricetulus sokolovi* |
| Cricetidae | NC_007936.1 | *Cricetulus griseus* |
| Cricetidae | NC_053822.1 | *Cricetulus barabensis* |
| Cricetidae | NC_031802.1 | *Cricetulus migratorius* |
| Cricetidae | NC_025330.1 | *Cricetulus longicaudatus* |

| | | |
|---|---|---|
| Cricetidae | NC_024592.1 | *Cricetulus kamensis* |
| Cricetidae | NC_034646.1 | *Dicrostonyx torquatus* |
| Cricetidae | NC_034313.1 | *Dicrostonyx groenlandicus* |
| Cricetidae | NC_034307.1 | *Dicrostonyx hudsonius* |
| Cricetidae | NC_056388.1 | *Dinaromys bogdanovi* |
| Cricetidae | NC_054160.1 | *Ellobius talpinus* |
| Cricetidae | NC_065401.1 | *Eothenomys eleusis* |
| Cricetidae | NC_030330.1 | *Eothenomys miletus* |
| Cricetidae | NC_027418.1 | *Eothenomys melanogaster* |
| Cricetidae | NC_013571.1 | *Eothenomys chinensis* |
| Cricetidae | NC_035612.1 | *Habromys ixtlani* |
| Cricetidae | NC_035599.1 | *Isthmomys pirrensis* |
| Cricetidae | NC_057534.1 | *Lasiopodomys brandtii* |
| Cricetidae | NC_025283.1 | *Lasiopodomys mandarinus* |
| Cricetidae | NC_013276.1 | *Mesocricetus auratus* |
| Cricetidae | NC_029477.1 | *Myodes rufocanus* |
| Cricetidae | NC_024538.1 | *Myodes glareolus* |
| Cricetidae | NC_016427.1 | *Myodes regulus* |
| Cricetidae | NC_040138.1 | *Neodon fuscus* |
| Cricetidae | NC_035503.1 | *Neodon sikimensis* |
| Cricetidae | NC_016055.1 | *Neodon irene* |
| Cricetidae | NC_039670.1 | *Neotoma magister* |
| Cricetidae | NC_035594.1 | *Neotoma mexicana* |

| | | |
|---|---|---|
| Cricetidae | NC_033356.1 | *Neotoma fuscipes* |
| Cricetidae | NC_035615.1 | *Neotomodon alstoni* |
| Cricetidae | NC_036035.1 | *Ondatra zibethicus* |
| Cricetidae | NC_029760.1 | *Onychomys leucogaster* |
| Cricetidae | NC_047188.1 | *Peromyscus eremicus* |
| Cricetidae | NC_039921.1 | *Peromyscus maniculatus bairdii* |
| Cricetidae | NC_037180.1 | *Peromyscus leucopus* |
| Cricetidae | NC_035614.1 | *Peromyscus crinitus* |
| Cricetidae | NC_035613.1 | *Peromyscus megalops* |
| Cricetidae | NC_035571.1 | *Peromyscus polionotus* |
| Cricetidae | NC_035598.1 | *Peromyscus pectoralis* |
| Cricetidae | NC_035596.1 | *Peromyscus aztecus* |
| Cricetidae | NC_035593.1 | *Peromyscus attwateri* |
| Cricetidae | NC_042823.1 | *Phodopus sungorus* |
| Cricetidae | NC_031809.1 | *Phodopus roborovskii* |
| Cricetidae | NC_035595.1 | *Podomys floridanus* |
| Cricetidae | NC_013563.1 | *Proedromys liangshanensis* |
| Cricetidae | NC_049036.1 | *Prometheomys schaposchnikowi* |
| Cricetidae | NC_035597.1 | *Reithrodontomys mexicanus* |
| Cricetidae | NC_013068.1 | *Tscherskia triton* |
| Crocidura | NC_056768.1 | *Crocidura russula* |
| Crocidura | NC_056167.1 | *Crocidura dongyangjiangensis* |
| Crocidura | NC_046831.1 | *Crocidura tanakae* |

| | | |
|---|---|---|
| Crocidura | NC_042762.1 | *Crocidura fuliginosa* |
| Crocidura | NC_029329.1 | *Crocidura lasiura* |
| Crocidura | NC_026204.2 | *Crocidura attenuata* |
| Crocidura | NC_027249.1 | *Crocidura beatus* |
| Crocidura | NC_027248.1 | *Crocidura mindorus* |
| Crocidura | NC_027247.1 | *Crocidura grayi* |
| Crocidura | NC_027246.1 | *Crocidura panayensis* |
| Crocidura | NC_027245.1 | *Crocidura negrina* |
| Crocidura | NC_027243.1 | *Crocidura palawanensis* |
| Crocidura | NC_027242.1 | *Crocidura orientalis* |
| Crocidura | NC_027244.1 | *Crocidura ninoyi* |
| Crocodylia | NC_001922.1 | *Alligator mississippiensis* |
| Crocodylia | NC_002744.2 | *Caiman crocodilus* |
| Crocodylia | NC_009732.1 | *Paleosuchus trigonatus* |
| Crocodylia | NC_009729.1 | *Paleosuchus palpebrosus* |
| Crocodylia | NC_010639.1 | *Mecistops cataphractus* |
| Crocodylia | NC_009728.1 | *Osteolaemus tetraspis* |
| Crocodylia | NC_008241.1 | *Gavialis gangeticus* |
| Crocodylia | NC_011074.1 | *Tomistoma schlegelii* |
| Crocodylus | NC_008143.1 | *Crocodylus porosus* |
| Crocodylus | NC_015238.2 | *Crocodylus johnsoni* |
| Crocodylus | NC_024513.1 | *Crocodylus rhombifer* |
| Crocodylus | NC_015651.1 | *Crocodylus novaeguineae* |

| | | |
|---|---|---|
| Crocodylus | NC_015648.1 | *Crocodylus intermedius* |
| Crocodylus | NC_015647.1 | *Crocodylus acutus* |
| Crocodylus | NC_015235.1 | *Crocodylus moreletii* |
| Crocodylus | NC_014706.1 | *Crocodylus palustris* |
| Crocodylus | NC_014670.1 | *Crocodylus mindorensis* |
| Crocodylus | NC_008795.1 | *Crocodylus siamensis* |
| Crocodylus | NC_008142.1 | *Crocodylus niloticus* |
| Cryptodira | NC_054232.1 | *Amyda cartilaginea* |
| Cryptodira | NC_021371.1 | *Apalone spinifera* |
| Cryptodira | NC_014054.1 | *Apalone ferox* |
| Cryptodira | NC_053617.1 | *Chitra vandijki* |
| Cryptodira | NC_026028.1 | *Chitra indica* |
| Cryptodira | NC_002780.1 | *Dogania subplana* |
| Cryptodira | NC_025494.1 | *Lissemys scutata* |
| Cryptodira | NC_012414.1 | *Lissemys punctata* |
| Cryptodira | NC_039559.1 | *Nilssonia nigricans* |
| Cryptodira | NC_013841.1 | *Palea steindachneri* |
| Cryptodira | NC_015825.1 | *Pelochelys cantorii* |
| Cryptodira | NC_017901.1 | *Rafetus swinhoei* |
| Cryptodira | NC_012833.1 | *Trionyx triunguis* |
| Cryptorhynchinae | NC_050893.1 | *Trigonopterus porg* |
| Cryptorhynchinae | NC_050892.1 | *Trigonopterus kotamobagensis* |
| Cryptorhynchinae | NC_050890.1 | *Trigonopterus singkawangensis* |

| | | |
|---|---|---|
| Cryptorhynchinae | NC_050889.1 | *Trigonopterus triradiatus* |
| Cryptorhynchinae | NC_050887.1 | *Trigonopterus tanimbarensis* |
| Cryptorhynchinae | NC_050886.1 | *Trigonopterus selaruensis* |
| Cryptorhynchinae | NC_050888.1 | *Trigonopterus jasminae* |
| Cryptorhynchinae | NC_026719.1 | *Eucryptorrhynchus chinensis* |
| Cucujiformia | NC_061195.1 | *Basiprionota bisignata* |
| Cucujiformia | NC_053935.1 | *Brontispa longissima* |
| Cucujiformia | NC_042198.1 | *Callispa bowringii* |
| Cucujiformia | NC_065028.1 | *Aoria nigripes* |
| Cucujiformia | NC_052915.1 | *Chrysolina aeruginosa* |
| Cucujiformia | NC_050933.1 | *Chrysomela vigintipunctata* |
| Cucujiformia | NC_036108.1 | *Gastrolina thoracica* |
| Cucujiformia | NC_045247.1 | *Gastrophysa polygoni* |
| Cucujiformia | NC_042500.1 | *Gonioctena intermedia* |
| Cucujiformia | NC_052916.1 | *Plagiodera versicolora* |
| Cucujiformia | NC_003372.1 | *Crioceris duodecimpunctata* |
| Cucujiformia | NC_057259.1 | *Physosmaragdina nigrifrons* |
| Cucujiformia | NC_054189.1 | *Basilepta fulvipes* |
| Cucujiformia | NC_057218.1 | *Colasposoma dauricum* |
| Cucujiformia | NC_033532.1 | *Paleosepharia posticata* |
| Cucujiformia | NC_041423.1 | *Podagricomela nigricollis* |
| Cucujiformia | NC_042828.1 | *Oomorphoides metallicus* |
| Cucujiformia | NC_048496.1 | *Paulhutchinsonia pilosicollis* |

| | | |
|---|---|---|
| Cucujiformia | NC_029515.1 | *Spiniphilus spinicornis* |
| Cucujiformia | NC_036267.1 | *Byturus ochraceus BMNH 677381* |
| Cucujiformia | NC_011324.1 | *Chaetosoma scaritides* |
| Cucujiformia | NC_044896.1 | *Idgia oculata* |
| Cucujiformia | NC_024271.1 | *Dastarcus helophoroides* |
| Cucujiformia | NC_035677.1 | *Acanthoscelides obtectus* |
| Cucujiformia | NC_047198.1 | *Bruchidius uberatus* |
| Cucujiformia | NC_053358.1 | *Callosobruchus maculatus* |
| Cucujiformia | NC_050294.1 | *Caryopemon giganteus* |
| Cucujiformia | NC_052914.1 | *Spondylis buprestoides* |
| Cucujiformia | NC_028332.1 | *Agasicles hygrophila* |
| Cucujiformia | NC_045929.1 | *Argopistes tsekooni* |
| Cucujiformia | NC_041169.1 | *Macrohaltica subplicata* |
| Cucujiformia | NC_045901.1 | *Phyllotreta striolata* |
| Cucujiformia | NC_053362.1 | *Psylliodes chlorophana* |
| Cucujiformia | NC_038090.1 | *Anastrangalia sequensi* |
| Cucujiformia | NC_033340.1 | *Trichodes sinae* |
| Cucujiformia | NC_062857.1 | *Tenebroides mauritanicus* |
| Cucujiformia | NC_041172.1 | *Henosepilachna vigintioctopunctata* |
| Cucujiformia | NC_023469.1 | *Henosepilachna pusillanima* |
| Cucujiformia | NC_064321.1 | *Epiverta chelonia* |
| Cucujiformia | NC_050855.1 | *Nephus oblongosignatus* |
| Cucujoidea | NC_051939.1 | *Cucujus kempi* |

| | | |
|---|---|---|
| Cucujoidea | NC_051938.1 | *Cucujus mniszechi* |
| Cucujoidea | NC_018350.1 | *Cucujus clavipes* |
| Cucujoidea | NC_051934.1 | *Palaestes abruptus* |
| Cucujoidea | NC_051937.1 | *Platisus moerosus* |
| Cucujoidea | NC_051936.1 | *Platisus angusticollis* |
| Cucujoidea | NC_051935.1 | *Thesaurus albertalleni* |
| Cucujoidea | NC_028204.1 | *Cryptolestes pusillus* |
| Cucujoidea | NC_028203.1 | *Cryptolestes ferrugineus* |
| Cucujoidea | NC_051933.1 | *Hymaea magna* |
| Cucujoidea | NC_011326.1 | *Priasilpha obscura* |
| Cucujoidea | NC_056919.1 | *Aulacochilus grouvellei* |
| Cucujoidea | NC_036266.1 | *Monotoma quadricollis* |
| Cucujoidea | NC_046036.1 | *Carpophilus dimidiatus* |
| Cucujoidea | NC_046035.1 | *Carpophilus pilosellus* |
| Cucujoidea | NC_036104.1 | *Aethina tumida* |
| Cucujoidea | NC_050852.1 | *Omosita colon* |
| Cucujoidea | NC_036273.1 | *Silvanus bidentatus* |
| Cuculiformes | NC_052811.1 | *Centropus unirufus* |
| Cuculiformes | NC_050049.1 | *Centropus bengalensis* |
| Cuculiformes | NC_052804.1 | *Piaya cayana* |
| Cuculiformes | NC_052800.1 | *Crotophaga sulcirostris* |
| Cuculiformes | NC_052776.1 | *Ceuthmochares aereus* |
| Cuculiformes | NC_028414.1 | *Cuculus poliocephalus* |

| | | |
|---|---|---|
| Cuculiformes | NC_060520.1 | *Eudynamys scolopaceus* |
| Cuculiformes | NC_011709.1 | *Eudynamys taitensis* |
| Culex | NC_065809.1 | *Culex vishnui* |
| Culex | NC_054323.1 | *Culex annulirostris* |
| Culex | NC_054318.1 | *Culex sitiens* |
| Culex | NC_054317.1 | *Culex orbostiensis* |
| Culex | NC_054316.1 | *Culex fergusoni* |
| Culex | NC_054315.1 | *Culex cylindricus* |
| Culex | NC_054314.1 | *Culex australicus* |
| Culex | NC_038160.1 | *Culex gelidus* |
| Culex | NC_037823.1 | *Culex nigripalpus* |
| Culex | NC_037828.1 | *Culex brami* |
| Culex | NC_037826.1 | *Culex chidesteri* |
| Culex | NC_037825.1 | *Culex lygrus* |
| Culex | NC_037822.1 | *Culex declarator* |
| Culex | NC_037819.1 | *Culex bilineatus* |
| Culex | NC_037812.1 | *Culex mollis* |
| Culex | NC_037797.1 | *Culex surinamensis* |
| Culex | NC_028616.1 | *Culex tritaeniorhynchus* |
| Culex | NC_015079.1 | *Culex pipiens pipiens* |
| Culex | NC_014574.1 | *Culex quinquefasciatus* |
| Culicinae | NC_065121.1 | *Aedes vexans* |
| Culicinae | NC_006817.1 | *Aedes albopictus* |

| | | |
|---|---|---|
| Culicinae | NC_054325.1 | *Aedes alternans* |
| Culicinae | NC_054320.1 | *Aedes rubrithorax* |
| Culicinae | NC_054319.1 | *Aedes alboannulatus* |
| Culicinae | NC_050044.1 | *Aedes flavopictus* |
| Culicinae | NC_046946.1 | *Aedes koreicus* |
| Culicinae | NC_035159.1 | *Aedes aegypti* |
| Culicinae | NC_025473.1 | *Aedes notoscriptus* |
| Culicinae | NC_057214.1 | *Haemagogus tropicalis* |
| Culicinae | NC_057213.1 | *Haemagogus spegazzinii* |
| Culicinae | NC_057212.1 | *Haemagogus leucocelaenus* |
| Culicinae | NC_057211.1 | *Haemagogus albomaculatus* |
| Culicinae | NC_028025.1 | *Haemagogus janthinomys* |
| Culicinae | NC_044656.1 | *Limatus flavisetosus* |
| Culicinae | NC_060639.1 | *Ochlerotatus fluviatilis* |
| Culicinae | NC_054326.1 | *Ochlerotatus vittiger* |
| Culicinae | NC_054321.1 | *Ochlerotatus nigrithorax* |
| Culicinae | NC_027494.1 | *Ochlerotatus vigilax* |
| Culicinae | NC_060642.1 | *Psorophora ferox* |
| Culicinae | NC_060641.1 | *Psorophora albipes* |
| Culicinae | NC_044658.1 | *Psorophora saeva* |
| Culicinae | NC_044659.1 | *Runchomyia reversa* |
| Culicinae | NC_044660.1 | *Sabethes undosus* |
| Culicinae | NC_037500.1 | *Sabethes glaucodaemon* |

| | | |
|---|---|---|
| Culicinae | NC_037499.1 | *Sabethes chloropterus* |
| Culicinae | NC_037498.1 | *Sabethes belisarioi* |
| Culicinae | NC_044661.1 | *Trichoprosopon pallidiventer* |
| Culicinae | NC_054322.1 | *Tripteroides tasmaniensis* |
| Culicinae | NC_044663.1 | *Wyeomyia confusa* |
| Culicinae | NC_054327.1 | *Culiseta inconspicua* |
| Culicinae | NC_060640.1 | *Coquillettidia nigricans* |
| Culicinae | NC_054313.1 | *Coquillettidia linealis* |
| Culicinae | NC_044655.1 | *Coquillettidia chrysonotum* |
| Culicinae | NC_047479.1 | *Mansonia uniformis* |
| Culicinae | NC_044657.1 | *Mansonia amazonensis* |
| Culicinae | NC_044662.1 | *Uranotaenia geometrica* |
| Culicomorpha | NC_029354.1 | *Dixella aestivalis* |
| Culicomorpha | NC_037796.1 | *Bironella hollandi* |
| Culicomorpha | NC_062634.1 | *Chagasia sp.* |
| Culicomorpha | NC_054324.1 | *Toxorhynchites speciosus* |
| Darevskia | NC_045934.1 | *Darevskia valentini* |
| Darevskia | NC_050005.1 | *Darevskia brauneri* |
| Darevskia | NC_050003.1 | *Darevskia daghestanica* |
| Darevskia | NC_050002.1 | *Darevskia clarkorum* |
| Darevskia | NC_050000.1 | *Darevskia praticola* |
| Darevskia | NC_049999.1 | *Darevskia caucasica* |
| Darevskia | NC_049998.1 | *Darevskia derjugini* |

| | | |
|---|---|---|
| Darevskia | NC_046012.1 | *Darevskia saxicola* |
| Darevskia | NC_046011.1 | *Darevskia rudis* |
| Darevskia | NC_046010.1 | *Darevskia portschinskii* |
| Darevskia | NC_046009.1 | *Darevskia parvula* |
| Darevskia | NC_046008.1 | *Darevskia mixta* |
| Darevskia | NC_050004.1 | *Darevskia chlorogaster* |
| Dermacentor | NC_062070.1 | *Dermacentor niveus* |
| Dermacentor | NC_062069.1 | *Dermacentor marginatus* |
| Dermacentor | NC_062068.1 | *Dermacentor steini* |
| Dermacentor | NC_061217.1 | *Dermacentor variabilis* |
| Dermacentor | NC_059724.1 | *Dermacentor auratus* |
| Dermacentor | NC_042764.1 | *Dermacentor everestianus* |
| Dermacentor | NC_023349.1 | *Dermacentor nitens Deni* |
| Dermacentor | NC_026552.1 | *Dermacentor silvarum* |
| Dermacentor | NC_028528.1 | *Dermacentor nuttalli* |
| Dermacentor | NC_062165.1 | *Dermacentor sinicus A40* |
| Dermacentor | NC_061057.1 | *Dermacentor andersoni* |
| Didelphidae | NC_057519.1 | *Didelphis pernigra* |
| Didelphidae | NC_057517.1 | *Didelphis imperfecta* |
| Didelphidae | NC_057516.1 | *Didelphis aurita* |
| Didelphidae | NC_057515.1 | *Didelphis albiventris* |
| Didelphidae | NC_001610.1 | *Didelphis virginiana* |
| Didelphidae | NC_057518.1 | *Didelphis marsupialis* |

| | | |
|---|---|---|
| Didelphidae | NC_054268.1 | *Gracilinanus agilis* |
| Didelphidae | NC_057520.1 | *Lutreolina crassicaudata* |
| Didelphidae | NC_006516.1 | *Metachirus nudicaudatus* |
| Didelphidae | NC_006299.1 | *Monodelphis domestica* |
| Didelphidae | NC_005825.1 | *Thylamys elegans* |
| Didelphidae | NC_029381.1 | *Tlacuatzin canescens* |
| Dionycha | NC_041120.1 | *Cheliceroides longipalpis* |
| Dionycha | NC_024878.1 | *Selenops bursarius* |
| Dionycha | NC_025557.1 | *Oxytate striatipes* |
| Dionycha | NC_061408.1 | *Plator insolens* |
| Dionycha | NC_057201.1 | *Phanuelus gladstone* |
| Dionycha | NC_027492.1 | *Carrhotus xanthogramma* |
| Dionycha | NC_060328.1 | *Phintella cavaleriei* |
| Dionycha | NC_061918.1 | *Asemonea sichuanensis* |
| Dionycha | NC_005942.1 | *Habronattus oregonensis* |
| Dionycha | NC_042829.1 | *Epeus alboguttatus* |
| Dionycha | NC_024877.1 | *Plexippus paykulli* |
| Dionycha | NC_024287.1 | *Telamonia vlijmi* |
| Dipodidae | NC_027499.1 | *Dipus sagitta* |
| Dipodidae | NC_027692.1 | *Stylodipus telum* |
| Dipodidae | NC_027500.1 | *Euchoreutes naso* |
| Dipodidae | NC_027579.1 | *Sicista concolor* |
| Dipodidae | NC_063779.1 | *Sicista strandi* |

| | | |
|---|---|---|
| Dipodidae | NC_027578.1 | *Eozapus setchuanus* |
| Diprotodontia | NC_008145.1 | *Distoechurus pennatus* |
| Diprotodontia | NC_008136.1 | *Lagorchestes hirsutus* |
| Diprotodontia | NC_008447.1 | *Lagostrophus fasciatus* |
| Diprotodontia | NC_039717.1 | *Macropus fuliginosus* |
| Diprotodontia | NC_001794.1 | *Macropus robustus* |
| Diprotodontia | NC_008134.1 | *Dactylopsila trivirgata* |
| Diprotodontia | NC_053898.1 | *Gymnobelideus leadbeateri* |
| Diprotodontia | NC_008135.1 | *Petaurus breviceps* |
| Diprotodontia | NC_056259.1 | *Phalanger orientalis* |
| Diprotodontia | NC_008137.1 | *Phalanger vestitus* |
| Diprotodontia | NC_003039.1 | *Trichosurus vulpecula* |
| Diprotodontia | NC_008133.1 | *Phascolarctos cinereus* |
| Diprotodontia | NC_006524.1 | *Potorous tridactylus* |
| Diprotodontia | NC_006519.1 | *Pseudocheirus peregrinus* |
| Diprotodontia | NC_006518.1 | *Tarsipes rostratus* |
| Diprotodontia | NC_003322.1 | *Vombatus ursinus* |
| Ditrysia | NC_029811.1 | *Mesophleps albilinella* |
| Ditrysia | NC_065403.1 | *Pectinophora gossypiella* |
| Ditrysia | NC_041123.1 | *Sitotroga cerealella* |
| Ditrysia | NC_029810.1 | *Dichomeris ustalella* |
| Ditrysia | NC_029844.1 | *Helcystogramma macroscopa* |
| Ditrysia | NC_057501.1 | *Phthorimaea operculella* |

| | | |
|---|---|---|
| Ditrysia | NC_029386.1 | *Tecia solanivora* |
| Ditrysia | NC_050874.1 | *Tuta absoluta* |
| Ditrysia | NC_048471.1 | *Scythris sinensis* |
| Ditrysia | NC_028168.1 | *Atrijuglans hetaohei* |
| Ditrysia | NC_046832.1 | *Eudarcia gwangneungensis* |
| Ditrysia | NC_058014.1 | *Meleonoma mirabilis* |
| Ditrysia | NC_063695.1 | *Issikiopteryx taipingensis* |
| Ditrysia | NC_053695.1 | *Casmara patrona* |
| Ditrysia | NC_026697.1 | *Promalactis suzukiella* |
| Ditrysia | NC_058015.1 | *Ripeacma umbellata* |
| Ditrysia | NC_031831.1 | *Stathmopoda auriferella* |
| Ditrysia | NC_046600.1 | *Caloptilia theivora* |
| Ditrysia | NC_061172.1 | *Conopomorpha sinensis* |
| Ditrysia | NC_061539.1 | *Phyllonorycter ringoniella* |
| Ditrysia | NC_057979.1 | *Corythoxestis sunosei* |
| Ditrysia | NC_053760.1 | *Phyllocnistis citrella* |
| Ditrysia | NC_047459.1 | *Dahlica ochrostigma* |
| Ditrysia | NC_057469.1 | *Acanthopsyche nigraplaga* |
| Dolichopodidae | NC_051891.1 | *Argyra pingwuensis* |
| Dolichopodidae | NC_053651.1 | *Asyndetus clavipes* |
| Dolichopodidae | NC_063610.1 | *Dolichopus ungulatus* |
| Dolichopodidae | NC_065147.1 | *Gymnopternus congruens* |
| Dolichopodidae | NC_046942.1 | *Hercostomus brevipilosus* |

| | | |
|---|---|---|
| Dolichopodidae | NC_060353.1 | *Lichtwardtia dentalis* |
| Dolichopodidae | NC_053650.1 | *Neurigona zhejiangensis* |
| Dolichopodidae | NC_053652.1 | *Xanthochlorus tibetensis* |
| Emberiza | NC_062087.1 | *Emberiza schoeniclus* |
| Emberiza | NC_037692.1 | *Emberiza leucocephalos* |
| Emberiza | NC_033338.1 | *Emberiza fucata* |
| Emberiza | NC_030368.1 | *Emberiza elegans* |
| Emberiza | NC_027251.1 | *Emberiza jankowskii* |
| Emberiza | NC_024925.1 | *Emberiza rutila* |
| Emberiza | NC_024924.1 | *Emberiza rustica* |
| Emberiza | NC_021445.1 | *Emberiza spodocephala* |
| Emberiza | NC_024524.1 | *Emberiza cioides* |
| Emberiza | NC_021408.1 | *Emberiza pusilla* |
| Emberiza | NC_015234.1 | *Emberiza tristrami* |
| Emberiza | NC_015233.1 | *Emberiza chrysophrys* |
| Empidoidea | NC_059903.1 | *Calohilara tibetensis* |
| Empidoidea | NC_053939.1 | *Empis separata* |
| Empidoidea | NC_064390.1 | *Hybos grossipes* |
| Empidoidea | NC_051551.1 | *Anthalia sp.* |
| Endopterygota | NC_044741.1 | *Bittacus strigosus* |
| Endopterygota | NC_015118.1 | *Bittacus pilicornis* |
| Endopterygota | NC_013180.1 | *Neopanorpa pulchra* |
| Endopterygota | NC_044742.1 | *Panorpa debilis* |

| | | |
|---|---|---|
| Epeorus | NC_065805.1 | *Epeorus psi* |
| Epeorus | NC_065804.1 | *Epeorus alexandri* |
| Epeorus | NC_065803.1 | *Epeorus rhithralis* |
| Epeorus | NC_065802.1 | *Epeorus bispinosus* |
| Epeorus | NC_065801.1 | *Epeorus aculeatus* |
| Epeorus | NC_065800.1 | *Epeorus unispinosus* |
| Epeorus | NC_065799.1 | *Epeorus gibbus* |
| Epeorus | NC_065659.1 | *Epeorus pellucidus* |
| Epeorus | NC_065658.1 | *Epeorus montanus* |
| Epeorus | NC_065657.1 | *Epeorus melli* |
| Epeorus | NC_065656.1 | *Epeorus bifurcatus* |
| Epeorus | NC_057491.1 | *Epeorus carinatus* |
| Epeorus | NC_057490.1 | *Epeorus dayongensis* |
| Epeorus | NC_039612.1 | *Epeorus herklotsi* |
| Epiprocta | NC_012644.1 | *Davidius lunatus* |
| Epiprocta | NC_060537.1 | *Gomphus vulgatissimus* |
| Epiprocta | NC_023232.1 | *Epiophlebia superstes* |
| Epiprocta | NC_050976.1 | *Anax parthenope* |
| Epiprocta | NC_031821.1 | *Anax imperator* |
| Epiprocta | NC_061968.1 | *Asiagomphus coreanus* |
| Epiprocta | NC_063611.1 | *Cordulegaster boltonii* |
| Epiprocta | NC_046756.1 | *Epophthalmia elegans* |
| Epiprocta | NC_041425.1 | *Macromia daimoji* |

| | | |
|---|---|---|
| Equus | NC_060655.1 | *Equus hemionus hemippus* |
| Equus | NC_001788.1 | *Equus asinus* |
| Equus | NC_001640.1 | *Equus caballus* |
| Equus | NC_018781.1 | *Equus burchellii chapmani* |
| Equus | NC_018782.1 | *Equus hemionus kulan* |
| Equus | NC_018780.1 | *Equus zebra hartmannae* |
| Equus | NC_020433.1 | *Equus kiang* |
| Equus | NC_016061.1 | *Equus hemionus* |
| Equus | NC_020432.2 | *Equus grevyi* |
| Equus | NC_020476.1 | *Equus zebra H11* |
| Equus | NC_044858.1 | *Equus burchellii quagga* |
| Equus | NC_018783.1 | *Equus ovodovi* |
| Eremias | NC_064329.1 | *Eremias scripta* |
| Eremias | NC_062143.1 | *Eremias szczerbaki* |
| Eremias | NC_060637.1 | *Eremias yarkandensis* |
| Eremias | NC_060561.1 | *Eremias nikolskii* |
| Eremias | NC_056302.1 | *Eremias dzungarica* |
| Eremias | NC_029878.1 | *Eremias stummeri* |
| Eremias | NC_025929.1 | *Eremias przewalskii* |
| Eremias | NC_025320.1 | *Eremias vermiculata* |
| Eremias | NC_025304.1 | *Eremias multiocellata* |
| Eremias | NC_011764.1 | *Eremias brenchleyi* |
| Eulipotyphla | NC_029762.1 | *Condylura cristata* |

| | | |
|---|---|---|
| Eulipotyphla | NC_008156.1 | *Galemys pyrenaicus* |
| Eulipotyphla | NC_063097.1 | *Mogera hainana* |
| Eulipotyphla | NC_029836.1 | *Mogera robusta* |
| Eulipotyphla | NC_005035.1 | *Mogera wogura* |
| Eulipotyphla | NC_056332.1 | *Parascaptor leucura* |
| Eulipotyphla | NC_025777.1 | *Scapanulus oweni* |
| Eulipotyphla | NC_049123.1 | *Scaptochirus moschatus* |
| Eulipotyphla | NC_053257.1 | *Talpa aquitania* |
| Eulipotyphla | NC_039630.1 | *Talpa occidentalis* |
| Eulipotyphla | NC_002391.1 | *Talpa europaea* |
| Eulipotyphla | NC_060485.1 | *Uropsilus investigator* |
| Eulipotyphla | NC_041144.1 | *Uropsilus andersoni* |
| Eulipotyphla | NC_031330.1 | *Uropsilus gracilis* |
| Eulipotyphla | NC_028150.1 | *Uropsilus sp. 4 FT-2015* |
| Eulipotyphla | NC_018598.1 | *Uropsilus sp. 1 FT-2014* |
| Eulipotyphla | NC_023244.1 | *Uropsilus soricipes* |
| Eulipotyphla | NC_005034.1 | *Urotrichus talpoides* |
| Eulipotyphla | NC_002080.2 | *Erinaceus europaeus* |
| Eulipotyphla | NC_005033.1 | *Hemiechinus auritus* |
| Eulipotyphla | NC_002808.1 | *Echinosorex gymnura* |
| Eulipotyphla | NC_010298.1 | *Hylomys suillus* |
| Eulipotyphla | NC_063830.1 | *Neohylomys hainanensis* |
| Eulipotyphla | NC_019626.1 | *Neotetracus sinensis* |

| | | |
|---|---|---|
| Eulipotyphla | NC_024563.1 | *Anourosorex squamipes* |
| Eulipotyphla | NC_042734.1 | *Blarina brevicauda* |
| Eulipotyphla | NC_042694.1 | *Blarina hylophaga* |
| Eulipotyphla | NC_057272.1 | *Blarinella griselda* |
| Eulipotyphla | NC_042750.1 | *Blarinella cf. quadraticauda* |
| Eulipotyphla | NC_041145.1 | *Blarinella wardi* |
| Eulipotyphla | NC_023950.1 | *Blarinella quadraticauda* |
| Eulipotyphla | NC_063622.1 | *Chimarrogale leander* |
| Eulipotyphla | NC_060870.1 | *Chodsigoa hypsibia* |
| Eulipotyphla | NC_053858.1 | *Chodsigoa parva* |
| Eulipotyphla | NC_056333.1 | *Episoriculus leucops* |
| Eulipotyphla | NC_029840.1 | *Episoriculus macrurus* |
| Eulipotyphla | NC_026131.1 | *Episoriculus caudatus* |
| Eulipotyphla | NC_003040.1 | *Episoriculus fumidus* |
| Eulipotyphla | NC_023351.1 | *Nectogale elegans* |
| Eulipotyphla | NC_025559.1 | *Neomys fodiens* |
| Eulipotyphla | NC_042749.1 | *Pantherina griselda* |
| Eulipotyphla | NC_052688.1 | *Soriculus nigrescens* |
| Eulipotyphla | NC_024604.1 | *Suncus murinus* |
| Eumantodea | NC_024028.1 | *Leptomantella albella* |
| Eumantodea | NC_037206.1 | *Sceptuchus simplex* |
| Eumantodea | NC_063851.1 | *Sinomantis denticulata* |
| Eumantodea | NC_037208.1 | *Eomantis yunnanensis* |

| | | |
|---|---|---|
| Eumantodea | NC_051490.1 | *Pliacanthopus bimaculatus* |
| Eumantodea | NC_037205.1 | *Tropidomantis tenera* |
| Felidae | NC_028300.1 | *Catopuma badia* |
| Felidae | NC_027115.1 | *Catopuma temminckii* |
| Felidae | NC_001700.1 | *Felis catus* |
| Felidae | NC_028310.1 | *Felis silvestris* |
| Felidae | NC_028309.1 | *Felis nigripes* |
| Felidae | NC_028308.1 | *Felis margarita* |
| Felidae | NC_028307.1 | *Felis chaus* |
| Felidae | NC_028322.1 | *Leopardus jacobita* |
| Felidae | NC_028321.1 | *Leopardus guigna* |
| Felidae | NC_028320.1 | *Leopardus geoffroyi* |
| Felidae | NC_028318.1 | *Leopardus wiedii* |
| Felidae | NC_028317.1 | *Leopardus tigrinus* |
| Felidae | NC_028315.1 | *Leopardus pardalis* |
| Felidae | NC_028314.1 | *Leopardus colocolo* |
| Felidae | NC_028316.1 | *Leptailurus serval* |
| Felidae | NC_028319.1 | *Lynx pardinus* |
| Felidae | NC_028313.1 | *Lynx canadensis* |
| Felidae | NC_014456.1 | *Lynx rufus* |
| Felidae | NC_008450.1 | *Neofelis nebulosa* |
| Felidae | NC_028323.1 | *Otocolobus manul* |
| Felidae | NC_010642.1 | *Panthera tigris* |

| | | |
|---|---|---|
| Felidae | NC_010641.1 | *Panthera pardus* |
| Felidae | NC_028302.1 | *Panthera leo* |
| Felidae | NC_022842.1 | *Panthera onca* |
| Felidae | NC_018053.1 | *Panthera leo persica* |
| Felidae | NC_014770.1 | *Panthera tigris amoyensis* |
| Felidae | NC_028303.1 | *Pardofelis marmorata* |
| Felidae | NC_028312.1 | *Prionailurus planiceps* |
| Felidae | NC_028305.1 | *Prionailurus viverrinus* |
| Felidae | NC_028304.1 | *Prionailurus rubiginosus* |
| Felidae | NC_028301.1 | *Prionailurus bengalensis* |
| Felidae | NC_016189.1 | *Prionailurus bengalensis euptilurus* |
| Felidae | NC_028299.1 | *Profelis aurata* |
| Felidae | NC_016470.1 | *Puma concolor* |
| Folivora | NC_037527.1 | *Bradypus tridactylus* |
| Folivora | NC_028554.1 | *Bradypus pygmaeus* |
| Folivora | NC_028501.1 | *Bradypus variegatus* |
| Folivora | NC_028555.1 | *Bradypus torquatus* |
| Folivora | NC_042752.1 | *Acratocnus ye* |
| Folivora | NC_027964.1 | *Choloepus hoffmanni* |
| Folivora | NC_006924.1 | *Choloepus didactylus* |
| Folivora | NC_042736.1 | *Megalonyx jeffersonii* |
| Folivora | NC_042754.1 | *Parocnus serus* |
| Folivora | NC_042737.1 | *Megatherium americanum* |

| | | |
|---|---|---|
| Folivora | NC_042753.1 | *Nothrotheriops shastensis* |
| Fordinae | NC_035314.1 | *Baizongia pistaciae* |
| Fordinae | NC_060658.1 | *Kaburagia rhusicola ovogallis* |
| Fordinae | NC_047419.1 | *Meitanaphis microgallis* |
| Fordinae | NC_035315.1 | *Meitanaphis elongallis* |
| Fordinae | NC_035312.1 | *Meitanaphis flavogallis* |
| Formicidae | NC_049088.1 | *Dolichoderus quadripunctatus* |
| Formicidae | NC_041075.1 | *Dolichoderus sibiricus* |
| Formicidae | NC_023093.1 | *Leptomyrmex pallens* |
| Formicidae | NC_045057.1 | *Linepithema humile* |
| Formicidae | NC_049860.1 | *Ochetellus glaber* |
| Formicidae | NC_065783.1 | *Tapinoma ibericum* |
| Formicidae | NC_046425.1 | *Acropyga pallida* |
| Formicidae | NC_046424.1 | *Acropyga myops* |
| Formicidae | NC_046423.1 | *Acropyga kinomurai* |
| Formicidae | NC_046422.1 | *Acropyga guianensis* |
| Formicidae | NC_046421.1 | *Acropyga fuhrmanni* |
| Formicidae | NC_046420.1 | *Acropyga goeldii* |
| Formicidae | NC_046426.1 | *Acropyga panamensis* |
| Formicidae | NC_046399.1 | *Acropyga smithii* |
| Formicidae | NC_046398.1 | *Acropyga sauteri* |
| Formicidae | NC_061037.1 | *Camponotus japonicus* |
| Formicidae | NC_042676.1 | *Camponotus concavus* |

| | | |
|---|---|---|
| Formicidae | NC_053900.1 | *Colobopsis nipponica* |
| Formicidae | NC_030790.1 | *Polyrhachis dives* |
| Formicidae | NC_060872.1 | *Cataglyphis aenescens* |
| Formicidae | NC_060873.1 | *Formica sinae* |
| Formicidae | NC_026711.1 | *Formica selysi* |
| Formicidae | NC_026132.1 | *Formica fusca* |
| Formicidae | NC_039576.1 | *Anoplolepis gracilipes* |
| Formicidae | NC_049861.1 | *Nylanderia flavipes* |
| Formicidae | NC_061556.1 | *Lepisiota frauenfeldi* |
| Formicidae | NC_041202.1 | *Cryptopone sauteri* |
| Formicidae | NC_042678.1 | *Ectomomyrmex javanus* |
| Fulgoroidea | NC_063540.1 | *Augilina tetraina* |
| Fulgoroidea | NC_063539.1 | *Augilina triaina* |
| Fulgoroidea | NC_063537.1 | *Augilodes binghami* |
| Fulgoroidea | NC_059809.1 | *Bambusicaliscelis fanjingensis* |
| Fulgoroidea | NC_059808.1 | *Bambusicaliscelis flavus* |
| Fulgoroidea | NC_063541.1 | *Caliscelis shandongensis* |
| Fulgoroidea | NC_063536.1 | *Cylindratus longicephalus* |
| Fulgoroidea | NC_063531.1 | *Neosymplana vittatum* |
| Fulgoroidea | NC_063538.1 | *Pseudosymplanella nigrifasciata* |
| Fulgoroidea | NC_063542.1 | *Symplanella unipuncta* |
| Fulgoroidea | NC_063533.1 | *Symplanella brevicephala* |
| Fulgoroidea | NC_063532.1 | *Symplanella nigricans* |

| | | |
|---|---|---|
| Fulgoroidea | NC_059811.1 | *Youtuus erythrus* |
| Fulgoroidea | NC_059810.1 | *Youtuus strigatus* |
| Fulgoroidea | NC_066441.1 | *Oecleopsis sinicus* |
| Fulgoroidea | NC_052690.1 | *Bambusiphaga taibaishana* |
| Fulgoroidea | NC_052689.1 | *Bambusiphaga furca* |
| Fulgoroidea | NC_037181.1 | *Changeondelphax velitchkovskyi* |
| Fulgoroidea | NC_052691.1 | *Chloriona tateyamana* |
| Fulgoroidea | NC_052692.1 | *Epeurysa nawaii* |
| Fulgoroidea | NC_052693.1 | *Ishiharodelphax matsuyamensis* |
| Fulgoroidea | NC_033388.1 | *Nilaparvata bakeri* |
| Fulgoroidea | NC_024627.1 | *Nilaparvata muiri* |
| Fulgoroidea | NC_021748.1 | *Nilaparvata lugens* |
| Fulgoroidea | NC_037182.1 | *Peregrinus maidis* |
| Fulgoroidea | NC_052694.1 | *Perkinsiella saccharicida* |
| Fulgoroidea | NC_042179.1 | *Saccharosydne procerus* |
| Fulgoroidea | NC_056127.1 | *Sogatella kolophon* |
| Fulgoroidea | NC_042180.1 | *Sogatella vibix* |
| Fulgoroidea | NC_021417.1 | *Sogatella furcifera* |
| Fulgoroidea | NC_052695.1 | *Tropidocephala brunnipennis* |
| Fulgoroidea | NC_012617.1 | *Geisha distinctissima* |
| Fulgoroidea | NC_053739.1 | *Lophops carinata* |
| Fulgoroidea | NC_063535.1 | *Symplana lii* |
| Fulgoroidea | NC_063534.1 | *Symplana brevistrata* |

| | | |
|---|---|---|
| Fulgoroidea | NC_060807.1 | *Pochazia confusa* |
| Fulgoroidea | NC_060806.1 | *Pochazia guttifera* |
| Fulgoroidea | NC_060730.1 | *Pochazia discreta* |
| Fulgoroidea | NC_060809.1 | *Ricania fumosa* |
| Fulgoroidea | NC_060808.1 | *Ricania simulans* |
| Fulgoroidea | NC_051496.1 | *Ricania shantungensis* |
| Fulgoroidea | NC_031369.1 | *Ricania speculum* |
| Fulgoroidea | NC_013706.1 | *Laodelphax striatellus* |
| Galliformes | NC_024618.1 | *Crax rubra* |
| Galliformes | NC_024617.1 | *Crax daubentoni* |
| Galliformes | NC_052778.1 | *Penelope pileata* |
| Galliformes | NC_007227.1 | *Alectura lathami* |
| Galliformes | NC_014180.1 | *Acryllium vulturinum* |
| Galliformes | NC_034374.1 | *Numida meleagris* |
| Galliformes | NC_029340.1 | *Callipepla squamata* |
| Galliformes | NC_024620.1 | *Colinus virginianus* |
| Galliformes | NC_052784.1 | *Odontophorus gujanensis* |
| Galliformes | NC_020585.1 | *Alectoris chukar* |
| Galliformes | NC_012453.1 | *Arborophila rufipectus* |
| Galliformes | NC_022683.1 | *Arborophila ardens* |
| Galliformes | NC_022684.1 | *Arborophila brunneopectus* |
| Galliformes | NC_020584.1 | *Arborophila rufogularis* |
| Galliformes | NC_020583.1 | *Bambusicola fytchii* |

| | | |
|---|---|---|
| Galliformes | NC_011816.1 | *Bambusicola thoracica* |
| Galliformes | NC_024619.1 | *Caloperdix oculeus* |
| Galliformes | NC_020590.1 | *Chrysolophus amherstiae* |
| Galliformes | NC_014576.1 | *Chrysolophus pictus* |
| Galliformes | NC_003408.1 | *Coturnix japonica* |
| Galliformes | NC_004575.1 | *Coturnix chinensis* |
| Galliformes | NC_026548.1 | *Crossoptilon mantchuricum* |
| Galliformes | NC_026547.1 | *Crossoptilon harmani* |
| Galliformes | NC_011817.1 | *Francolinus pintadeanus* |
| Galliformes | NC_007238.1 | *Gallus varius* |
| Galliformes | NC_007240.1 | *Gallus sonneratii* |
| Galliformes | NC_007239.1 | *Gallus lafayetii* |
| Galliformes | NC_034001.1 | *Haematortyx sanguiniceps* |
| Galliformes | NC_018033.1 | *Ithaginis cruentus* |
| Galliformes | NC_040850.1 | *Lophophorus impejanus* |
| Galliformes | NC_020589.1 | *Lophophorus sclateri* |
| Galliformes | NC_013979.1 | *Lophophorus lhuysii* |
| Galliformes | NC_023779.1 | *Lophura swinhoii* |
| Galliformes | NC_012895.1 | *Lophura nycthemera* |
| Galliformes | NC_010781.1 | *Lophura ignita* |
| Galliformes | NC_024533.1 | *Pavo cristatus* |
| Galliformes | NC_012897.1 | *Pavo muticus* |
| Galliformes | NC_023940.1 | *Perdix hodgsoniae* |

| | | |
|---|---|---|
| Galliformes | NC_020588.1 | *Perdix dauurica* |
| Galliformes | NC_015526.1 | *Phasianus colchicus* |
| Galliformes | NC_010778.1 | *Phasianus versicolor* |
| Galliformes | NC_044743.1 | *Polyplectron malacense* |
| Galliformes | NC_024615.1 | *Polyplectron napoleonis* |
| Galliformes | NC_023264.1 | *Polyplectron germaini* |
| Galliformes | NC_012900.1 | *Polyplectron bicalcaratum* |
| Galliformes | NC_024616.1 | *Ptilopachus petrosus* |
| Galliformes | NC_020587.1 | *Pucrasia macrolopha* |
| Galliformes | NC_010767.1 | *Syrmaticus soemmerringi ijimae* |
| Galliformes | NC_010770.1 | *Syrmaticus reevesii* |
| Galliformes | NC_010774.1 | *Syrmaticus humiae* |
| Galliformes | NC_027279.1 | *Tetraogallus himalayensis* |
| Galliformes | NC_023939.1 | *Tetraogallus tibetanus* |
| Galliformes | NC_020613.1 | *Tetraophasis szechenyii* |
| Galliformes | NC_018034.1 | *Tetraophasis obscurus* |
| Galliformes | NC_020586.1 | *Tragopan temminckii* |
| Galliformes | NC_013619.1 | *Tragopan caboti* |
| Galliformes | NC_010195.2 | *Meleagris gallopavo* |
| Galliformes | NC_034002.1 | *Lagopus muta* |
| Galliformes | NC_024554.1 | *Lyrurus tetrix* |
| Galliformes | NC_043950.1 | *Tetrao parvirostris kamtschaticus* |
| Galliformes | NC_025318.1 | *Tetrastes sewerzowi* |

| | | |
|---|---|---|
| Galliformes | NC_020591.1 | *Tetrastes bonasia* |
| Galloanserae | NC_023969.1 | *Aix galericulata* |
| Galloanserae | NC_052827.1 | *Asarcornis scutulata* |
| Galloanserae | NC_010965.1 | *Cairina moschata* |
| Galloanserae | NC_045373.1 | *Mareca strepera* |
| Galloanserae | NC_040986.1 | *Mergus merganser* |
| Galloanserae | NC_016723.1 | *Mergus squamatus* |
| Galloanserae | NC_052807.1 | *Chauna torquata* |
| Galloanserae | NC_005933.1 | *Anseranas semipalmata* |
| Galloanserae | NC_059801.1 | *Aythya nyroca* |
| Galloanserae | NC_024602.1 | *Aythya ferina* |
| Galloanserae | NC_024595.1 | *Aythya fuligula* |
| Galloanserae | NC_000877.1 | *Aythya americana* |
| Galloanserae | NC_024922.1 | *Netta rufina* |
| Galloanserae | NC_012844.1 | *Dendrocygna javanica* |
| Galloanserae | NC_024640.1 | *Tadorna ferruginea* |
| Garrulax | NC_065197.1 | *Garrulax courtoisi* |
| Garrulax | NC_060673.1 | *Garrulax chinensis* |
| Garrulax | NC_037464.1 | *Garrulax albogularis* |
| Garrulax | NC_034373.1 | *Garrulax elliotii* |
| Garrulax | NC_034353.1 | *Garrulax formosus* |
| Garrulax | NC_029402.1 | *Garrulax affinis* |
| Garrulax | NC_028186.1 | *Garrulax sannio* |

| | | |
|---|---|---|
| Garrulax | NC_028082.1 | *Garrulax poecilorhynchus* |
| Garrulax | NC_027657.1 | *Garrulax ocellatus* |
| Garrulax | NC_026068.1 | *Garrulax perspicillatus* |
| Garrulax | NC_024553.1 | *Garrulax cineraceus* |
| Garrulax | NC_020429.1 | *Garrulax canorus* |
| Gebiidea | NC_043831.1 | *Axianassa australis* |
| Gebiidea | NC_043832.1 | *Laomedia healyi* |
| Gebiidea | NC_041157.1 | *Thalassina squamifera* |
| Gebiidea | NC_019608.1 | *Thalassina kelanang* |
| Gebiidea | NC_019606.1 | *Austinogebia edulis* |
| Gebiidea | NC_041158.1 | *Upogebia bowerbankii* |
| Gebiidea | NC_025943.1 | *Upogebia yokoyai* |
| Gebiidea | NC_020023.1 | *Upogebia pusilla* |
| Gebiidea | NC_019607.1 | *Upogebia major* |
| Gekkonidae | NC_020039.1 | *Cnemaspis limi* |
| Gekkonidae | NC_052836.1 | *Gekko subpalmatus* |
| Gekkonidae | NC_035304.1 | *Gekko hokouensis* |
| Gekkonidae | NC_027191.1 | *Gekko chinensis* |
| Gekkonidae | NC_028035.1 | *Gekko japonicus* |
| Gekkonidae | NC_018050.1 | *Gekko swinhonis* |
| Gekkonidae | NC_007627.1 | *Gekko gecko* |
| Gekkonidae | NC_008772.1 | *Gekko vittatus* |
| Gekkonidae | NC_066456.1 | *Hemidactylus ulii* |

| | | |
|---|---|---|
| Gekkonidae | NC_066455.1 | *Hemidactylus mandebensis* |
| Gekkonidae | NC_066454.1 | *Hemidactylus farasani* |
| Gekkonidae | NC_066453.1 | *Hemidactylus almakhwah* |
| Gekkonidae | NC_025938.1 | *Hemidactylus bowringii* |
| Gekkonidae | NC_012902.2 | *Hemidactylus frenatus* |
| Gekkonidae | NC_025782.1 | *Lepidodactylus lugubris* |
| Gekkonidae | NC_028326.1 | *Paroedura picta* |
| Gekkonidae | NC_025783.1 | *Uroplatus ebenaui* |
| Gekkonidae | NC_025779.1 | *Uroplatus fimbriatus* |
| Geometridae | NC_034804.1 | *Abraxas suspecta* |
| Geometridae | NC_061234.1 | *Amraica recursaria* |
| Geometridae | NC_024824.1 | *Apocheima cinerarium* |
| Geometridae | NC_061644.1 | *Biston regalis* |
| Geometridae | NC_057638.1 | *Biston thoracicaria* |
| Geometridae | NC_030769.1 | *Biston perclara* |
| Geometridae | NC_030632.1 | *Biston thibetaria* |
| Geometridae | NC_027111.1 | *Biston suppressaria* |
| Geometridae | NC_020004.1 | *Biston panterinaria* |
| Geometridae | NC_062108.1 | *Chorodna fulgurita* |
| Geometridae | NC_062170.1 | *Cleora fraterna* |
| Geometridae | NC_061237.1 | *Cotta incongruaria* |
| Geometridae | NC_057121.1 | *Ectropis grisescens* |
| Geometridae | NC_036717.1 | *Ectropis obliqua* |

| | | |
|---|---|---|
| Geometridae | NC_047212.1 | *Erannis ankeraria* |
| Geometridae | NC_061236.1 | *Hydatocapnia marginata* |
| Geometridae | NC_027948.1 | *Jankowskia athleta* |
| Geometridae | NC_061239.1 | *Luxiaria mitorrhaphes* |
| Geometridae | NC_061233.1 | *Menophra senilis* |
| Geometridae | NC_061235.1 | *Ophthalmitis albosignaria* |
| Geometridae | NC_010522.1 | *Phthonandria atrilineata* |
| Geometridae | NC_061036.1 | *Xanthabraxas hemionata* |
| Geometridae | NC_056092.1 | *Iotaphora admirabilis* |
| Geometridae | NC_061238.1 | *Lophophelma iterans* |
| Geometridae | NC_061231.1 | *Pingasa rufofasciata* |
| Geometridae | NC_062174.1 | *Tanaorhinus viridiluteata* |
| Geometridae | NC_027723.1 | *Operophtera brumata* |
| Geometridae | NC_057249.1 | *Pasiphila chloerata* |
| Geometridae | NC_066404.1 | *Psychophora sabini* |
| Geometridae | NC_061232.1 | *Somatina indicataria* |
| Glires | NC_057103.1 | *Ochotona hyperborea* |
| Glires | NC_005358.1 | *Ochotona princeps* |
| Glires | NC_052873.1 | *Ochotona coreana* |
| Glires | NC_044120.1 | *Ochotona dauurica* |
| Glires | NC_039987.1 | *Ochotona koslowi* |
| Glires | NC_037186.1 | *Ochotona erythrotis* |
| Glires | NC_011029.1 | *Ochotona curzoniae* |

| | | |
|---|---|---|
| Glires | NC_003033.1 | *Ochotona collaris* |
| Glires | NC_064132.1 | *Brachylagus idahoensis* |
| Glires | NC_001913.1 | *Oryctolagus cuniculus* |
| Glires | NC_066654.1 | *Romerolagus diazi* |
| Gonodactyloidea | NC_065860.1 | *Gonodactylaceus falcatus* |
| Gonodactyloidea | NC_053854.1 | *Gonodactylaceus randalli* |
| Gonodactyloidea | NC_060311.1 | *Gonodactylus smithii* |
| Gonodactyloidea | NC_007442.1 | *Gonodactylus chiragra* |
| Gonodactyloidea | NC_060590.1 | *Neogonodactylus oerstedii* |
| Gonodactyloidea | NC_060589.1 | *Neogonodactylus bredini* |
| Gonodactyloidea | NC_063582.1 | *Odontodactylus havanensis* |
| Gonodactyloidea | NC_063584.1 | *Mesacturoides brevisquamatus* |
| Gonodactyloidea | NC_050686.1 | *Taku spinosocarinatus* |
| Grapsoidea | NC_052834.1 | *Chasmagnathus convexus* |
| Grapsoidea | NC_047209.1 | *Chiromantes eulimene* |
| Grapsoidea | NC_042142.1 | *Chiromantes haematocheir* |
| Grapsoidea | NC_041212.1 | *Chiromantes dehaani* |
| Grapsoidea | NC_033866.1 | *Clistocoeloma sinense* |
| Grapsoidea | NC_050045.1 | *Cyclograpsus intermedius* |
| Grapsoidea | NC_063149.1 | *Episesarma lafondii* |
| Grapsoidea | NC_011598.1 | *Eriocheir hepuensis* |
| Grapsoidea | NC_011597.1 | *Eriocheir japonica* |
| Grapsoidea | NC_006992.1 | *Eriocheir sinensis* |

| | | |
|---|---|---|
| Grapsoidea | NC_038179.1 | *Gaetice depressus* |
| Grapsoidea | NC_065158.1 | *Helicana japonica* |
| Grapsoidea | NC_034995.1 | *Helicana wuana* |
| Grapsoidea | NC_033865.1 | *Helice latimera* |
| Grapsoidea | NC_065995.1 | *Hemigrapsus sinensis* |
| Grapsoidea | NC_040976.1 | *Metaplax longipes* |
| Grapsoidea | NC_040977.1 | *Nanosesarma minutum* |
| Grapsoidea | NC_041211.1 | *Neoeriocheir leptognathus* |
| Grapsoidea | NC_061931.1 | *Parasesarma eumolpe* |
| Grapsoidea | NC_039990.1 | *Parasesarma affine* |
| Grapsoidea | NC_038066.1 | *Parasesarma pictum* |
| Grapsoidea | NC_030046.2 | *Parasesarma tripectinis* |
| Grapsoidea | NC_042685.1 | *Pseudohelice subquadrata* |
| Grapsoidea | NC_056882.1 | *Varuna litterata* |
| Grapsoidea | NC_037155.1 | *Varuna yui* |
| Grapsoidea | NC_057477.1 | *Cardisoma armatum* |
| Grapsoidea | NC_039105.1 | *Cardisoma carnifex* |
| Grapsoidea | NC_057475.1 | *Gecarcoidea lalandii* |
| Grapsoidea | NC_039811.2 | *Gecarcoidea natalis* |
| Grapsoidea | NC_057301.1 | *Grapsus albolineatus* |
| Grapsoidea | NC_029724.1 | *Grapsus tenuicrustatus* |
| Grapsoidea | NC_042152.1 | *Metopograpsus frontalis* |
| Grapsoidea | NC_038178.1 | *Metopograpsus quadridentatus* |

| | | |
|---|---|---|
| Grapsoidea | NC_039109.1 | *Pachygrapsus marmoratus* |
| Grapsoidea | NC_021754.1 | *Pachygrapsus crassipes* |
| Grapsoidea | NC_063602.1 | *Plagusia squamosa* |
| Grapsoidea | NC_013480.1 | *Xenograpsus testudinatus* |
| Gruiformes | NC_024593.1 | *Amaurornis phoenicurus* |
| Gruiformes | NC_023982.1 | *Amaurornis akool* |
| Gruiformes | NC_039814.1 | *Atlantisia rogersi* |
| Gruiformes | NC_012143.1 | *Coturnicops exquisitus* |
| Gruiformes | NC_063128.1 | *Coturnicops noveboracensis* |
| Gruiformes | NC_025501.1 | *Eulabeornis castaneoventris* |
| Gruiformes | NC_025500.1 | *Fulica atra* |
| Gruiformes | NC_028408.1 | *Gallicrex cinerea* |
| Gruiformes | NC_015236.1 | *Gallinula chloropus* |
| Gruiformes | NC_041577.1 | *Gallirallus striatus* |
| Gruiformes | NC_025507.1 | *Gallirallus philippensis* |
| Gruiformes | NC_012140.1 | *Gallirallus okinawae* |
| Gruiformes | NC_056095.1 | *Laterallus spilonota* |
| Gruiformes | NC_025502.1 | *Lewinia muelleri* |
| Gruiformes | NC_054210.1 | *Nesotrochis steganinos* |
| Gruiformes | NC_010092.1 | *Porphyrio hochstetteri* |
| Gruiformes | NC_053843.1 | *Porzana pusilla* |
| Gruiformes | NC_037406.1 | *Porzana paykullii* |
| Gruiformes | NC_012142.1 | *Rallina eurizonoides sepiaria* |

| | | |
|---|---|---|
| Gruiformes | NC_041578.1 | *Rallus aquaticus* |
| Gruiformes | NC_052808.1 | *Zapornia atra* |
| Gruiformes | NC_052795.1 | *Aramus guarauna* |
| Gruiformes | NC_020570.1 | *Balearica pavonina* |
| Gruiformes | NC_020569.1 | *Balearica regulorum* |
| Gruiformes | NC_025499.1 | *Heliornis fulica* |
| Gruiformes | NC_052775.1 | *Ardeotis kori* |
| Gruiformes | NC_014046.1 | *Otis tarda* |
| Gruiformes | NC_052785.1 | *Psophia crepitans* |
| Gruiformes | NC_010091.1 | *Rhynochetos jubatus* |
| Grus | NC_021368.1 | *Grus vipio* |
| Grus | NC_020582.1 | *Grus canadensis* |
| Grus | NC_020581.1 | *Grus antigone* |
| Grus | NC_020580.1 | *Grus rubicunda* |
| Grus | NC_020579.1 | *Grus nigricollis* |
| Grus | NC_020578.1 | *Grus monacha* |
| Grus | NC_020577.1 | *Grus grus* |
| Grus | NC_020576.1 | *Grus americana* |
| Grus | NC_020575.1 | *Grus japonensis* |
| Grus | NC_020574.1 | *Grus leucogeranus* |
| Grus | NC_020571.1 | *Grus carunculatus* |
| Gryllidea | NC_037914.1 | *Cardiodactylus muiri* |
| Gryllidea | NC_041236.1 | *Xenogryllus marmoratus* |

| | | |
|---|---|---|
| Gryllidea | NC_057195.1 | *Gryllodes sigillatus* |
| Gryllidea | NC_057053.1 | *Gryllus veletis* |
| Gryllidea | NC_057052.1 | *Gryllus lineaticeps* |
| Gryllidea | NC_053546.1 | *Gryllus bimaculatus* |
| Gryllidea | NC_033985.1 | *Loxoblemmus doenitzi* |
| Gryllidea | NC_028619.1 | *Teleogryllus oceanicus* |
| Gryllidea | NC_011823.1 | *Teleogryllus emma* |
| Gryllidea | NC_060317.1 | *Turanogryllus eous* |
| Gryllidea | NC_030762.1 | *Velarifictorus hemelytrus* |
| Gryllidea | NC_039667.1 | *Ornebius kanetataki* |
| Gryllidea | NC_039666.1 | *Ornebius bimaculatus* |
| Gryllidea | NC_039739.1 | *Ornebius fuscicercis* |
| Gryllidea | NC_011301.1 | *Myrmecophilus manni* |
| Gryllidea | NC_045847.1 | *Dianemobius furumagiensis* |
| Gryllidea | NC_045846.1 | *Dianemobius fascipes* |
| Gryllidea | NC_045848.1 | *Polionemobius taprobanensis* |
| Gryllidea | NC_034799.1 | *Oecanthus sinensis* |
| Gryllidea | NC_034797.1 | *Truljalia hibinonis* |
| Gryllidea | NC_045841.1 | *Homoeoxipha nigripes* |
| Gryllidea | NC_050742.1 | *Natula pravdini* |
| Gryllidea | NC_053543.1 | *Svistella anhuiensis* |
| Gryllidea | NC_032077.1 | *Trigonidium sjostedti* |
| Gryllidea | NC_039664.1 | *Cacoplistes rogenhoferi* |

| | | |
|---|---|---|
| Gryllidea | NC_039665.1 | *Meloimorpha japonica* |
| Gymnophiona | NC_023508.1 | *Caecilia gracilis* |
| Gymnophiona | NC_023507.1 | *Caecilia tentaculata* |
| Gymnophiona | NC_020137.1 | *Caecilia volcani* |
| Gymnophiona | NC_020142.1 | *Oscaecilia ochrocephala* |
| Gymnophiona | NC_021369.1 | *Chikila fulleri* |
| Gymnophiona | NC_020138.1 | *Dermophis mexicanus* |
| Gymnophiona | NC_020155.1 | *Geotrypetes seraphini* |
| Gymnophiona | NC_020139.1 | *Gymnopis multiplicata* |
| Gymnophiona | NC_023518.1 | *Schistometopum gregorii* |
| Gymnophiona | NC_020154.1 | *Boulengerula taitana* |
| Gymnophiona | NC_020136.1 | *Boulengerula boulengeri* |
| Gymnophiona | NC_019586.1 | *Herpele squalostoma* |
| Gymnophiona | NC_023511.1 | *Ichthyophis bombayensis* |
| Gymnophiona | NC_006404.1 | *Ichthyophis bannanicus* |
| Gymnophiona | NC_006302.1 | *Ichthyophis glutinosus* |
| Gymnophiona | NC_023519.1 | *Uraeotyphlus gansi* |
| Gymnophiona | NC_006305.1 | *Uraeotyphlus cf. oxyurus* |
| Gymnophiona | NC_006301.1 | *Gegeneophis ramaswamii* |
| Gymnophiona | NC_023510.1 | *Grandisonia sechellensis* |
| Gymnophiona | NC_020140.1 | *Hypogeophis rostratus* |
| Gymnophiona | NC_023512.1 | *Indotyphlus maharashtraensis* |
| Gymnophiona | NC_023517.1 | *Praslinia cooperi* |

| | | |
|---|---|---|
| Gymnophiona | NC_006303.1 | *Rhinatrema bivittatum* |
| Gymnophiona | NC_019596.1 | *Crotaphatrema lamottei* |
| Gymnophiona | NC_006304.1 | *Scolecomorphus vittatus* |
| Gymnophiona | NC_023513.1 | *Luetkenotyphlus brasiliensis* |
| Gymnophiona | NC_026067.1 | *Luscinia cyanura* |
| Gymnophiona | NC_015074.1 | *Luscinia calliope* |
| Gymnophiona | NC_023515.1 | *Microcaecilia unicolor* |
| Gymnophiona | NC_023514.1 | *Microcaecilia dermatophaga* |
| Gymnophiona | NC_020141.1 | *Microcaecilia sp. PZ-2009* |
| Gymnophiona | NC_007911.1 | *Siphonops annulatus* |
| Gymnophiona | NC_023509.1 | *Chthonerpeton indistinctum* |
| Gymnophiona | NC_023516.1 | *Potomotyphlus kaupii* |
| Gymnophiona | NC_002471.1 | *Typhlonectes natans* |
| Hadziida | NC_019653.1 | *Metacrangonyx repens* |
| Hadziida | NC_019660.1 | *Metacrangonyx remyi* |
| Hadziida | NC_019659.1 | *Metacrangonyx panousei* |
| Hadziida | NC_019658.1 | *Metacrangonyx longicaudus* |
| Hadziida | NC_019657.1 | *Metacrangonyx spinicaudatus* |
| Hadziida | NC_019656.1 | *Metacrangonyx ilvanus* |
| Hadziida | NC_019655.1 | *Metacrangonyx goulmimensis* |
| Hadziida | NC_019654.1 | *Metacrangonyx dominicanus* |
| Hadziida | NC_013032.1 | *Metacrangonyx longipes* |
| Haemaphysalis | NC_064124.1 | *Haemaphysalis nepalensis* |

| | | |
|---|---|---|
| Haemaphysalis | NC_062164.1 | *Haemaphysalis colasbelcouri* |
| Haemaphysalis | NC_062163.1 | *Haemaphysalis mageshimaensis* |
| Haemaphysalis | NC_062162.1 | *Haemaphysalis cornigera* |
| Haemaphysalis | NC_062161.1 | *Haemaphysalis kitaokai* |
| Haemaphysalis | NC_062160.1 | *Haemaphysalis yeni* |
| Haemaphysalis | NC_062159.1 | *Haemaphysalis campanulata* |
| Haemaphysalis | NC_062158.1 | *Haemaphysalis doenitzi* |
| Haemaphysalis | NC_062067.1 | *Haemaphysalis qinghaiensis* |
| Haemaphysalis | NC_062066.1 | *Haemaphysalis tibetensis* |
| Haemaphysalis | NC_062065.1 | *Haemaphysalis danieli* |
| Haemaphysalis | NC_062064.1 | *Haemaphysalis punctata* |
| Haemaphysalis | NC_062063.1 | *Haemaphysalis sulcata* |
| Haemaphysalis | NC_058312.1 | *Haemaphysalis montgomeryi yunnan* |
| Haemaphysalis | NC_039765.1 | *Haemaphysalis hystricis* |
| Haemaphysalis | NC_020335.1 | *Haemaphysalis inermis* |
| Haemaphysalis | NC_041076.1 | *Haemaphysalis bancrofti* |
| Haemaphysalis | NC_037493.1 | *Haemaphysalis longicornis* |
| Haemaphysalis | NC_037246.1 | *Haemaphysalis japonica* |
| Haemaphysalis | NC_034785.1 | *Haemaphysalis concinna* |
| Haemaphysalis | NC_020334.1 | *Haemaphysalis formosensis* |
| Haemaphysalis | NC_005292.1 | *Haemaphysalis flava* |
| Harpalinae | NC_066084.1 | *Galerita orientalis* |
| Harpalinae | NC_066080.1 | *Harpalus griseus* |

| | | |
|---|---|---|
| Harpalinae | NC_066079.1 | *Harpalus anxius* |
| Harpalinae | NC_066078.1 | *Harpalus discrepans* |
| Harpalinae | NC_046953.1 | *Harpalus pensylvanicus* |
| Harpalinae | NC_045094.1 | *Harpalus sinicus* |
| Harpalinae | NC_030592.1 | *Abax parallelepipedus* |
| Harpalinae | NC_044760.1 | *Pterostichus niger* |
| Harpalinae | NC_036268.1 | *Amara communis* |
| Henophidia | NC_014343.1 | *Anilius scytale* |
| Henophidia | NC_007401.1 | *Cylindrophis ruffus* |
| Henophidia | NC_042397.1 | *Malayopython reticulatus* |
| Henophidia | NC_021479.1 | *Python bivittatus* |
| Henophidia | NC_015812.1 | *Python molurus molurus* |
| Henophidia | NC_007399.1 | *Python regius* |
| Henophidia | NC_012573.1 | *Tropidophis haetianus* |
| Henophidia | NC_007402.1 | *Xenopeltis unicolor* |
| Henophidia | NC_007398.1 | *Boa constrictor* |
| Henophidia | NC_063114.1 | *Chilabothrus argentum* |
| Hepialoidea | NC_060512.1 | *Ahamus yushuensis* |
| Hepialoidea | NC_018095.1 | *Ahamus yunnanensis* |
| Hepialoidea | NC_029873.1 | *Endoclita signifer* |
| Hepialoidea | NC_028348.1 | *Hepialus xiaojinensis* |
| Hepialoidea | NC_024424.1 | *Napialus hunanensis* |
| Hepialoidea | NC_044770.1 | *Thitarodes damxungensis* |

| | | |
|---|---|---|
| Hepialoidea | NC_032649.1 | *Thitarodes sejilaensis* |
| Hepialoidea | NC_026903.1 | *Thitarodes gonggaensis* |
| Hepialoidea | NC_023530.1 | *Thitarodes pui* |
| Hepialoidea | NC_018094.1 | *Thitarodes renzhiensis* |
| Hepialoidea | NC_040921.1 | *Triodia sylvina* |
| Hesperiidae | NC_053717.1 | *Abraximorpha davidii* |
| Hesperiidae | NC_060817.1 | *Capila translucida* |
| Hesperiidae | NC_060818.1 | *Celaenorrhinus aspersus* |
| Hesperiidae | NC_022853.1 | *Celaenorrhinus maculosa* |
| Hesperiidae | NC_016704.1 | *Ctenoptilum vasava* |
| Hesperiidae | NC_024648.1 | *Daimio tethys* |
| Hesperiidae | NC_060819.1 | *Gerosis phisara* |
| Hesperiidae | NC_060820.1 | *Mooreana trichoneura* |
| Hesperiidae | NC_060821.1 | *Pseudocoladenia dea* |
| Hesperiidae | NC_060822.1 | *Satarupa nymphalis* |
| Hesperiidae | NC_060823.1 | *Tagiades menaka* |
| Hesperiidae | NC_036220.1 | *Tagiades litigiosa* |
| Hesperiidae | NC_042214.1 | *Astictopterus jama* |
| Hesperiidae | NC_042215.1 | *Isoteinon lamprospilus* |
| Hesperiidae | NC_029136.1 | *Parnara guttata* |
| Hesperiidae | NC_056138.1 | *Pelopidas mathias* |
| Hesperiidae | NC_034676.1 | *Burara striata* |
| Hesperiidae | NC_024647.1 | *Choaspes benjaminii* |

| | | |
|---|---|---|
| Hesperiidae | NC_045249.1 | *Hasora badra* |
| Hesperiidae | NC_027263.1 | *Hasora anura* |
| Hesperiidae | NC_027170.1 | *Hasora vitta* |
| Hesperiidae | NC_048454.1 | *Malaza empyreus* |
| Hesperiidae | NC_050387.1 | *Malaza fastuosus* |
| Hesperiidae | NC_048455.1 | *Malaza carmides* |
| Hesperiidae | NC_042216.1 | *Notocrypta curvifascia* |
| Hesperiidae | NC_029826.1 | *Lerema accius* |
| Hesperiidae | NC_018048.1 | *Ochlodes venata* |
| Hesperiidae | NC_039946.1 | *Apostictopterus fuliginosus* |
| Hesperiidae | NC_039947.1 | *Barca bicolor* |
| Hesperiidae | NC_024646.1 | *Carterocephalus silvicola* |
| Hesperiidae | NC_028506.1 | *Heteropterus morpheus* |
| Hesperiidae | NC_045924.1 | *Leptalina unicolor* |
| Hesperiidae | NC_060824.1 | *Erynnis popoviana* |
| Hesperiidae | NC_021427.1 | *Erynnis montanus* |
| Hesperiidae | NC_034231.1 | *Euschemon rafflesia* |
| Hesperiidae | NC_030192.1 | *Pyrgus maculatus* |
| Hesperiidae | NC_024650.1 | *Potanthus flavus* |
| Hesperiidae | NC_048456.1 | *Rachelia extrusus* |
| Hesperiidae | NC_024649.1 | *Lobocla bifasciatus* |
| Heterotermes | NC_030034.1 | *Heterotermes validus* |
| Heterotermes | NC_030029.1 | *Heterotermes cf. occiduus* |

| | | |
|---|---|---|
| Heterotermes | NC_030028.1 | *Heterotermes cf. occiduus* |
| Heterotermes | NC_030027.1 | *Heterotermes nr. tenuis* |
| Heterotermes | NC_030026.1 | *Heterotermes malabaricus* |
| Heterotermes | NC_030024.1 | *Heterotermes cf. paradoxus* |
| Heterotermes | NC_030023.1 | *Heterotermes cf. paradoxus* |
| Heterotermes | NC_030033.1 | *Heterotermes vagus* |
| Heterotermes | NC_030032.1 | *Heterotermes tenuis* |
| Heterotermes | NC_030031.1 | *Heterotermes tenuior* |
| Heterotermes | NC_030030.1 | *Heterotermes platycephalus* |
| Heterotermes | NC_030025.1 | *Heterotermes crinitus* |
| Heterotermes | NC_018127.1 | *Heterotermes sp.* |
| Heterotremata | NC_020314.1 | *Austinograea alayseae* |
| Heterotremata | NC_020312.1 | *Austinograea rodriguezensis* |
| Heterotremata | NC_027414.1 | *Gandalfus puia* |
| Heterotremata | NC_013713.1 | *Gandalfus yunohana* |
| Heterotremata | NC_035300.1 | *Segonzacia mesatlantica* |
| Heterotremata | NC_006281.1 | *Callinectes sapidus* |
| Heterotremata | NC_060621.1 | *Charybdis hellerii* |
| Heterotremata | NC_037695.1 | *Charybdis bimaculata* |
| Heterotremata | NC_036132.1 | *Charybdis natator* |
| Heterotremata | NC_024632.1 | *Charybdis feriata* |
| Heterotremata | NC_013246.1 | *Charybdis japonica* |
| Heterotremata | NC_057302.1 | *Lissocarcinus arkati* |

| | | |
|---|---|---|
| Heterotremata | NC_037173.1 | *Monomia gladiator* |
| Heterotremata | NC_040124.1 | *Portunus gracilimanus* |
| Heterotremata | NC_028225.1 | *Portunus sanguinolentus* |
| Heterotremata | NC_026209.1 | *Portunus pelagicus* |
| Heterotremata | NC_005037.1 | *Portunus trituberculatus* |
| Heterotremata | NC_012567.1 | *Scylla tranquebarica* |
| Heterotremata | NC_012572.1 | *Scylla paramamosain* |
| Heterotremata | NC_012569.1 | *Scylla olivacea* |
| Heterotremata | NC_012565.1 | *Scylla serrata* |
| Heterotremata | NC_039640.1 | *Thalamita sima* |
| Heterotremata | NC_053638.1 | *Matuta victor* |
| Heterotremata | NC_039351.1 | *Matuta planipes* |
| Heterotremata | NC_039639.1 | *Orithyia sinica* |
| Heterotremata | NC_047195.1 | *Calappa bilineata* |
| Heterotremata | NC_061949.1 | *Myra affinis* |
| Heterotremata | NC_030047.1 | *Pyrhila pisum* |
| Heterotremata | NC_049029.1 | *Daldorfia horrida* |
| Heterotremata | NC_042695.1 | *Ovalipes punctatus* |
| Hippoboscoidea | NC_037368.1 | *Melophagus ovinus* |
| Hippoboscoidea | NC_044702.1 | *Paradyschiria parvula* |
| Hippoboscoidea | NC_044652.1 | *Paratrichobius longicrus* |
| Hirundo | NC_050299.1 | *Hirundo albigularis* |
| Hirundo | NC_050293.1 | *Hirundo aethiopica* |

| | | |
|---|---|---|
| Hirundo | NC_050291.1 | *Hirundo nigrita* |
| Hirundo | NC_050290.1 | *Hirundo neoxena* |
| Hirundo | NC_050288.1 | *Hirundo dimidiata* |
| Hirundo | NC_050287.1 | *Hirundo angolensis* |
| Hirundo | NC_050286.1 | *Hirundo smithii* |
| Hirundo | NC_050278.1 | *Hirundo atrocaerulea* |
| Hirundo | NC_048497.1 | *Hirundo tahitica* |
| Hirundo | NC_036345.1 | *Sterna hirundo* |
| Hirundo | NC_050297.1 | *Hirundo rustica tytleri* |
| Hirundo | NC_050295.1 | *Hirundo rustica transitiva* |
| Homininae | NC_037853.1 | *Gorilla beringei* |
| Homininae | NC_011137.1 | *Homo sapiens neanderthalensis* |
| Homininae | NC_013993.1 | *Homo sp. Altai* |
| Homininae | NC_012920.1 | *Homo sapiens* |
| Homininae | NC_023100.1 | *Homo heidelbergensis* |
| Homininae | NC_001644.1 | *Pan paniscus* |
| Homininae | NC_001643.1 | *Pan troglodytes* |
| Hominoidea | NC_033883.1 | *Hoolock leuconedys* |
| Hominoidea | NC_033882.1 | *Hoolock leuconedys* |
| Hominoidea | NC_014045.1 | *Hylobates pileatus* |
| Hominoidea | NC_014042.1 | *Hylobates agilis* |
| Hominoidea | NC_002082.1 | *Hylobates lar* |
| Hominoidea | NC_021957.1 | *Nomascus leucogenys* |

| | | |
|---|---|---|
| Hominoidea | NC_018753.1 | *Nomascus gabriellae* |
| Hominoidea | NC_014047.1 | *Symphalangus syndactylus* |
| Hominoidea | NC_002083.1 | *Pongo abelii* |
| Hominoidea | NC_001646.1 | *Pongo pygmaeus* |
| Hydropsychoidea | NC_036952.1 | *Cheumatopsyche speciosa* |
| Hydropsychoidea | NC_036156.1 | *Hydromanicus wulaianus* |
| Hydropsychoidea | NC_060325.1 | *Hydropsyche fryeri* |
| Hydropsychoidea | NC_036951.1 | *Hydropsyche orris* |
| Hydropsychoidea | NC_036950.1 | *Hydropsyche simulans* |
| Hydropsychoidea | NC_036953.1 | *Potamyia flava* |
| Hymenopodidae | NC_057070.1 | *Amorphoscelis hainana* |
| Hymenopodidae | NC_030268.1 | *Anaxarcha zhengi* |
| Hymenopodidae | NC_045876.1 | *Psychomantis borneensis* |
| Hymenopodidae | NC_037234.1 | *Creobroter jiangxiensis* |
| Hymenopodidae | NC_030267.1 | *Creobroter gemmatus* |
| Hymenopodidae | NC_037235.1 | *Sibylla pretiosa* |
| Hymenoptera | NC_012689.1 | *Orussus occidentalis* |
| Hymenoptera | NC_040123.1 | *Tremex columba* |
| Hymenoptera | NC_042445.1 | *Labriocimbex sinicus* |
| Hymenoptera | NC_050309.1 | *Praia tianmunica* |
| Hymenoptera | NC_029733.1 | *Trichiosoma anthracinum* |
| Hymenoptera | NC_045902.1 | *Macroxyela ferruginea* |
| Hymenoptera | NC_057632.1 | *Sinopoppia nigroflagella* |

| | | |
|---|---|---|
| Hymenoptera | NC_057102.1 | *Analcellicampa danfengensis* |
| Hymenoptera | NC_059910.1 | *Neostromboceros nipponicus* |
| Hymenoptera | NC_058200.1 | *Conaspidia wangi* |
| Hynobius | NC_045210.1 | *Hynobius unisacculus* |
| Hynobius | NC_026033.1 | *Hynobius nigrescens* |
| Hynobius | NC_026032.1 | *Hynobius kimurae* |
| Hynobius | NC_023789.1 | *Hynobius maoershanensis* |
| Hynobius | NC_020650.1 | *Hynobius nebulosus* |
| Hynobius | NC_020649.1 | *Hynobius yiwuensis* |
| Hynobius | NC_013825.1 | *Hynobius yangi* |
| Hynobius | NC_013762.1 | *Hynobius guabangshanensis* |
| Hynobius | NC_010224.1 | *Hynobius quelpaertensis* |
| Hynobius | NC_009335.1 | *Hynobius arisanensis* |
| Hynobius | NC_008084.1 | *Hynobius formosanus* |
| Hynobius | NC_008076.1 | *Hynobius amjiensis* |
| Hynobius | NC_008079.1 | *Hynobius leechii* |
| Hypsiglena | NC_013992.1 | *Hypsiglena torquata* |
| Hypsiglena | NC_013987.1 | *Hypsiglena slevini* |
| Hypsiglena | NC_013982.1 | *Hypsiglena sp.* |
| Hypsiglena | NC_013977.1 | *Hypsiglena chlorophaea chlorophaea* |
| Hypsiglena | NC_013975.1 | *Hypsiglena jani texana* |
| Hypsiglena | NC_013984.1 | *Hypsiglena ochrorhyncha klauberi* |
| Hypsiglena | NC_013983.1 | *Hypsiglena ochrorhyncha nuchalata* |

| | | |
|---|---|---|
| Hypsiglena | NC_013980.1 | *Hypsiglena ochrorhyncha ochrorhyncha* |
| Hypsiglena | NC_024164.1 | *Hypsiglena unaocularus* |
| Hystricomorpha | NC_027742.1 | *Fukomys damarensis* |
| Hystricomorpha | NC_015112.1 | *Heterocephalus glaber* |
| Hystricomorpha | NC_047305.1 | *Geocapromys ingrahami* |
| Hystricomorpha | NC_000884.1 | *Cavia porcellus* |
| Hystricomorpha | NC_046949.1 | *Cavia aperea* |
| Hystricomorpha | NC_021386.1 | *Chinchilla lanigera* |
| Hystricomorpha | NC_020659.1 | *Ctenomys leucodon* |
| Hystricomorpha | NC_020658.1 | *Ctenomys sociabilis* |
| Hystricomorpha | NC_029876.1 | *Dactylomys dactylinus* |
| Hystricomorpha | NC_037778.1 | *Echimys saturnus* |
| Hystricomorpha | NC_037779.1 | *Makalata macrura* |
| Hystricomorpha | NC_037781.1 | *Pattonomys semivillosus* |
| Hystricomorpha | NC_037780.1 | *Pattonomys punctatus* |
| Hystricomorpha | NC_037783.1 | *Toromys rhipidurus* |
| Hystricomorpha | NC_037782.1 | *Toromys albiventris* |
| Hystricomorpha | NC_021387.1 | *Coendou insidiosus* |
| Hystricomorpha | NC_050263.1 | *Hystrix brachyura* |
| Hystricomorpha | NC_035820.1 | *Trichys fasciculata* |
| Hystricomorpha | NC_030184.1 | *Laonastes aenigmamus* |
| Hystricomorpha | NC_035866.1 | *Myocastor coypus* |
| Hystricomorpha | NC_020661.1 | *Octodon degus* |

| | | |
|---|---|---|
| Hystricomorpha | NC_020660.1 | *Spalacopus cyanus* |
| Hystricomorpha | NC_020792.1 | *Tympanoctomys barrerae* |
| Hystricomorpha | NC_002658.1 | *Thryonomys swinderianus* |
| Isotomidae | NC_037610.1 | *Cryptopygus terranovus* |
| Isotomidae | NC_010533.1 | *Cryptopygus antarcticus* |
| Isotomidae | NC_024155.1 | *Folsomotoma octooculata* |
| Isotomidae | NC_056909.1 | *Kaylathalia klovstadi* |
| Ixodes | NC_062061.1 | *Ixodes ovatus* |
| Ixodes | NC_062629.1 | *Ixodes confusus* |
| Ixodes | NC_062626.1 | *Ixodes barkeri* |
| Ixodes | NC_062628.1 | *Ixodes fecialis* |
| Ixodes | NC_062627.1 | *Ixodes woyliei* |
| Ixodes | NC_062625.1 | *Ixodes australiensis* |
| Ixodes | NC_062633.1 | *Ixodes trichosuri* |
| Ixodes | NC_062632.1 | *Ixodes myrmecobii* |
| Ixodes | NC_062631.1 | *Ixodes hirsti* |
| Ixodes | NC_062630.1 | *Ixodes cornuatus* |
| Ixodes | NC_062157.1 | *Ixodes kuntzi* |
| Ixodes | NC_062062.1 | *Ixodes nuttallianus* |
| Ixodes | NC_062060.1 | *Ixodes simplex* |
| Ixodes | NC_062059.1 | *Ixodes sinensis* |
| Ixodes | NC_061226.1 | *Ixodes granulatus* |
| Ixodes | NC_061225.1 | *Ixodes acutitarsus* |

| | | |
|---|---|---|
| Ixodes | NC_058244.1 | *Ixodes vespertilionis* |
| Ixodes | NC_058242.1 | *Ixodes nipponensis* |
| Ixodes | NC_041086.1 | *Ixodes tasmani* |
| Ixodes | NC_023831.1 | *Ixodes pavlovskyi* |
| Ixodes | NC_018369.2 | *Ixodes ricinus* |
| Ixodes | NC_006078.1 | *Ixodes uriae* |
| Ixodes | NC_005293.1 | *Ixodes holocyclus* |
| Ixodes | NC_004370.1 | *Ixodes persulcatus* |
| Junonia | NC_064714.1 | *Junonia zonalis* |
| Junonia | NC_064713.1 | *Junonia westermanni* |
| Junonia | NC_064712.1 | *Junonia hierta* |
| Junonia | NC_064708.1 | *Junonia oenone* |
| Junonia | NC_064707.1 | *Junonia sophia* |
| Junonia | NC_064706.1 | *Junonia terea* |
| Junonia | NC_064705.1 | *Junonia touhilimasa* |
| Junonia | NC_064704.1 | *Junonia villida* |
| Junonia | NC_064703.1 | *Junonia nigrosuffusa* |
| Junonia | NC_064702.1 | *Junonia natalica* |
| Junonia | NC_064701.1 | *Junonia intermedia* |
| Junonia | NC_064700.1 | *Junonia cytora* |
| Junonia | NC_064699.1 | *Junonia hedonia* |
| Junonia | NC_064698.1 | *Junonia goudoti* |
| Junonia | NC_064697.1 | *Junonia erigone* |

| | | |
|---|---|---|
| Junonia | NC_064696.1 | *Junonia chorimene* |
| Junonia | NC_064695.1 | *Junonia atlites* |
| Junonia | NC_064693.1 | *Junonia adulatrix* |
| Junonia | NC_064692.1 | *Junonia evarete* |
| Junonia | NC_024407.1 | *Junonia almana* |
| Junonia | NC_022697.1 | *Junonia orithya* |
| Junonia | NC_033861.1 | *Junonia iphita* |
| Junonia | NC_028324.1 | *Junonia lemonias* |
| Junonia | NC_037710.1 | *Junonia rhadama* |
| Junonia | NC_034663.1 | *Junonia litoralis* |
| Lacertibaenia | NC_059775.1 | *Acanthodactylus aureus* |
| Lacertibaenia | NC_059773.1 | *Acanthodactylus erythrurus* |
| Lacertibaenia | NC_059772.1 | *Acanthodactylus boskianus* |
| Lacertibaenia | NC_059782.1 | *Acanthodactylus schmidti* |
| Lacertibaenia | NC_059781.1 | *Acanthodactylus guineensis* |
| Lacertibaenia | NC_059780.1 | *Algyroides nigropunctatus* |
| Lacertibaenia | NC_059777.1 | *Australolacerta australis* |
| Lacertibaenia | NC_028440.1 | *Lacerta bilineata* |
| Lacertibaenia | NC_021766.1 | *Lacerta agilis* |
| Lacertibaenia | NC_008328.1 | *Lacerta viridis viridis* |
| Lacertibaenia | NC_059779.1 | *Meroles squamulosus* |
| Lacertibaenia | NC_059774.1 | *Mesalina olivieri* |
| Lacertibaenia | NC_059778.1 | *Pedioplanis laticeps* |

| | | |
|---|---|---|
| Lacertibaenia | NC_011606.1 | *Phoenicolacerta kulzeri* |
| Lacertibaenia | NC_011607.1 | *Podarcis muralis* |
| Lacertibaenia | NC_011609.1 | *Podarcis siculus* |
| Lacertibaenia | NC_030209.1 | *Takydromus amurensis* |
| Lacertibaenia | NC_022703.1 | *Takydromus sexlineatus* |
| Lacertibaenia | NC_018777.1 | *Takydromus wolteri* |
| Lacertibaenia | NC_026867.1 | *Zootoca vivipara* |
| Lacertibaenia | NC_059771.1 | *Gallotia atlantica* |
| Lacertibaenia | NC_059776.1 | *Psammodromus algirus* |
| Lampyridae | NC_044776.1 | *Abscondita terminalis* |
| Lampyridae | NC_039706.1 | *Abscondita anceyi* |
| Lampyridae | NC_035755.1 | *Aquatica lateralis* |
| Lampyridae | NC_035060.1 | *Aquatica ficta* |
| Lampyridae | NC_035061.1 | *Aquatica wuhana* |
| Lampyridae | NC_032062.1 | *Asymmetricata circumdata* |
| Lampyridae | NC_058281.1 | *Curtos fulvocapitalis* |
| Lampyridae | NC_044789.1 | *Curtos bilineatus* |
| Lampyridae | NC_051869.1 | *Diaphanes citrinus* |
| Lampyridae | NC_044793.1 | *Diaphanes pectinealis* |
| Lampyridae | NC_044791.1 | *Diaphanes mendax* |
| Lampyridae | NC_044787.1 | *Diaphanes nubilus* |
| Lampyridae | NC_044778.1 | *Emeia pseudosauteri* |
| Lampyridae | NC_050947.1 | *Hotaria unmunsana* |

| | | |
|---|---|---|
| Lampyridae | NC_039700.1 | *Inflata indica* |
| Lampyridae | NC_044786.1 | *Lamprigera yunnana* |
| Lampyridae | NC_038225.1 | *Luciola curtithorax* |
| Lampyridae | NC_027176.1 | *Luciola substriata* |
| Lampyridae | NC_022472.1 | *Luciola cruciata* |
| Lampyridae | NC_036353.1 | *Pteroptyx maipo* |
| Lampyridae | NC_057261.1 | *Pygoluciola qingyu* |
| Lampyridae | NC_044792.1 | *Pyrocoelia thibetana* |
| Lampyridae | NC_044790.1 | *Pyrocoelia praetexta* |
| Lampyridae | NC_003970.1 | *Pyrocoelia rufa* |
| Lampyridae | NC_044788.1 | *Vesta saturnalis* |
| Lepadoidea | NC_066685.1 | *Paralepas cf. quadrata* |
| Lepadoidea | NC_062431.1 | *Lepas anatifera* |
| Lepadoidea | NC_026576.1 | *Lepas anserifera* |
| Lepadoidea | NC_025295.1 | *Lepas australis* |
| Lepadoidea | NC_066679.1 | *Glyptelasma gigas* |
| Lepadoidea | NC_043898.1 | *Glyptelasma annandalei* |
| Lepadoidea | NC_066683.1 | *Octolasmis warwickii* |
| Lepadoidea | NC_066680.1 | *Poecilasma litum* |
| Lepilemur | NC_034731.1 | *Lepilemur seali* |
| Lepilemur | NC_034727.1 | *Lepilemur septentrionalis* |
| Lepilemur | NC_034721.1 | *Lepilemur dorsalis* |
| Lepilemur | NC_034739.1 | *Lepilemur microdon* |

| | | |
|---|---|---|
| Lepilemur | NC_034738.1 | *Lepilemur hollandorum* |
| Lepilemur | NC_034737.1 | *Lepilemur scottorum* |
| Lepilemur | NC_034736.1 | *Lepilemur jamesi* |
| Lepilemur | NC_034735.1 | *Lepilemur tymerlachsoni* |
| Lepilemur | NC_034734.1 | *Lepilemur sahamalazensis* |
| Lepilemur | NC_034733.1 | *Lepilemur ahmansoni* |
| Lepilemur | NC_034732.1 | *Lepilemur wrighti* |
| Lepilemur | NC_034730.1 | *Lepilemur aeeclis* |
| Lepilemur | NC_034729.1 | *Lepilemur grewcocki* |
| Lepilemur | NC_034726.1 | *Lepilemur betsileo* |
| Lepilemur | NC_034725.1 | *Lepilemur milanoii* |
| Lepilemur | NC_034724.1 | *Lepilemur otto* |
| Lepilemur | NC_034723.1 | *Lepilemur petteri* |
| Lepilemur | NC_034722.1 | *Lepilemur randrianasoloi* |
| Lepilemur | NC_034720.1 | *Lepilemur edwardsi* |
| Lepilemur | NC_034719.1 | *Lepilemur leucopus* |
| Lepilemur | NC_034718.1 | *Lepilemur fleuretae* |
| Lepilemur | NC_034717.1 | *Lepilemur ankaranensis* |
| Lepilemur | NC_021953.1 | *Lepilemur ruficaudatus* |
| Lepilemur | NC_014453.1 | *Lepilemur hubbardorum* |
| Lepus | NC_065133.1 | *Lepus tibetanus* |
| Lepus | NC_050983.1 | *Lepus oiostolus* |
| Lepus | NC_050569.1 | *Lepus yarkandensis* |

| | | |
|---|---|---|
| Lepus | NC_044769.1 | *Lepus arcticus* |
| Lepus | NC_025902.1 | *Lepus hainanus* |
| Lepus | NC_025748.1 | *Lepus tolai* |
| Lepus | NC_025316.1 | *Lepus sinensis* |
| Lepus | NC_024259.1 | *Lepus coreanus* |
| Lepus | NC_015841.1 | *Lepus capensis* |
| Lepus | NC_004028.1 | *Lepus europaeus* |
| Lepus | NC_024042.1 | *Lepus granatensis* |
| Lepus | NC_024043.1 | *Lepus americanus* |
| Lepus | NC_024040.1 | *Lepus timidus* |
| Lepus | NC_024041.1 | *Lepus townsendii* |
| Lethe | NC_050916.1 | *Lethe verma* |
| Lethe | NC_050915.1 | *Lethe uemurai* |
| Lethe | NC_050914.1 | *Lethe titania* |
| Lethe | NC_050913.1 | *Lethe syrcis* |
| Lethe | NC_050912.1 | *Lethe satyrina* |
| Lethe | NC_050911.1 | *Lethe oculatissima* |
| Lethe | NC_050910.1 | *Lethe nigrifascia* |
| Lethe | NC_050909.1 | *Lethe marginalis* |
| Lethe | NC_050908.1 | *Lethe helle* |
| Lethe | NC_050907.1 | *Lethe hayashii* |
| Lethe | NC_050906.1 | *Lethe baucis* |
| Lethe | NC_050905.1 | *Lethe baileyi* |

| | | |
|---|---|---|
| Lethe | NC_028507.1 | *Lethe albolineata* |
| Limenitidinae | NC_039696.1 | *Adelpha iphiclus* |
| Limenitidinae | NC_039885.1 | *Adelpha bredowii* |
| Limenitidinae | NC_039858.1 | *Adelpha ethelda* |
| Limenitidinae | NC_039882.1 | *Auzakia danava* |
| Limenitidinae | NC_039861.1 | *Chalinga pratti* |
| Limenitidinae | NC_050060.1 | *Lelecella limenitoides* |
| Limenitidinae | NC_039877.1 | *Litinga cottini* |
| Limenitidinae | NC_039874.1 | *Litinga mimica* |
| Limenitidinae | NC_060742.1 | *Neptis thisbe* |
| Limenitidinae | NC_060741.1 | *Neptis obscurior* |
| Limenitidinae | NC_060739.1 | *Neptis raddei* |
| Limenitidinae | NC_038154.1 | *Neptis alwina* |
| Limenitidinae | NC_024401.1 | *Neptis soma* |
| Limenitidinae | NC_024419.1 | *Neptis philyra* |
| Limenitidinae | NC_025759.1 | *Neptis clinia* |
| Limenitidinae | NC_024402.1 | *Pantoporia hordonia* |
| Limenitidinae | NC_039698.1 | *Parasarpa zayla* |
| Limenitidinae | NC_039875.1 | *Parasarpa albomaculata* |
| Limenitidinae | NC_024405.1 | *Parasarpa dudu* |
| Limenitidinae | NC_036332.1 | *Phaedyma columella* |
| Limenitidinae | NC_039887.1 | *Tacola larymna* |
| Limenitidinae | NC_039864.1 | *Tacola eulimene* |

| | | |
|---|---|---|
| Limenitis | NC_065070.1 | *Limenitis sulpitia* |
| Limenitis | NC_039884.1 | *Limenitis lorquini* |
| Limenitis | NC_039881.1 | *Limenitis cleophas* |
| Limenitis | NC_039878.1 | *Limenitis populi* |
| Limenitis | NC_039871.1 | *Limenitis ciocolatina* |
| Limenitis | NC_039869.1 | *Limenitis arthemis* |
| Limenitis | NC_039868.1 | *Limenitis archippus* |
| Limenitis | NC_039867.1 | *Limenitis elwesi* |
| Limenitis | NC_039866.1 | *Limenitis camilla* |
| Limenitis | NC_039865.1 | *Limenitis reducta* |
| Limenitis | NC_039863.1 | *Limenitis glorifica* |
| Limenitis | NC_039862.1 | *Limenitis weidemeyerii* |
| Limenitis | NC_034754.1 | *Limenitis helmanni* |
| Limenitis | NC_034235.1 | *Limenitis amphyssa* |
| Limenitis | NC_034234.1 | *Limenitis moltrechti* |
| Limenitis | NC_034233.1 | *Limenitis sydyi* |
| Lonchura | NC_036401.1 | *Lonchura grandis* |
| Lonchura | NC_036396.1 | *Lonchura leucosticta* |
| Lonchura | NC_029475.1 | *Lonchura striata* |
| Lonchura | NC_028036.1 | *Lonchura punctulata* |
| Lorisiformes | NC_021949.1 | *Galago moholi* |
| Lorisiformes | NC_012761.1 | *Galago senegalensis* |
| Lorisiformes | NC_012762.1 | *Otolemur crassicaudatus* |

| | | |
|---|---|---|
| Lorisiformes | NC_021955.1 | *Loris lydekkerianus* |
| Lorisiformes | NC_012763.1 | *Loris tardigradus* |
| Lorisiformes | NC_040292.1 | *Nycticebus coucang insularis* |
| Lorisiformes | NC_033381.1 | *Nycticebus pygmaeus* |
| Lorisiformes | NC_021958.1 | *Nycticebus bengalensis* |
| Lucanidae | NC_045068.1 | *Cyclommatus strigiceps vitalisi* |
| Lucanidae | NC_054278.1 | *Dorcus koreanus* |
| Lucanidae | NC_043928.1 | *Dorcus hansi* |
| Lucanidae | NC_045124.1 | *Dorcus tenuihirsutus* |
| Lucanidae | NC_045123.1 | *Dorcus ursulus* |
| Lucanidae | NC_045102.1 | *Figulus binodulus* |
| Lucanidae | NC_060603.1 | *Lucanus prometheus* |
| Lucanidae | NC_044961.1 | *Lucanus fortunei* |
| Lucanidae | NC_038212.1 | *Macrodorcas seguyi* |
| Lucanidae | NC_039652.1 | *Neolucanus maximus* |
| Lucanidae | NC_065111.1 | *Nigidius sinicus* |
| Lucanidae | NC_063663.1 | *Nigidius miwai* |
| Lucanidae | NC_044962.1 | *Prismognathus prossi* |
| Lucanidae | NC_065362.1 | *Prosopocoilus laterotarsus* |
| Lucanidae | NC_050851.1 | *Prosopocoilus astacoides* |
| Lucanidae | NC_027580.1 | *Prosopocoilus gracilis* |
| Lucanidae | NC_036038.1 | *Prosopocoilus confucius* |
| Lucanidae | NC_044096.1 | *Serrognathus platymelus* |

| | | |
|---|---|---|
| Luperina | NC_063914.1 | *Monolepta signata* |
| Luperina | NC_045838.1 | *Monolepta occifluvis* |
| Luperina | NC_039711.1 | *Monolepta quadriguttata* |
| Luperina | NC_057489.1 | *Monolepta hieroglyphica* |
| Lycaenidae | NC_065068.1 | *Curetis acuta* |
| Lycaenidae | NC_023087.1 | *Lycaena phlaeas* |
| Lycaenidae | NC_020779.1 | *Cupido argiades* |
| Lycaenidae | NC_065461.1 | *Jamides bochus* |
| Lycaenidae | NC_029763.1 | *Shijimiaeoides divina* |
| Lycaenidae | NC_063699.1 | *Ahlbergia frivaldszkyi* |
| Lycaenidae | NC_063698.1 | *Ahlbergia ferrea* |
| Lycaenidae | NC_007976.1 | *Coreana raphaelis* |
| Lycosoidea | NC_058921.1 | *Phoneutria boliviensis* |
| Lycosoidea | NC_065748.1 | *Lycosa singoriensis* |
| Lycosoidea | NC_065747.1 | *Lycosa shansia* |
| Lycosoidea | NC_064110.1 | *Pardosa pusiola* |
| Lycosoidea | NC_025223.1 | *Pardosa laura* |
| Lycosoidea | NC_026123.1 | *Wadicosa fidelis* |
| Lycosoidea | NC_046736.1 | *Oxyopes hupingensis* |
| Lycosoidea | NC_025224.1 | *Oxyopes sertatus* |
| Lycosoidea | NC_053648.1 | *Oxyopes licenti* |
| Lycosoidea | NC_031355.1 | *Dolomedes angustivirgatus* |
| Macaca | NC_012670.1 | *Macaca fascicularis* |

| | | |
|---|---|---|
| Macaca | NC_005943.1 | *Macaca mulatta* |
| Macaca | NC_037466.1 | *Macaca mulatta vestita* |
| Macaca | NC_037437.1 | *Macaca sinica* |
| Macaca | NC_031156.1 | *Macaca leucogenys* |
| Macaca | NC_027604.1 | *Macaca leonina* |
| Macaca | NC_027449.1 | *Macaca cyclopis* |
| Macaca | NC_026976.1 | *Macaca nemestrina* |
| Macaca | NC_026120.1 | *Macaca nigra* |
| Macaca | NC_025513.1 | *Macaca fuscata* |
| Macaca | NC_025222.1 | *Macaca tonkeana* |
| Macaca | NC_025221.1 | *Macaca silenus* |
| Macaca | NC_025201.1 | *Macaca arctoides* |
| Macaca | NC_023795.1 | *Macaca assamensis* |
| Macaca | NC_011519.1 | *Macaca thibetana* |
| Macaca | NC_002764.1 | *Macaca sylvanus* |
| Macroglossinae | NC_062879.1 | *Theretra clotho* |
| Macroglossinae | NC_062187.1 | *Theretra alecto* |
| Macrotermes | NC_034288.1 | *Macrotermes yunnanensis* |
| Macrotermes | NC_034110.1 | *Macrotermes gilvus* |
| Macrotermes | NC_034050.1 | *Macrotermes falciger* |
| Macrotermes | NC_034046.1 | *Macrotermes carbonarius* |
| Macrotermes | NC_034127.1 | *Macrotermes muelleri* |
| Macrotermes | NC_034078.1 | *Macrotermes annandalei* |

| | | |
|---|---|---|
| Macrotermes | NC_034054.1 | *Macrotermes vitrialatus* |
| Macrotermes | NC_018599.1 | *Macrotermes barneyi* |
| Macrotermes | NC_025522.1 | *Macrotermes natalensis* |
| Macrotermes | NC_018128.1 | *Macrotermes subhyalinus* |
| Majoidea | NC_052726.1 | *Chionoecetes japonicus* |
| Majoidea | NC_035425.1 | *Maja squinado* |
| Majoidea | NC_035424.1 | *Maja crispata* |
| Majoidea | NC_025518.1 | *Damithrax spinosissimus* |
| Majoidea | NC_057204.1 | *Oregonia gracilis* |
| Majoidea | NC_057485.1 | *Scyra compressipes* |
| Manis | NC_060494.1 | *Manis pentadactyla pentadactyla* |
| Manis | NC_036434.1 | *Manis culionensis* |
| Manis | NC_036433.1 | *Manis crassicaudata* |
| Manis | NC_036064.1 | *Manis gigantea* |
| Manis | NC_026781.1 | *Manis javanica* |
| Manis | NC_026780.1 | *Manis tricuspis* |
| Manis | NC_025769.1 | *Manis temminckii* |
| Mantidae | NC_065758.1 | *Hierodula maculata* |
| Mantidae | NC_065746.1 | *Hierodula zhangi* |
| Mantidae | NC_065745.1 | *Hierodula chinensis* |
| Mantidae | NC_048984.1 | *Hierodula membranacea* |
| Mantidae | NC_034283.1 | *Hierodula patellifera* |
| Mantidae | NC_029326.1 | *Hierodula formosana* |

| | | |
|---|---|---|
| Mantidae | NC_007702.1 | *Tamolanica tamolana* |
| Mantidae | NC_037207.1 | *Schizocephala bicornis* |
| Mantidae | NC_030265.1 | *Mantis religiosa* |
| Mantidae | NC_065761.1 | *Statilia flavobrunnea* |
| Mantidae | NC_065760.1 | *Statilia maculata* |
| Mantidae | NC_037204.1 | *Sphodromantis lineola* |
| Mantidae | NC_065759.1 | *Tenodera angustipennis* |
| Mantidae | NC_030266.1 | *Tenodera sinensis* |
| Martes | NC_020664.1 | *Martes pennanti* |
| Martes | NC_020643.1 | *Martes foina* |
| Martes | NC_020642.1 | *Martes americana* |
| Martes | NC_012141.1 | *Martes flavigula* |
| Martes | NC_011579.1 | *Martes zibellina* |
| Martes | NC_021749.1 | *Martes martes* |
| Martes | NC_009678.1 | *Martes melampus* |
| Megachiroptera | NC_046901.1 | *Casinycteris ophiodon* |
| Megachiroptera | NC_046900.1 | *Casinycteris campomaanensis* |
| Megachiroptera | NC_046899.1 | *Casinycteris argynnis* |
| Megachiroptera | NC_046913.1 | *Hypsignathus monstrosus* |
| Megachiroptera | NC_046923.1 | *Nanonycteris veldkampii* |
| Megachiroptera | NC_046925.1 | *Plerotes anchietae* |
| Megachiroptera | NC_046932.1 | *Scotonycteris zenkeri* |
| Megachiroptera | NC_046931.1 | *Scotonycteris zenkeri occidentalis* |

| | | |
|---|---|---|
| Megachiroptera | NC_046930.1 | *Scotonycteris bergmansi* |
| Megachiroptera | NC_046933.1 | *Sphaerias blanfordi* |
| Megachiroptera | NC_046904.1 | *Eonycteris spelaea* |
| Megachiroptera | NC_046914.1 | *Macroglossus sobrinus* |
| Meliphagoidea | NC_029139.1 | *Gerygone igata* |
| Meliphagoidea | NC_053100.1 | *Origma solitaria* |
| Meliphagoidea | NC_053097.1 | *Dasyornis broadbenti* |
| Meliphagoidea | NC_053095.1 | *Malurus elegans* |
| Meliphagoidea | NC_024873.1 | *Malurus melanocephalus* |
| Meliphagoidea | NC_051552.1 | *Acanthorhynchus tenuirostris* |
| Meliphagoidea | NC_019664.1 | *Epthianura albifrons* |
| Meliphagoidea | NC_053098.1 | *Grantiella picta* |
| Meliphagoidea | NC_062733.1 | *Lichenostomus cassidix* |
| Meliphagoidea | NC_029144.1 | *Prosthemadera novaeseelandiae* |
| Meliphagoidea | NC_053099.1 | *Pardalotus punctatus* |
| Merostomata | NC_019623.1 | *Carcinoscorpius rotundicauda* |
| Merostomata | NC_003057.1 | *Limulus polyphemus* |
| Merostomata | NC_052701.1 | *Tachypleus gigas* |
| Merostomata | NC_012574.1 | *Tachypleus tridentatus* |
| Metatheria | NC_056379.1 | *Antechinus flavipes Adam Ant* |
| Metatheria | NC_050988.1 | *Antechinus arktos AA100* |
| Metatheria | NC_007630.1 | *Dasyurus hallucatus* |
| Metatheria | NC_045217.1 | *Murexia melanurus* |

| | | |
|---|---|---|
| Metatheria | NC_006523.1 | *Phascogale tapoatafa* |
| Metatheria | NC_061372.1 | *Phascolosorex doriae* |
| Metatheria | NC_018788.1 | *Sarcophilus harrisii* |
| Metatheria | NC_007631.1 | *Sminthopsis crassicaudata* |
| Metatheria | NC_006517.1 | *Sminthopsis douglasi* |
| Metatheria | NC_011949.1 | *Myrmecobius fasciatus* |
| Metatheria | NC_011944.1 | *Thylacinus cynocephalus* |
| Metatheria | NC_005826.1 | *Dromiciops gliroides* |
| Metatheria | NC_006522.1 | *Notoryctes typhlops* |
| Metatheria | NC_005828.1 | *Caenolestes fuliginosus* |
| Metatheria | NC_005829.1 | *Rhyncholestes raphanurus* |
| Metatheria | NC_002746.1 | *Isoodon macrourus* |
| Metatheria | NC_006520.1 | *Macrotis lagotis* |
| Metatheria | NC_007632.1 | *Echymipera rufescens australis* |
| Microcebus | NC_035630.1 | *Microcebus simmonsi* |
| Microcebus | NC_035629.1 | *Microcebus berthae* |
| Microcebus | NC_035603.1 | *Microcebus margotmarshae* |
| Microcebus | NC_035602.1 | *Microcebus lehilahytsara* |
| Microcebus | NC_035600.1 | *Microcebus danfossi* |
| Microcebus | NC_035564.1 | *Microcebus jollyae* |
| Microcebus | NC_035563.1 | *Microcebus mittermeieri* |
| Microcebus | NC_035560.1 | *Microcebus tavaratra* |
| Microcebus | NC_035559.1 | *Microcebus bongolavensis* |

| | | |
|---|---|---|
| Microcebus | NC_035557.1 | *Microcebus myoxinus* |
| Microcebus | NC_035556.1 | *Microcebus ravelobensis* |
| Microcebus | NC_035555.1 | *Microcebus tanosi* |
| Microcebus | NC_035554.1 | *Microcebus arnholdi* |
| Microcebus | NC_035601.1 | *Microcebus mamiratra* |
| Microcebus | NC_035561.1 | *Microcebus sambiranensis* |
| Microcebus | NC_035558.1 | *Microcebus griseorufus* |
| Microcebus | NC_035562.1 | *Microcebus rufus* |
| Microcebus | NC_028718.1 | *Microcebus murinus* |
| Microchiroptera | NC_002626.1 | *Chalinolobus tuberculatus* |
| Microchiroptera | NC_057092.1 | *Eudiscopus denticulus* |
| Microchiroptera | NC_029939.1 | *Hypsugo alaschanicus* |
| Microchiroptera | NC_061569.1 | *Kerivoula minuta* |
| Microchiroptera | NC_061567.1 | *Kerivoula papillosa* |
| Microchiroptera | NC_050995.1 | *Lasionycteris noctivagans* |
| Microchiroptera | NC_016873.1 | *Lasiurus borealis* |
| Microchiroptera | NC_025949.1 | *Murina leucogaster* |
| Microchiroptera | NC_021119.1 | *Murina ussuriensis* |
| Microchiroptera | NC_060309.1 | *Nyctalus aviator* |
| Microchiroptera | NC_041160.1 | *Nyctalus plancyi* |
| Microchiroptera | NC_029191.1 | *Pipistrellus coromandra* |
| Microchiroptera | NC_005436.1 | *Pipistrellus abramus* |
| Microchiroptera | NC_027977.1 | *Plecotus macrobullaris* |

| | | |
|---|---|---|
| Microchiroptera | NC_016872.1 | *Plecotus rafinesquii* |
| Microchiroptera | NC_015484.1 | *Plecotus auritus* |
| Microchiroptera | NC_065469.1 | *Submyotodon moupinensis* |
| Microchiroptera | NC_033347.1 | *Vespertilio murinus* |
| Microchiroptera | NC_024558.1 | *Vespertilio sinensis* |
| Microchiroptera | NC_022421.1 | *Brachyphylla cavernarum* |
| Microchiroptera | NC_066073.1 | *Carollia brevicauda* |
| Microchiroptera | NC_065677.1 | *Carollia castanea* |
| Microchiroptera | NC_022422.1 | *Carollia perspicillata* |
| Microchiroptera | NC_022426.1 | *Rhinophylla pumilio* |
| Microchiroptera | NC_022423.1 | *Desmodus rotundus* |
| Microchiroptera | NC_037133.1 | *Diaemus youngi* |
| Microchiroptera | NC_037138.1 | *Diphylla ecaudata* |
| Microchiroptera | NC_048476.1 | *Furipterus horrens* |
| Microchiroptera | NC_061963.1 | *Hipposideros pomona* |
| Microchiroptera | NC_061570.1 | *Hipposideros larvatus* |
| Microchiroptera | NC_061568.1 | *Hipposideros cervinus* |
| Microchiroptera | NC_018540.1 | *Hipposideros armiger* |
| Microchiroptera | NC_037134.1 | *Hsunycteris thomasi* |
| Microchiroptera | NC_065688.1 | *Lonchophylla concava* |
| Microchiroptera | NC_065684.1 | *Lonchophylla robusta* |
| Microchiroptera | NC_057635.1 | *Macroderma gigas* |
| Microchiroptera | NC_044489.1 | *Miniopterus fuliginosus* |

| | | |
|---|---|---|
| Microchiroptera | NC_036331.1 | *Tadarida latouchei* |
| Microchiroptera | NC_036330.1 | *Tadarida teniotis* |
| Microchiroptera | NC_033353.1 | *Pteronotus personatus* |
| Microchiroptera | NC_023368.1 | *Pteronotus parnellii* |
| Microchiroptera | NC_022425.1 | *Pteronotus rubiginosus* |
| Microchiroptera | NC_006925.1 | *Mystacina tuberculata* |
| Microchiroptera | NC_037137.1 | *Noctilio leporinus* |
| Microchiroptera | NC_036421.1 | *Saccopteryx leptura* |
| Microhylidae | NC_045110.1 | *Microhyla fissipes* |
| Microhylidae | NC_039176.1 | *Microhyla taraiensis* |
| Microhylidae | NC_038130.1 | *Microhyla mixtura* |
| Microhylidae | NC_030049.1 | *Microhyla butleri* |
| Microhylidae | NC_024547.1 | *Microhyla pulchra* |
| Microhylidae | NC_006406.1 | *Microhyla heymonsi* |
| Microhylidae | NC_010233.1 | *Microhyla okinavensis* |
| Microhylidae | NC_009422.1 | *Microhyla ornata* |
| Microtus | NC_059928.1 | *Microtus montebelli* |
| Microtus | NC_057558.1 | *Microtus thomasi* |
| Microtus | NC_057557.1 | *Microtus chrotorrhinus* |
| Microtus | NC_057556.1 | *Microtus cabrerae* |
| Microtus | NC_049220.1 | *Microtus richardsoni* |
| Microtus | NC_041250.1 | *Microtus agrestis* |
| Microtus | NC_038176.1 | *Microtus arvalis* |

| | | |
|---|---|---|
| Microtus | NC_027945.1 | *Microtus ochrogaster* |
| Microtus | NC_015243.1 | *Microtus fortis calamorum* |
| Microtus | NC_015241.1 | *Microtus fortis fortis* |
| Microtus | NC_008064.1 | *Microtus levis* |
| Microtus | NC_003041.1 | *Microtus kikuchii* |
| Muridae | NC_049122.1 | *Apodemus sylvaticus* |
| Muridae | NC_019585.1 | *Apodemus latronum* |
| Muridae | NC_019584.1 | *Apodemus draco* |
| Muridae | NC_017599.1 | *Apodemus chevrieri* |
| Muridae | NC_016662.1 | *Apodemus chejuensis* |
| Muridae | NC_016428.1 | *Apodemus agrarius* |
| Muridae | NC_016060.1 | *Apodemus peninsulae* |
| Muridae | NC_053802.1 | *Arvicanthis rufinus* |
| Muridae | NC_053801.1 | *Arvicanthis somalicus* |
| Muridae | NC_057104.1 | *Bandicota bengalensis* |
| Muridae | NC_028335.1 | *Bandicota indica* |
| Muridae | NC_036730.1 | *Berylmys berdmorei* |
| Muridae | NC_036724.1 | *Bunomys penitus* |
| Muridae | NC_049121.1 | *Chiropodomys gliroides* |
| Muridae | NC_053812.1 | *Dasymys incomtus* |
| Muridae | NC_053808.1 | *Dephomys defua* |
| Muridae | NC_053805.1 | *Desmomys harringtoni* |
| Muridae | NC_053815.1 | *Golunda ellioti* |

| | | |
|---|---|---|
| Muridae | NC_053797.1 | *Grammomys dolichurus* |
| Muridae | NC_053798.1 | *Grammomys selousi* |
| Muridae | NC_053796.1 | *Grammomys surdaster* |
| Muridae | NC_036725.1 | *Halmaheramys bokimekot* |
| Muridae | NC_049120.1 | *Heimyscus fumosus* |
| Muridae | NC_053810.1 | *Hybomys trivirgatus* |
| Muridae | NC_053806.1 | *Hybomys univittatus* |
| Muridae | NC_053809.1 | *Hybomys planifrons* |
| Muridae | NC_065075.1 | *Hylomyscus parvus* |
| Muridae | NC_065074.1 | *Hylomyscus kerbispeterhansi* |
| Muridae | NC_065072.1 | *Hylomyscus alleni* |
| Muridae | NC_065071.1 | *Hylomyscus aeta* |
| Muridae | NC_065073.1 | *Hylomyscus endorobae* |
| Muridae | NC_053804.1 | *Lamottemys okuensis* |
| Muridae | NC_014696.1 | *Leggadina lakedownensis* |
| Muridae | NC_053800.1 | *Lemniscomys barbarus* |
| Muridae | NC_053799.1 | *Lemniscomys rosalia* |
| Muridae | NC_036731.1 | *Lenothrix canus* |
| Muridae | NC_035819.1 | *Leopoldamys sabanus* |
| Muridae | NC_025670.1 | *Leopoldamys edwardsi* |
| Muridae | NC_065079.1 | *Mastomys angolensis* |
| Muridae | NC_065077.1 | *Mastomys kollmannspergeri* |
| Muridae | NC_036995.1 | *Mastomys natalensis* |

| | | |
|---|---|---|
| Muridae | NC_036018.1 | *Mastomys coucha* |
| Muridae | NC_065076.1 | *Mastomys erythroleucus* |
| Muridae | NC_056988.1 | *Maxomys ochraceiventer* |
| Muridae | NC_049119.1 | *Maxomys whiteheadi* |
| Muridae | NC_036732.1 | *Maxomys surifer* |
| Muridae | NC_049118.1 | *Melomys burtoni* |
| Muridae | NC_053816.1 | *Micaelamys namaquensis* |
| Muridae | NC_060316.1 | *Micromys erythrotis* |
| Muridae | NC_027932.1 | *Micromys minutus* |
| Muridae | NC_053803.1 | *Mylomys dybowskii* |
| Muridae | NC_065081.1 | *Myomyscus verreauxii* |
| Muridae | NC_065080.1 | *Myomyscus brockmani* |
| Muridae | NC_065402.1 | *Niviventer lotipes* |
| Muridae | NC_060500.1 | *Niviventer andersoni* |
| Muridae | NC_056776.1 | *Niviventer fulvescens* |
| Muridae | NC_019617.1 | *Niviventer excelsior* |
| Muridae | NC_035822.1 | *Niviventer cremoriventer* |
| Muridae | NC_023960.1 | *Niviventer confucianus* |
| Muridae | NC_053811.1 | *Otomys typus* |
| Muridae | NC_049116.1 | *Otomys irroratus* |
| Muridae | NC_036726.1 | *Paruromys dominator* |
| Muridae | NC_065085.1 | *Praomys morio* |
| Muridae | NC_065084.1 | *Praomys minor* |

| | | |
|---|---|---|
| Muridae | NC_065083.1 | *Praomys jacksoni* |
| Muridae | NC_065082.1 | *Praomys hartwigi* |
| Muridae | NC_053817.1 | *Praomys delectorum* |
| Muridae | NC_049115.1 | *Praomys rostratus* |
| Muridae | NC_014698.1 | *Pseudomys chapmani* |
| Muridae | NC_065078.1 | *Serengetimys pernanus* |
| Muridae | NC_065337.1 | *Solomys ponceleti* |
| Muridae | NC_051519.1 | *Stenocephalemys zimai* |
| Muridae | NC_051518.1 | *Stenocephalemys albocaudata* |
| Muridae | NC_051517.1 | *Stenocephalemys griseicauda* |
| Muridae | NC_051516.1 | *Stenocephalemys sokolovi* |
| Muridae | NC_051515.1 | *Stenocephalemys ruppi* |
| Muridae | NC_051514.1 | *Stenocephalemys albipes* |
| Muridae | NC_053807.1 | *Stochomys longicaudatus* |
| Muridae | NC_036729.1 | *Sundamys annandalei* |
| Muridae | NC_036727.1 | *Sundamys maxi* |
| Muridae | NC_036728.1 | *Sundamys infraluteus* |
| Muridae | NC_053814.1 | *Thallomys paedulcus* |
| Muridae | NC_053813.1 | *Thamnomys kempi* |
| Muridae | NC_065086.1 | *Zelotomys hildegardeae* |
| Mus | NC_049117.1 | *Mus baoulei* |
| Mus | NC_036680.1 | *Mus pahari* |
| Mus | NC_010650.1 | *Mus terricolor* |

| | | |
|---|---|---|
| Mus | NC_005089.1 | *Mus musculus* |
| Mus | NC_030342.1 | *Mus famulus* |
| Mus | NC_025952.1 | *Mus spretus* |
| Mus | NC_025287.1 | *Mus fragilicauda* |
| Mus | NC_025270.1 | *Mus cookii* |
| Mus | NC_025269.1 | *Mus cervicolor* |
| Mus | NC_025268.1 | *Mus caroli* |
| Mus | NC_006915.1 | *Mus musculus molossinus* |
| Mus | NC_012387.1 | *Mus musculus castaneus* |
| Muscicapidae | NC_030603.1 | *Copsychus saularis* |
| Muscicapidae | NC_052839.1 | *Cossypha semirufa* |
| Muscicapidae | NC_015232.1 | *Cyanoptila cyanomelana* |
| Muscicapidae | NC_058320.1 | *Ficedula hyperythra* |
| Muscicapidae | NC_021621.1 | *Ficedula albicollis* |
| Muscicapidae | NC_015802.1 | *Ficedula zanthopygia* |
| Muscicapidae | NC_052841.1 | *Melaenornis chocolatinus* |
| Muscicapidae | NC_045375.1 | *Muscicapa latirostris* |
| Muscicapidae | NC_045374.1 | *Muscicapa sibirica* |
| Muscicapidae | NC_045181.1 | *Muscicapa griseisticta* |
| Muscicapidae | NC_039538.1 | *Niltava davidi* |
| Muscicapidae | NC_040290.1 | *Oenanthe isabellina* |
| Muscicapidae | NC_046943.1 | *Paradoxornis heudei* |
| Muscicapidae | NC_039536.1 | *Paradoxornis gularis* |

| | | |
|---|---|---|
| Muscicapidae | NC_028436.1 | *Paradoxornis fulvifrons* |
| Muscicapidae | NC_028437.1 | *Paradoxornis nipalensis* |
| Muscicapidae | NC_024539.1 | *Paradoxornis webbianus* |
| Muscicapidae | NC_053917.1 | *Phoenicurus frontalis* |
| Muscicapidae | NC_026066.1 | *Phoenicurus auroreus* |
| Mustela | NC_056132.1 | *Mustela lutreola* |
| Mustela | NC_034330.1 | *Mustela itatsi* |
| Mustela | NC_028013.1 | *Mustela eversmannii* |
| Mustela | NC_025516.1 | *Mustela erminea* |
| Mustela | NC_024942.1 | *Mustela nigripes* |
| Mustela | NC_021751.1 | *Mustela altaica* |
| Mustela | NC_023210.1 | *Mustela kathiah* |
| Mustela | NC_020640.1 | *Mustela frenata* |
| Mustela | NC_020639.1 | *Mustela nivalis* |
| Mustela | NC_020638.1 | *Mustela putorius* |
| Mustelidae | NC_053973.1 | *Galictis vittata* |
| Mustelidae | NC_065092.1 | *Ictonyx libycus* |
| Mustelidae | NC_053979.1 | *Ictonyx striatus* |
| Mustelidae | NC_053975.1 | *Poecilogale albinucha* |
| Mustelidae | NC_054246.1 | *Vormela peregusna* |
| Mustelidae | NC_020644.1 | *Melogale moschata* |
| Mustelidae | NC_046484.1 | *Aonyx capensis* |
| Mustelidae | NC_035814.1 | *Aonyx cinerea* |

| | | |
|---|---|---|
| Mustelidae | NC_009692.1 | *Enhydra lutris* |
| Mustelidae | NC_046485.1 | *Hydrictis maculicollis* |
| Mustelidae | NC_035810.1 | *Lutra sumatrana* |
| Mustelidae | NC_035811.1 | *Lutrogale perspicillata* |
| Mustelidae | NC_020645.1 | *Arctonyx collaris* |
| Mustelidae | NC_039173.1 | *Meles leucurus* |
| Mustelidae | NC_009677.1 | *Meles anakuma* |
| Mustelidae | NC_053981.1 | *Mellivora capensis* |
| Mustelidae | NC_020646.1 | *Taxidea taxus* |
| Myomorpha | NC_065673.1 | *Acomys mullah* |
| Myomorpha | NC_020758.1 | *Acomys cahirinus* |
| Myomorpha | NC_042667.1 | *Gerbilliscus leucogaster* |
| Myomorpha | NC_034314.1 | *Meriones tamariscinus* |
| Myomorpha | NC_027684.1 | *Meriones meridianus* |
| Myomorpha | NC_027683.1 | *Meriones libycus* |
| Myomorpha | NC_023263.1 | *Meriones unguiculatus* |
| Myomorpha | NC_037509.1 | *Psammomys obesus* |
| Myomorpha | NC_033382.1 | *Typhlomys cinereus* |
| Myomorpha | NC_039104.1 | *Rhizomys sumatrensis* |
| Myomorpha | NC_026124.1 | *Rhizomys sinensis* |
| Myomorpha | NC_021478.1 | *Rhizomys pruinosus* |
| Myomorpha | NC_058275.1 | *Tachyoryctes macrocephalus* |
| Myomorpha | NC_048987.1 | *Eospalax smithii* |

| | | |
|---|---|---|
| Myomorpha | NC_047427.1 | *Eospalax rufescens* |
| Myomorpha | NC_045904.1 | *Eospalax fontanierii* |
| Myomorpha | NC_021129.1 | *Eospalax cansus* |
| Myomorpha | NC_018535.1 | *Eospalax rothschildi* |
| Myomorpha | NC_018098.1 | *Eospalax baileyi* |
| Myomorpha | NC_026915.1 | *Myospalax aspalax* |
| Myomorpha | NC_026034.1 | *Myospalax psilurus* |
| Myomorpha | NC_020755.1 | *Nannospalax judaei* |
| Myomorpha | NC_020754.1 | *Nannospalax galili* |
| Myomorpha | NC_020757.1 | *Nannospalax golani* |
| Myomorpha | NC_020756.1 | *Spalax carmeli* |
| Myomorpha | NC_025746.1 | *Akodon montensis* |
| Myomorpha | NC_061914.1 | *Holochilus sciureus* |
| Myomorpha | NC_039723.1 | *Oligoryzomys stramineus* |
| Myomorpha | NC_035572.1 | *Sigmodon hispidus* |
| Myomorpha | NC_025747.1 | *Wiedomys cerradensis* |
| Myotis | NC_060697.1 | *Myotis aurascens* |
| Myotis | NC_056773.1 | *Myotis petax* |
| Myotis | NC_056111.1 | *Myotis ricketti* |
| Myotis | NC_041638.1 | *Myotis frater* |
| Myotis | NC_036328.1 | *Myotis martiniquensis* |
| Myotis | NC_036327.1 | *Myotis albescens* |
| Myotis | NC_036326.1 | *Myotis volans* |

| | | |
|---|---|---|
| Myotis | NC_036325.1 | *Myotis horsfieldii* |
| Myotis | NC_036324.1 | *Myotis atacamensis* |
| Myotis | NC_036323.1 | *Myotis thysanodes* |
| Myotis | NC_036321.1 | *Myotis leibii* |
| Myotis | NC_036319.1 | *Myotis yumanensis* |
| Myotis | NC_036318.1 | *Myotis nigricans* |
| Myotis | NC_036317.1 | *Myotis riparius* |
| Myotis | NC_036316.1 | *Myotis oxyotus* |
| Myotis | NC_036314.1 | *Myotis keaysi* |
| Myotis | NC_036313.1 | *Myotis evotis* |
| Myotis | NC_036312.1 | *Myotis dominicensis* |
| Myotis | NC_034227.1 | *Myotis bechsteinii* |
| Myotis | NC_029849.1 | *Myotis lucifugus* |
| Myotis | NC_029422.1 | *Myotis muricola* |
| Myotis | NC_029346.1 | *Myotis myotis* |
| Myotis | NC_029342.1 | *Myotis bombinus* |
| Myotis | NC_015828.1 | *Myotis formosus* |
| Myotis | NC_022698.1 | *Myotis ikonnikovi* |
| Myotis | NC_022694.1 | *Myotis macrodactylus* |
| Myotis | NC_025308.1 | *Myotis brandtii* |
| Myotis | NC_025568.1 | *Myotis davidii* |
| Myotis | NC_049871.1 | *Myotis septentrionalis* |
| Myotis | NC_036315.1 | *Myotis ruber* |

| | | |
|---|---|---|
| Myotis | NC_036320.1 | *Myotis auriculus* |
| Myrmeleontidae | NC_064385.1 | *Austrogymnocnemia maculata* |
| Myrmeleontidae | NC_032298.1 | *Bullanga florida* |
| Myrmeleontidae | NC_064383.1 | *Dendroleon similis* |
| Myrmeleontidae | NC_064386.1 | *Distoleon nigricans* |
| Myrmeleontidae | NC_025905.1 | *Epacanthaclisis banksi* |
| Myrmeleontidae | NC_064382.1 | *Gatzara nigrivena* |
| Myrmeleontidae | NC_064384.1 | *Glenoleon pulchellus* |
| Myrmeleontidae | NC_061035.1 | *Layahima valida* |
| Myrmeleontidae | NC_061034.1 | *Layahima weiweii* |
| Myrmeleontidae | NC_061033.1 | *Layahima wuzhishana* |
| Myrmeleontidae | NC_047285.1 | *Myrmeleon formicarius* |
| Myrmeleontidae | NC_024826.1 | *Myrmeleon immanis* |
| Myrmeleontidae | NC_064381.1 | *Nepsalus petrophilus* |
| Myrmeleontidae | NC_064380.1 | *Nepsalus decorosus* |
| Myrmeleontidae | NC_064379.1 | *Nepsalus decorillus* |
| Myrmeleontidae | NC_064378.1 | *Nepsalus caelestis* |
| Myrmicinae | NC_030541.1 | *Wasmannia auropunctata* |
| Myrmicinae | NC_062352.1 | *Crematogaster matsumurai* |
| Myrmicinae | NC_015075.1 | *Pristomyrmex punctatus* |
| Myrmicinae | NC_026133.1 | *Myrmica scabrinodis* |
| Myrmicinae | NC_060647.1 | *Messor structor* |
| Myrmicinae | NC_060604.1 | *Carebara diversa* |

| | | |
|---|---|---|
| Myrmicinae | NC_051486.1 | *Monomorium pharaonis* |
| Myrmicinae | NC_014672.1 | *Solenopsis invicta* |
| Myrmicinae | NC_014677.1 | *Solenopsis richteri* |
| Myrmicinae | NC_014669.1 | *Solenopsis geminata* |
| Myrmicinae | NC_062436.1 | *Stenamma impar* |
| Myrmicinae | NC_030176.1 | *Vollenhovia emeryi* |
| Nasutitermes | NC_034146.1 | *Nasutitermes macrocephalus* |
| Nasutitermes | NC_034115.1 | *Nasutitermes exitiosus* |
| Nasutitermes | NC_034045.1 | *Nasutitermes octopilis* |
| Nasutitermes | NC_034034.1 | *Nasutitermes matangensis* |
| Nasutitermes | NC_034026.1 | *Nasutitermes banksi* |
| Nasutitermes | NC_034023.1 | *Nasutitermes longirostris* |
| Nasutitermes | NC_034020.1 | *Nasutitermes diabolus* |
| Nasutitermes | NC_034108.1 | *Nasutitermes arborum* |
| Nasutitermes | NC_034093.1 | *Nasutitermes similis* |
| Nasutitermes | NC_034080.1 | *Nasutitermes neoparvus* |
| Nasutitermes | NC_034042.1 | *Nasutitermes lujae* |
| Nasutitermes | NC_026115.1 | *Nasutitermes corniger* |
| Nasutitermes | NC_018131.1 | *Nasutitermes triodiae* |
| Nasutitermes | NC_062440.1 | *Nasutitermes tiantongensis* |
| Nasutitermes | NC_034040.1 | *Nasutitermes graveolus* |
| Nasutitermes | NC_034117.1 | *Nasutitermes latifrons* |
| Nasutitermes | NC_034060.1 | *Nasutitermes longipennis* |

| | | |
|---|---|---|
| Nematocera | NC_016202.1 | *Protoplasa fitchii* |
| Nematocera | NC_054180.1 | *Ptychoptera qinggouensis* |
| Nematocera | NC_050315.1 | *Ptychoptera minuta* |
| Nemouridae | NC_060429.1 | *Amphinemura claviloba* |
| Nemouridae | NC_059860.1 | *Amphinemura wui* |
| Nemouridae | NC_057056.1 | *Amphinemura bulla* |
| Nemouridae | NC_044749.1 | *Amphinemura yao* |
| Nemouridae | NC_044748.1 | *Amphinemura longispina* |
| Nemouridae | NC_044751.1 | *Indonemoura nohirae* |
| Nemouridae | NC_044750.1 | *Indonemoura jacobsoni* |
| Nemouridae | NC_064323.1 | *Malenka flexura* |
| Nemouridae | NC_044720.1 | *Mesonemoura tritaenia* |
| Nemouridae | NC_044719.1 | *Mesonemoura metafiligera* |
| Nemouridae | NC_044755.1 | *Sphaeronemoura acutispina* |
| Nemouridae | NC_057512.1 | *Sphaeronemoura elephas* |
| Nemouridae | NC_044754.1 | *Sphaeronemoura grandicauda* |
| Neobatrachia | NC_047224.1 | *Anaxyrus americanus* |
| Neobatrachia | NC_020048.1 | *Bufo tibetanus* |
| Neobatrachia | NC_027686.1 | *Bufo stejnegeri* |
| Neobatrachia | NC_005794.2 | *Bufo melanostictus* |
| Neobatrachia | NC_009886.1 | *Bufo japonicus* |
| Neobatrachia | NC_008410.1 | *Bufo gargarizans* |
| Neobatrachia | NC_050665.1 | *Bufotes variabilis* |

| | | |
|---|---|---|
| Neobatrachia | NC_047225.1 | *Bufotes pewzowi* |
| Neobatrachia | NC_046047.1 | *Bufotes zamdaensis* |
| Neobatrachia | NC_028424.1 | *Bufotes raddei* |
| Neobatrachia | NC_062077.1 | *Bufotes turanensis* |
| Neobatrachia | NC_037378.1 | *Melanophryniscus moreirae* |
| Neobatrachia | NC_066225.1 | *Rhinella marina* |
| Neobatrachia | NC_019998.1 | *Heleophryne regis* |
| Neobatrachia | NC_063649.1 | *Dryophytes femoralis* |
| Neobatrachia | NC_063648.1 | *Dryophytes andersonii* |
| Neobatrachia | NC_045917.1 | *Dryophytes versicolor* |
| Neobatrachia | NC_010232.1 | *Dryophytes japonicus* |
| Neobatrachia | NC_032380.1 | *Dryophytes suweonensis* |
| Neobatrachia | NC_062326.1 | *Hyla sanchiangensis* |
| Neobatrachia | NC_029410.1 | *Hyla ussuriensis* |
| Neobatrachia | NC_026524.1 | *Hyla tsinlingensis* |
| Neobatrachia | NC_025309.1 | *Hyla annectans* |
| Neobatrachia | NC_006403.1 | *Hyla chinensis* |
| Neobatrachia | NC_019999.1 | *Lechriodus melanopyga* |
| Neobatrachia | NC_020001.1 | *Sooglossus thomasseti* |
| Neobatrachia | NC_020002.1 | *Telmatobius bolivianus* |
| Neobatrachia | NC_030333.1 | *Telmatobius chusmisensis* |
| Neobatrachia | NC_037857.1 | *Anomaloglossus blanci* |
| Neobatrachia | NC_037855.1 | *Anomaloglossus dewynteri* |

| | | |
|---|---|---|
| Neobatrachia | NC_037854.1 | *Anomaloglossus surinamensis* |
| Neobatrachia | NC_030054.1 | *Anomaloglossus baeobatrachus* |
| Neobatrachia | NC_037856.1 | *Anomaloglossus degranvillei* |
| Neobatrachia | NC_037380.1 | *Phyllobates terribilis* |
| Neobatrachia | NC_037379.1 | *Hyloxalus subpunctatus* |
| Neobatrachia | NC_036493.1 | *Bokermannohyla alvarengai* |
| Neobatrachia | NC_041426.1 | *Pseudis tocantins* |
| Neobatrachia | NC_056366.1 | *Leptodactylus fallax* |
| Neobatrachia | NC_039411.1 | *Kaloula verrucosa* |
| Neobatrachia | NC_020044.1 | *Kaloula borealis* |
| Neobatrachia | NC_029409.1 | *Kaloula rugifera* |
| Neobatrachia | NC_006405.1 | *Kaloula pulchra* |
| Neognathae | NC_031871.1 | *Alopecoenas salamonis* |
| Neognathae | NC_031869.1 | *Caloenas nicobarica* |
| Neognathae | NC_031870.1 | *Caloenas maculata* |
| Neognathae | NC_013978.1 | *Columba livia* |
| Neognathae | NC_048989.1 | *Columba hodgsonii* |
| Neognathae | NC_031868.1 | *Columba jouyi* |
| Neognathae | NC_031867.1 | *Columba rupestris* |
| Neognathae | NC_031866.1 | *Didunculus strigirostris* |
| Neognathae | NC_042502.1 | *Ectopistes migratorius* |
| Neognathae | NC_047279.1 | *Geopelia cuneata* |
| Neognathae | NC_015207.1 | *Geotrygon violacea* |

| | | |
|---|---|---|
| Neognathae | NC_036613.1 | *Goura victoria* |
| Neognathae | NC_031865.1 | *Goura cristata* |
| Neognathae | NC_027947.1 | *Goura scheepmakeri* |
| Neognathae | NC_013244.1 | *Hemiphaga novaeseelandiae* |
| Neognathae | NC_015190.1 | *Leptotila verreauxi* |
| Neognathae | NC_036612.1 | *Otidiphaps nobilis* |
| Neognathae | NC_059796.1 | *Streptopelia tranquebarica* |
| Neognathae | NC_037513.1 | *Streptopelia decaocto* |
| Neognathae | NC_031447.1 | *Streptopelia orientalis* |
| Neognathae | NC_026459.1 | *Streptopelia chinensis* |
| Neognathae | NC_062674.1 | *Treron sphenurus* |
| Neognathae | NC_059795.1 | *Treron curvirostra* |
| Neognathae | NC_036611.1 | *Trugon terrestris* |
| Neognathae | NC_031863.1 | *Zenaida macroura* |
| Neognathae | NC_015203.1 | *Zenaida auriculata* |
| Neognathae | NC_057089.1 | *Falco subbuteo* |
| Neognathae | NC_000878.1 | *Falco peregrinus* |
| Neognathae | NC_039842.1 | *Falco amurensis* |
| Neognathae | NC_029359.1 | *Falco rusticolus* |
| Neognathae | NC_025579.1 | *Falco columbarius* |
| Neognathae | NC_026715.1 | *Falco cherrug* |
| Neognathae | NC_011307.1 | *Falco tinnunculus* |
| Neognathae | NC_008547.1 | *Falco sparverius* |

| | | |
|---|---|---|
| Neognathae | NC_039736.1 | *Asio otus* |
| Neognathae | NC_027606.1 | *Asio flammeus* |
| Neognathae | NC_038220.1 | *Bubo scandiacus* |
| Neognathae | NC_034296.1 | *Glaucidium cuculoides* |
| Neognathae | NC_033967.1 | *Ninox strenua* |
| Neognathae | NC_029384.1 | *Ninox scutulata* |
| Neognathae | NC_005932.1 | *Ninox novaeseelandiae* |
| Neognathae | NC_041422.1 | *Otus sunia* |
| Neognathae | NC_028163.1 | *Otus bakkamoena* |
| Neognathae | NC_028162.1 | *Otus scops* |
| Neognathae | NC_038218.1 | *Strix uralensis* |
| Neognathae | NC_023946.1 | *Ciconia nigra* |
| Neognathae | NC_002196.1 | *Ciconia boyciana* |
| Neognathae | NC_052814.1 | *Pterocles burchelli* |
| Neognathae | NC_002783.2 | *Pterocnemia pennata* |
| Neognathae | NC_052772.1 | *Colius striatus* |
| Neognathae | NC_052817.1 | *Urocolius indicus* |
| Neognathae | NC_031864.1 | *Raphus cucullatus* |
| Neognathae | NC_044672.1 | *Caracara plancus* |
| Neognathae | NC_044673.1 | *Caracara cheriway* |
| Neognathae | NC_044674.1 | *Caracara creightoni* |
| Neognathae | NC_052801.1 | *Herpetotheres cachinnans* |
| Neognathae | NC_008548.1 | *Micrastur gilvicollis* |

| | | |
|---|---|---|
| Neognathae | NC_031897.1 | *Phalcoboenus australis* |
| Neognathae | NC_052779.1 | *Bucco capensis* |
| Neognathae | NC_007007.1 | *Gavia stellata* |
| Neognathae | NC_041165.1 | *Gavia arctica* |
| Neognathae | NC_008139.1 | *Gavia pacifica* |
| Neognathae | NC_052806.1 | *Corythaixoides concolor* |
| Neognathae | NC_027934.1 | *Phoenicopterus ruber* |
| Neognathae | NC_010089.1 | *Phoenicopterus roseus* |
| Neognathae | NC_008140.1 | *Podiceps cristatus* |
| Neognathae | NC_024594.1 | *Tachybaptus ruficollis* |
| Neognathae | NC_010095.1 | *Tachybaptus novaehollandiae* |
| Neognathae | NC_023787.1 | *Phodilus badius* |
| Neognathae | NC_052802.1 | *Trogon melanurus* |
| Neognathae | NC_011714.1 | *Trogon viridis* |
| Neognathae | NC_052796.1 | *Rhinopomastus cyanomelas* |
| Neognathae | NC_028178.1 | *Upupa epops* |
| Nesophrosyne | NC_066173.1 | *Nesophrosyne maritima* |
| Nesophrosyne | NC_066170.1 | *Nesophrosyne sp. 48* |
| Nesophrosyne | NC_066169.1 | *Nesophrosyne sp. 242* |
| Nesophrosyne | NC_066168.1 | *Nesophrosyne sp. 58* |
| Nesophrosyne | NC_066167.1 | *Nesophrosyne sp. 21* |
| Nesophrosyne | NC_066165.1 | *Nesophrosyne sp. 126* |
| Nesophrosyne | NC_066164.1 | *Nesophrosyne sp. 29* |

| | | |
|---|---|---|
| Nesophrosyne | NC_066163.1 | *Nesophrosyne sp. 23* |
| Nesophrosyne | NC_066161.1 | *Nesophrosyne montium* |
| Nesophrosyne | NC_066160.1 | *Nesophrosyne sp. 281* |
| Neuropterida | NC_035504.1 | *Archichauliodes deceptor* |
| Neuropterida | NC_024657.1 | *Dysmicohermes ingens* |
| Neuropterida | NC_025282.1 | *Neochauliodes fraternus* |
| Neuropterida | NC_025281.1 | *Neochauliodes rotundatus* |
| Neuropterida | NC_023444.1 | *Neochauliodes bowringi* |
| Neuropterida | NC_018772.1 | *Neochauliodes punctatolosus* |
| Neuropterida | NC_023462.1 | *Acanthacorydalis orientalis* |
| Neuropterida | NC_033349.1 | *Chloronia mirifica* |
| Neuropterida | NC_011276.1 | *Corydalus cornutus* |
| Neuropterida | NC_027852.1 | *Neoneuromus tonkinensis* |
| Neuropterida | NC_027851.1 | *Nevromus exterior* |
| Neuropterida | NC_011524.1 | *Protohermes concolorus* |
| Neuropterida | NC_013256.1 | *Sialis hamata* |
| Neuropterida | NC_021428.1 | *Ascalohybris subjacens* |
| Neuropterida | NC_011277.1 | *Ascaloptynx appendiculatus* |
| Neuropterida | NC_015609.1 | *Libelloides macaronius* |
| Neuropterida | NC_039948.1 | *Suhpalacsa longialata* |
| Neuropterida | NC_051001.1 | *Hemerobius spodipennis* |
| Neuropterida | NC_042677.1 | *Micromus angulatus* |
| Neuropterida | NC_028153.1 | *Neuronema laminatum* |

| | | |
|---|---|---|
| Neuropterida | NC_011278.1 | *Polystoechotes punctatus* |
| Neuropterida | NC_026228.1 | *Polytremis nascens* |
| Neuropterida | NC_026990.1 | *Polytremis jigongi* |
| Neuropterida | NC_013257.1 | *Ditaxis biseriata* |
| Neuropterida | NC_039773.1 | *Euclimacia badia* |
| Neuropterida | NC_039772.1 | *Eumantispa harmandi* |
| Neuropterida | NC_039771.1 | *Mantispa japonica* |
| Neuropterida | NC_024825.1 | *Nymphes myrmeleonoides* |
| Neuropterida | NC_053271.1 | *Gryposmylus pennyi* |
| Neuropterida | NC_050653.1 | *Osmylus fulvicephalus* |
| Neuropterida | NC_046578.1 | *Thaumatosmylus hainanus* |
| Neuropterida | NC_021415.1 | *Thyridosmylus langii* |
| Neuropterida | NC_023363.1 | *Rapisma zayuanum* |
| Neuropterida | NC_013251.1 | *Mongoloraphidia harmandi* |
| Neuropterida | NC_015095.1 | *Apochrysa matsumurae* |
| Neuropterida | NC_019618.1 | *Chrysopa pallens* |
| Neuropterida | NC_030341.1 | *Chrysoperla externa* |
| Neuropterida | NC_015093.1 | *Chrysoperla nipponensis* |
| Neuropterida | NC_057219.1 | *Conwentzia sinica* |
| Neuropterida | NC_061560.1 | *Semidalis macleodi* |
| Noctuidae | NC_046049.1 | *Anarta trifolii* |
| Noctuidae | NC_062880.1 | *Condica illecta* |
| Noctuidae | NC_062101.1 | *Condica capensis* |

| | | |
|---|---|---|
| Noctuidae | NC_046524.1 | *Lacanobia aliena* |
| Noctuidae | NC_059936.1 | *Leiometopon simyrides* |
| Noctuidae | NC_061705.1 | *Mamestra brassicae* |
| Noctuidae | NC_062113.1 | *Melanchra persicariae* |
| Noctuidae | NC_064392.1 | *Mythimna unipuncta* |
| Noctuidae | NC_057500.1 | *Mythimna loreyi* |
| Noctuidae | NC_023118.1 | *Mythimna separata* |
| Noctuidae | NC_034938.1 | *Protegira songi* |
| Noctuidae | NC_015835.1 | *Sesamia inferens* |
| Noctuidae | NC_061562.1 | *Spodoptera depravata* |
| Noctuidae | NC_054179.1 | *Spodoptera exempta* |
| Noctuidae | NC_053623.1 | *Spodoptera littoralis* |
| Noctuidae | NC_027836.1 | *Spodoptera frugiperda* |
| Noctuidae | NC_022676.1 | *Spodoptera litura* |
| Noctuidae | NC_019622.1 | *Spodoptera exigua* |
| Noctuidae | NC_062103.1 | *Tiracola aureata* |
| Noctuidae | NC_062090.1 | *Tiracola plagiata* |
| Noctuidae | NC_062172.1 | *Acronicta major* |
| Noctuidae | NC_062117.1 | *Acronicta rumicis* |
| Noctuidae | NC_061640.1 | *Sphragifera sigillata* |
| Noctuidae | NC_064395.1 | *Xanthodes albago* |
| Noctuidae | NC_062099.1 | *Xanthodes intersepta* |
| Noctuidae | NC_062110.1 | *Eucarta virgo* |

| | | |
|---|---|---|
| Noctuidae | NC_062181.1 | *Cucullia pustulata* |
| Noctuidae | NC_061507.1 | *Helicoverpa zea* |
| Noctuidae | NC_014668.1 | *Helicoverpa armigera* |
| Noctuidae | NC_035890.1 | *Helicoverpa assulta* |
| Noctuidae | NC_023791.1 | *Helicoverpa punctigera* |
| Noctuidae | NC_028539.1 | *Heliothis subflexa* |
| Noctuidae | NC_061565.1 | *Pyrrhia umbra* |
| Noctuidae | NC_062112.1 | *Cosmia restituta* |
| Noctuidae | NC_062171.1 | *Actebia praecox* |
| Noctuidae | NC_065464.1 | *Agrotis exclamationis* |
| Noctuidae | NC_065463.1 | *Agrotis tokionis* |
| Noctuidae | NC_062121.1 | *Agrotis munda* |
| Noctuidae | NC_062115.1 | *Agrotis trifurca* |
| Noctuidae | NC_022689.1 | *Agrotis segetum* |
| Noctuidae | NC_022185.1 | *Agrotis ipsilon* |
| Noctuidae | NC_059038.1 | *Anaplectoides virens* |
| Noctuidae | NC_062100.1 | *Athetis thoracica* |
| Noctuidae | NC_046525.1 | *Athetis pallidipennis* |
| Noctuidae | NC_036057.1 | *Athetis lepigone HB25* |
| Noctuidae | NC_025774.1 | *Striacosta albicosta* |
| Noctuidae | NC_065462.1 | *Xestia c-nigrum* |
| Noctuidae | NC_062120.1 | *Abrostola triplasia* |
| Noctuidae | NC_066154.1 | *Chrysodeixis acuta* |

| | | |
|---|---|---|
| Noctuidae | NC_053742.1 | *Ctenoplusia albostriata* |
| Noctuidae | NC_025760.1 | *Ctenoplusia limbirena* |
| Noctuidae | NC_021410.1 | *Ctenoplusia agnata* |
| Noctuidae | NC_053855.1 | *Diachrysia nadeja* |
| Noctuidae | NC_045936.1 | *Trichoplusia ni* |
| Noctuidae | NC_062107.1 | *Actinotia intermediata* |
| Noctuidae | NC_061564.1 | *Imosca coreana* |
| Noctuidae | NC_062119.1 | *Niphonyx segregata* |
| Nolidae | NC_066709.1 | *Blenina donans* |
| Nolidae | NC_064396.1 | *Camptoloma kishidai* |
| Nolidae | NC_066086.1 | *Carea varipes* |
| Nolidae | NC_026842.1 | *Gabala argentata* |
| Nolidae | NC_062184.1 | *Pseudoips prasinana* |
| Nolidae | NC_045120.1 | *Sinna extrema* |
| Nolidae | NC_062116.1 | *Earias clorana* |
| Nolidae | NC_062104.1 | *Eligma narcissus* |
| Nolidae | NC_026841.1 | *Risoba prominens* |
| Notodontidae | NC_062123.1 | *Syntypistis chambae* |
| Notodontidae | NC_062111.1 | *Peridea elzet* |
| Notodontidae | NC_061645.1 | *Pheosia rimosa* |
| Notodontidae | NC_016067.1 | *Phalera flavescens* |
| Notodontidae | NC_041140.1 | *Clostera anastomosis* |
| Notodontidae | NC_034740.1 | *Clostera anachoreta* |

| | | |
|---|---|---|
| Notodontidae | NC_062182.1 | *Neocerura liturata* |
| Notodontidae | NC_011128.1 | *Ochrogaster lunifer* |
| Notostraca | NC_044654.1 | *Lepidurus arcticus* |
| Notostraca | NC_044646.1 | *Lepidurus apus lubbocki* |
| Notostraca | NC_044781.1 | *Triops granarius* |
| Notostraca | NC_006079.1 | *Triops longicaudatus* |
| Nymphalinae | NC_061261.1 | *Hypolimnas misippus* |
| Nymphalinae | NC_026072.1 | *Hypolimnas bolina* |
| Nymphalinae | NC_064725.1 | *Hypolimnas usambara* |
| Nymphalinae | NC_064724.1 | *Hypolimnas anthedon* |
| Nymphalinae | NC_050691.1 | *Kallimoides rumia* |
| Nymphalinae | NC_064723.1 | *Precis tugela* |
| Nymphalinae | NC_064722.1 | *Precis pelarga* |
| Nymphalinae | NC_064721.1 | *Precis octavia* |
| Nymphalinae | NC_064719.1 | *Precis eurodoce* |
| Nymphalinae | NC_064718.1 | *Precis cuama* |
| Nymphalinae | NC_064717.1 | *Precis ceryne* |
| Nymphalinae | NC_064710.1 | *Precis archesia* |
| Nymphalinae | NC_041242.1 | *Precis andremiaja* |
| Nymphalinae | NC_064726.1 | *Protogoniomorpha parhassus* |
| Nymphalinae | NC_064709.1 | *Protogoniomorpha temora* |
| Nymphalinae | NC_064727.1 | *Salamis cacta* |
| Nymphalinae | NC_041243.1 | *Salamis anteva* |

| | | |
|---|---|---|
| Nymphalinae | NC_064711.1 | *Yoma algina* |
| Nymphalinae | NC_024403.1 | *Yoma sabina* |
| Obtectomera | NC_034279.1 | *Brahmaea hearseyi* |
| Obtectomera | NC_038106.1 | *Prismosticta fenestrata* |
| Obtectomera | NC_038010.1 | *Prismostictoides unihyala* |
| Obtectomera | NC_038084.1 | *Ganisa cyanogrisea* |
| Obtectomera | NC_036347.1 | *Dendrolimus kikuchii* |
| Obtectomera | NC_039841.1 | *Dendrolimus superans* |
| Obtectomera | NC_039840.1 | *Dendrolimus houi* |
| Obtectomera | NC_027157.1 | *Dendrolimus tabulaeformis* |
| Obtectomera | NC_025763.1 | *Dendrolimus spectabilis* |
| Obtectomera | NC_031507.1 | *Euthrix laeta* |
| Obtectomera | NC_062175.1 | *Trabala vishnou* |
| Obtectomera | NC_018133.1 | *Actias selene* |
| Obtectomera | NC_045899.1 | *Actias luna* |
| Obtectomera | NC_044744.1 | *Antheraea pernyi* |
| Obtectomera | NC_030270.1 | *Antheraea assama* |
| Obtectomera | NC_027071.1 | *Antheraea frithi* |
| Obtectomera | NC_012739.1 | *Antheraea yamamai* |
| Obtectomera | NC_012727.1 | *Eriogyna pyretorum* |
| Obtectomera | NC_036765.1 | *Neoris haraldi* |
| Obtectomera | NC_059700.1 | *Rhodinia fugax* |
| Obtectomera | NC_063568.1 | *Saturnia japonica* |

| | | |
|---|---|---|
| Obtectomera | NC_052913.1 | *Pterodecta felderi* |
| Obtectomera | NC_045248.1 | *Epicopeia hainesii* |
| Obtectomera | NC_050856.1 | *Macrosoma conifera* |
| Obtectomera | NC_038082.1 | *Andraca olivacea* |
| Obtectomera | NC_032694.1 | *Andraca theae* |
| Obtectomera | NC_038083.1 | *Comparmustilia sphingiformis* |
| Obtectomera | NC_038085.1 | *Mustilia undulosa* |
| Obtectomera | NC_038105.1 | *Mustilizans hepatica* |
| Obtectomera | NC_038086.1 | *Oberthueria jiatongae* |
| Obtectomera | NC_021770.1 | *Attacus atlas* |
| Obtectomera | NC_061325.1 | *Epiphora bauhiniae* |
| Obtectomera | NC_024270.1 | *Samia canningi* |
| Obtectomera | NC_066089.1 | *Manduca quinquemaculata* |
| Obtectomera | NC_010266.1 | *Manduca sexta* |
| Obtectomera | NC_037445.1 | *Psilogramma increta* |
| Obtectomera | NC_020780.1 | *Sphinx morio* |
| Obtectomera | NC_065770.1 | *Agnidra scabiosa* |
| Obtectomera | NC_065769.1 | *Pseudalbara parvula* |
| Obtectomera | NC_061643.1 | *Tethea albicostata* |
| Ochthebius | NC_052906.1 | *Ochthebius viridis* |
| Ochthebius | NC_052905.1 | *Ochthebius uniformis* |
| Ochthebius | NC_052904.1 | *Ochthebius sculptoides* |
| Ochthebius | NC_052903.1 | *Ochthebius scopuli* |

| | | |
|---|---|---|
| Ochthebius | NC_052902.1 | *Ochthebius salinarius* |
| Ochthebius | NC_052901.1 | *Ochthebius nobilis* |
| Ochthebius | NC_052900.1 | *Ochthebius mediterraneus* |
| Ochthebius | NC_052898.1 | *Ochthebius lividipennis* |
| Ochthebius | NC_052897.1 | *Ochthebius himalayae* |
| Ochthebius | NC_052896.1 | *Ochthebius hasegawai* |
| Ochthebius | NC_052894.1 | *Ochthebius glaber* |
| Ochthebius | NC_052893.1 | *Ochthebius deletus* |
| Ochthebius | NC_052892.1 | *Ochthebius capicola* |
| Ochthebius | NC_052891.1 | *Ochthebius atriceps* |
| Ochthebius | NC_052890.1 | *Ochthebius minoicus* |
| Ochthebius | NC_052887.1 | *Ochthebius remotus* |
| Ochthebius | NC_052886.1 | *Ochthebius puncticollis* |
| Ochthebius | NC_052885.1 | *Ochthebius plesiotypus* |
| Ochthebius | NC_052888.1 | *Ochthebius quadricollis* |
| Ochthebius | NC_052895.1 | *Ochthebius griotes* |
| Ocypodidae | NC_042401.1 | *Austruca lactea* |
| Ocypodidae | NC_039111.1 | *Cranuca inversa* |
| Ocypodidae | NC_038177.1 | *Gelasimus borealis* |
| Ocypodidae | NC_058243.1 | *Tubuca arcuata* |
| Ocypodidae | NC_039107.1 | *Tubuca capricornis* |
| Ocypodidae | NC_039106.1 | *Tubuca polita* |
| Ocypodidae | NC_046797.1 | *Ocypode stimpsoni* |

| | | |
|---|---|---|
| Odontoceti | NC_005274.1 | *Berardius bairdii* |
| Odontoceti | NC_060610.1 | *Cephalorhynchus commersonii* |
| Odontoceti | NC_020696.1 | *Cephalorhynchus heavisidii* |
| Odontoceti | NC_012061.1 | *Delphinus capensis* |
| Odontoceti | NC_036415.1 | *Delphinus delphis* |
| Odontoceti | NC_019588.1 | *Feresa attenuata* |
| Odontoceti | NC_019578.2 | *Globicephala macrorhynchus* |
| Odontoceti | NC_019441.1 | *Globicephala melas* |
| Odontoceti | NC_012062.1 | *Grampus griseus* |
| Odontoceti | NC_005273.1 | *Hyperoodon ampullatus* |
| Odontoceti | NC_034348.1 | *Indopacetus pacificus* |
| Odontoceti | NC_037848.1 | *Lagenodelphis hosei* |
| Odontoceti | NC_050265.1 | *Lagenorhynchus acutus* |
| Odontoceti | NC_035426.1 | *Lagenorhynchus obliquidens* |
| Odontoceti | NC_005278.1 | *Lagenorhynchus albirostris* |
| Odontoceti | NC_042218.1 | *Mesoplodon bidens* |
| Odontoceti | NC_021974.2 | *Mesoplodon densirostris* |
| Odontoceti | NC_036997.1 | *Mesoplodon stejnegeri* |
| Odontoceti | NC_027593.1 | *Mesoplodon ginkgodens* |
| Odontoceti | NC_023830.1 | *Mesoplodon grayi* |
| Odontoceti | NC_042217.1 | *Mesoplodon mirus* |
| Odontoceti | NC_021434.2 | *Mesoplodon europaeus* |
| Odontoceti | NC_019591.1 | *Orcaella heinsohni* |

| | | |
|---|---|---|
| Odontoceti | NC_019590.1 | *Orcaella brevirostris* |
| Odontoceti | NC_064558.1 | *Orcinus orca* |
| Odontoceti | NC_019589.1 | *Peponocephala electra* |
| Odontoceti | NC_019577.1 | *Pseudorca crassidens* |
| Odontoceti | NC_045404.1 | *Sousa teuszii* |
| Odontoceti | NC_012057.1 | *Sousa chinensis* |
| Odontoceti | NC_060612.1 | *Stenella frontalis* |
| Odontoceti | NC_060611.1 | *Stenella clymene* |
| Odontoceti | NC_012051.1 | *Stenella attenuata* |
| Odontoceti | NC_012053.1 | *Stenella coeruleoalba* |
| Odontoceti | NC_032301.1 | *Stenella longirostris* |
| Odontoceti | NC_042761.1 | *Steno bredanensis* |
| Odontoceti | NC_012059.1 | *Tursiops truncatus* |
| Odontoceti | NC_022805.1 | *Tursiops australis* |
| Odontoceti | NC_012058.1 | *Tursiops aduncus* |
| Odontoceti | NC_021435.1 | *Ziphius cavirostris* |
| Odontoceti | NC_005276.1 | *Inia geoffrensis* |
| Odontoceti | NC_007629.1 | *Lipotes vexillifer* |
| Odontoceti | NC_034236.1 | *Delphinapterus leucas* |
| Odontoceti | NC_005279.1 | *Monodon monoceros* |
| Odontoceti | NC_026456.1 | *Neophocaena asiaeorientalis* |
| Odontoceti | NC_021461.1 | *Neophocaena phocaenoides* |
| Odontoceti | NC_053752.1 | *Phocoena dioptrica* |

| | | |
|---|---|---|
| Odontoceti | NC_053751.1 | *Phocoena sinus* |
| Odontoceti | NC_053750.1 | *Phocoena spinipinnis* |
| Odontoceti | NC_053753.1 | *Phocoenoides dalli* |
| Odontoceti | NC_041303.1 | *Kogia sima* |
| Odontoceti | NC_005272.1 | *Kogia breviceps* |
| Odontoceti | NC_002503.2 | *Physeter catodon* |
| Odontoceti | NC_057433.1 | *Platanista gangetica* |
| Odontoceti | NC_005275.1 | *Platanista minor* |
| Odontoceti | NC_005277.1 | *Pontoporia blainvillei* |
| Oestroidea | NC_063664.1 | *Blaesoxipha lapidosa* |
| Oestroidea | NC_041073.1 | *Oxysarcodexia varia* |
| Oestroidea | NC_041072.1 | *Oxysarcodexia thornax* |
| Oestroidea | NC_041071.1 | *Oxysarcodexia terminalis* |
| Oestroidea | NC_041070.1 | *Oxysarcodexia avuncula* |
| Oestroidea | NC_041079.1 | *Peckia collusor* |
| Oestroidea | NC_041078.1 | *Peckia australis* |
| Oestroidea | NC_041077.1 | *Peckia resona* |
| Oestroidea | NC_026196.1 | *Ravinia pernix* |
| Olethreutinae | NC_022865.1 | *Retinia pseudotsugaicola* |
| Olethreutinae | NC_019619.1 | *Rhyacionia leptotubula* |
| Olethreutinae | NC_014294.1 | *Spilonota lechriaspis* |
| Olethreutinae | NC_020003.2 | *Cydia pomonella* |
| Olethreutinae | NC_024582.1 | *Grapholita dimorpha* |

| | | |
|---|---|---|
| Olethreutinae | NC_014806.1 | *Grapholita molesta* |
| Olethreutinae | NC_064065.1 | *Thaumatotibia leucotreta* |
| Olethreutinae | NC_046051.1 | *Celypha flavipalpana* |
| Olethreutinae | NC_029193.1 | *Lobesia botrana* |
| Orbiculariae | NC_032402.1 | *Araneus angulatus* |
| Orbiculariae | NC_025634.1 | *Araneus ventricosus* |
| Orbiculariae | NC_064399.1 | *Araniella displicata* |
| Orbiculariae | NC_044695.1 | *Argiope perforata* |
| Orbiculariae | NC_024281.1 | *Argiope bruennichi* |
| Orbiculariae | NC_044696.1 | *Cyclosa japonica* |
| Orbiculariae | NC_027682.1 | *Cyclosa argenteoalba* |
| Orbiculariae | NC_028077.1 | *Cyrtarachne nagasakiensis* |
| Orbiculariae | NC_028078.1 | *Hypsosinga pygmaea* |
| Orbiculariae | NC_044653.1 | *Neoscona multiplicans* |
| Orbiculariae | NC_044101.1 | *Neoscona scylla* |
| Orbiculariae | NC_029756.1 | *Neoscona adianta* |
| Orbiculariae | NC_029755.1 | *Neoscona nautica* |
| Orbiculariae | NC_026290.1 | *Neoscona theisi* |
| Orbiculariae | NC_061750.1 | *Leucauge wulingensis* |
| Orbiculariae | NC_028068.1 | *Tetragnatha nitens* |
| Orbiculariae | NC_025775.1 | *Tetragnatha maxillosa* |
| Orconectes | NC_029721.1 | *Orconectes sanbornii* |
| Orconectes | NC_029720.1 | *Orconectes rusticus* |

| | | |
|---|---|---|
| Orconectes | NC_026561.1 | *Orconectes limosus* |
| Orconectes | NC_033508.1 | *Orconectes luteus* |
| Orconectes | NC_030768.1 | *Orconectes punctimanus* |
| Ornithodoros | NC_039688.1 | *Ornithodoros sonrai* |
| Ornithodoros | NC_039857.1 | *Ornithodoros coriaceus California* |
| Ornithodoros | NC_039832.1 | *Ornithodoros hermsi* |
| Ornithodoros | NC_039831.1 | *Ornithodoros parkeri* |
| Ornithodoros | NC_039830.1 | *Ornithodoros tholozani* |
| Ornithodoros | NC_039829.1 | *Ornithodoros turicata Kansas* |
| Ornithodoros | NC_037525.1 | *Ornithodoros savignyi* |
| Ornithodoros | NC_033857.1 | *Ornithodoros zumpti* |
| Ornithodoros | NC_023373.1 | *Ornithodoros brasiliensis* |
| Ornithodoros | NC_023372.1 | *Ornithodoros rostratus* |
| Ornithodoros | NC_005820.1 | *Ornithodoros porcinus* |
| Ornithodoros | NC_004357.1 | *Ornithodoros moubata* |
| Otariidae | NC_063561.1 | *Arctocephalus australis* |
| Otariidae | NC_008420.1 | *Arctocephalus townsendi* |
| Otariidae | NC_004023.1 | *Arctocephalus forsteri* |
| Otariidae | NC_008415.3 | *Callorhinus ursinus* |
| Otariidae | NC_004030.2 | *Eumetopias jubatus* |
| Otariidae | NC_008419.1 | *Neophoca cinerea* |
| Otariidae | NC_049152.1 | *Otaria byronia* |
| Otariidae | NC_008418.1 | *Phocarctos hookeri* |

| | | |
|---|---|---|
| Otariidae | NC_062331.1 | *Zalophus wollebaeki* |
| Otariidae | NC_058016.1 | *Zalophus japonicus* |
| Otariidae | NC_008416.1 | *Zalophus californianus* |
| Paguroidea | NC_024202.1 | *Lithodes nintokuae* |
| Paguroidea | NC_057304.1 | *Pagurus similis* |
| Palaemonoidea | NC_046034.1 | *Anchistus australis* |
| Palaemonoidea | NC_012217.1 | *Macrobrachium lanchesteri* |
| Palaemonoidea | NC_027602.1 | *Macrobrachium bullatum* |
| Palaemonoidea | NC_015073.1 | *Macrobrachium nipponense* |
| Palaemonoidea | NC_006880.1 | *Macrobrachium rosenbergii* |
| Palaemonoidea | NC_050266.1 | *Palaemon serratus* |
| Palaemonoidea | NC_050168.1 | *Palaemon adspersus* |
| Palaemonoidea | NC_045090.1 | *Palaemon sinensis* |
| Palaemonoidea | NC_039373.1 | *Palaemon capensis* |
| Palaemonoidea | NC_038117.1 | *Palaemon annandalei* |
| Palaemonoidea | NC_029240.1 | *Palaemon gravieri* |
| Palaemonoidea | NC_027601.1 | *Palaemon serenus* |
| Palaemonoidea | NC_061664.1 | *Periclimenes brevicarpalis* |
| Palaeognathae | NC_052824.1 | *Apteryx rowi* |
| Palaeognathae | NC_013806.1 | *Apteryx owenii* |
| Palaeognathae | NC_002782.2 | *Apteryx haastii* |
| Palaeognathae | NC_002784.1 | *Dromaius novaehollandiae* |
| Palaeognathae | NC_002672.1 | *Dinornis giganteus* |

| | | |
|---|---|---|
| Palaeognathae | NC_002779.1 | *Anomalopteryx didiformis* |
| Palaeognathae | NC_002673.1 | *Emeus crassus* |
| Palaeognathae | NC_000846.1 | *Rhea americana* |
| Palaeognathae | NC_002785.1 | *Struthio camelus* |
| Pamphagidae | NC_064211.1 | *Haplotropis brunneriana* |
| Pamphagidae | NC_023535.1 | *Humphaplotropis culaishanensis* |
| Pamphagidae | NC_046559.1 | *Filchnerella tenggerensis* |
| Pamphagidae | NC_046558.1 | *Filchnerella qilianshanensis* |
| Pamphagidae | NC_024923.1 | *Filchnerella beicki* |
| Pamphagidae | NC_020329.1 | *Filchnerella helanshanensis* |
| Pamphagidae | NC_052733.1 | *Filchnerella rubrimargina* |
| Pamphagidae | NC_020330.1 | *Pseudotmethis rubimarginis* |
| Pamphagidae | NC_025904.1 | *Asiotmethis jubatus* |
| Pamphagidae | NC_020328.1 | *Asiotmethis zacharjini* |
| Pamphagidae | NC_014610.1 | *Thrinchus schrenkii* |
| Panheteroptera | NC_037370.1 | *Aphelocheirus jendeki* |
| Panheteroptera | NC_012838.1 | *Nerthra indica* |
| Panheteroptera | NC_039588.1 | *Micronecta sahlbergii* |
| Panheteroptera | NC_028629.1 | *Paraplea frontalis* |
| Panheteroptera | NC_056774.1 | *Lethocerus indicus* |
| Panheteroptera | NC_012822.1 | *Helotrephes sp.* |
| Panheteroptera | NC_012845.1 | *Ilyocoris cimicoides* |
| Panheteroptera | NC_012817.1 | *Laccotrephes robustus* |

| | | |
|---|---|---|
| Panheteroptera | NC_028084.1 | *Nepa hoffmanni* |
| Panheteroptera | NC_012819.1 | *Enithares tibialis* |
| Panheteroptera | NC_045886.1 | *Notonecta montandoni* |
| Panheteroptera | NC_036671.1 | *Notonecta chinensis* |
| Pannota | NC_057973.1 | *Cincticostella fusca* |
| Pannota | NC_050279.1 | *Ephemerella sp. Yunnan-2018* |
| Pannota | NC_050282.1 | *Serratella zapekinae* |
| Pannota | NC_050281.1 | *Serratella sp. Yunnan-2018* |
| Pannota | NC_050280.1 | *Serratella sp. Liaoning-2019* |
| Pannota | NC_057972.1 | *Torleya mikhaili* |
| Pannota | NC_050284.1 | *Torleya nepalica* |
| Pannota | NC_050283.1 | *Torleya grandiforceps* |
| Pannota | NC_065870.1 | *Vietnamella sinensis* |
| Papilio | NC_066472.1 | *Papilio dialis* |
| Papilio | NC_059755.1 | *Papilio thoas* |
| Papilio | NC_043911.1 | *Papilio memnon* |
| Papilio | NC_025757.1 | *Papilio helenus* |
| Papilio | NC_034356.1 | *Papilio rex* |
| Papilio | NC_034317.1 | *Papilio protenor* |
| Papilio | NC_023978.1 | *Papilio syfanius* |
| Papilio | NC_021411.1 | *Papilio maackii* |
| Papilio | NC_029244.1 | *Papilio xuthus* |
| Papilio | NC_014055.1 | *Papilio maraho* |

| | | |
|---|---|---|
| Papilio | NC_027252.1 | *Papilio glaucus* |
| Papilio | NC_027506.1 | *Papilio demoleus* |
| Papilio | NC_024742.1 | *Papilio polytes* |
| Papilio | NC_018047.1 | *Papilio machaon* |
| Papilio | NC_018040.1 | *Papilio bianor* |
| Papilio | NC_053770.1 | *Papilio paris* |
| Papilio | NC_037874.1 | *Papilio slateri* |
| Papilio | NC_034355.1 | *Papilio dardanus* |
| Papilionidae | NC_034837.1 | *Graphium leechi* |
| Papilionidae | NC_026910.1 | *Graphium chironides* |
| Papilionidae | NC_024098.1 | *Graphium timur* |
| Papilionidae | NC_037867.1 | *Lamproptera meges* |
| Papilionidae | NC_023953.1 | *Lamproptera curius* |
| Papilionidae | NC_037871.1 | *Mimoides lysithous* |
| Papilionidae | NC_027108.1 | *Teinopalpus imperialis* |
| Papilionidae | NC_014398.1 | *Teinopalpus aureus* |
| Papilioninae | NC_024564.1 | *Atrophaneura alcinous* |
| Papilioninae | NC_037868.1 | *Losaria neptunus* |
| Papilioninae | NC_037870.1 | *Ornithoptera priamus* |
| Papilioninae | NC_037869.1 | *Ornithoptera richmondia* |
| Papilioninae | NC_034280.1 | *Pachliopta aristolochiae* |
| Papilioninae | NC_037875.1 | *Trogonoptera brookiana* |
| Papilioninae | NC_060569.1 | *Troides aeacus formosanus* |

| | | |
|---|---|---|
| Papilionoidea | NC_066397.1 | *Apatura iris* |
| Papilionoidea | NC_065069.1 | *Apatura nycteis* |
| Papilionoidea | NC_016062.1 | *Apatura ilia* |
| Papilionoidea | NC_015537.1 | *Apatura metis* |
| Papilionoidea | NC_026569.1 | *Chitoria ulupi* |
| Papilionoidea | NC_027109.1 | *Euripus nyctelius* |
| Papilionoidea | NC_028086.1 | *Herona marathus* |
| Papilionoidea | NC_064982.1 | *Hestina assimilis* |
| Papilionoidea | NC_063472.1 | *Hestina persimilis* |
| Papilionoidea | NC_063473.1 | *Hestinalis nama* |
| Papilionoidea | NC_014224.1 | *Sasakia charonda* |
| Papilionoidea | NC_022134.1 | *Sasakia funebris* |
| Papilionoidea | NC_065067.1 | *Sephisa princeps* |
| Papilionoidea | NC_021090.1 | *Timelaea maculata* |
| Papilionoidea | NC_026070.1 | *Cethosia biblis* |
| Papilionoidea | NC_024404.1 | *Abrota ganga* |
| Papilionoidea | NC_065216.1 | *Bassarona dunya* |
| Papilionoidea | NC_024400.1 | *Dophla evelina* |
| Papilionoidea | NC_024396.1 | *Euthalia irrubescens* |
| Papilionoidea | NC_024399.1 | *Lexias dirtea* |
| Papilionoidea | NC_024416.1 | *Tanaecia julii* |
| Papilionoidea | NC_064983.1 | *Argynnis paphia* |
| Papilionoidea | NC_064981.1 | *Argynnis anadyomene* |

| | | |
|---|---|---|
| Papilionoidea | NC_015988.1 | *Argynnis hyperbius* |
| Papilionoidea | NC_024415.1 | *Argynnis childreni* |
| Papilionoidea | NC_016419.1 | *Fabriciana nerippe* |
| Papilionoidea | NC_050261.1 | *Issoria eugenia* |
| Papilionoidea | NC_018030.1 | *Issoria lathonia* |
| Papilionoidea | NC_015480.1 | *Calinaga davidis* |
| Papilionoidea | NC_057684.1 | *Baeotus beotus* |
| Papilionoidea | NC_065309.1 | *Danaus genutia* |
| Papilionoidea | NC_024532.1 | *Danaus chrysippus* |
| Papilionoidea | NC_021452.1 | *Danaus plexippus* |
| Papilionoidea | NC_024428.1 | *Ideopsis similis* |
| Papilionoidea | NC_063101.1 | *Parantica swinhoei* |
| Papilionoidea | NC_063100.1 | *Parantica melaneus* |
| Papilionoidea | NC_042670.1 | *Parantica aglea* |
| Papilionoidea | NC_024412.1 | *Parantica sita N1822* |
| Papilionoidea | NC_024605.1 | *Tirumala limniace* |
| Papilionoidea | NC_027516.1 | *Heliconius clysonymus* |
| Papilionoidea | NC_026564.1 | *Heliconius sara* |
| Papilionoidea | NC_026463.1 | *Heliconius ismenius* |
| Papilionoidea | NC_024864.1 | *Heliconius cydno* |
| Papilionoidea | NC_024744.1 | *Heliconius hecale* |
| Papilionoidea | NC_024741.1 | *Heliconius pachinus* |
| Papilionoidea | NC_060330.1 | *Catacroptera cloanthe* |

| | | |
|---|---|---|
| Papilionoidea | NC_052705.1 | *Doleschallia melana* |
| Papilionoidea | NC_054269.1 | *Kallima paralekta* |
| Papilionoidea | NC_016196.1 | *Kallima inachus* |
| Papilionoidea | NC_050692.1 | *Mallika jacksoni* |
| Papilionoidea | NC_016724.1 | *Libythea celtis* |
| Papilionoidea | NC_050649.1 | *Araschnia levana* |
| Papilionoidea | NC_038157.1 | *Vanessa indica* |
| Papilionoidea | NC_024413.1 | *Bhagadatta austenia* |
| Papilionoidea | NC_024417.1 | *Parthenos sylvia* |
| Papilionoidea | NC_021746.1 | *Abisara fylloides* |
| Papilionoidea | NC_058608.1 | *Dodona maculosa* |
| Papilionoidea | NC_053566.1 | *Dodona eugenes* |
| Papilionoidea | NC_046592.1 | *Zemeros flegyas* |
| Papilionoidea | NC_025551.1 | *Hamadryas epinome* |
| Papilionoidea | NC_060432.1 | *Polyura narcaeus* |
| Papilionoidea | NC_026073.1 | *Polyura nepenthes* |
| Papilionoidea | NC_024408.1 | *Polyura arja* |
| Papilionoidea | NC_026071.1 | *Cyrestis thyodamas* |
| Papilionoidea | NC_024409.1 | *Dichorragia nesimachus* |
| Papilionoidea | NC_060638.1 | *Stibochiona nicea* |
| Papilionoidea | NC_024863.1 | *Euploea midamus* |
| Papilionoidea | NC_016720.1 | *Euploea mulciber* |
| Papilionoidea | NC_024414.1 | *Euploea core* |

| | | |
|---|---|---|
| Papilionoidea | NC_018029.1 | *Melitaea cinxia* |
| Papilionoidea | NC_046470.1 | *Mellicta ambigua* |
| Papilionoidea | NC_026061.1 | *Elymnias hypermnestra* |
| Papilionoidea | NC_024406.1 | *Melanitis phedima* |
| Papilionoidea | NC_026060.1 | *Callerebia suroia* |
| Papilionoidea | NC_056106.1 | *Ypthima baldus* |
| Papilionoidea | NC_024420.1 | *Ypthima akragas* |
| Papilionoidea | NC_024571.1 | *Apodemia mormo* |
| Parastacoidea | NC_026215.1 | *Astacopsis gouldi* |
| Parastacoidea | NC_022847.1 | *Engaeus lengana* |
| Parastacoidea | NC_029407.1 | *Engaewa subcoerulea* |
| Parastacoidea | NC_029395.1 | *Engaewa walpolea* |
| Parastacoidea | NC_026214.1 | *Euastacus spinifer* |
| Parastacoidea | NC_023811.1 | *Euastacus yarraensis* |
| Parastacoidea | NC_026575.1 | *Euastacus armatus* |
| Parastacoidea | NC_023810.1 | *Geocharax gracilis* |
| Parastacoidea | NC_030531.1 | *Gramastacus insolitus* |
| Paridae | NC_053076.1 | *Anthoscopus minutus* |
| Paridae | NC_040875.1 | *Parus major* |
| Paridae | NC_028187.1 | *Parus monticolus* |
| Paridae | NC_026223.1 | *Periparus ater* |
| Paridae | NC_026911.1 | *Poecile palustris* |
| Paridae | NC_024867.1 | *Poecile atricapilla* |

| | | |
|---|---|---|
| Paridae | NC_014341.1 | *Pseudopodoces humilis* |
| Paridae | NC_021641.1 | *Remiz consobrinus* |
| Paridae | NC_026793.1 | *Sylviparus modestus* |
| Parnassiinae | NC_065029.1 | *Parnassius glacialis* |
| Parnassiinae | NC_047306.1 | *Parnassius mercurius* |
| Parnassiinae | NC_041148.1 | *Parnassius apollonius* |
| Parnassiinae | NC_014053.1 | *Parnassius bremeri* |
| Parnassiinae | NC_026864.1 | *Parnassius epaphus* |
| Parnassiinae | NC_026457.1 | *Parnassius cephalus* |
| Parnassiinae | NC_024727.1 | *Parnassius apollo* |
| Parnassiinae | NC_037863.1 | *Bhutanitis mansfieldi* |
| Parnassiinae | NC_027672.1 | *Luehdorfia chinensis* |
| Passeriformes | NC_051004.1 | *Acanthisitta chloris* |
| Passeriformes | NC_051462.1 | *Gymnorhina tibicen* |
| Passeriformes | NC_053057.1 | *Bombycilla garrulus* |
| Passeriformes | NC_053061.1 | *Phainopepla nitens* |
| Passeriformes | NC_051037.1 | *Cinclus mexicanus* |
| Passeriformes | NC_053101.1 | *Climacteris rufus* |
| Passeriformes | NC_051008.1 | *Cephalopterus ornatus* |
| Passeriformes | NC_053052.1 | *Oxyruncus cristatus* |
| Passeriformes | NC_045370.1 | *Rupicola peruvianus* |
| Passeriformes | NC_051012.1 | *Campylorhamphus procurvoides* |
| Passeriformes | NC_037154.1 | *Lepidocolaptes angustirostris* |

| | | |
|---|---|---|
| Passeriformes | NC_051470.1 | *Xiphorhynchus elegans* |
| Passeriformes | NC_053051.1 | *Serilophus lunatus* |
| Passeriformes | NC_000879.1 | *Smithornis sharpei* |
| Passeriformes | NC_051026.1 | *Formicarius rufipectus* |
| Passeriformes | NC_051009.1 | *Grallaria varia* |
| Passeriformes | NC_053074.1 | *Furnarius figulus* |
| Passeriformes | NC_051010.1 | *Sclerurus mexicanus* |
| Passeriformes | NC_024868.1 | *Hyliota flavigaster* |
| Passeriformes | NC_053102.1 | *Atrichornis clamosus* |
| Passeriformes | NC_007883.1 | *Menura novaehollandiae* |
| Passeriformes | NC_053053.1 | *Toxostoma redivivum* |
| Passeriformes | NC_031348.1 | *Moho braccatus* |
| Passeriformes | NC_029140.1 | *Notiomystis cincta* |
| Passeriformes | NC_027731.1 | *Arremon aurantiirostris* |
| Passeriformes | NC_053084.1 | *Spizella passerina* |
| Passeriformes | NC_053110.1 | *Zonotrichia albicollis* |
| Passeriformes | NC_053071.1 | *Neodrepanis coruscans* |
| Passeriformes | NC_053080.1 | *Chaetops frenatus* |
| Passeriformes | NC_053078.1 | *Picathartes gymnocephalus* |
| Passeriformes | NC_062944.1 | *Picoides kizuki* |
| Passeriformes | NC_027936.1 | *Picoides pubescens* |
| Passeriformes | NC_053111.1 | *Lepidothrix coronata* |
| Passeriformes | NC_051463.1 | *Pitta sordida* |

| | | |
|---|---|---|
| Passeriformes | NC_062819.1 | *Ailuroedus buccoides* |
| Passeriformes | NC_051014.1 | *Regulus satrapa* |
| Passeriformes | NC_024866.1 | *Regulus calendula* |
| Passeriformes | NC_051020.1 | *Rhabdornis inornatus* |
| Passeriformes | NC_053064.1 | *Scytalopus superciliaris* |
| Passeriformes | NC_051007.1 | *Chaetorhynchus papuensis* |
| Passeriformes | NC_051029.1 | *Rhipidura dahli* |
| Passeriformes | NC_029145.1 | *Rhipidura fuliginosa* |
| Passeriformes | NC_051513.1 | *Sitta villosa* |
| Passeriformes | NC_042731.1 | *Sitta nagaensis* |
| Passeriformes | NC_042730.1 | *Sitta himalayensis* |
| Passeriformes | NC_024870.1 | *Sitta carolinensis* |
| Passeriformes | NC_053059.1 | *Sitta europaea* |
| Passeriformes | NC_045387.1 | *Siva cyanouroptera* |
| Passeriformes | NC_053081.1 | *Tichodroma muraria* |
| Passeriformes | NC_042191.1 | *Culicicapa ceylonensis* |
| Passeriformes | NC_051466.1 | *Rhegmatorhina hoffmannsi* |
| Passeriformes | NC_051011.1 | *Sakesphorus luctuosus* |
| Passeriformes | NC_051013.1 | *Catharus fuscescens* |
| Passeriformes | NC_033536.1 | *Monticola gularis* |
| Passeriformes | NC_031352.1 | *Myadestes myadestinus* |
| Passeriformes | NC_054298.1 | *Zoothera aurea* |
| Passeriformes | NC_053055.1 | *Certhia brachydactyla* |

| | | |
|---|---|---|
| Passeriformes | NC_051031.1 | *Polioptila caerulea* |
| Passeriformes | NC_029482.1 | *Campylorhynchus brunneicapillus* |
| Passeriformes | NC_022840.1 | *Campylorhynchus zonatus* |
| Passeriformes | NC_024673.1 | *Henicorhina leucosticta* |
| Passeriformes | NC_051032.1 | *Thryothorus ludovicianus* |
| Passeroidea | NC_018801.1 | *Agelaius phoeniceus* |
| Passeroidea | NC_018802.1 | *Amblyramphus holosericeus* |
| Passeroidea | NC_018827.1 | *Euphagus cyanocephalus* |
| Passeroidea | NC_018795.1 | *Gnorimopsar chopi* |
| Passeroidea | NC_018812.1 | *Gymnomystax mexicanus* |
| Passeroidea | NC_018810.1 | *Macroagelaius imthurni* |
| Passeroidea | NC_051468.1 | *Molothrus ater* |
| Passeroidea | NC_018811.1 | *Molothrus badius* |
| Passeroidea | NC_018806.1 | *Molothrus aeneus* |
| Passeroidea | NC_042414.1 | *Montifringilla henrici* |
| Passeroidea | NC_025913.1 | *Montifringilla adamsi* |
| Passeroidea | NC_025914.1 | *Montifringilla taczanowskii* |
| Passeroidea | NC_022815.1 | *Montifringilla ruficollis* |
| Passeroidea | NC_018794.1 | *Nesopsar nigerrimus* |
| Passeroidea | NC_018797.1 | *Oreopsar bolivianus* |
| Passeroidea | NC_028441.1 | *Padda oryzivora* |
| Passeroidea | NC_029344.1 | *Passer ammodendri* |
| Passeroidea | NC_024821.1 | *Passer montanus* |

| | | |
|---|---|---|
| Passeroidea | NC_025611.1 | *Passer domesticus* |
| Passeroidea | NC_054361.1 | *Prunella rubeculoides* |
| Passeroidea | NC_035747.1 | *Prunella fulvescens* |
| Passeroidea | NC_031819.1 | *Prunella strophiata* |
| Passeroidea | NC_027284.1 | *Prunella montanella* |
| Passeroidea | NC_053082.1 | *Prunella himalayana* |
| Passeroidea | NC_018809.1 | *Pseudoleistes guirahuro* |
| Passeroidea | NC_018805.1 | *Pseudoleistes virescens* |
| Passeroidea | NC_025912.1 | *Pyrgilauda blanfordi* |
| Passeroidea | NC_025915.1 | *Pyrgilauda davidiana* |
| Passeroidea | NC_051021.1 | *Quiscalus mexicanus* |
| Passeroidea | NC_018803.1 | *Quiscalus quiscula* |
| Passeroidea | NC_018804.1 | *Xanthopsar flavus* |
| Passeroidea | NC_057295.1 | *Dicaeum agile* |
| Passeroidea | NC_057294.1 | *Dicaeum concolor* |
| Passeroidea | NC_051023.1 | *Dicaeum eximium* |
| Passeroidea | NC_027285.1 | *Acanthis flammea* |
| Passeroidea | NC_031349.1 | *Akialoa obscura* |
| Passeroidea | NC_051538.1 | *Carpodacus pulcherrimus* |
| Passeroidea | NC_040975.1 | *Carpodacus rubicilloides* |
| Passeroidea | NC_025597.1 | *Carpodacus erythrinus* |
| Passeroidea | NC_025607.1 | *Carpodacus roseus* |
| Passeroidea | NC_041094.1 | *Chloris sinica* |

| | | |
|---|---|---|
| Passeroidea | NC_052840.1 | *Crithagra tristriata* |
| Passeroidea | NC_031374.1 | *Eophona migratoria* |
| Passeroidea | NC_025613.1 | *Haemorhous cassinii* |
| Passeroidea | NC_025610.1 | *Haemorhous mexicanus* |
| Passeroidea | NC_025624.1 | *Hemignathus stejnegeri* |
| Passeroidea | NC_025622.1 | *Hemignathus parvus* |
| Passeroidea | NC_025608.1 | *Hemignathus flavus* |
| Passeroidea | NC_025600.1 | *Hesperiphona vespertina* |
| Passeroidea | NC_025602.1 | *Himatione sanguinea* |
| Passeroidea | NC_025604.1 | *Leucosticte brandti* |
| Passeroidea | NC_025615.1 | *Leucosticte arctoa* |
| Passeroidea | NC_051015.1 | *Loxia leucoptera* |
| Passeroidea | NC_025623.1 | *Loxia curvirostra* |
| Passeroidea | NC_025605.1 | *Loxops caeruleirostris* |
| Passeroidea | NC_025612.1 | *Loxops coccineus* |
| Passeroidea | NC_025598.1 | *Loxops mana* |
| Passeroidea | NC_025617.1 | *Melamprosops phaeosoma* |
| Passeroidea | NC_057247.1 | *Mycerobas carnipes* |
| Passeroidea | NC_025628.1 | *Oreomystis bairdi* |
| Passeroidea | NC_025601.1 | *Paroreomyza montana* |
| Passeroidea | NC_025609.1 | *Pinicola enucleator* |
| Passeroidea | NC_025630.1 | *Pseudonestor xanthophrys* |
| Passeroidea | NC_031353.1 | *Psittirostra psittacea* |

| | | |
|---|---|---|
| Passeroidea | NC_037521.1 | *Serinus canaria* |
| Passeroidea | NC_025595.1 | *Serinus albogularis* |
| Passeroidea | NC_025621.1 | *Serinus dorsostriatus* |
| Passeroidea | NC_025594.1 | *Uragus sibiricus* |
| Passeroidea | NC_053079.1 | *Urocynchramus pylzowi* |
| Passeroidea | NC_025620.1 | *Vestiaria coccinea* |
| Passeroidea | NC_053089.1 | *Melanocharis versteri* |
| Passeroidea | NC_061698.1 | *Motacilla tschutschensis taivana* |
| Passeroidea | NC_029703.1 | *Motacilla lugens* |
| Passeroidea | NC_029229.1 | *Motacilla alba* |
| Passeroidea | NC_027933.1 | *Motacilla cinerea* |
| Passeroidea | NC_027241.1 | *Aethopyga gouldiae* |
| Passeroidea | NC_051024.1 | *Leptocoma aspasia* |
| Passeroidea | NC_053093.1 | *Oreocharis arfaki* |
| Passeroidea | NC_041111.1 | *Cardellina canadensis* |
| Passeroidea | NC_051027.1 | *Setophaga kirtlandii* |
| Passeroidea | NC_053083.1 | *Calcarius ornatus* |
| Passeroidea | NC_031845.1 | *Melophus lathami* |
| Passeroidea | NC_007897.1 | *Taeniopygia guttata* |
| Passeroidea | NC_053065.1 | *Vidua macroura* |
| Passeroidea | NC_000880.1 | *Vidua chalybeata* |
| Passeroidea | NC_031157.1 | *Fringilla polatzeki* |
| Passeroidea | NC_025599.1 | *Fringilla coelebs* |

| | | |
|---|---|---|
| Passeroidea | NC_053066.1 | *Peucedramus taeniatus* |
| Passeroidea | NC_051038.1 | *Ploceus nigricollis* |
| Pecora | NC_020679.1 | *Antilocapra americana* |
| Pecora | NC_020681.1 | *Axis porcinus* |
| Pecora | NC_020682.1 | *Blastocerus dichotomus* |
| Pecora | NC_039093.1 | *Capreolus pygargus tianschanicus* |
| Pecora | NC_024819.1 | *Dama mesopotamica* |
| Pecora | NC_018358.1 | *Elaphurus davidianus* |
| Pecora | NC_020711.1 | *Hippocamelus antisensis* |
| Pecora | NC_065788.1 | *Mazama nana* |
| Pecora | NC_065375.1 | *Mazama temama* |
| Pecora | NC_024812.1 | *Mazama nemorivaga* |
| Pecora | NC_020719.1 | *Mazama americana* |
| Pecora | NC_020721.1 | *Mazama rufina* |
| Pecora | NC_020720.1 | *Mazama gouazoupira* |
| Pecora | NC_065787.1 | *Mazama bororo* |
| Pecora | NC_057642.1 | *Muntiacus vaginalis* |
| Pecora | NC_048506.1 | *Muntiacus gongshanensis* |
| Pecora | NC_041100.1 | *Muntiacus feae* |
| Pecora | NC_036430.1 | *Muntiacus putaoensis* |
| Pecora | NC_016920.1 | *Muntiacus vuquangensis* |
| Pecora | NC_004577.1 | *Muntiacus crinifrons* |
| Pecora | NC_008491.1 | *Muntiacus reevesi micrurus* |

| | | |
|---|---|---|
| Pecora | NC_015247.1 | *Odocoileus virginianus* |
| Pecora | NC_020729.1 | *Odocoileus hemionus* |
| Pecora | NC_020766.1 | *Ozotoceros bezoarticus* |
| Pecora | NC_020739.1 | *Pudu mephistophiles* |
| Pecora | NC_020740.1 | *Pudu puda* |
| Pecora | NC_007703.1 | *Rangifer tarandus* |
| Pecora | NC_045060.1 | *Rucervus duvaucelii branderi* |
| Pecora | NC_020743.1 | *Rucervus duvaucelii* |
| Pecora | NC_014701.1 | *Rucervus eldi* |
| Pecora | NC_031835.1 | *Rusa unicolor* |
| Pecora | NC_020745.1 | *Rusa timorensis* |
| Pecora | NC_020744.1 | *Rusa alfredi* |
| Pecora | NC_008414.3 | *Rusa unicolor swinhoei* |
| Pecora | NC_024820.1 | *Giraffa camelopardalis* |
| Pecora | NC_020730.1 | *Okapia johnstoni* |
| Pecora | NC_056097.1 | *Moschus cupreus* |
| Pecora | NC_042604.1 | *Moschus leucogaster* |
| Pecora | NC_020093.1 | *Moschus chrysogaster* |
| Pecora | NC_020017.1 | *Moschus anhuiensis* |
| Pecora | NC_013753.1 | *Moschus moschiferus* |
| Pecora | NC_012694.1 | *Moschus berezovskii* |
| Pecora | NC_011821.1 | *Hydropotes inermis* |
| Pecora | NC_050383.1 | *Elaphodus cephalophus cephalophus* |

| | | |
|---|---|---|
| Pelecaniformes | NC_040004.1 | *Ardea insignis* |
| Pelecaniformes | NC_025919.1 | *Ardea purpurea* |
| Pelecaniformes | NC_025918.1 | *Ardea intermedia* |
| Pelecaniformes | NC_025916.1 | *Ardea modesta* |
| Pelecaniformes | NC_025900.1 | *Ardea cinerea* |
| Pelecaniformes | NC_008551.1 | *Ardea novaehollandiae* |
| Pelecaniformes | NC_025921.1 | *Ardeola bacchus* |
| Pelecaniformes | NC_025923.1 | *Botaurus stellaris* |
| Pelecaniformes | NC_025917.1 | *Bubulcus ibis* |
| Pelecaniformes | NC_025922.1 | *Butorides striata* |
| Pelecaniformes | NC_025920.1 | *Egretta sacra* |
| Pelecaniformes | NC_023981.1 | *Egretta garzetta* |
| Pelecaniformes | NC_009736.1 | *Egretta eulophotes* |
| Pelecaniformes | NC_028195.1 | *Gorsachius melanolophus* |
| Pelecaniformes | NC_028194.1 | *Gorsachius goisagi* |
| Pelecaniformes | NC_028193.1 | *Gorsachius magnificus* |
| Pelecaniformes | NC_025925.1 | *Ixobrychus sinensis* |
| Pelecaniformes | NC_025924.1 | *Ixobrychus eurhythmus* |
| Pelecaniformes | NC_015077.1 | *Ixobrychus cinnamomeus* |
| Pelecaniformes | NC_053867.1 | *Anhinga melanogaster* |
| Pelecaniformes | NC_052797.1 | *Fregata magnificens* |
| Pelecaniformes | NC_027275.1 | *Phaethon lepturus* |
| Pelecaniformes | NC_007979.1 | *Phaethon rubricauda* |

| | | |
|---|---|---|
| Pelecaniformes | NC_027267.1 | *Phalacrocorax carbo* |
| Pelecaniformes | NC_062945.1 | *Sula dactylatra* |
| Pelecaniformes | NC_027504.1 | *Eudocimus ruber* |
| Pelecaniformes | NC_050835.1 | *Mesembrinibis cayennensis* |
| Pelecaniformes | NC_008132.1 | *Nipponia nippon* |
| Pelecaniformes | NC_012772.1 | *Platalea leucorodia* |
| Pelecaniformes | NC_010962.1 | *Platalea minor* |
| Pelecaniformes | NC_013146.1 | *Threskiornis aethiopicus* |
| Penaeoidea | NC_040139.1 | *Metapenaeopsis barbata* |
| Penaeoidea | NC_029457.1 | *Metapenaeopsis dalei* |
| Penaeoidea | NC_042173.1 | *Metapenaeus joyneri* |
| Penaeoidea | NC_039179.1 | *Metapenaeus affinis* |
| Penaeoidea | NC_026834.1 | *Metapenaeus ensis* |
| Penaeoidea | NC_038069.1 | *Parapenaeopsis hungerfordi* |
| Penaeoidea | NC_030277.1 | *Parapenaeopsis hardwickii* |
| Penaeoidea | NC_009626.1 | *Penaeus vannamei* |
| Penaeoidea | NC_040140.1 | *Penaeus latisulcatus* |
| Penaeoidea | NC_002184.1 | *Penaeus monodon* |
| Penaeoidea | NC_050695.1 | *Trachypenaeus curvirostris* |
| Penaeoidea | NC_039154.1 | *Aristeus virilis* |
| Penaeoidea | NC_039171.1 | *Benthonectes filipes* |
| Penaeoidea | NC_039170.1 | *Gennadas parvus* |
| Penaeoidea | NC_039168.1 | *Sicyonia lancifer* |

| | | |
|---|---|---|
| Penaeoidea | NC_039167.1 | *Sicyonia japonica* |
| Penaeoidea | NC_039172.1 | *Gordonella aff. paravillosa* |
| Penaeoidea | NC_039169.1 | *Hymenopenaeus neptunus* |
| Penaeoidea | NC_039964.1 | *Pleoticus muelleri* |
| Penaeoidea | NC_030280.1 | *Solenocera crassicornis* |
| Pentastomida | NC_037187.1 | *Armillifer grandis* |
| Pentastomida | NC_032061.1 | *Armillifer agkistrodontis* |
| Pentastomida | NC_039399.1 | *Linguatula serrata* |
| Pentastomida | NC_051998.1 | *Linguatula arctica* |
| Pentatomomorpha | NC_061654.1 | *Camptopus lateralis* |
| Pentatomomorpha | NC_061653.1 | *Daclera levana* |
| Pentatomomorpha | NC_061682.1 | *Grypocephalus pallipectus* |
| Pentatomomorpha | NC_061739.1 | *Leptocorisa lepida* |
| Pentatomomorpha | NC_061738.1 | *Leptocorisa acuta* |
| Pentatomomorpha | NC_061737.1 | *Leptocorisa chinensis* |
| Pentatomomorpha | NC_061680.1 | *Leptocorisa costalis* |
| Pentatomomorpha | NC_061655.1 | *Melanacanthus marginatus* |
| Pentatomomorpha | NC_061652.1 | *Paramarcius puncticeps* |
| Pentatomomorpha | NC_061681.1 | *Planusocoris schaeferi* |
| Pentatomomorpha | NC_012462.1 | *Riptortus pedestris* |
| Pentatomomorpha | NC_042441.1 | *Antilochus coquebertii* |
| Pentatomomorpha | NC_042440.1 | *Antilochus russus* |
| Pentatomomorpha | NC_042439.1 | *Dindymus rubiginosus* |

| | | |
|---|---|---|
| Pentatomomorpha | NC_042438.1 | *Dysdercus decussatus* |
| Pentatomomorpha | NC_042437.1 | *Dysdercus evanescens* |
| Pentatomomorpha | NC_012421.1 | *Dysdercus cingulatus* |
| Pentatomomorpha | NC_042436.1 | *Euscopus rufipes* |
| Pentatomomorpha | NC_042435.1 | *Melamphaus faber* |
| Pentatomomorpha | NC_042434.1 | *Melamphaus rubrocinctus* |
| Pentatomomorpha | NC_042431.1 | *Pyrrhopeplus carduelis* |
| Pentatomomorpha | NC_042430.1 | *Pyrrhopeplus posthumus* |
| Pentatomomorpha | NC_012446.1 | *Aeschyntelus notatus* |
| Pentatomomorpha | NC_061753.1 | *Liorhyssus hyalinus* |
| Pentatomomorpha | NC_046898.1 | *Myrmus lateralis* |
| Pentatomomorpha | NC_012888.1 | *Stictopleurus subviridis* |
| Pentatomomorpha | NC_042433.1 | *Physopelta cincticollis* |
| Pentatomomorpha | NC_042432.1 | *Physopelta slanbuschii* |
| Pentatomomorpha | NC_012432.1 | *Physopelta gutta* |
| Pentatomomorpha | NC_012456.1 | *Hydaropsis longirostris* |
| Perissodactyla | NC_001808.1 | *Ceratotherium simum* |
| Perissodactyla | NC_012681.1 | *Coelodonta antiquitatis* |
| Perissodactyla | NC_012684.1 | *Dicerorhinus sumatrensis* |
| Perissodactyla | NC_012682.1 | *Diceros bicornis* |
| Perissodactyla | NC_001779.1 | *Rhinoceros unicornis* |
| Perissodactyla | NC_012683.1 | *Rhinoceros sondaicus* |
| Perissodactyla | NC_063943.1 | *Tapirus bairdii* |

| | | |
|---|---|---|
| Perissodactyla | NC_053962.1 | *Tapirus terrestris* |
| Perissodactyla | NC_023838.1 | *Tapirus indicus* |
| Perlidae | NC_022843.1 | *Dinocras cephalotes* |
| Perlidae | NC_028076.1 | *Kamimuria chungnanshana* |
| Perlidae | NC_060671.1 | *Neoperla bimaculata* |
| Perlidae | NC_060586.1 | *Oyamia seminigra* |
| Perlidae | NC_056285.1 | *Oyamia nigribasis* |
| Perlidae | NC_057280.1 | *Paragnetina indentata* |
| Perlidae | NC_053853.1 | *Togoperla limbata* |
| Petroicidae | NC_053105.1 | *Drymodes brunneopygia* |
| Petroicidae | NC_027230.1 | *Eopsaltria georgiana* |
| Petroicidae | NC_019665.1 | *Eopsaltria australis* |
| Petroicidae | NC_029142.1 | *Petroica macrocephala* |
| Petroicidae | NC_029141.1 | *Petroica australis* |
| Petroicidae | NC_019668.1 | *Petroica phoenicea* |
| Petroicidae | NC_019667.1 | *Petroica goodenovii* |
| Petroicidae | NC_019666.1 | *Petroica boodang* |
| Petroicidae | NC_024871.1 | *Tregellasia leucops* |
| Petroicidae | NC_027231.1 | *Tregellasia capito* |
| Phlaeothripidae | NC_053761.1 | *Gynaikothrips ficorum* |
| Phlaeothripidae | NC_060726.1 | *Psephenothrips eriobotryae* |
| Phocidae | NC_008427.1 | *Cystophora cristata* |
| Phocidae | NC_008426.1 | *Erignathus barbatus* |

| | | |
|---|---|---|
| Phocidae | NC_001602.1 | *Halichoerus grypus* |
| Phocidae | NC_008425.1 | *Hydrurga leptonyx* |
| Phocidae | NC_008424.1 | *Leptonychotes weddellii* |
| Phocidae | NC_008423.1 | *Lobodon carcinophaga* |
| Phocidae | NC_008422.1 | *Mirounga leonina* |
| Phocidae | NC_001325.1 | *Phoca vitulina* |
| Phocidae | NC_008430.1 | *Phoca largha* |
| Phocidae | NC_008429.1 | *Phoca groenlandica* |
| Phocidae | NC_008428.1 | *Phoca fasciata* |
| Phocidae | NC_008431.1 | *Pusa caspica* |
| Phocidae | NC_008432.1 | *Pusa sibirica* |
| Phocidae | NC_008433.1 | *Pusa hispida* |
| Phrynocephalus | NC_061391.1 | *Phrynocephalus helioscopus cameranoi* |
| Phrynocephalus | NC_061390.1 | *Phrynocephalus helioscopus varius* |
| Phrynocephalus | NC_057064.1 | *Phrynocephalus nasatus* |
| Phrynocephalus | NC_020340.1 | *Phrynocephalus axillaris* |
| Phrynocephalus | NC_029186.1 | *Phrynocephalus albolineatus* |
| Phrynocephalus | NC_026454.1 | *Phrynocephalus forsythii* |
| Phrynocephalus | NC_025640.1 | *Phrynocephalus grumgrzimailoi* |
| Phrynocephalus | NC_025639.1 | *Phrynocephalus helioscopus* |
| Phrynocephalus | NC_024875.1 | *Phrynocephalus guinanensis* |
| Phrynocephalus | NC_024654.1 | *Phrynocephalus versicolor* |
| Phrynocephalus | NC_022719.1 | *Phrynocephalus przewalskii* |

| | | |
|---|---|---|
| Phyllostomidae | NC_065676.1 | *Anoura geoffroyi* |
| Phyllostomidae | NC_065675.1 | *Anoura cultrata* |
| Phyllostomidae | NC_022420.1 | *Anoura caudifer* |
| Phyllostomidae | NC_065681.1 | *Artibeus hartii* |
| Phyllostomidae | NC_016871.1 | *Artibeus lituratus* |
| Phyllostomidae | NC_002009.1 | *Artibeus jamaicensis* |
| Phyllostomidae | NC_065679.1 | *Chiroderma salvini* |
| Phyllostomidae | NC_065683.1 | *Choeroniscus minor* |
| Phyllostomidae | NC_037132.1 | *Chrotopterus auritus* |
| Phyllostomidae | NC_065680.1 | *Dermanura rava* |
| Phyllostomidae | NC_041639.1 | *Ectophylla alba* |
| Phyllostomidae | NC_065682.1 | *Glossophaga soricina* |
| Phyllostomidae | NC_037139.1 | *Glyphonycteris daviesi* |
| Phyllostomidae | NC_066832.1 | *Leptonycteris curasoae* |
| Phyllostomidae | NC_066831.1 | *Leptonycteris yerbabuenae* |
| Phyllostomidae | NC_066830.1 | *Leptonycteris nivalis* |
| Phyllostomidae | NC_037135.1 | *Lonchorhina aurita* |
| Phyllostomidae | NC_065678.1 | *Lophostoma brasiliense* |
| Phyllostomidae | NC_022424.1 | *Lophostoma silvicolum* |
| Phyllostomidae | NC_037136.1 | *Macrotus californicus* |
| Phyllostomidae | NC_065685.1 | *Micronycteris hirsuta* |
| Phyllostomidae | NC_022419.1 | *Micronycteris megalotis* |
| Phyllostomidae | NC_065690.1 | *Phyllostomus discolor* |

| | | |
|---|---|---|
| Phyllostomidae | NC_065686.1 | *Platyrrhinus matapalensis* |
| Phyllostomidae | NC_065691.1 | *Sturnira ludovici* |
| Phyllostomidae | NC_022427.1 | *Sturnira tildae* |
| Phyllostomidae | NC_065687.1 | *Sturnira bakeri* |
| Phyllostomidae | NC_022428.1 | *Tonatia saurophila* |
| Phyllostomidae | NC_022429.1 | *Vampyrum spectrum* |
| Phymatidae | NC_037733.1 | *Amblythyreus gestroi* |
| Phymatidae | NC_037734.1 | *Carcinochelis bannaensis* |
| Phymatidae | NC_036012.1 | *Carcinocoris binghami* |
| Phymatidae | NC_036013.1 | *Cnizocoris sinensis* |
| Phymatidae | NC_036011.1 | *Phymata americana* |
| Piciformes | NC_034278.1 | *Campephilus imperialis* |
| Piciformes | NC_028020.1 | *Campephilus guatemalensis* |
| Piciformes | NC_042683.1 | *Dendrocopos darjellensis* |
| Piciformes | NC_041121.1 | *Dendrocopos canicapillus* |
| Piciformes | NC_029862.1 | *Dendrocopos leucotos* |
| Piciformes | NC_028174.1 | *Dendrocopos major* |
| Piciformes | NC_061952.1 | *Dryobates minor* |
| Piciformes | NC_008546.1 | *Dryocopus pileatus* |
| Piciformes | NC_056091.1 | *Jynx torquilla* |
| Piciformes | NC_039890.1 | *Jynx ruficollis* |
| Piciformes | NC_039537.1 | *Picumnus innominatus* |
| Piciformes | NC_045372.1 | *Picus canus* |

| | | |
|---|---|---|
| Piciformes | NC_028019.1 | *Sasia ochracea* |
| Piciformes | NC_040005.1 | *Indicator xanthonotus* |
| Piciformes | NC_039889.1 | *Indicator maculatus* |
| Piciformes | NC_039892.1 | *Prodotiscus insignis* |
| Piciformes | NC_039891.1 | *Megalaima virens* |
| Piciformes | NC_052781.1 | *Psilopogon haemacephalus* |
| Piciformes | NC_008549.1 | *Pteroglossus azara flavirostris* |
| Pieridae | NC_022687.1 | *Catopsilia pomona* |
| Pieridae | NC_053715.1 | *Colias fieldii* |
| Pieridae | NC_027253.1 | *Colias erate* |
| Pieridae | NC_032285.1 | *Eurema blanda* |
| Pieridae | NC_022685.1 | *Eurema hecabe* |
| Pieridae | NC_026837.1 | *Gonepteryx mahaguru* |
| Pieridae | NC_026046.1 | *Gonepteryx rhamni* |
| Pieridae | NC_022686.1 | *Leptidea morsei* |
| Pieridae | NC_057569.1 | *Anthocharis scolymus* |
| Pieridae | NC_025274.1 | *Anthocharis bambusarum* |
| Pieridae | NC_058011.1 | *Aporia largeteaui* |
| Pieridae | NC_033890.1 | *Aporia martineti* |
| Pieridae | NC_033888.1 | *Aporia bieti* |
| Pieridae | NC_033889.1 | *Aporia hippia* |
| Pieridae | NC_025273.1 | *Aporia intercostata* |
| Pieridae | NC_018346.1 | *Aporia crataegi* |

| | | |
|---|---|---|
| Pieridae | NC_053734.1 | *Appias albina* |
| Pieridae | NC_053273.1 | *Appias lyncida* |
| Pieridae | NC_053272.1 | *Appias nero* |
| Pieridae | NC_039715.1 | *Appias remedios* |
| Pieridae | NC_045189.1 | *Baltia butleri* |
| Pieridae | NC_047457.1 | *Delias pasithoe* |
| Pieridae | NC_020428.1 | *Delias hyparete* |
| Pieridae | NC_033891.1 | *Mesapia peloria* |
| Pieridae | NC_061692.1 | *Pieris napi napi* |
| Pieridae | NC_015895.1 | *Pieris rapae* |
| Pieridae | NC_026532.1 | *Pieris canidia* |
| Pieridae | NC_010568.1 | *Pieris melete* |
| Pieridae | NC_047456.1 | *Pontia edusa* |
| Pieridae | NC_045191.1 | *Pontia callidice* |
| Pieridae | NC_053274.1 | *Prioneris clemanthe* |
| Pitheciidae | NC_024630.1 | *Callicebus lugens* |
| Pitheciidae | NC_021965.1 | *Callicebus cupreus* |
| Pitheciidae | NC_019801.1 | *Callicebus donacophilus* |
| Pitheciidae | NC_064162.1 | *Cacajao calvus ucayalii* |
| Pitheciidae | NC_064154.1 | *Cacajao calvus calvus* |
| Pitheciidae | NC_064153.1 | *Cacajao ayresi* |
| Pitheciidae | NC_064152.1 | *Cacajao melanocephalus* |
| Pitheciidae | NC_021967.1 | *Cacajao calvus* |

| | | |
|---|---|---|
| Pitheciidae | NC_064208.1 | *Plecturocebus moloch* |
| Pitheciidae | NC_064207.1 | *Plecturocebus miltoni* |
| Pitheciidae | NC_064206.1 | *Plecturocebus hoffmannsi* |
| Pitheciidae | NC_064205.1 | *Plecturocebus grovesi* |
| Pitheciidae | NC_064204.1 | *Plecturocebus dubius U* |
| Pitheciidae | NC_064203.1 | *Plecturocebus cinerascens* |
| Pitheciidae | NC_064201.1 | *Plecturocebus brunneus UF* |
| Pitheciidae | NC_064200.1 | *Plecturocebus bernhardi* |
| Pitheciidae | NC_064199.1 | *Cheracebus lucifer* |
| Pitheciidae | NC_064198.1 | *Cheracebus torquatus* |
| Pitheciidae | NC_064197.1 | *Cheracebus regulus* |
| Pitheciidae | NC_021946.1 | *Chiropotes albinasus* |
| Pitheciidae | NC_024629.1 | *Chiropotes israelita* |
| Pitheciidae | NC_064161.1 | *Pithecia albicans* |
| Pitheciidae | NC_064160.1 | *Pithecia mittermeieri* |
| Pitheciidae | NC_064159.1 | *Pithecia hirsuta* |
| Platypezoidea | NC_056116.1 | *Diplonevra peregrina* |
| Platypezoidea | NC_056115.1 | *Diplonevra funebris* |
| Platypezoidea | NC_023794.1 | *Megaselia scalaris* |
| Platyrrhini | NC_064163.1 | *Aotus vociferans* |
| Platyrrhini | NC_021939.1 | *Aotus azarai* |
| Platyrrhini | NC_019799.1 | *Aotus lemurinus* |
| Platyrrhini | NC_018116.1 | *Aotus nancymaae* |

| | | |
|---|---|---|
| Platyrrhini | NC_064196.1 | *Alouatta juara* |
| Platyrrhini | NC_064195.1 | *Alouatta belzebul* |
| Platyrrhini | NC_064193.1 | *Alouatta palliata* |
| Platyrrhini | NC_064189.1 | *Alouatta discolor* |
| Platyrrhini | NC_064188.1 | *Alouatta seniculus puruensis* |
| Platyrrhini | NC_064187.1 | *Alouatta macconnelli* |
| Platyrrhini | NC_064185.1 | *Alouatta caraya* |
| Platyrrhini | NC_064186.1 | *Alouatta guariba clamitans* |
| Platyrrhini | NC_024628.1 | *Callimico goeldii* |
| Platyrrhini | NC_050682.1 | *Callithrix aurita* |
| Platyrrhini | NC_030788.1 | *Callithrix penicillata* |
| Platyrrhini | NC_021941.1 | *Callithrix geoffroyi* |
| Platyrrhini | NC_021942.1 | *Callithrix pygmaea* |
| Platyrrhini | NC_027658.1 | *Callithrix kuhlii* |
| Platyrrhini | NC_025586.1 | *Callithrix jacchus* |
| Platyrrhini | NC_064170.1 | *Cebuella niveiventris* |
| Platyrrhini | NC_064171.1 | *Leontocebus nigricollis* |
| Platyrrhini | NC_064181.1 | *Leontocebus fuscicollis* |
| Platyrrhini | NC_064177.1 | *Leontocebus fuscicollis weddelli* |
| Platyrrhini | NC_037878.1 | *Leontopithecus chrysopygus* |
| Platyrrhini | NC_021952.1 | *Leontopithecus rosalia* |
| Platyrrhini | NC_064164.1 | *Mico humilis* |
| Platyrrhini | NC_064175.1 | *Mico argentatus* |

| | | |
|---|---|---|
| Platyrrhini | NC_064174.1 | *Mico humeralifer* |
| Platyrrhini | NC_064179.1 | *Saguinus midas* |
| Platyrrhini | NC_064178.1 | *Saguinus labiatus rufiventer* |
| Platyrrhini | NC_064166.1 | *Saguinus inustus* |
| Platyrrhini | NC_064180.1 | *Saguinus mystax* |
| Platyrrhini | NC_064176.1 | *Saguinus bicolor* |
| Platyrrhini | NC_064172.1 | *Saguinus geoffroyi* |
| Platyrrhini | NC_021960.1 | *Saguinus oedipus* |
| Platyrrhini | NC_064168.1 | *Saimiri ustus* |
| Platyrrhini | NC_064183.1 | *Saimiri macrodon* |
| Platyrrhini | NC_064182.1 | *Saimiri cassiquiarensis* |
| Platyrrhini | NC_064173.1 | *Saimiri oerstedii* |
| Platyrrhini | NC_021966.1 | *Saimiri boliviensis* |
| Platyrrhini | NC_064194.1 | *Ateles geoffroyi* |
| Platyrrhini | NC_064192.1 | *Ateles marginatus* |
| Platyrrhini | NC_064191.1 | *Ateles paniscus* |
| Platyrrhini | NC_019800.1 | *Ateles belzebuth* |
| Platyrrhini | NC_021951.1 | *Lagothrix lagotricha* |
| Platyrrhini | NC_064169.1 | *Cebus unicolor* |
| Platyrrhini | NC_064165.1 | *Cebus olivaceus* |
| Platyrrhini | NC_021961.1 | *Cebus xanthosternos* |
| Platyrrhini | NC_064167.1 | *Sapajus apella macrocephalus* |
| Plecoptera | NC_042205.1 | *Diamphipnoa annulata* |

| | | |
|---|---|---|
| Plecoptera | NC_027698.1 | *Apteroperla tikumana* |
| Plecoptera | NC_064508.1 | *Capnia yunnana* |
| Plecoptera | NC_034661.1 | *Capnia zijinshana* |
| Plecoptera | NC_042206.1 | *Neonemura barrosi* |
| Plecoptera | NC_029248.1 | *Pteronarcella badia* |
| Plecoptera | NC_006133.1 | *Pteronarcys princeps* |
| Plecoptera | NC_034809.1 | *Styloperla spinicercia* |
| Plecoptera | NC_041105.1 | *Scopura longa* |
| Plecoptera | NC_042200.1 | *Neuroperla schedingi* |
| Plecoptera | NC_053557.1 | *Paraleuctra cercia* |
| Plecoptera | NC_053558.1 | *Perlomyia isobeae* |
| Plecoptera | NC_042207.1 | *Rhopalopsole bulbifera* |
| Plecoptera | NC_061407.1 | *Nemoura longicercia* |
| Plecoptera | NC_057513.1 | *Nemoura meniscata* |
| Plecoptera | NC_034939.1 | *Nemoura nankinensis* |
| Plecoptera | NC_060430.1 | *Protonemura datongensis* |
| Plecoptera | NC_050322.1 | *Protonemura meyeri* |
| Plecoptera | NC_044753.1 | *Protonemura orbiculata* |
| Plecoptera | NC_044752.1 | *Protonemura kohnoae* |
| Plecoptera | NC_065772.1 | *Strophopteryx fasciata* |
| Plecoptera | NC_037897.1 | *Taeniopteryx ugola* |
| Plecoptera | NC_037754.1 | *Suwallia teleckojensis* |
| Plecoptera | NC_065830.1 | *Cryptoperla kawasawai* |

| | | |
|---|---|---|
| Plecoptera | NC_065831.1 | *Peltoperlopsis sagittata* |
| Plecoptera | NC_038189.1 | *Soliperla sp. ZTC-2018* |
| Plecoptera | NC_053852.1 | *Acroneuria carolinensis* |
| Plecoptera | NC_026104.1 | *Acroneuria hainana* |
| Plecoptera | NC_057436.1 | *Flavoperla hatakeyamae* |
| Plecoptera | NC_057281.1 | *Perlesta teaysia* |
| Plecoptera | NC_038167.1 | *Isoperla eximia* |
| Plecoptera | NC_038190.1 | *Isoperla bilineata* |
| Plecoptera | NC_059845.1 | *Arcynopteryx dichroa* |
| Plecoptera | NC_038168.1 | *Pseudomegarcys japonica* |
| Pleocyemata | NC_031154.1 | *Callianassa ceramica* |
| Pleocyemata | NC_020025.1 | *Corallianassa coutierei* |
| Pleocyemata | NC_034298.1 | *Nihonotrypaea harmandi* |
| Pleocyemata | NC_020351.1 | *Nihonotrypaea japonica* |
| Pleocyemata | NC_019610.1 | *Nihonotrypaea thermophila* |
| Pleocyemata | NC_024651.1 | *Paraglypturus tonganus* |
| Pleocyemata | NC_026225.2 | *Trypaea australiensis* |
| Pleocyemata | NC_019609.1 | *Neaxius glyptocercus* |
| Pleocyemata | NC_043834.1 | *Strahlaxius plectrorhynchus* |
| Pleocyemata | NC_038166.1 | *Spongiocaris panglao* |
| Pleocyemata | NC_061665.1 | *Stenopus scutellatus* |
| Pleocyemata | NC_018097.1 | *Stenopus hispidus* |
| Plethodontidae | NC_006340.1 | *Batrachoseps attenuatus* |

| | | |
|---|---|---|
| Plethodontidae | NC_028184.1 | *Batrachoseps nigriventris* |
| Plethodontidae | NC_006337.1 | *Desmognathus wrighti* |
| Plethodontidae | NC_006339.1 | *Desmognathus fuscus* |
| Plethodontidae | NC_035494.1 | *Eurycea cirrigera* |
| Plethodontidae | NC_006329.1 | *Eurycea bislineata* |
| Plethodontidae | NC_028297.1 | *Gyrinophilus palleucus* |
| Plethodontidae | NC_006341.1 | *Gyrinophilus porphyriticus* |
| Plethodontidae | NC_006342.1 | *Hemidactylium scutatum* |
| Plethodontidae | NC_006345.1 | *Hydromantes brunus* |
| Plethodontidae | NC_006326.1 | *Oedipina poelzi* |
| Plethodontidae | NC_006344.1 | *Phaeognathus hubrichti* |
| Plethodontidae | NC_006335.1 | *Plethodon elongatus* |
| Plethodontidae | NC_006332.1 | *Pseudotriton ruber* |
| Plethodontidae | NC_006336.1 | *Thorius n. sp.* |
| Poduromorpha | NC_046523.1 | *Ceratophysella communis* |
| Poduromorpha | NC_005438.1 | *Gomphiocephalus hodgsoni* |
| Poduromorpha | NC_057969.1 | *Friesea gretae* |
| Poduromorpha | NC_056239.1 | *Friesea propria* |
| Poduromorpha | NC_010535.1 | *Friesea antarctica* |
| Poduromorpha | NC_054366.1 | *Yuukianura szeptyckii* |
| Poduromorpha | NC_011195.1 | *Bilobella aurantiaca* |
| Poduromorpha | NC_053646.1 | *Allonychiurus kimi* |
| Poduromorpha | NC_002735.1 | *Tetrodontophora bielanensis* |

| | | |
|---|---|---|
| Poduromorpha | NC_056908.1 | *Tullbergia mixta* |
| Prioninae | NC_038089.1 | *Aegosoma sinicum* |
| Prioninae | NC_037698.1 | *Callipogon relictus* |
| Prioninae | NC_045407.1 | *Megopis sinica* |
| Prioninae | NC_037927.1 | *Dorysthenes paradoxus* |
| Proboscidea | NC_035230.1 | *Elephas antiquus* |
| Proboscidea | NC_005129.2 | *Elephas maximus* |
| Proboscidea | NC_000934.1 | *Loxodonta africana* |
| Proboscidea | NC_020759.1 | *Loxodonta cyclotis* |
| Proboscidea | NC_035800.1 | *Mammut americanum* |
| Proboscidea | NC_007596.2 | *Mammuthus primigenius* |
| Proboscidea | NC_015529.1 | *Mammuthus columbi* |
| Procellariiformes | NC_026190.1 | *Phoebastria albatrus* |
| Procellariiformes | NC_026189.1 | *Phoebastria immutabilis* |
| Procellariiformes | NC_026188.1 | *Phoebastria nigripes* |
| Procellariiformes | NC_007172.2 | *Thalassarche melanophrys* |
| Procellariiformes | NC_052777.1 | *Oceanites oceanicus* |
| Procellariiformes | NC_052809.1 | *Pelecanoides urinatrix* |
| Procellariiformes | NC_057528.1 | *Ardenna pacifica* |
| Procellariiformes | NC_057527.1 | *Ardenna carneipes* |
| Procellariiformes | NC_043899.1 | *Daption capense* |
| Proechimys | NC_039551.1 | *Proechimys trinitatis* |
| Proechimys | NC_039550.1 | *Proechimys steerei* |

| | | |
|---|---|---|
| Proechimys | NC_039548.1 | *Proechimys pattoni* |
| Proechimys | NC_039547.1 | *Proechimys mincae* |
| Proechimys | NC_039546.1 | *Proechimys kulinae* |
| Proechimys | NC_039545.1 | *Proechimys goeldii* |
| Proechimys | NC_039544.1 | *Proechimys gardneri* |
| Proechimys | NC_039444.1 | *Proechimys simonsi* |
| Proechimys | NC_039420.1 | *Proechimys roberti* |
| Proechimys | NC_039419.1 | *Proechimys hoplomyoides* |
| Proechimys | NC_039409.1 | *Proechimys echinothrix* |
| Proechimys | NC_039370.1 | *Proechimys quadruplicatus* |
| Proechimys | NC_039102.1 | *Proechimys semispinosus calidior* |
| Proechimys | NC_039103.1 | *Proechimys semispinosus* |
| Proechimys | NC_039101.1 | *Proechimys gularis* |
| Proechimys | NC_039100.1 | *Proechimys brevicauda* |
| Proechimys | NC_039099.1 | *Proechimys cuvieri* |
| Proechimys | NC_020657.1 | *Proechimys longicaudatus* |
| Proechimys | NC_039098.1 | *Proechimys guyannensis* |
| Proechimys | NC_039549.1 | *Proechimys poliopus* |
| Proechimys | NC_039369.1 | *Proechimys guairae* |
| Protobothrops | NC_030182.1 | *Protobothrops tokarensis* |
| Protobothrops | NC_030181.1 | *Protobothrops flavoviridis* |
| Protobothrops | NC_029166.1 | *Protobothrops kaulbacki* |
| Protobothrops | NC_029165.1 | *Protobothrops himalayanus* |

| | | |
|---|---|---|
| Protobothrops | NC_026052.1 | *Protobothrops mangshanensis* |
| Protobothrops | NC_026051.1 | *Protobothrops maolanensis* |
| Protobothrops | NC_022473.1 | *Protobothrops dabieshanensis* |
| Protobothrops | NC_021412.1 | *Protobothrops mucrosquamatus* |
| Protobothrops | NC_022695.1 | *Protobothrops cornutus* |
| Psittaciformes | NC_045369.1 | *Agapornis lilianae* |
| Psittaciformes | NC_011708.1 | *Agapornis roseicollis* |
| Psittaciformes | NC_063712.1 | *Amazona auropalliata* |
| Psittaciformes | NC_034679.1 | *Amazona ventralis* |
| Psittaciformes | NC_033336.1 | *Amazona aestiva* |
| Psittaciformes | NC_027840.1 | *Amazona ochrocephala* |
| Psittaciformes | NC_052783.1 | *Amazona guildingii* |
| Psittaciformes | NC_047199.1 | *Ara chloropterus* |
| Psittaciformes | NC_045076.1 | *Ara macao* |
| Psittaciformes | NC_029319.1 | *Ara ararauna* |
| Psittaciformes | NC_026029.1 | *Ara glaucogularis* |
| Psittaciformes | NC_037895.1 | *Ara tricolor* |
| Psittaciformes | NC_027839.1 | *Ara militaris* |
| Psittaciformes | NC_045371.1 | *Aratinga nenday* |
| Psittaciformes | NC_026039.1 | *Aratinga solstitialis* |
| Psittaciformes | NC_015530.1 | *Brotogeris cyanoptera* |
| Psittaciformes | NC_059712.1 | *Cacatua goffiniana* |
| Psittaciformes | NC_059711.1 | *Cacatua galerita* |

| | | |
|---|---|---|
| Psittaciformes | NC_059710.1 | *Cacatua alba* |
| Psittaciformes | NC_040142.1 | *Cacatua pastinator* |
| Psittaciformes | NC_020592.1 | *Cacatua moluccensis* |
| Psittaciformes | NC_020593.1 | *Calyptorhynchus lathami* |
| Psittaciformes | NC_020595.1 | *Calyptorhynchus latirostris* |
| Psittaciformes | NC_027841.1 | *Coracopsis vasa* |
| Psittaciformes | NC_052711.1 | *Diopsittaca nobilis* |
| Psittaciformes | NC_027842.1 | *Eclectus roratus* |
| Psittaciformes | NC_040154.1 | *Eolophus roseicapillus* |
| Psittaciformes | NC_015197.1 | *Eupsittula pertinax chrysogenys* |
| Psittaciformes | NC_027843.1 | *Forpus passerinus* |
| Psittaciformes | NC_026031.1 | *Guaruba guarouba* |
| Psittaciformes | NC_027844.1 | *Myiopsitta monachus* |
| Psittaciformes | NC_019804.1 | *Neophema chrysogaster* |
| Psittaciformes | NC_027845.1 | *Nestor notabilis* |
| Psittaciformes | NC_015192.1 | *Nymphicus hollandicus* |
| Psittaciformes | NC_029161.1 | *Orthopsittaca manilata* |
| Psittaciformes | NC_044184.1 | *Pionites leucogaster* |
| Psittaciformes | NC_044083.1 | *Poicephalus senegalus* |
| Psittaciformes | NC_029322.1 | *Primolius maracana* |
| Psittaciformes | NC_025742.1 | *Primolius couloni* |
| Psittaciformes | NC_027846.1 | *Prioniturus luconensis* |
| Psittaciformes | NC_041257.1 | *Psittacara leucophthalmus* |

| | | |
|---|---|---|
| Psittaciformes | NC_026042.1 | *Psittacara rubritorquis* |
| Psittaciformes | NC_021764.1 | *Psittacara brevipes* |
| Psittaciformes | NC_020325.1 | *Psittacara acuticaudatus acuticaudatus* |
| Psittaciformes | NC_054153.1 | *Psittacula cyanocephala* |
| Psittaciformes | NC_045379.1 | *Psittacula roseata* |
| Psittaciformes | NC_045378.1 | *Psittacula alexandri* |
| Psittaciformes | NC_042765.1 | *Psittacula eupatria* |
| Psittaciformes | NC_027848.1 | *Psittrichas fulgidus* |
| Psittaciformes | NC_028404.1 | *Pyrrhura rupicola* |
| Psittaciformes | NC_021771.1 | *Rhynchopsitta terrisi* |
| Psittaciformes | NC_005931.1 | *Strigops habroptilus* |
| Psittaciformes | NC_009134.1 | *Melopsittacus undulatus* |
| Psittaciformes | NC_031358.1 | *Psephotellus pulcherrimus* |
| Psittaciformes | NC_066152.1 | *Trichoglossus moluccanus* |
| Psittaciformes | NC_066151.1 | *Trichoglossus forsteni* |
| Psoroptidia | NC_041090.1 | *Trouessartia rubecula* |
| Psoroptidia | NC_024675.1 | *Psoroptes cuniculi* |
| Psoroptidia | NC_012218.1 | *Dermatophagoides pteronyssinus* |
| Psoroptidia | NC_013184.1 | *Dermatophagoides farinae* |
| Psychodoidea | NC_026898.1 | *Nyssomyia umbratilis* |
| Psychodoidea | NC_028042.1 | *Phlebotomus papatasi* |
| Psychodoidea | NC_028041.1 | *Phlebotomus chinensis* |
| Pteropodidae | NC_061537.1 | *Balionycteris maculata* |

| | | |
|---|---|---|
| Pteropodidae | NC_046902.1 | *Cynopterus sphinx* |
| Pteropodidae | NC_026465.1 | *Cynopterus brachyotis* |
| Pteropodidae | NC_046903.1 | *Eidolon helvum* |
| Pteropodidae | NC_046915.1 | *Megaerops niphanae* |
| Pteropodidae | NC_046924.1 | *Nyctimene cephalotes* |
| Pteropodidae | NC_046926.1 | *Pteropus ornatus* |
| Pteropodidae | NC_026542.1 | *Pteropus vampyrus* |
| Pteropodidae | NC_023122.1 | *Pteropus alecto* |
| Pteropodidae | NC_002619.1 | *Pteropus scapulatus* |
| Pteropodidae | NC_002612.1 | *Pteropus dasymallus* |
| Pteropodidae | NC_007393.1 | *Rousettus aegyptiacus* |
| Pteropodidae | NC_046934.1 | *Rousettus lanosus* |
| Pteropodidae | NC_046929.1 | *Rousettus obliviosus* |
| Pteropodidae | NC_046928.1 | *Rousettus madagascariensis* |
| Pteropodidae | NC_046927.1 | *Rousettus leschenaultii* |
| Pteropodidae | NC_045044.1 | *Rousettus amplexicaudatus* |
| Pteropodinae | NC_046909.1 | *Epomophorus minor* |
| Pteropodinae | NC_046907.1 | *Epomophorus wahlbergi* |
| Pteropodinae | NC_046906.1 | *Epomophorus labiatus* |
| Pteropodinae | NC_046905.1 | *Epomophorus crypturus* |
| Pteropodinae | NC_029375.1 | *Epomophorus gambianus* |
| Pteropodinae | NC_046908.1 | *Epomophorus minimus* |
| Pteropodinae | NC_046912.1 | *Epomops franqueti* |

| | | |
|---|---|---|
| Pteropodinae | NC_046911.1 | *Epomops dobsonii* |
| Pteropodinae | NC_046910.1 | *Epomops buettikoferi* |
| Pteropodinae | NC_046917.1 | *Megaloglossus woermanni* |
| Pteropodinae | NC_046916.1 | *Megaloglossus azagnyi* |
| Pteropodinae | NC_046922.1 | *Myonycteris torquata* |
| Pteropodinae | NC_046921.1 | *Myonycteris relicta* |
| Pteropodinae | NC_046920.1 | *Myonycteris leptodon* |
| Pteropodinae | NC_046919.1 | *Myonycteris angolensis* |
| Pyralidae | NC_061244.1 | *Acrobasis inouei* |
| Pyralidae | NC_058009.1 | *Aglossa dimidiata* |
| Pyralidae | NC_028443.1 | *Amyelois transitella* |
| Pyralidae | NC_061242.1 | *Dioryctria rubella* |
| Pyralidae | NC_061240.1 | *Dusungwua basinigra* |
| Pyralidae | NC_061642.1 | *Endotricha kuznetzovi* |
| Pyralidae | NC_037501.1 | *Endotricha consocia* |
| Pyralidae | NC_061241.1 | *Endotricha olivacealis* |
| Pyralidae | NC_039716.1 | *Ephestia elutella* |
| Pyralidae | NC_022476.1 | *Ephestia kuehniella* |
| Pyralidae | NC_037175.1 | *Euzophera pyriella* |
| Pyralidae | NC_030508.1 | *Hypsopygia regina* |
| Pyralidae | NC_035242.1 | *Meroptera pravella* |
| Pyralidae | NC_061247.1 | *Orybina regalis* |
| Pyralidae | NC_066226.1 | *Perula sp.* |

| | | |
|---|---|---|
| Pyralidae | NC_027961.1 | *Plodia interpunctella* |
| Pyralidae | NC_047303.1 | *Pyralis farinalis* |
| Pyralidae | NC_024535.1 | *Lista haraldusalis* |
| Pyralidae | NC_061246.1 | *Orthaga euadrusalis* |
| Pyralidae | NC_046504.1 | *Orthaga olivacea* |
| Pyralidae | NC_061604.1 | *Achroia grisella* |
| Pyralidae | NC_053657.1 | *Cathayia obliquella* |
| Pyralidae | NC_028532.1 | *Galleria mellonella* |
| Pyralidae | NC_062173.1 | *Lamoria adaptella* |
| Pyralidae | NC_054356.1 | *Paralipsa gularis* |
| Pyrgomorphoidea | NC_046552.1 | *Atractomorpha psittacina* |
| Pyrgomorphoidea | NC_011824.1 | *Atractomorpha sinensis* |
| Pyrgomorphoidea | NC_014450.1 | *Mekongiana xiangchengensis* |
| Pyrgomorphoidea | NC_023921.1 | *Mekongiella kingdoni* |
| Pyrgomorphoidea | NC_014451.1 | *Mekongiella xizangensis* |
| Pyrgomorphoidea | NC_045930.1 | *Tagasta indica* |
| Rana | NC_061371.1 | *Rana hanluica* |
| Rana | NC_061370.1 | *Rana longicrus* |
| Rana | NC_058599.1 | *Rana johnsi* |
| Rana | NC_056272.1 | *Rana uenoi* |
| Rana | NC_035805.1 | *Rana omeimontis* |
| Rana | NC_035804.1 | *Rana kukunoris* |
| Rana | NC_035803.1 | *Rana chaochiaoensis* |

| | | |
|---|---|---|
| Rana | NC_027236.1 | *Rana sylvatica* |
| Rana | NC_030042.1 | *Rana amurensis* |
| Rana | NC_028521.1 | *Rana huanrensis* |
| Rana | NC_028296.1 | *Rana draytonii* |
| Rana | NC_028283.1 | *Rana okaloosae* |
| Rana | NC_023529.1 | *Rana cf. chensinensis* |
| Rana | NC_023528.1 | *Rana dybowskii* |
| Rana | NC_022696.1 | *Rana catesbeiana* |
| Rana | NC_016059.1 | *Rana chosenica* |
| Rana | NC_009264.1 | *Rana plancyi* |
| Rana | NC_002805.1 | *Rana nigromaculata* |
| Ranoidea | NC_044901.1 | *Amolops granulosus* |
| Ranoidea | NC_029250.1 | *Amolops loloensis* |
| Ranoidea | NC_025591.1 | *Amolops wuyiensis* |
| Ranoidea | NC_024180.1 | *Amolops mantzorum* |
| Ranoidea | NC_023949.1 | *Amolops ricketti* |
| Ranoidea | NC_009423.1 | *Amolops tormotus* |
| Ranoidea | NC_018771.1 | *Babina adenopleura* |
| Ranoidea | NC_025226.1 | *Glandirana tientaiensis* |
| Ranoidea | NC_057198.1 | *Hylarana latouchii* |
| Ranoidea | NC_024748.1 | *Hylarana guentheri* |
| Ranoidea | NC_065297.1 | *Odorrana jingdongensis* |
| Ranoidea | NC_053712.1 | *Odorrana exiliversabilis* |

| | | |
|---|---|---|
| Ranoidea | NC_050884.1 | *Odorrana graminea* |
| Ranoidea | NC_034984.1 | *Odorrana hainanensis* |
| Ranoidea | NC_034983.1 | *Odorrana wuchuanensis* |
| Ranoidea | NC_027827.1 | *Odorrana schmackeri* |
| Ranoidea | NC_024603.1 | *Odorrana margaretae* |
| Ranoidea | NC_029201.1 | *Pelophylax cf. bedriagae* |
| Ranoidea | NC_029200.1 | *Pelophylax bedriagae* |
| Ranoidea | NC_029199.1 | *Pelophylax cf. terentievi* |
| Ranoidea | NC_026896.1 | *Pelophylax shqipericus* |
| Ranoidea | NC_026895.1 | *Pelophylax kurtmuelleri* |
| Ranoidea | NC_026894.1 | *Pelophylax epeiroticus* |
| Ranoidea | NC_026893.1 | *Pelophylax cypriensis* |
| Ranoidea | NC_025575.1 | *Pelophylax cretensis* |
| Ranoidea | NC_029754.1 | *Fejervarya multistriata* |
| Ranoidea | NC_012647.1 | *Fejervarya cancrivora* |
| Ranoidea | NC_007440.2 | *Limnonectes fujianensis* |
| Ranoidea | NC_039094.1 | *Nanorana ventripunctata* |
| Ranoidea | NC_026789.1 | *Nanorana parkeri* |
| Ranoidea | NC_024272.1 | *Nanorana taihangnica* |
| Ranoidea | NC_016119.1 | *Nanorana pleskei* |
| Ranoidea | NC_056269.1 | *Quasipaa exilispinosa* |
| Ranoidea | NC_024843.1 | *Quasipaa yei* |
| Ranoidea | NC_021937.1 | *Quasipaa boulengeri* |

| | | |
|---|---|---|
| Ranoidea | NC_013270.1 | *Quasipaa spinosa* |
| Ranoidea | NC_014685.1 | *Occidozyga martensii* |
| Ranoidea | NC_057992.1 | *Phrynoglossus myanhessei* |
| Rattus | NC_012374.1 | *Rattus rattus* |
| Rattus | NC_049042.1 | *Rattus korinchi* |
| Rattus | NC_049040.1 | *Rattus hoogerwerfi* |
| Rattus | NC_046686.1 | *Rattus andamanensis* |
| Rattus | NC_040919.1 | *Rattus nitidus* |
| Rattus | NC_035621.1 | *Rattus baluensis* |
| Rattus | NC_001665.2 | *Rattus norvegicus* |
| Rattus | NC_029888.1 | *Rattus tiomanicus* |
| Rattus | NC_023347.1 | *Rattus niobe* |
| Rattus | NC_014855.1 | *Rattus leucopus* |
| Rattus | NC_014871.1 | *Rattus sordidus* |
| Rattus | NC_014867.1 | *Rattus fuscipes* |
| Rattus | NC_014864.1 | *Rattus villosissimus* |
| Rattus | NC_014861.1 | *Rattus tunneyi* |
| Rattus | NC_014858.1 | *Rattus lutreolus* |
| Rattus | AC_000022.2 | *Rattus norvegicus Wistar* |
| Rattus | NC_012461.1 | *Rattus praetor* |
| Rattus | NC_012389.1 | *Rattus exulans* |
| Rattus | NC_011638.1 | *Rattus tanezumi* |
| Rattus | NC_049041.1 | *Rattus sp.* |

| | | |
|---|---|---|
| Reduvioidea | NC_050325.1 | *Triatoma huehuetenanguensis* |
| Reduvioidea | NC_050327.1 | *Triatoma mazzottii* |
| Reduvioidea | NC_050326.1 | *Triatoma lecticularia* |
| Reduvioidea | NC_042881.1 | *Triatoma migrans* |
| Reduvioidea | NC_035547.1 | *Triatoma infestans* |
| Reduvioidea | NC_002609.1 | *Triatoma dimidiata* |
| Reduvioidea | NC_050329.1 | *Triatoma sanguisuga* |
| Reduvioidea | NC_050324.1 | *Triatoma mexicana* |
| Reduvioidea | NC_060486.1 | *Neocentrocnemis stali* |
| Reduvioidea | NC_024745.1 | *Brontostoma colossus* |
| Reduvioidea | NC_026672.1 | *Peirates lepturoides* |
| Reduvioidea | NC_026671.1 | *Peirates turpis* |
| Reduvioidea | NC_026670.1 | *Peirates atromaculatus* |
| Reduvioidea | NC_026669.1 | *Peirates fulvescens* |
| Reduvioidea | NC_024264.1 | *Peirates arcuatus* |
| Reduvioidea | NC_020143.1 | *Sirthenea flavipes* |
| Reduvioidea | NC_060487.1 | *Physoderes impexa* |
| Reduvioidea | NC_037369.1 | *Acanthaspis ruficeps* |
| Reduvioidea | NC_037735.1 | *Acanthaspis cincticrus* |
| Reduvioidea | NC_037736.1 | *Inara alboguttata* |
| Reduvioidea | NC_037746.1 | *Reduvius gregoryi* |
| Reduvioidea | NC_035756.1 | *Reduvius tenebrosus* |
| Reduvioidea | NC_037737.1 | *Tapeinus singularis* |

| | | |
|---|---|---|
| Reduvioidea | NC_012823.1 | *Valentia hoffmanni* |
| Reduvioidea | NC_037745.1 | *Canthesancus helluo* |
| Reduvioidea | NC_022816.1 | *Oncocephalus breviscutum* |
| Reduvioidea | NC_042682.1 | *Panstrongylus rufotuberculatus* |
| Reduvioidea | NC_043846.1 | *Rhodnius pictipes* |
| Reduvioidea | NC_050328.1 | *Rhodnius prolixus* |
| Reduvioidea | NC_015842.1 | *Agriosphodrus dohrni* |
| Reduvioidea | NC_037743.1 | *Rhynocoris incertis* |
| Reduvioidea | NC_037744.1 | *Scipinia horrida* |
| Reduvioidea | NC_056990.1 | *Sclomina erinacea* |
| Reduvioidea | NC_053357.1 | *Sycanus croceovittatus* |
| Reduvioidea | NC_037738.1 | *Velinus nodipes* |
| Reticulitermes | NC_062664.1 | *Reticulitermes parvus* |
| Reticulitermes | NC_062663.1 | *Reticulitermes luofunicus* |
| Reticulitermes | NC_062662.1 | *Reticulitermes dabieshanensis* |
| Reticulitermes | NC_062660.1 | *Reticulitermes affinis* |
| Reticulitermes | NC_053728.1 | *Reticulitermes ovatilabrum* |
| Reticulitermes | NC_045240.1 | *Reticulitermes lucifugus* |
| Reticulitermes | NC_045231.1 | *Reticulitermes tibialis* |
| Reticulitermes | NC_042419.1 | *Reticulitermes leptomandibularis* |
| Reticulitermes | NC_031162.1 | *Reticulitermes flaviceps* |
| Reticulitermes | NC_030262.1 | *Reticulitermes labralis* |
| Reticulitermes | NC_030035.1 | *Reticulitermes grassei* |

| | | |
|---|---|---|
| Reticulitermes | NC_026695.1 | *Reticulitermes aculabialis* |
| Reticulitermes | NC_025567.1 | *Reticulitermes chinensis* |
| Reticulitermes | NC_009499.1 | *Reticulitermes santonensis* |
| Reticulitermes | NC_009501.1 | *Reticulitermes hageni* |
| Reticulitermes | NC_009500.1 | *Reticulitermes virginicus* |
| Reticulitermes | NC_009498.1 | *Reticulitermes flavipes* |
| Reticulitermes | NC_062661.1 | *Reticulitermes citrinus* |
| Reticulitermes | NC_030036.1 | *Reticulitermes nelsonae* |
| Rhacophoridae | NC_061400.1 | *Gracixalus yunnanensis* |
| Rhacophoridae | NC_062354.1 | *Polypedates impresus* |
| Rhacophoridae | NC_062356.1 | *Polypedates leucomystax* |
| Rhacophoridae | NC_062355.1 | *Polypedates mutus* |
| Rhacophoridae | NC_043955.1 | *Polypedates megacephalus* |
| Rhacophoridae | NC_042797.1 | *Polypedates braueri* |
| Rhacophoridae | NC_062878.1 | *Rhacophorus chenfui* |
| Rhacophoridae | NC_046387.1 | *Rhacophorus omeimontis* |
| Rhacophoridae | NC_027452.1 | *Rhacophorus dennysi* |
| Rhacophoridae | NC_007178.1 | *Rhacophorus schlegelii* |
| Rhinolophus | NC_061981.1 | *Rhinolophus siamensis* |
| Rhinolophus | NC_061980.1 | *Rhinolophus paradoxolophus* |
| Rhinolophus | NC_061979.1 | *Rhinolophus marshalli* |
| Rhinolophus | NC_061978.1 | *Rhinolophus huananus* |
| Rhinolophus | NC_061262.1 | *Rhinolophus philippinensis* |

| | | |
|---|---|---|
| Rhinolophus | NC_053269.1 | *Rhinolophus affinis* |
| Rhinolophus | NC_046021.1 | *Rhinolophus pusillus* |
| Rhinolophus | NC_036419.1 | *Rhinolophus yunnanensis* |
| Rhinolophus | NC_034306.1 | *Rhinolophus thomasi* |
| Rhinolophus | NC_026460.1 | *Rhinolophus macrotis* |
| Rhinolophus | NC_020326.1 | *Rhinolophus ferrumequinum quelpartis* |
| Rhinolophus | NC_018539.1 | *Rhinolophus luctus* |
| Rhinolophus | NC_005433.1 | *Rhinolophus monoceros* |
| Rhinolophus | NC_005434.1 | *Rhinolophus pumilus* |
| Rhinolophus | NC_011304.1 | *Rhinolophus formosae* |
| Rodentia | NC_033912.1 | *Castor canadensis* |
| Rodentia | NC_050264.1 | *Muscardinus avellanarius* |
| Rodentia | NC_059783.1 | *Eliomys quercinus* |
| Rodentia | NC_023780.1 | *Ratufa bicolor* |
| Ruminantia | NC_020714.1 | *Hyemoschus aquaticus* |
| Ruminantia | NC_037993.1 | *Moschiola indica* |
| Ruminantia | NC_035821.1 | *Tragulus napu* |
| Ruminantia | NC_020753.1 | *Tragulus kanchil* |
| Salamandridae | NC_017870.1 | *Echinotriton andersoni* |
| Salamandridae | NC_028278.1 | *Notophthalmus perstriatus* |
| Salamandridae | NC_054199.1 | *Pachytriton granulosus* |
| Salamandridae | NC_053711.1 | *Pachytriton brevipes* |
| Salamandridae | NC_029345.1 | *Pachytriton feii* |

| | | |
|---|---|---|
| Salamandridae | NC_062887.1 | *Paramesotriton fuzhongensis* |
| Salamandridae | NC_053903.1 | *Paramesotriton aurantius* |
| Salamandridae | NC_037713.1 | *Paramesotriton deloustali* |
| Salamandridae | NC_035008.1 | *Paramesotriton chinensis* |
| Salamandridae | NC_006407.1 | *Paramesotriton hongkongensis* |
| Salamandridae | NC_015796.1 | *Triturus pygmaeus* |
| Salamandridae | NC_015795.1 | *Triturus marmoratus* |
| Salamandridae | NC_015794.1 | *Triturus macedonicus* |
| Salamandridae | NC_015792.1 | *Triturus karelinii* |
| Salamandridae | NC_015791.1 | *Triturus dobrogicus* |
| Salamandridae | NC_015790.1 | *Triturus cristatus* |
| Salamandridae | NC_015788.1 | *Triturus carnifex* |
| Salamandridae | NC_029231.1 | *Tylototriton kweichowensis* |
| Salamandridae | NC_027507.1 | *Tylototriton wenxianensis* |
| Salamandridae | NC_027505.1 | *Tylototriton shanjing* |
| Salamandridae | NC_027421.1 | *Tylototriton taliangensis* |
| Salamandridae | NC_017871.1 | *Tylototriton verrucosus* |
| Sarcophaga | NC_060327.1 | *Sarcophaga gracilior* |
| Sarcophaga | NC_057591.1 | *Sarcophaga tsinanensis* |
| Sarcophaga | NC_053729.1 | *Sarcophaga pauciseta* |
| Sarcophaga | NC_053686.1 | *Sarcophaga aegyptica* |
| Sarcophaga | NC_053685.1 | *Sarcophaga shnitnikovi* |
| Sarcophaga | NC_053683.1 | *Sarcophaga pingi* |

| | | |
|---|---|---|
| Sarcophaga | NC_053682.1 | *Sarcophaga minor* |
| Sarcophaga | NC_053681.1 | *Sarcophaga schuetzei* |
| Sarcophaga | NC_053680.1 | *Sarcophaga cetu* |
| Sarcophaga | NC_053678.1 | *Sarcophaga jacobsoni* |
| Sarcophaga | NC_053675.1 | *Sarcophaga plotnikovi* |
| Sarcophaga | NC_053674.1 | *Sarcophaga diminuta* |
| Sarcophaga | NC_053673.1 | *Sarcophaga macroauriculata* |
| Sarcophaga | NC_053671.1 | *Sarcophaga anchoriformis* |
| Sarcophaga | NC_053669.1 | *Sarcophaga kentejana* |
| Sarcophaga | NC_053667.1 | *Sarcophaga graciliforceps* |
| Sarcophaga | NC_053665.1 | *Sarcophaga carnaria* |
| Sarcophaga | NC_053664.1 | *Sarcophaga depressifrons* |
| Sarcophaga | NC_053666.1 | *Sarcophaga josephi* |
| Sarcophaga | NC_051537.1 | *Sarcophaga kanoi* |
| Sarcophaga | NC_051536.1 | *Sarcophaga scopariiformis* |
| Sarcophaga | NC_047405.1 | *Sarcophaga tuberosa* |
| Sarcophaga | NC_047404.1 | *Sarcophaga brevicornis* |
| Sarcophaga | NC_042759.1 | *Sarcophaga princeps* |
| Sarcophaga | NC_041069.1 | *Sarcophaga ruficornis* |
| Sarcophaga | NC_039827.1 | *Sarcophaga antilope* |
| Sarcophaga | NC_039826.1 | *Sarcophaga dux* |
| Sarcophaga | NC_017605.1 | *Sarcophaga impatiens* |
| Sarcophaga | NC_036428.1 | *Sarcophaga formosensis* |

| | | |
|---|---|---|
| Sarcophaga | NC_036107.1 | *Sarcophaga misera* |
| Sarcophaga | NC_026112.1 | *Sarcophaga melanura* |
| Sarcophaga | NC_025944.1 | *Sarcophaga africa* |
| Sarcophaga | NC_028413.1 | *Sarcophaga albiceps* |
| Sarcophaga | NC_026667.1 | *Sarcophaga crassipalpis* |
| Sarcophaga | NC_025574.1 | *Sarcophaga portschinskyi* |
| Sarcophaga | NC_025573.1 | *Sarcophaga similis* |
| Sarcophaga | NC_023532.1 | *Sarcophaga peregrina* |
| Saturniidae | NC_061295.1 | *Bunaea alcinoe* |
| Saturniidae | NC_061324.1 | *Eochroa trimenii* |
| Saturniidae | NC_061326.1 | *Gonimbrasia tyrrhea* |
| Saturniidae | NC_046032.1 | *Gonimbrasia belina* |
| Saturniidae | NC_061331.1 | *Gonimbrasia cytherea* |
| Saturniidae | NC_046033.1 | *Gynanisa maja* |
| Saturniidae | NC_061327.1 | *Heniocha dyops* |
| Saturniidae | NC_061296.1 | *Heniocha apollonia* |
| Saturniidae | NC_061297.1 | *Holocerina smilax* |
| Saturniidae | NC_061328.1 | *Ludia delegorguei* |
| Saturniidae | NC_061330.1 | *Nudaurelia wahlbergi* |
| Saturniidae | NC_061329.1 | *Vegetia grimmia* |
| Saturniidae | NC_061298.1 | *Vegetia ducalis* |
| Satyrini | NC_046491.1 | *Coenonympha amaryllis* |
| Satyrini | NC_024551.1 | *Triphysa phryne* |

| | | |
|---|---|---|
| Satyrini | NC_024411.1 | *Neope pulaha* |
| Satyrini | NC_026838.1 | *Ninguta schrenckii* |
| Satyrini | NC_065030.1 | *Mycalesis francisca* |
| Satyrini | NC_039986.1 | *Lasiommata deidamia* |
| Satyrini | NC_063460.1 | *Lopinga achine* |
| Satyrini | NC_028505.1 | *Davidina armandi* |
| Satyrini | NC_014587.1 | *Hipparchia autonoe* |
| Satyrini | NC_046889.1 | *Oeneis urda* |
| Scandentia | NC_065160.1 | *Tupaia nicobarica* |
| Scandentia | NC_054191.1 | *Tupaia montana* |
| Scandentia | NC_002521.1 | *Tupaia belangeri* |
| Scandentia | NC_050992.1 | *Tupaia tana* |
| Scandentia | NC_050993.1 | *Tupaia minor* |
| Scarabaeiformia | NC_062860.1 | *Clinterocera nigra* |
| Scarabaeiformia | NC_065313.1 | *Coenochilus striatus* |
| Scarabaeiformia | NC_063846.1 | *Gametis jucunda* |
| Scarabaeiformia | NC_063847.1 | *Glycyphana fulvistemma* |
| Scarabaeiformia | NC_030778.1 | *Osmoderma opicum* |
| Scarabaeiformia | NC_063849.1 | *Trichius succinctus* |
| Scarabaeiformia | NC_059757.1 | *Eophileurus chinensis* |
| Scarabaeiformia | NC_066495.1 | *Eupatorus hardwickei* |
| Scarabaeiformia | NC_066494.1 | *Eupatorus sukkiti* |
| Scarabaeiformia | NC_065036.1 | *Eupatorus gracilicornis* |

| | | |
|---|---|---|
| Scarabaeiformia | NC_059756.1 | *Oryctes rhinoceros* |
| Scarabaeiformia | NC_062856.1 | *Trichogomphus mongol* |
| Scarabaeiformia | NC_065312.1 | *Apogonia cf. basalis* |
| Scarabaeiformia | NC_065311.1 | *Apogonia splendida* |
| Scarabaeiformia | NC_046890.1 | *Cheirotonus gestroi* |
| Scarabaeiformia | NC_023246.1 | *Cheirotonus jansoni* |
| Scarabaeiformia | NC_054285.1 | *Polyphylla gracilicornis* |
| Scarabaeiformia | NC_013252.1 | *Rhopaea magnicornis* |
| Scarabaeiformia | NC_065314.1 | *Sophrops subrugatus* |
| Scarabaeiformia | NC_060602.1 | *Ophrygonius sp.* |
| Scarabaeiformia | NC_065310.1 | *Anomala russiventris* |
| Scarabaeiformia | NC_056126.1 | *Popillia mutans* |
| Scarabaeiformia | NC_038115.1 | *Popillia japonica* |
| Scarabaeiformia | NC_036157.1 | *Sinodendron yunnanense* |
| Sciaroidea | NC_061662.1 | *Bradysia odoriphaga* |
| Sciaroidea | NC_046768.1 | *Dolichosciara megumiae* |
| Sciaroidea | NC_053636.1 | *Pnyxia scabiei* |
| Sciaroidea | NC_046767.1 | *Sciara ruficauda* |
| Sciaroidea | NC_046769.1 | *Trichosia lengersdorfi* |
| Sciaroidea | NC_016204.1 | *Arachnocampa flava* |
| Sciaroidea | NC_050318.1 | *Acnemia nitidicollis* |
| Sciaroidea | NC_060624.1 | *Allodia protenta* |
| Sciaroidea | NC_060623.1 | *Allodia pyxidiiformis* |

| | | |
|---|---|---|
| Sciaroidea | NC_060622.1 | *Allodia zaitzevi* |
| Scincidae | NC_058309.1 | *Asymblepharus himalayanus* |
| Scincidae | NC_041147.1 | *Isopachys gyldenstolpei* |
| Scincidae | NC_054206.1 | *Scincella reevesii* |
| Scincidae | NC_048521.1 | *Scincella modesta* |
| Scincidae | NC_030779.1 | *Scincella huanrenensis* |
| Scincidae | NC_030776.1 | *Scincella vandenburghi* |
| Scincidae | NC_045408.1 | *Sphenomorphus indicus* |
| Scincidae | NC_041124.1 | *Sphenomorphus incognitus* |
| Scincidae | NC_066473.1 | *Tropidophorus hainanus* |
| Scincidae | NC_050664.1 | *Tropidophorus hangnam* |
| Scincidae | NC_063644.1 | *Eutropis multifasciata* |
| Scincidae | NC_057221.1 | *Ateuchosaurus chinensis* |
| Scincidae | NC_045232.1 | *Plestiodon tunganus* |
| Scincidae | NC_029352.1 | *Plestiodon chinensis* |
| Scincidae | NC_024576.1 | *Plestiodon elegans* |
| Scincidae | NC_000888.1 | *Plestiodon egregius* |
| Sciurinae | NC_050025.1 | *Guerlinguetus aestuans* |
| Sciurinae | NC_050010.1 | *Guerlinguetus brasiliensis* |
| Sciurinae | NC_050029.1 | *Hadrosciurus spadiceus* |
| Sciurinae | NC_050028.1 | *Hadrosciurus pyrrhinus* |
| Sciurinae | NC_050027.1 | *Hadrosciurus igniventris* |
| Sciurinae | NC_050031.1 | *Microsciurus sabanillae* |

| | | |
|---|---|---|
| Sciurinae | NC_050030.1 | *Microsciurus flaviventer* |
| Sciurinae | NC_050022.1 | *Microsciurus similis* |
| Sciurinae | NC_050020.1 | *Microsciurus mimulus* |
| Sciurinae | NC_050021.1 | *Microsciurus otinus* |
| Sciurinae | NC_065807.1 | *Sciurus anomalus* |
| Sciurinae | NC_050033.1 | *Sciurus lis* |
| Sciurinae | NC_002369.1 | *Sciurus vulgaris* |
| Sciurinae | NC_050024.1 | *Simosciurus stramineus* |
| Sciurinae | NC_050023.1 | *Simosciurus nebouxii* |
| Sciurinae | NC_050026.1 | *Glaucomys volans* |
| Sciurinae | NC_031847.1 | *Hylopetes alboniger* |
| Sciurinae | NC_026443.1 | *Hylopetes phayrei* |
| Sciurinae | NC_023922.1 | *Petaurista alborufus* |
| Sciurinae | NC_023089.1 | *Petaurista hainana* |
| Sciurinae | NC_033902.1 | *Petaurista yunanensis* |
| Sciurinae | NC_019612.1 | *Pteromys volans* |
| Setisura | NC_063607.1 | *Electrogena lateralis* |
| Setisura | NC_065660.1 | *Heptagenia ngi* |
| Setisura | NC_005924.1 | *Heptathela hangzhouensis* |
| Setisura | NC_065661.1 | *Notacanthurus lamellosus* |
| Setisura | NC_065662.1 | *Paegniodes cupulatus* |
| Setisura | NC_011359.1 | *Parafronurus youi* |
| Setisura | NC_042163.1 | *Isonychia kiangsinensis* |

| | | |
|---|---|---|
| Silphidae | NC_059900.1 | *Nicrophorus nepalensis* |
| Silphidae | NC_045874.1 | *Diamesus osculans* |
| Silphidae | NC_056196.1 | *Necrodes littoralis* |
| Silphidae | NC_018352.1 | *Necrophila americana* |
| Silphidae | NC_061362.1 | *Oiceoptoma thoracicum* |
| Sinopodisma | NC_056238.1 | *Sinopodisma qinlingensis* |
| Sinopodisma | NC_052716.1 | *Sinopodisma rostellocerca* |
| Sinopodisma | NC_051867.1 | *Sinopodisma pieli* |
| Sinopodisma | NC_046562.1 | *Sinopodisma lofaoshana* |
| Sinopodisma | NC_046547.1 | *Sinopodisma wudangshanensis* |
| Sinopodisma | NC_046546.1 | *Sinopodisma funiushana* |
| Sinopodisma | NC_033906.1 | *Sinopodisma wulingshanensis* |
| Sinopodisma | NC_033905.1 | *Sinopodisma houshana* |
| Sinopodisma | NC_032303.1 | *Sinopodisma tsinlingensis* |
| Smerinthinae | NC_046728.1 | *Adhemarius dariensis* |
| Smerinthinae | NC_046713.1 | *Adhemarius dentoni* |
| Smerinthinae | NC_062106.1 | *Ambulyx liturata* |
| Smerinthinae | NC_046715.1 | *Ambulyx substrigilis* |
| Smerinthinae | NC_046714.1 | *Ambulyx dohertyi* |
| Smerinthinae | NC_046717.1 | *Amplypterus panopus* |
| Smerinthinae | NC_046716.1 | *Amplypterus mansoni* |
| Smerinthinae | NC_046718.1 | *Barbourion lemaii* |
| Smerinthinae | NC_046719.1 | *Batocnema coquerelii* |

| | | |
|---|---|---|
| Smerinthinae | NC_046722.1 | *Orecta lycidas* |
| Smerinthinae | NC_046726.1 | *Protambulyx strigilis* |
| Smerinthinae | NC_046725.1 | *Protambulyx ockendeni* |
| Smerinthinae | NC_046724.1 | *Protambulyx eurycles* |
| Smerinthinae | NC_046723.1 | *Protambulyx astygonus* |
| Smerinthinae | NC_046727.1 | *Trogolegnum pseudambulyx* |
| Smerinthinae | NC_046720.1 | *Clanis bilineata* |
| Smerinthinae | NC_046721.1 | *Leucophlebia lineata* |
| Smerinthinae | NC_066088.1 | *Marumba saishiuana* |
| Smerinthinae | NC_039166.1 | *Parum colligata* |
| Sorex | NC_064993.1 | *Sorex thibetanus* |
| Sorex | NC_044107.1 | *Sorex daphaenodon* |
| Sorex | NC_042196.1 | *Sorex minutissimus* |
| Sorex | NC_037174.1 | *Sorex sinalis* |
| Sorex | NC_034808.1 | *Sorex roboratus* |
| Sorex | NC_027963.1 | *Sorex araneus* |
| Sorex | NC_025327.1 | *Sorex tundrensis* |
| Sorex | NC_025278.1 | *Sorex cylindricauda* |
| Sorex | NC_005435.1 | *Sorex unguiculatus* |
| Sorex | NC_037859.1 | *Sorex gracillimus* |
| Sphenisciformes | NC_045377.1 | *Aptenodytes patagonicus* |
| Sphenisciformes | NC_027938.1 | *Aptenodytes forsteri* |
| Sphenisciformes | NC_054275.1 | *Eudyptes chrysolophus* |

| | | |
|---|---|---|
| Sphenisciformes | NC_008138.1 | *Eudyptes chrysocome* |
| Sphenisciformes | NC_004538.1 | *Eudyptula minor* |
| Sphenisciformes | NC_037702.1 | *Pygoscelis papua* |
| Sphenisciformes | NC_021137.1 | *Pygoscelis adeliae* |
| Sphenisciformes | NC_021474.1 | *Pygoscelis antarcticus* |
| Sphenisciformes | NC_036337.1 | *Spheniscus humboldti* |
| Sphenisciformes | NC_036297.1 | *Spheniscus mendiculus* |
| Sphenisciformes | NC_036264.1 | *Spheniscus magellanicus* |
| Sphenisciformes | NC_022817.1 | *Spheniscus demersus* |
| Staphyliniformia | NC_036270.1 | *Hydrochus carinatus* |
| Staphyliniformia | NC_018349.1 | *Tropisternus sp.* |
| Staphyliniformia | NC_028612.1 | *Sphaeridium bipustulatum* |
| Staphyliniformia | NC_063613.1 | *Sphaeridium lunatum* |
| Stratiomyidae | NC_035232.1 | *Hermetia illucens* |
| Stratiomyidae | NC_054304.1 | *Nasimyia megacephala* |
| Stratiomyidae | NC_053880.1 | *Parastratiosphecomyia szechuanensis* |
| Stratiomyidae | NC_054251.1 | *Tinda javana* |
| Stratiomyidae | NC_060842.1 | *Ptecticus tenebrifer* |
| Strepsirrhini | NC_021940.1 | *Avahi laniger* |
| Strepsirrhini | NC_065434.1 | *Cheirogaleus medius* |
| Strepsirrhini | NC_035655.1 | *Cheirogaleus major* |
| Strepsirrhini | NC_035605.1 | *Cheirogaleus sibreei* |
| Strepsirrhini | NC_035604.1 | *Cheirogaleus crossleyi* |

| | | |
|---|---|---|
| Strepsirrhini | NC_021948.1 | *Eulemur rufus* |
| Strepsirrhini | NC_012771.1 | *Eulemur macaco macaco* |
| Strepsirrhini | NC_012769.1 | *Eulemur fulvus mayottensis* |
| Strepsirrhini | NC_026098.1 | *Eulemur rubriventer* |
| Strepsirrhini | NC_010300.1 | *Eulemur mongoz* |
| Strepsirrhini | NC_021950.1 | *Hapalemur griseus* |
| Strepsirrhini | NC_035656.1 | *Mirza coquereli* |
| Strepsirrhini | NC_035606.1 | *Mirza zaza* |
| Strepsirrhini | NC_035609.1 | *Phaner parienti* |
| Strepsirrhini | NC_035608.1 | *Phaner pallescens* |
| Strepsirrhini | NC_035607.1 | *Phaner electromontis* |
| Strepsirrhini | NC_021959.1 | *Prolemur simus* |
| Strepsirrhini | NC_026084.1 | *Propithecus diadema* |
| Strepsirrhini | NC_011053.1 | *Propithecus coquereli* |
| Strepsirrhini | NC_028210.1 | *Propithecus verreauxi* |
| Strepsirrhini | NC_027740.1 | *Propithecus tattersalli* |
| Strepsirrhini | NC_026086.1 | *Propithecus edwardsi* |
| Strepsirrhini | NC_065304.1 | *Propithecus candidus* |
| Strepsirrhini | NC_012773.1 | *Varecia variegata variegata* |
| Strepsirrhini | NC_026085.1 | *Varecia variegata* |
| Strepsirrhini | NC_010299.1 | *Daubentonia madagascariensis* |
| Strepsirrhini | NC_026088.1 | *Megaladapis edwardsi* |
| Strepsirrhini | NC_026090.1 | *Palaeopropithecus ingens* |

| | | |
|---|---|---|
| Sturnidae | NC_015613.1 | *Acridotheres cristatellus* |
| Sturnidae | NC_053077.1 | *Buphagus erythrorhynchus* |
| Sturnidae | NC_015898.1 | *Gracula religiosa* |
| Sturnidae | NC_053050.1 | *Leucopsar rothschildi* |
| Sturnidae | NC_029360.1 | *Sturnus vulgaris* |
| Sturnidae | NC_014455.1 | *Sturnus sericeus* |
| Sturnidae | NC_020423.1 | *Sturnus nigricollis* |
| Sturnidae | NC_015237.1 | *Sturnus cineraceus* |
| Sturnidae | NC_015195.1 | *Sturnus tristis* |
| Suina | NC_008830.1 | *Phacochoerus africanus* |
| Suina | NC_043879.1 | *Porcula salvania* |
| Suina | NC_020737.1 | *Potamochoerus porcus* |
| Suina | NC_000845.1 | *Sus scrofa* |
| Suina | NC_039090.1 | *Sus scrofa cristatus* |
| Suina | NC_026992.1 | *Sus barbatus* |
| Suina | NC_024860.1 | *Sus celebensis* |
| Suina | NC_023541.1 | *Sus cebifrons* |
| Suina | NC_012095.1 | *Sus scrofa domesticus* |
| Suina | NC_014692.1 | *Sus scrofa taiwanensis* |
| Sylvioidea | NC_053069.1 | *Erpornis zantholeuca* |
| Sylvioidea | NC_050259.1 | *Erythrogenys gravivox dedekensi* |
| Sylvioidea | NC_040006.1 | *Erythrogenys gravivox* |
| Sylvioidea | NC_045401.1 | *Fulvetta ruficapilla* |

| | | |
|---|---|---|
| Sylvioidea | NC_039396.1 | *Ixos mcclellandii* |
| Sylvioidea | NC_053072.1 | *Mystacornis crossleyi* |
| Sylvioidea | NC_035626.1 | *Napothera epilepidota* |
| Sylvioidea | NC_053062.1 | *Nicator chloris* |
| Sylvioidea | NC_052842.1 | *Parophasma galinieri* |
| Sylvioidea | NC_029769.1 | *Pomatorhinus ruficollis* |
| Sylvioidea | NC_051464.1 | *Pomatostomus ruficeps* |
| Sylvioidea | NC_053108.1 | *Pteruthius melanotis* |
| Sylvioidea | NC_053063.1 | *Pycnonotus jocosus* |
| Sylvioidea | NC_031830.1 | *Pycnonotus xanthorrhous* |
| Sylvioidea | NC_024730.1 | *Pycnonotus melanicterus* |
| Sylvioidea | NC_013483.2 | *Pycnonotus taivanus* |
| Sylvioidea | NC_029321.1 | *Spizixos semitorques* |
| Sylvioidea | NC_020604.1 | *Tachycineta meyeni* |
| Sylvioidea | NC_020603.1 | *Tachycineta leucorrhoa* |
| Sylvioidea | NC_020602.1 | *Tachycineta albiventer* |
| Sylvioidea | NC_020601.1 | *Tachycineta albilinea* |
| Sylvioidea | NC_020600.1 | *Tachycineta stolzmanni* |
| Sylvioidea | NC_020599.1 | *Tachycineta cyaneoviridis* |
| Sylvioidea | NC_020598.1 | *Tachycineta euchrysea* |
| Sylvioidea | NC_020597.1 | *Tachycineta thalassina* |
| Sylvioidea | NC_020596.1 | *Tachycineta bicolor* |
| Sylvioidea | NC_040991.1 | *Yuhina nigrimenta* |

| | | |
|---|---|---|
| Sylvioidea | NC_029462.1 | *Yuhina diademata* |
| Sylvioidea | NC_024268.1 | *Aegithalos glaucogularis* |
| Sylvioidea | NC_024267.1 | *Aegithalos bonvaloti* |
| Sylvioidea | NC_057960.1 | *Alauda gulgula* |
| Sylvioidea | NC_020425.1 | *Alauda arvensis* |
| Sylvioidea | NC_046419.1 | *Alaudala cheleensis* |
| Sylvioidea | NC_048469.1 | *Calandrella cinerea* |
| Sylvioidea | NC_048470.1 | *Eremophila alpestris* |
| Sylvioidea | NC_025616.1 | *Eremopsaltria mongolica* |
| Sylvioidea | NC_036760.1 | *Melanocorypha mongolica* |
| Sylvioidea | NC_024107.1 | *Cecropis daurica* |
| Sylvioidea | NC_050298.1 | *Delichon urbicum* |
| Sylvioidea | NC_020605.1 | *Progne chalybea* |
| Sylvioidea | NC_020427.1 | *Leiothrix lutea* |
| Sylvioidea | NC_015114.1 | *Leiothrix argentauris* |
| Sylvioidea | NC_030588.1 | *Minla ignotincta* |
| Sylvioidea | NC_041141.1 | *Trochalopteron milnei* |
| Sylvioidea | NC_059853.1 | *Locustella pleskei* |
| Sylvioidea | NC_060852.1 | *Phylloscopus collybita* |
| Sylvioidea | NC_060851.1 | *Phylloscopus trochilus* |
| Sylvioidea | NC_046416.1 | *Phylloscopus fuscatus* |
| Sylvioidea | NC_024726.1 | *Phylloscopus inornatus* |
| Sylvioidea | NC_064505.1 | *Phylloscopus burkii* |

| | | |
|---|---|---|
| Sylvioidea | NC_053060.1 | *Rhadina sibilatrix* |
| Sylvioidea | NC_053103.1 | *Horornis vulcanius* |
| Sylvioidea | NC_053106.1 | *Panurus biarmicus* |
| Sylvioidea | NC_059907.1 | *Sinosuthora conspicillata* |
| Sylvioidea | NC_053073.1 | *Hypocryptadius cinnamomeus* |
| Sylvioidea | NC_029146.1 | *Zosterops lateralis* |
| Sylvioidea | NC_027942.1 | *Zosterops erythropleurus* |
| Sylvioidea | NC_032059.1 | *Zosterops poliogastrus* |
| Sylvioidea | NC_059932.1 | *Zosterops japonicus* |
| Sylvioidea | NC_032058.1 | *Zosterops abyssinicus* |
| Sylvioidea | NC_046418.1 | *Acrocephalus orientalis* |
| Sylvioidea | NC_010227.1 | *Acrocephalus scirpaceus* |
| Syrphidae | NC_052908.1 | *Eristalinus quinquestriatus* |
| Syrphidae | NC_042910.1 | *Eristalinus tabanoides* |
| Syrphidae | NC_042908.1 | *Eristalinus barclayi* |
| Syrphidae | NC_042911.1 | *Eristalinus aeneus* |
| Syrphidae | NC_042909.1 | *Eristalinus vicarians* |
| Syrphidae | NC_050932.1 | *Eristalis cerealis* |
| Syrphidae | NC_041143.1 | *Eristalis tenax* |
| Syrphidae | NC_049124.1 | *Phytomia zonata* |
| Syrphidae | NC_050314.1 | *Orthonevra geniculata* |
| Syrphidae | NC_064403.1 | *Sphegina orientalis* |
| Syrphidae | NC_062964.1 | *Sphegina sp.* |

| | | |
|---|---|---|
| Syrphidae | NC_052907.1 | *Ferdinandea cuprea* |
| Syrphidae | NC_063606.1 | *Tropidia scita DM968* |
| Syrphinae | NC_061032.1 | *Melanostoma mellinum* |
| Syrphinae | NC_051975.1 | *Melanostoma orientale* |
| Syrphinae | NC_050968.1 | *Melanostoma scalare* |
| Syrphinae | NC_056282.1 | *Platycheirus albimanus* |
| Syrphinae | NC_036481.1 | *Episyrphus balteatus* |
| Syrphinae | NC_036482.1 | *Eupeodes corollae* |
| Syrphinae | NC_008754.1 | *Simosyrphus grandicornis* |
| Syrphinae | NC_056283.1 | *Syrphus torvus* |
| Syrphinae | NC_054190.1 | *Syrphus ribesii* |
| Syrphinae | NC_050969.1 | *Syrphus vitripennis* |
| Tabaninae | NC_008756.1 | *Cydistomyia duplonotata* |
| Tabaninae | NC_059934.1 | *Haematopota vexativa Henan* |
| Tabaninae | NC_030000.1 | *Atylotus miser* |
| Tabaninae | NC_062705.1 | *Tabanus chrysurus* |
| Tarsiiformes | NC_012774.1 | *Carlito syrichta* |
| Tarsiiformes | NC_024053.1 | *Tarsius wallacei* |
| Tarsiiformes | NC_024052.1 | *Tarsius dentatus* |
| Tarsiiformes | NC_024051.1 | *Tarsius lariang* |
| Tarsiiformes | NC_002811.1 | *Tarsius bancanus* |
| Tenthredinidae | NC_057612.1 | *Allantus togatus* |
| Tenthredinidae | NC_024664.1 | *Allantus luctifer* |

| | | |
|---|---|---|
| Tenthredinidae | NC_062840.1 | *Cladiucha huangbki* |
| Tenthredinidae | NC_062839.1 | *Cladiucha magnoliae* |
| Tenthredinidae | NC_062838.1 | *Cladiucha punctata* |
| Tenthredinidae | NC_056796.1 | *Megabeleses magnoliae* |
| Tenthredinidae | NC_056795.1 | *Megabeleses liriodendrovorax* |
| Tephritidae | NC_061932.1 | *Dacus vijaysegarani* |
| Tephritidae | NC_053984.1 | *Dacus trimacula* |
| Tephritidae | NC_043843.1 | *Dacus conopsoides* |
| Tephritidae | NC_032690.1 | *Dacus longicornis* |
| Tephritidae | NC_062801.1 | *Zeugodacus caudatus* |
| Tephritidae | NC_062140.1 | *Zeugodacus strigifinis* |
| Tephritidae | NC_061658.1 | *Zeugodacus scutellaris* |
| Tephritidae | NC_016056.1 | *Zeugodacus cucurbitae* |
| Tephritidae | NC_052852.1 | *Zeugodacus cilifer* |
| Tephritidae | NC_049063.1 | *Zeugodacus proprediaphora* |
| Tephritidae | NC_000857.1 | *Ceratitis capitata* |
| Tephritidae | NC_053847.1 | *Ceratitis rosa* |
| Tephritidae | NC_053846.1 | *Ceratitis quilicii* |
| Tephritidae | NC_035497.1 | *Ceratitis fasciventris* |
| Tephritidae | NC_052851.1 | *Felderimyia fuscipennis* |
| Tephritidae | NC_020463.1 | *Procecidochares utilis* |
| Tephritidae | NC_047184.1 | *Tephritis femoralis* |
| Tephritidae | NC_034912.1 | *Anastrepha fraterculus* |

| | | |
|---|---|---|
| Tephritidae | NC_061399.1 | *Rhagoletis cerasi* |
| Tephritidae | NC_053982.1 | *Acidiella diversa* |
| Termitidae | NC_034135.1 | *Acholotermes chirotus* |
| Termitidae | NC_034089.1 | *Aciculitermes aciculatus* |
| Termitidae | NC_034107.1 | *Aciculitermes maymyoensis* |
| Termitidae | NC_034122.1 | *Acidnotermes praus* |
| Termitidae | NC_034025.1 | *Agnathotermes crassinasus* |
| Termitidae | NC_034079.1 | *Allodontermes schultzei* |
| Termitidae | NC_034022.1 | *Alyscotermes kilimandjaricus* |
| Termitidae | NC_034099.1 | *Amalotermes phaeocephalus* |
| Termitidae | NC_034092.1 | *Anhangatermes macarthuri* |
| Termitidae | NC_034123.1 | *Anoplotermes parvus* |
| Termitidae | NC_034120.1 | *Anoplotermes janus* |
| Termitidae | NC_034057.1 | *Araujotermes parvellus* |
| Termitidae | NC_034132.1 | *Astalotermes murcus* |
| Termitidae | NC_034069.1 | *Ateuchotermes retifaciens* |
| Termitidae | NC_034148.1 | *Atlantitermes oculatissimus* |
| Termitidae | NC_034102.1 | *Atlantitermes snyderi* |
| Termitidae | NC_034147.1 | *Bulbitermes singaporiensis* |
| Termitidae | NC_034059.1 | *Bulbitermes laticephalus* |
| Termitidae | NC_034029.1 | *Bulbitermes makhamensis* |
| Termitidae | NC_034141.1 | *Coatitermes kartaboensis* |
| Termitidae | NC_034144.1 | *Compositermes vindai* |

| | | |
|---|---|---|
| Termitidae | NC_034044.1 | *Constrictotermes cyphergaster* |
| Termitidae | NC_034064.1 | *Euhamitermes hamatus* |
| Termitidae | NC_034070.1 | *Havilanditermes proatripennis* |
| Termitidae | NC_034134.1 | *Hirtitermes hirtiventris* |
| Termitidae | NC_036047.1 | *Hospitalitermes medioflavus* |
| Termitidae | NC_034074.1 | *Hospitalitermes hospitalis* |
| Termitidae | NC_034129.1 | *Humutermes krishnai* |
| Termitidae | NC_034037.1 | *Hypotermes makhamensis* |
| Termitidae | NC_034083.1 | *Jugositermes tuberculatus* |
| Termitidae | NC_034047.1 | *Leucopitermes leucops* |
| Termitidae | NC_034094.1 | *Longustitermes manni* |
| Termitidae | NC_034072.1 | *Microtermes obesi* |
| Termitidae | NC_034088.1 | *Occasitermes occasus* |
| Termitidae | NC_034130.1 | *Odontotermes longignathus* |
| Termitidae | NC_034028.1 | *Odontotermes hainanensis* |
| Termitidae | NC_034027.1 | *Odontotermes obesus* |
| Termitidae | NC_034106.1 | *Odontotermes javanicus* |
| Termitidae | NC_034061.1 | *Odontotermes minutus* |
| Termitidae | NC_034035.1 | *Odontotermes mathuri* |
| Termitidae | NC_034087.1 | *Oriensubulitermes inanis* |
| Termitidae | NC_034032.1 | *Patawatermes nigripunctatus* |
| Termitidae | NC_034137.1 | *Patawatermes turricola* |
| Termitidae | NC_034114.1 | *Postsubulitermes parviconstrictus* |

| | | |
|---|---|---|
| Termitidae | NC_034126.1 | *Protermes prorepens* |
| Termitidae | NC_034077.1 | *Pseudacanthotermes militaris* |
| Termitidae | NC_034024.1 | *Pseudacanthotermes spiniger* |
| Termitidae | NC_034111.1 | *Rubeotermes jheringi* |
| Termitidae | NC_034140.1 | *Ruptitermes arboreus* |
| Termitidae | NC_034103.1 | *Sphaerotermes sphaerothorax* |
| Termitidae | NC_034073.1 | *Trichotermes ducis* |
| Termitidae | NC_034098.1 | *Tumulitermes pastinator* |
| Termitidae | NC_034051.1 | *Tumulitermes recalvus* |
| Termitinae | NC_034075.1 | *Amitermes dentatus* |
| Termitinae | NC_034062.1 | *Amitermes meridionalis* |
| Termitinae | NC_034038.1 | *Amitermes capito* |
| Termitinae | NC_034124.1 | *Amitermes obeuntis* |
| Termitinae | NC_034031.1 | *Apilitermes longiceps* |
| Termitinae | NC_034097.1 | *Cavitermes tuberosus* |
| Termitinae | NC_034136.1 | *Cephalotermes rectangularis* |
| Termitinae | NC_034113.1 | *Crenetermes albotarsalis* |
| Termitinae | NC_034041.1 | *Crepititermes verruculosus* |
| Termitinae | NC_034056.1 | *Cubitermes oblectatus* |
| Termitinae | NC_034033.1 | *Cubitermes fulvus* |
| Termitinae | NC_026113.1 | *Cubitermes ugandensis* |
| Termitinae | NC_034109.1 | *Cubitermes sulcifrons* |
| Termitinae | NC_034096.1 | *Cylindrotermes parvignathus* |

| | | |
|---|---|---|
| Termitinae | NC_018129.1 | *Drepanotermes sp.* |
| Termitinae | NC_034149.1 | *Ephelotermes taylori* |
| Termitinae | NC_034019.1 | *Ephelotermes melachoma* |
| Termitinae | NC_034116.1 | *Foraminitermes rhinoceros* |
| Termitinae | NC_034131.1 | *Furculitermes cubitalis* |
| Termitinae | NC_034063.1 | *Furculitermes winifredae* |
| Termitinae | NC_034128.1 | *Furculitermes longilabius* |
| Termitinae | NC_034082.1 | *Furculitermes soyeri* |
| Termitinae | NC_034139.1 | *Globitermes sulphureus* |
| Termitinae | NC_034095.1 | *Globitermes globosus* |
| Termitinae | NC_034118.1 | *Inquilinitermes inquilinus* |
| Termitinae | NC_034058.1 | *Labritermes buttelreepeni* |
| Termitinae | NC_034105.1 | *Lophotermes septentrionalis* |
| Termitinae | NC_018130.1 | *Macrognathotermes errator* |
| Termitinae | NC_034142.1 | *Microcerotermes serrula* |
| Termitinae | NC_034084.1 | *Microcerotermes havilandi* |
| Termitinae | NC_034067.1 | *Microcerotermes fuscotibialis* |
| Termitinae | NC_034065.1 | *Microcerotermes baluchistanicus* |
| Termitinae | NC_034104.1 | *Microcerotermes nervosus* |
| Termitinae | NC_034036.1 | *Microcerotermes crassus* |
| Termitinae | NC_034021.1 | *Microcerotermes newmani* |
| Termitinae | NC_026114.1 | *Microcerotermes parvus* |
| Termitinae | NC_034133.1 | *Microcerotermes progrediens* |

| | | |
|---|---|---|
| Termitinae | NC_034085.1 | *Mirocapritermes connectens* |
| Termitinae | NC_034053.1 | *Neocapritermes angusticeps* |
| Termitinae | NC_026116.1 | *Neocapritermes taracua* |
| Termitinae | NC_034145.1 | *Noditermes cristifrons* |
| Termitinae | NC_034076.1 | *Ophiotermes grandilabius* |
| Termitinae | NC_034068.1 | *Ophiotermes mirandus* |
| Termitinae | NC_034048.1 | *Orientotermes emersoni* |
| Termitinae | NC_034125.1 | *Orthotermes depressifrons* |
| Termitinae | NC_034100.1 | *Orthotermes mansuetus* |
| Termitinae | NC_039398.1 | *Pericapritermes nitobei* |
| Termitinae | NC_034112.1 | *Pericapritermes dolichocephalus* |
| Termitinae | NC_034090.1 | *Planicapritermes planiceps* |
| Termitinae | NC_034071.1 | *Proboscitermes tubuliferus* |
| Termitinae | NC_034119.1 | *Procapritermes martyni* |
| Termitinae | NC_034138.1 | *Procubitermes undulans* |
| Termitinae | NC_034039.1 | *Prohamitermes mirabilis* |
| Termitinae | NC_034081.1 | *Promirotermes pygmaeus* |
| Termitinae | NC_034091.1 | *Spinitermes trispinosus* |
| Termitinae | NC_034121.1 | *Termes comis* |
| Termitinae | NC_034049.1 | *Termes fatalis* |
| Termitinae | NC_034043.1 | *Termes rostratus* |
| Termitinae | NC_026117.1 | *Termes hospes* |
| Termitinae | NC_034143.1 | *Thoracotermes macrothorax* |

| | | |
|---|---|---|
| Termitinae | NC_034052.1 | *Tuberculitermes bycanistes* |
| Testudines | NC_015989.1 | *Chelus fimbriata* |
| Testudines | NC_041289.1 | *Elseya novaeguineae* |
| Testudines | NC_041288.1 | *Elseya dentata* |
| Testudines | NC_041287.1 | *Elseya albagula* |
| Testudines | NC_041283.1 | *Elseya lavarackorum* |
| Testudines | NC_041282.1 | *Elseya irwini* |
| Testudines | NC_026047.1 | *Elseya branderhorsti* |
| Testudines | NC_041290.1 | *Elseya schultzei* |
| Testudines | NC_030217.1 | *Elusor macrurus* |
| Testudines | NC_042476.1 | *Emydura tanybaraga* |
| Testudines | NC_042473.1 | *Emydura victoriae* |
| Testudines | NC_041302.1 | *Emydura macquarii* |
| Testudines | NC_036346.1 | *Mesoclemmys hogei* |
| Testudines | NC_042475.1 | *Myuchelys latisternum* |
| Testudines | NC_042474.1 | *Myuchelys georgesi* |
| Testudines | NC_035748.1 | *Myuchelys bellii* |
| Testudines | NC_026050.1 | *Platemys platycephala* |
| Testudines | NC_035731.1 | *Pseudemydura umbrina* |
| Testudines | NC_042247.1 | *Rheodytes leukops* |
| Testudines | NC_001947.1 | *Pelomedusa subrufa* |
| Testudines | NC_026049.1 | *Pelusios castaneus* |
| Testudinoidea | NC_028438.1 | *Aldabrachelys gigantea* |

| | | |
|---|---|---|
| Testudinoidea | NC_041096.1 | *Geochelone elegans* |
| Testudinoidea | NC_007696.1 | *Indotestudo forstenii* |
| Testudinoidea | NC_007695.1 | *Indotestudo elongata* |
| Testudinoidea | NC_007700.1 | *Malacochersus tornieri* |
| Testudinoidea | NC_007693.1 | *Manouria emys* |
| Testudinoidea | NC_011815.1 | *Manouria impressa* |
| Testudinoidea | NC_007699.1 | *Testudo kleinmanni* |
| Testudinoidea | NC_007697.1 | *Testudo horsfieldii* |
| Testudinoidea | NC_007692.1 | *Testudo graeca* |
| Testudinoidea | NC_002073.3 | *Chrysemys picta* |
| Testudinoidea | NC_023890.1 | *Chrysemys picta bellii* |
| Testudinoidea | NC_031300.1 | *Malaclemys terrapin terrapin* |
| Testudinoidea | NC_063096.1 | *Pseudemys peninsularis* |
| Testudinoidea | NC_011573.1 | *Trachemys scripta* |
| Testudinoidea | NC_062409.1 | *Cuora mccordi* |
| Testudinoidea | NC_022857.1 | *Cuora trifasciata* |
| Testudinoidea | NC_017885.1 | *Cuora bourreti* |
| Testudinoidea | NC_014769.1 | *Cuora amboinensis* |
| Testudinoidea | NC_014401.1 | *Cuora pani* |
| Testudinoidea | NC_014102.1 | *Cuora galbinifrons* |
| Testudinoidea | NC_018793.1 | *Cyclemys dentata* |
| Testudinoidea | NC_026027.1 | *Cyclemys pulchristriata* |
| Testudinoidea | NC_023221.1 | *Cyclemys tcheponensis* |

| | | |
|---|---|---|
| Testudinoidea | NC_010970.1 | *Cyclemys atripons* |
| Testudinoidea | NC_044641.1 | *Geoemyda spengleri* |
| Testudinoidea | NC_032297.1 | *Heosemys grandis* |
| Testudinoidea | NC_026024.1 | *Heosemys depressa* |
| Testudinoidea | NC_020668.1 | *Heosemys annandalii* |
| Testudinoidea | NC_031432.1 | *Mauremys leprosa* |
| Testudinoidea | NC_016685.1 | *Mauremys sinensis* |
| Testudinoidea | NC_029369.1 | *Mauremys nigricans* |
| Testudinoidea | NC_029183.1 | *Mauremys rivulata* |
| Testudinoidea | NC_017875.1 | *Mauremys annamensis* |
| Testudinoidea | NC_016951.1 | *Mauremys japonica* |
| Testudinoidea | NC_015101.1 | *Mauremys megalocephala* |
| Testudinoidea | NC_009330.1 | *Mauremys mutica* |
| Testudinoidea | NC_020665.1 | *Notochelys platynota* |
| Testudinoidea | NC_016691.1 | *Sacalia bealei* |
| Testudinoidea | NC_011819.1 | *Sacalia quadriocellata* |
| Testudinoidea | NC_032300.1 | *Batagur trivittata* |
| Testudinoidea | NC_065016.1 | *Geoclemys hamiltonii* |
| Tetrigoidea | NC_018542.1 | *Alulatettix yunnanensis* |
| Tetrigoidea | NC_046540.1 | *Ergatettix dorsifera* |
| Tetrigoidea | NC_046542.1 | *Euparatettix variabilis* |
| Tetrigoidea | NC_046541.1 | *Euparatettix bimaculatus* |
| Tetrigoidea | NC_063118.1 | *Teredorus bashanensis* |

| | | |
|---|---|---|
| Tetrigoidea | NC_063117.1 | *Teredorus hainanensis* |
| Tetrigoidea | NC_046412.1 | *Tetrix ruyuanensis* |
| Tetrigoidea | NC_018543.1 | *Tetrix japonica* |
| Tettigoniidea | NC_028059.1 | *Cyphoderris monstrosa* |
| Tettigoniidea | NC_021397.1 | *Tarragoilus diuturnus* |
| Tettigoniidea | NC_028060.1 | *Camptonotus carolinensis* |
| Tettigoniidea | NC_033998.1 | *Homogryllacris anelytra* |
| Tettigoniidea | NC_033994.1 | *Phryganogryllacris xiai* |
| Tettigoniidea | NC_028058.1 | *Stenopelmatus fuscus* |
| Tettigoniidea | NC_035553.1 | *Pteranabropsis crenatis* |
| Tettigoniidea | NC_035552.1 | *Pteranabropsis carnarius* |
| Tettigoniidea | NC_035420.1 | *Pteranabropsis carli* |
| Tettigoniidea | NC_028063.1 | *Henicus brevimucronatus* |
| Tettigonioidea | NC_045212.1 | *Acosmetura nigrogeniculata* |
| Tettigonioidea | NC_065298.1 | *Alloxiphidiopsis emarginata* |
| Tettigonioidea | NC_033981.1 | *Decma fissa* |
| Tettigonioidea | NC_033853.1 | *Pseudocosmetura anjiensis* |
| Tettigonioidea | NC_033982.1 | *Pseudokuzicus pieli* |
| Tettigonioidea | NC_048466.1 | *Shoveliteratura triangula* |
| Tettigonioidea | NC_039981.1 | *Xiphidiopsis gurneyi* |
| Tettigonioidea | NC_040974.1 | *Xizicus maculatus* |
| Tettigonioidea | NC_018765.1 | *Xizicus fascipes* |
| Tettigonioidea | NC_065467.1 | *Anelytra multicurvata* |

| | | |
|---|---|---|
| Tettigonioidea | NC_065466.1 | *Anelytra obtusa* |
| Tettigonioidea | NC_042666.1 | *Isophya major* |
| Tettigonioidea | NC_042665.1 | *Poecilimon luschani* |
| Tettigonioidea | NC_011813.1 | *Deracantha onos* |
| Tettigonioidea | NC_033984.1 | *Zichya baranovi* |
| Tettigonioidea | NC_033987.1 | *Conanalus pieli* |
| Tettigonioidea | NC_045065.1 | *Conocephalus maculatus* |
| Tettigonioidea | NC_033988.1 | *Conocephalus melaenus* |
| Tettigonioidea | NC_053383.1 | *Euconocephalus nasutus* |
| Tettigonioidea | NC_033992.1 | *Pseudorhynchus acuminatus* |
| Tettigonioidea | NC_033990.1 | *Pseudorhynchus crassiceps* |
| Tettigonioidea | NC_033991.1 | *Ruspolia lineosa* |
| Tettigonioidea | NC_009876.1 | *Ruspolia dubia* |
| Tettigonioidea | NC_031652.1 | *Ducetia japonica* |
| Tettigonioidea | NC_033995.1 | *Kuwayamaea chinensis* |
| Tettigonioidea | NC_033993.1 | *Holochlora fruhstorferi* |
| Tettigonioidea | NC_046548.1 | *Ruidocollaris convexipennis* |
| Tettigonioidea | NC_028160.1 | *Ruidocollaris obscura* |
| Tettigonioidea | NC_033996.1 | *Lipotactes tripyrga* |
| Tettigonioidea | NC_033999.1 | *Hexacentrus unicolor* |
| Tettigonioidea | NC_033983.1 | *Hexacentrus japonicus* |
| Tettigonioidea | NC_021380.1 | *Mecopoda elongata* |
| Tettigonioidea | NC_021379.1 | *Mecopoda niponensis* |

| | | |
|---|---|---|
| Tettigonioidea | NC_034994.1 | *Sinochlora szechwanensis* |
| Tettigonioidea | NC_034757.1 | *Phaneroptera nigroantennata* |
| Tettigonioidea | NC_033997.1 | *Phyllomimus sinicus* |
| Tettigonioidea | NC_028158.1 | *Phyllomimus detersus* |
| Tettigonioidea | NC_034773.1 | *Pseudophyllus titan* |
| Tettigonioidea | NC_009967.1 | *Anabrus simplex* |
| Tettigonioidea | NC_046894.1 | *Anterastes babadaghi* |
| Tettigonioidea | NC_011200.1 | *Gampsocleis gratiosa* |
| Tettigonioidea | NC_033986.1 | *Metrioptera bonneti* |
| Thraupidae | NC_025606.1 | *Chlorophanes spiza* |
| Thraupidae | NC_062466.1 | *Diglossa brunneiventris* |
| Thraupidae | NC_039770.1 | *Geospiza magnirostris* |
| Thraupidae | NC_051469.1 | *Nesospiza acunhae* |
| Thraupidae | NC_053645.1 | *Rhodinocichla rosea* |
| Thraupidae | NC_037153.1 | *Sicalis olivascens* |
| Thraupidae | NC_051465.1 | *Sporophila hypoxantha* |
| Thraupidae | NC_035673.1 | *Sporophila maximiliani* |
| Thraupidae | NC_045366.1 | *Saltator similis* |
| Thraupidae | NC_025596.1 | *Thraupis episcopus* |
| Thripidae | NC_037839.1 | *Dendrothrips minowai* |
| Thripidae | NC_050743.1 | *Pseudodendrothrips mori* |
| Thripidae | NC_035510.1 | *Anaphothrips obscurus* |
| Thripidae | NC_018370.1 | *Frankliniella occidentalis* |

| | | |
|---|---|---|
| Thripidae | NC_025241.1 | *Scirtothrips dorsalis* |
| Thripidae | NC_058008.1 | *Thrips hawaiiensis* |
| Thripidae | NC_039437.1 | *Thrips palmi* |
| Tinamiformes | NC_052825.1 | *Crypturellus cinnamomeus* |
| Tinamiformes | NC_052773.1 | *Crypturellus soui* |
| Tinamiformes | NC_052774.1 | *Crypturellus undulatus* |
| Tinamiformes | NC_002772.2 | *Eudromia elegans* |
| Tinamiformes | NC_052818.1 | *Nothocercus julius* |
| Tinamiformes | NC_052819.1 | *Nothocercus nigrocapillus* |
| Tinamiformes | NC_052826.1 | *Nothoprocta perdicaria* |
| Tinamiformes | NC_052820.1 | *Nothoprocta ornata* |
| Tinamiformes | NC_027260.1 | *Tinamus guttatus* |
| Tinamiformes | NC_002781.3 | *Tinamus major* |
| Tingoidea | NC_037146.1 | *Agramma hupehanum* |
| Tingoidea | NC_044420.1 | *Corythucha marmorata* |
| Tingoidea | NC_022922.1 | *Corythucha ciliata* |
| Tingoidea | NC_037833.1 | *Cysteochila chiniana* |
| Tingoidea | NC_037834.1 | *Dictyla platyoma* |
| Tingoidea | NC_037835.1 | *Metasalis populi* |
| Tingoidea | NC_046031.1 | *Neoplerochila paliatseasi* |
| Tingoidea | NC_037148.1 | *Phatnoma laciniatum* |
| Tingoidea | NC_025299.1 | *Pseudacysta perseae* |
| Tingoidea | NC_037836.1 | *Tingis cardui* |

| | | |
|---|---|---|
| Tipuloidea | NC_030519.1 | *Symplecta hybrida* |
| Tipuloidea | NC_057072.1 | *Conosia irrorata* |
| Tipuloidea | NC_057085.1 | *Epiphragma mediale* |
| Tipuloidea | NC_044484.1 | *Limonia phragmitidis* |
| Tipuloidea | NC_053795.1 | *Tanyptera hebeiensis* |
| Tortricidae | NC_037395.1 | *Choristoneura fumiferana* |
| Tortricidae | NC_019996.1 | *Choristoneura longicellana* |
| Tortricidae | NC_037396.1 | *Choristoneura murinana* |
| Tortricidae | NC_037397.1 | *Choristoneura rosaceana* |
| Tortricidae | NC_039422.1 | *Choristoneura pinus pinus* |
| Tortricidae | NC_037393.1 | *Choristoneura occidentalis* |
| Tortricidae | NC_039421.1 | *Choristoneura conflictana* |
| Tortricidae | NC_037394.1 | *Choristoneura biennis* |
| Tortricidae | NC_021396.1 | *Adoxophyes orana* |
| Tortricidae | NC_008141.1 | *Adoxophyes honmai* |
| Tortricidae | NC_060406.1 | *Cochylidia moguntiana* |
| Tortricidae | NC_060405.1 | *Cochylimorpha cultana* |
| Tortricidae | NC_018754.1 | *Acleris fimbriana* |
| Tortricidae | NC_054267.1 | *Cerace xanthocosma* |
| Toxicofera | NC_012829.1 | *Basiliscus vittatus* |
| Toxicofera | NC_012831.1 | *Gambelia wislizenii* |
| Toxicofera | NC_044125.1 | *Anolis punctatus* |
| Toxicofera | NC_028031.1 | *Amblyrhynchus cristatus* |

| | | |
|---|---|---|
| Toxicofera | NC_028030.1 | *Conolophus subcristatus* |
| Toxicofera | NC_027089.1 | *Cyclura pinguis* |
| Toxicofera | NC_044899.1 | *Iguana delicatissima* |
| Toxicofera | NC_057244.1 | *Liolaemus parthenos* |
| Toxicofera | NC_057243.1 | *Liolaemus millcayac* |
| Toxicofera | NC_057242.1 | *Liolaemus darwinii* |
| Toxicofera | NC_012836.1 | *Chalarodon madagascariensis* |
| Toxicofera | NC_012827.1 | *Oplurus grandidieri* |
| Toxicofera | NC_012839.1 | *Polychrus marmoratus* |
| Toxicofera | NC_012834.1 | *Leiocephalus personatus* |
| Toxicofera | NC_014174.1 | *Brookesia decaryi* |
| Toxicofera | NC_008777.1 | *Furcifer oustaleti* |
| Toxicofera | NC_014176.1 | *Trioceros melleri* |
| Toxicofera | NC_041001.1 | *Holbrookia lacerata* |
| Toxicofera | NC_036492.1 | *Phrynosoma blainvillii* |
| Toxicofera | NC_005960.1 | *Sceloporus occidentalis* |
| Toxicofera | NC_026308.1 | *Urosaurus nigricaudus* |
| Toxicofera | NC_027261.1 | *Uta stansburiana* |
| Trachypithecus | NC_056330.1 | *Trachypithecus laotum* |
| Trachypithecus | NC_056326.1 | *Trachypithecus phayrei* |
| Trachypithecus | NC_056325.1 | *Trachypithecus mauritius* |
| Trachypithecus | NC_056324.1 | *Trachypithecus geei* |
| Trachypithecus | NC_034795.1 | *Trachypithecus poliocephalus* |

| | | |
|---|---|---|
| Trachypithecus | NC_024529.1 | *Trachypithecus pileatus* |
| Trachypithecus | NC_023971.1 | *Trachypithecus cristatus* |
| Trachypithecus | NC_023970.1 | *Trachypithecus francoisi* |
| Trachypithecus | NC_019583.1 | *Trachypithecus johnii* |
| Trachypithecus | NC_019582.1 | *Trachypithecus vetulus* |
| Trachypithecus | NC_019581.1 | *Trachypithecus shortridgei* |
| Trachypithecus | NC_019580.1 | *Trachypithecus germaini* |
| Trachypithecus | NC_019579.1 | *Trachypithecus hatinhensis* |
| Trachypithecus | NC_006900.1 | *Trachypithecus obscurus* |
| Turdus | NC_059847.1 | *Turdus dissimilis* |
| Turdus | NC_057250.1 | *Turdus ruficollis* |
| Turdus | NC_052843.1 | *Turdus abyssinicus* |
| Turdus | NC_046948.1 | *Turdus cardis* |
| Turdus | NC_041095.1 | *Turdus kessleri* |
| Turdus | NC_028273.1 | *Turdus eunomus* |
| Turdus | NC_028188.1 | *Turdus merula* |
| Turdus | NC_028179.1 | *Turdus rufiventris* |
| Turdus | NC_029147.1 | *Turdus philomelos* |
| Turdus | NC_024872.1 | *Turdus migratorius* |
| Tylopoda | NC_009849.1 | *Camelus dromedarius* |
| Tylopoda | NC_009629.2 | *Camelus ferus* |
| Tylopoda | NC_009628.2 | *Camelus bactrianus* |
| Tylopoda | NC_011822.1 | *Lama guanicoe* |

| | | |
|---|---|---|
| Typhlatya | NC_036335.1 | *Typhlatya miravetensis* |
| Typhlatya | NC_035410.1 | *Typhlatya arfeae* |
| Typhlatya | NC_035408.1 | *Typhlatya dzilamensis* |
| Typhlatya | NC_035407.1 | *Typhlatya consobrina* |
| Typhlatya | NC_035405.1 | *Typhlatya monae* |
| Typhlatya | NC_035403.1 | *Typhlatya mitchelli* |
| Typhlatya | NC_035402.1 | *Typhlatya galapagensis* |
| Typhlatya | NC_035401.1 | *Typhlatya iliffei* |
| Typhlatya | NC_035400.1 | *Typhlatya pearsei* |
| Typhlatya | NC_035399.1 | *Typhlatya taina* |
| Typhlatya | NC_035409.1 | *Typhlatya garciai* |
| Tyrannidae | NC_007975.1 | *Cnemotriccus fuscatus* |
| Tyrannidae | NC_024682.1 | *Mionectes oleagineus* |
| Tyrannidae | NC_053075.1 | *Neopipo cinnamomea* |
| Tyrannidae | NC_053085.1 | *Onychorhynchus coronatus* |
| Tyrannidae | NC_051035.1 | *Pachyramphus minor* |
| Tyrannidae | NC_053107.1 | *Sapayoa aenigma* |
| Tyrannidae | NC_051005.1 | *Tachuris rubrigastra* |
| Tyrannidae | NC_051025.1 | *Tyrannus savana* |
| Unipeltata | NC_063586.1 | *Alima pacifica* |
| Unipeltata | NC_006916.1 | *Harpiosquilla harpax* |
| Unipeltata | NC_053568.1 | *Lophosquilla costata* |
| Unipeltata | NC_014342.1 | *Oratosquilla oratoria* |

| | | |
|---|---|---|
| Unipeltata | NC_063585.1 | *Squilla biformis* |
| Unipeltata | NC_006081.1 | *Squilla mantis* |
| Unipeltata | NC_007444.1 | *Squilla empusa* |
| Unipeltata | NC_027178.1 | *Squilloides leptosquilla* |
| Unipeltata | NC_007443.1 | *Lysiosquillina maculata* |
| Unipeltata | NC_063583.1 | *Hemisquilla californiensis* |
| Ursidae | NC_009492.1 | *Ailuropoda melanoleuca* |
| Ursidae | NC_011116.1 | *Arctodus simus* |
| Ursidae | NC_030174.1 | *Arctotherium sp.* |
| Ursidae | NC_009968.1 | *Helarctos malayanus* |
| Ursidae | NC_009970.1 | *Melursus ursinus* |
| Ursidae | NC_009969.1 | *Tremarctos ornatus* |
| Ursidae | NC_008753.1 | *Ursus thibetanus mupinensis* |
| Ursidae | NC_003428.1 | *Ursus maritimus* |
| Ursidae | NC_003427.1 | *Ursus arctos* |
| Ursidae | NC_011118.1 | *Ursus thibetanus thibetanus* |
| Ursidae | NC_011117.1 | *Ursus thibetanus ussuricus* |
| Ursidae | NC_011112.1 | *Ursus spelaeus* |
| Ursidae | NC_009971.1 | *Ursus thibetanus* |
| Ursidae | NC_009331.1 | *Ursus thibetanus formosanus* |
| Ursidae | NC_003426.1 | *Ursus americanus* |
| Vermilingua | NC_028564.1 | *Cyclopes didactylus* |
| Vermilingua | NC_028572.1 | *Myrmecophaga tridactyla* |

| | | |
|---|---|---|
| Vermilingua | NC_028574.1 | *Tamandua mexicana* |
| Vermilingua | NC_004032.1 | *Tamandua tetradactyla* |
| Vespoidea | NC_048883.1 | *Antodynerus aff. limbatus* |
| Vespoidea | NC_039949.1 | *Orancistrocerus aterrimus* |
| Viperidae | NC_009768.1 | *Agkistrodon piscivorus* |
| Viperidae | NC_035638.1 | *Agkistrodon contortrix* |
| Viperidae | NC_039649.1 | *Bothrops diporus* |
| Viperidae | NC_030760.1 | *Bothrops jararaca* |
| Viperidae | NC_039648.1 | *Bothrops pubescens* |
| Viperidae | NC_041524.1 | *Crotalus adamanteus* |
| Viperidae | NC_010223.1 | *Deinagkistrodon acutus* |
| Viperidae | NC_064056.1 | *Gloydius rubromaculatus* |
| Viperidae | NC_036234.1 | *Gloydius strauchi* |
| Viperidae | NC_029424.1 | *Gloydius shedaoensis* |
| Viperidae | NC_026553.1 | *Gloydius ussuriensis* |
| Viperidae | NC_025666.1 | *Gloydius saxatilis* |
| Viperidae | NC_025560.1 | *Gloydius intermedius* |
| Viperidae | NC_011390.1 | *Gloydius blomhoffi brevicaudus* |
| Viperidae | NC_007397.1 | *Ovophis okinavensis* |
| Viperidae | NC_029494.1 | *Trimeresurus sichuanensis* |
| Viperidae | NC_022820.1 | *Trimeresurus albolabris* |
| Viperidae | NC_030781.1 | *Azemiops feae* |
| Viperidae | NC_013479.1 | *Causus defilippi* |

| | | |
|---|---|---|
| Viperidae | NC_011391.1 | *Daboia russellii* |
| Viperidae | NC_060592.1 | *Echis coloratus* |
| Viperidae | NC_044966.1 | *Macrovipera schweizeri* |
| Xanthoidea | NC_049030.1 | *Carpilius maculatus* |
| Xanthoidea | NC_006891.1 | *Pseudocarcinus gigas* |
| Xanthoidea | NC_039110.1 | *Epixanthus frontalis* |
| Xanthoidea | NC_037201.1 | *Atergatis floridus* |
| Xanthoidea | NC_037172.1 | *Atergatis integerrimus* |
| Xanthoidea | NC_042208.1 | *Etisus anaglyptus* |
| Xanthoidea | NC_029726.1 | *Leptodius sanguineus* |
| Xanthoidea | NC_057473.1 | *Macromedaeus distinguendus* |
| Xenopus | NC_006839.1 | *Xenopus tropicalis* |
| Xenopus | NC_001573.1 | *Xenopus laevis* |
| Xenopus | NC_044888.1 | *Xenopus fischbergi* |
| Xenopus | NC_044887.1 | *Xenopus muelleri* |
| Xenopus | NC_044886.1 | *Xenopus clivii* |
| Xenopus | NC_044885.1 | *Xenopus longipes* |
| Xenopus | NC_044884.1 | *Xenopus eysoole* |
| Xenopus | NC_044883.1 | *Xenopus kobeli* |
| Xenopus | NC_044882.1 | *Xenopus ruwenzoriensis* |
| Xenopus | NC_044881.1 | *Xenopus vestitus* |
| Xenopus | NC_044880.1 | *Xenopus lenduensis* |
| Xenopus | NC_044879.1 | *Xenopus itombwensis* |

| | | |
|---|---|---|
| Xenopus | NC_044878.1 | *Xenopus andrei* |
| Xenopus | NC_044877.1 | *Xenopus boumbaensis* |
| Xenopus | NC_044876.1 | *Xenopus amieti* |
| Xenopus | NC_044875.1 | *Xenopus wittei* |
| Xenopus | NC_044874.1 | *Xenopus allofraseri* |
| Xenopus | NC_044873.1 | *Xenopus pygmaeus* |
| Xenopus | NC_044872.1 | *Xenopus parafraseri* |
| Xenopus | NC_044871.1 | *Xenopus gilli* |
| Xenopus | NC_044870.1 | *Xenopus poweri* |
| Xenopus | NC_044869.1 | *Xenopus petersii* |
| Xenopus | NC_044868.1 | *Xenopus largeni* |
| Xenopus | NC_044867.1 | *Xenopus epitropicalis* |
| Xenopus | NC_044866.1 | *Xenopus mellotropicalis* |
| Xenopus | NC_044865.1 | *Xenopus calcaratus* |
| Xenopus | NC_018775.1 | *Xenopus victorianus* |
| Xenopus | NC_018776.1 | *Xenopus borealis* |
| Xerinae | NC_031210.1 | *Callospermophilus lateralis* |
| Xerinae | NC_026706.1 | *Cynomys ludovicianus* |
| Xerinae | NC_026705.1 | *Cynomys leucurus* |
| Xerinae | NC_027278.1 | *Ictidomys tridecemlineatus* |
| Xerinae | NC_048490.1 | *Marmota vancouverensis* |
| Xerinae | NC_042243.1 | *Marmota flaviventris* |
| Xerinae | NC_018367.1 | *Marmota himalayana* |

| | | |
|---|---|---|
| Xerinae | NC_059784.1 | *Spermophilus citellus* |
| Xerinae | NC_027283.1 | *Spermophilus dauricus* |
| Xerinae | NC_032375.1 | *Tamias striatus* |
| Xerinae | NC_032373.1 | *Tamias dorsalis* |
| Xerinae | NC_032372.1 | *Tamias canipes* |
| Xerinae | NC_032371.1 | *Tamias rufus* |
| Xerinae | NC_032370.1 | *Tamias quadrivittatus* |
| Xerinae | NC_032376.1 | *Tamias umbrinus* |
| Xerinae | NC_025277.1 | *Tamias sibiricus* |
| Xerinae | NC_059785.1 | *Urocitellus parryii* |
| Xerinae | NC_031209.1 | *Urocitellus richardsonii* |
| Yponomeutoidea | NC_018547.1 | *Leucoptera malifoliella* |
| Yponomeutoidea | NC_037944.1 | *Lyonetia clerkella* |
| Yponomeutoidea | NC_064061.1 | *Acrolepiopsis assectella* |
| Yponomeutoidea | NC_039687.1 | *Plutella australiana* |
| Yponomeutoidea | NC_025322.1 | *Plutella xylostella* |
| Yponomeutoidea | NC_064059.1 | *Plutella armoraciae* |
| Yponomeutoidea | NC_064060.1 | *Plutella porrectella* |
| Yponomeutoidea | NC_025948.1 | *Prays oleae P38* |
| Zygaenidae | NC_046467.1 | *Amesia sanguiflua* |
| Zygaenidae | NC_038208.1 | *Eterusia aedea* |
| Zygaenidae | NC_039447.1 | *Histia rhodope* |
| Zygaenidae | NC_037909.1 | *Pidorus atratus* |

| | | |
|---|---|---|
| Zygaenidae | NC_025761.1 | *Rhodopsona rubiginosa* |
| Zygentoma | NC_005437.1 | *Tricholepidion gertschi* |
| Zygentoma | NC_046478.1 | *Ctenolepisma villosa* |
| Zygentoma | NC_047445.1 | *Lepisma saccharina* |
| Zygentoma | NC_053634.1 | *Neoasterolepisma foreli* |
| Zygentoma | NC_006080.1 | *Thermobia domestica* |

Table 2: Table of every taxon, their included species and respective NCBI accession number in `euka`'s reference database.
